# Distance decay 2.0 – a global synthesis of taxonomic and functional turnover in ecological communities

**DOI:** 10.1101/2021.03.17.435827

**Authors:** Caio Graco-Roza, Sonja Aarnio, Nerea Abrego, Alicia T. R. Acosta, Janne Alahuhta, Jan Altman, Claudia Angiolini, Jukka Aroviita, Fabio Attorre, Lars Baastrup-Spohr, José Juan Barrera-Alba, Jonathan Belmaker, Idoia Biurrun, Gianmaria Bonari, Helge Bruelheide, Sabina Burrascano, Marta Carboni, Pedro Cardoso, José Carlos Carvalho, Giuseppe Castaldelli, Morten Christensen, Gilsineia Correa, Iwona Dembicz, Jürgen Dengler, Jiri Dolezal, Patricia Domingos, Tibor Erös, Carlos E. L. Ferreira, Goffredo Filibeck, Sergio R. Floeter, Alan Friedlander, Johanna Gammal, Anna Gavioli, Martin M. Gossner, Itai Granot, Riccardo Guarino, Camilla Gustafsson, Brian Hayden, Siwen He, Jacob Heilmann-Clausen, Jani Heino, John T. Hunter, Vera Lucia de Moraes Huszar, Monika Janišová, Jenny Jyrkänkallio-Mikkola, Kimmo Kahilainen, Julia Kemppinen, Łukasz Kozub, Carla Kruk, Michel Kulbiki, Anna Kuzemko, Peter Christian le Roux, Aleksi Lehikoinen, Domênica Teixeira de Lima, Angel Lopes-Urrutia, Balázs A. Lukács, Miska Luoto, Stefano Mammola, Marcelo Manzi Marinho, Luciana da Silva Menezes, Marco Milardi, Marcela Miranda, Gleyci Aparecida Oliveira Moser, Joerg Mueller, Pekka Niittynen, Alf Norkko, Arkadiusz Nowak, Jean Ometto, Otso Ovaskainen, Gerhard E. Overbeck, Felipe Siqueira Pacheco, Virpi Pajunen, Salza Palpurina, Félix Picazo, Juan Antonio Campos Prieto, Ivan F. Rodil, Francesco Maria Sabatini, Shira Salingré, Michele de Sanctis, Angel M. Segura, Lucia Helena Sampaio da Silva, Zora Dajic Stevanovic, Grzegorz Swacha, Anette Teittinen, Kimmo T. Tolonen, Ioannis Tsiripidis, Leena Virta, Beixin Wang, Jianjun Wang, Wolfgang Weisser, Yuan Xu, Janne Soininen

## Abstract

Understanding the variation in community composition and species abundances, i.e., β-diversity, is at the heart of community ecology. A common approach to examine β-diversity is to evaluate directional turnover in community composition by measuring the decay in the similarity among pairs of communities along spatial or environmental distances. We provide the first global synthesis of taxonomic and functional distance decay along spatial and environmental distance by analysing 149 datasets comprising different types of organisms and environments. We modelled an exponential distance decay for each dataset using generalized linear models and extracted r^2^ and slope to analyse the strength and the rate of the decay. We studied whether taxonomic or functional similarity has stronger decay across the spatial and environmental distances. We also unveiled the factors driving the rate of decay across the datasets, including latitude, spatial extent, realm, and organismal features. Taxonomic distance decay was stronger along spatial and environmental distances compared with functional distance decay. The rate of taxonomic spatial distance decay was the fastest in the datasets from mid-latitudes while the rate of functional decay increased with latitude. Overall, datasets covering larger spatial extents showed a lower rate of decay along spatial distances but a higher rate of decay along environmental distances. Marine ecosystems had the slowest rate of decay. This synthesis is an important step towards a more holistic understanding of patterns and drivers of taxonomic and functional β-diversity.

## Introduction

Biodiversity on Earth is shrinking^1^. Understanding its distribution is therefore paramount to inform conservation efforts, and to evaluate the links between biodiversity, ecosystem functioning, ecosystem services and human well-being^2,3^. The variation in the occurrence and abundance of species in space and time, i.e., β-diversity, is at the heart of community ecology and biogeography as it provides a direct link between local (α) and regional (γ) diversity^4,5^. Moreover, β-diversity has become an essential currency in spatial^6,7^ and temporal^8^ comparisons of biodiversity patterns and their underlying drivers. β-diversity is also informative in the context of biodiversity conservation and practical management decisions in rapidly changing environments^9,10^.

A common approach to examine spatial β-diversity is to consider directional turnover in community composition with distance, i.e., distance decay ^4,11^. The similarity among the pairs of biological communities typically decreases (“decays”) with increasing spatial or environmental distance ^11,12^. This pattern stems mainly from dispersal limitation (related to physical barriers and dispersal ability^13^) and species-specific responses to spatially structured environmental variation (related to environmental filters and evolutionary processes^14^) and is well-documented in observational^15–17^ and theoretical studies^18^ as well as meta-analyses^19^. Such studies offer interesting insights into the patterns and drivers of spatial taxonomic β-diversity and often provide information about the effects of environmental changes on ecosystem processes and associated functionality. Even if the patterns and drivers of taxonomic β-diversity are relatively well-documented in the biogeographic literature, it is much less understood whether the same patterns occur for functional β-diversity^20–22^. Therefore, functional biogeography emerges as a field to solve questions related to the distribution of forms and functions of individuals, populations, communities, ecosystems, and biomes across spatial scales^23^.

Understanding functional diversity relies on trait-based approaches, which are built on the idea that the environment selects species based on their ecological requirements, and that functional traits capture these requirements better than species identity^24^. Thus, a trait-based approach should reflect the functional response of biotic communities to environmental gradients better than an approach based on species’ taxonomic identities only, and better predict how biotic communities respond to environmental changes^25^. Even if functional diversity has been investigated widely at the α-diversity level^26,27^, our understanding of functional β-diversity is much more limited and fragmented^28–32^. Comparing the patterns of functional and taxonomic β-diversity across different biotic groups, ecosystems and geographic contexts has the potential to greatly contribute to a better mechanistic understanding of the drivers behind the spatial variation in ecosystem functionality and shed further light on how environmental change may affect ecological communities.

Niche filtering along environmental gradients induces coupling of taxonomic and functional diversity patterns because dominant functional strategies dictate along the environmental gradient^33,34^. However, high taxonomic β-diversity does not necessarily mean high functional β-diversity^25,35^ (Fig. 1a), and the gain or loss of species does not inform about variations in functional β-diversity whenever trait redundancy is high^36^. For example, taxonomic homogenization does not lead to functional homogenization if the newly introduced species in the assemblages are functionally similar to each other^30,37,38^. The most pressing question is whether functional features explain more of the distance decay along environmental gradients than species identities, as suggested by some earlier studies^39–43^.

**Figure 1.**
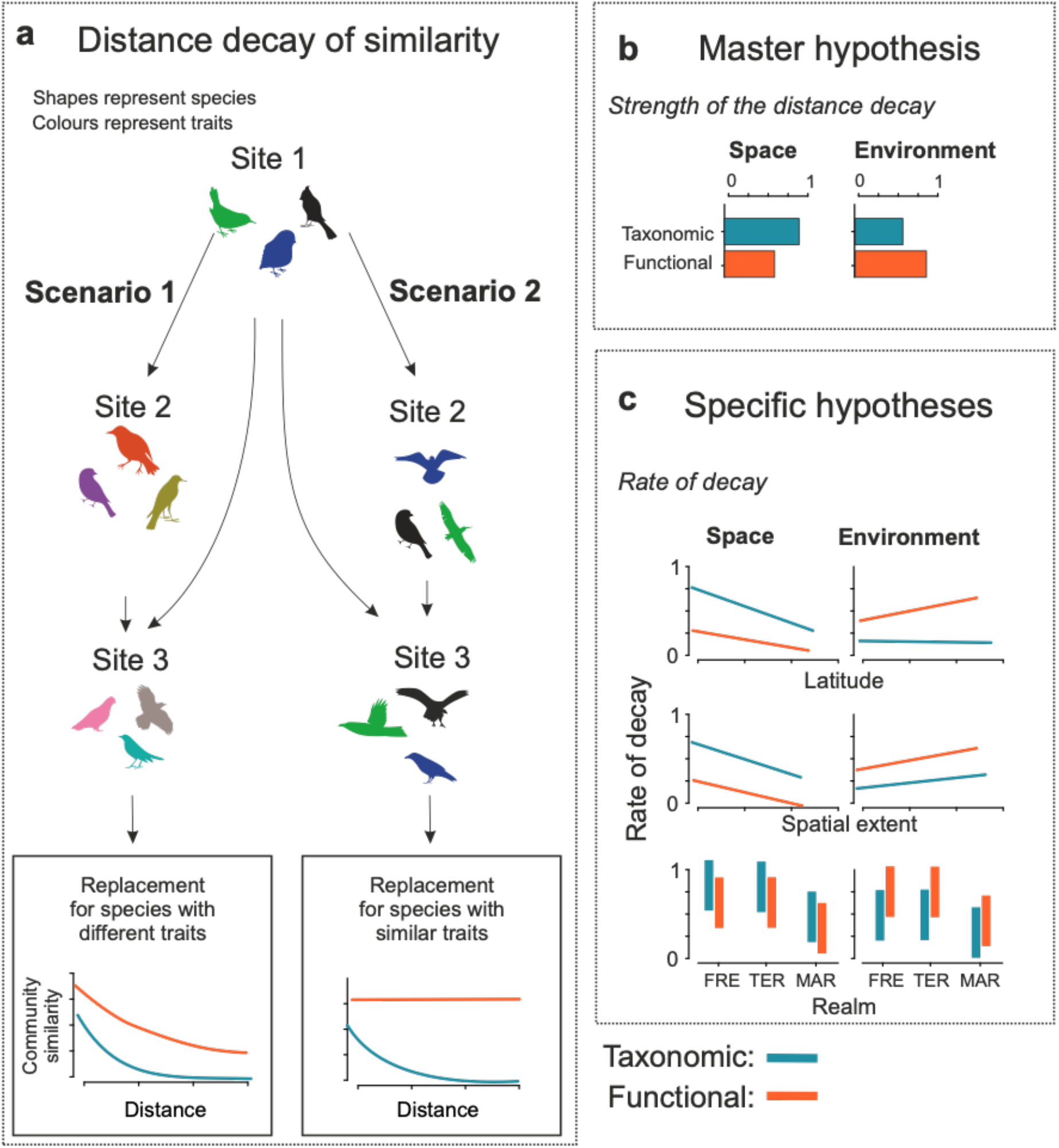
(a) Taxonomic and functional distance decay. Two scenarios of distance decay of taxonomic and functional similarities along spatial and environmental distances. In scenario 1 (for simplicity, we consider here replacement only), the replacement occurs among species that have different traits (i.e., colours), which leads to both taxonomic and functional distance decay. In scenario 2, the replacement occurs among species that have similar traits, which leads to zero functional distance decay measured by the slope. (b) Master hypothesis: spatial distance decay is stronger for taxonomic similarities than for functional similarities, while environmental distance decay is stronger for functional similarities. (c) Specific hypotheses (higher values indicate steeper slopes) across datasets: <u>Latitude</u>: spatial distance decay is flatter in the datasets from higher latitude and more notably for taxonomic similarities than for functional similarities. Environmental distance decay is steeper in datasets from higher latitude for functional similarities, while it does not vary notably with latitude for taxonomic similarities. <u>Spatial extent:</u> Both taxonomic and functional spatial distance decay are flatter in the datasets covering larger spatial extent, while environmental distance decay is steeper in datasets covering larger extent. <u>Realm:</u> Marine ecosystems show flatter spatial and environmental distance decay than terrestrial and freshwater systems. FRE= freshwater systems, TER = terrestrial systems, MAR = marine systems.

### Hypotheses

Since the emergence of the first comprehensive distance decay meta-analysis^19^, our understanding of community turnover along spatial and environmental gradients has increased notably. Here, based on existing ecological literature and theory, and as an initial step towards synthesising knowledge, we tested four hypotheses concerning the differences between taxonomic and functional distance decay along the spatial and environmental distances. The master hypothesis is that the distance decay along spatial gradients is stronger for taxonomic similarity than for functional similarity (**H_1a_**). This is because spatial factors relate with taxonomic more than functional composition as a result of dispersal processes, dispersal history and speciation^42^. Such a hypothesis should be valid when functional traits do not comprise dispersal related traits. In contrast, distance decay along environmental gradients is stronger for functional similarity than for taxonomic similarity because functional composition should respond more strongly to environmental variation^27,39,40,42^ (**H_1b_**) (Fig. 1b).

#### Latitudinal gradients

We also generalize the effects of major geographic and environmental factors in the three hypotheses, which are tested across the datasets. For example, latitudinal effect has been recognized as a relevant factor in meta-analyses^44^ and case studies^45,46^, and these studies suggest that β-diversity should decrease with increasing latitude (Fig. 1c). This is indicated by the faster latitudinal decline in γ- diversity than in α-diversity^47,48^, and the slopes of the species-area relationships (*proxy* for turnover) decrease with latitude^49^. Moreover, Rapoport’s rule^50^ postulates that species range sizes are larger at high latitudes leading to lower β-diversity. Therefore, we hypothesize that the rate of taxonomic distance decay along spatial gradients is generally slower in the datasets that originate from higher latitudes (**H_2a_**). In contrast, functional distance decay may show faster rates in the datasets from higher latitudes. This is because the high diversity of tropical areas stems mainly from niche overlap^51^, which increases the functional redundancy within communities and reduces the functional turnover^52^. Regarding the environmental gradients, large-scale environmental heterogeneity tends to increase towards poles^19,53,54^, leading to a faster rate of functional distance decay along environmental gradients at higher latitudes (**H_2b_**). An alternative hypothesis is that extreme climatic conditions at high latitudes decrease functional diversity because abiotic filtering limits the number of possible ecological strategies found in a biotic community^55,56^, resulting in relatively slow rate of functional distance decay.

#### Spatial extent

Distance decay is also likely to be affected by the spatial extent of a given study^57^. It has been shown that distance decay has a power-law shape at spatial extents that do not exceed regional species pools and exponential shape when extent encompasses multiple species pools^12^. This suggests that the slope of the relationship becomes flatter with increasing spatial extent^11,19^, mainly because regional species diversity is limited with a certain upper boundary^58^. Furthermore, environmental heterogeneity affects the diversity of species^59^ and functional traits at regional level^60,61^, but such effects are likely to be scale-dependent^62–64^. To summarize, we hypothesize that the rate of distance decay along spatial gradients is generally slower in the datasets covering larger spatial extent (**H_3a_**). In contrast, we hypothesize that the rate of distance decay along environmental gradients is generally faster when spatial extent is larger, especially for functional similarities, which are considered more sensitive to environmental variation (**H_3b_**).

#### Realms

We also expect that the patterns of distance decay vary among the realms. In general, marine ecosystems are environmentally more homogeneous than terrestrial or freshwater ecosystems, at least in the open ocean^65^, and typically show weaker dispersal barriers than terrestrial or freshwater ecosystems^66^. Therefore, we hypothesize that the datasets from marine ecosystems have generally slower rate of taxonomic and functional distance decay than the other ecosystems (**H_4_**).

Here, we tested these hypotheses using datasets that cover a wide range of biotic groups from unicellular diatoms to vascular plants, fungi, invertebrates, fish, birds, amphibians and mammals, and that originate from marine, terrestrial and freshwater ecosystems spanning broad latitudinal gradients (Fig. 2). To account for major biological differences in biotic groups, we also investigated if distance decay varied among different sized taxa or among taxa with different dispersal mode^67,68^. By using such a comprehensive, multi-realm and multi-taxon dataset, we will explore patterns at more general level, compared with case studies that have examined both taxonomic and functional β-diversity, but only considered a single or few biotic groups.

**Figure 2.**
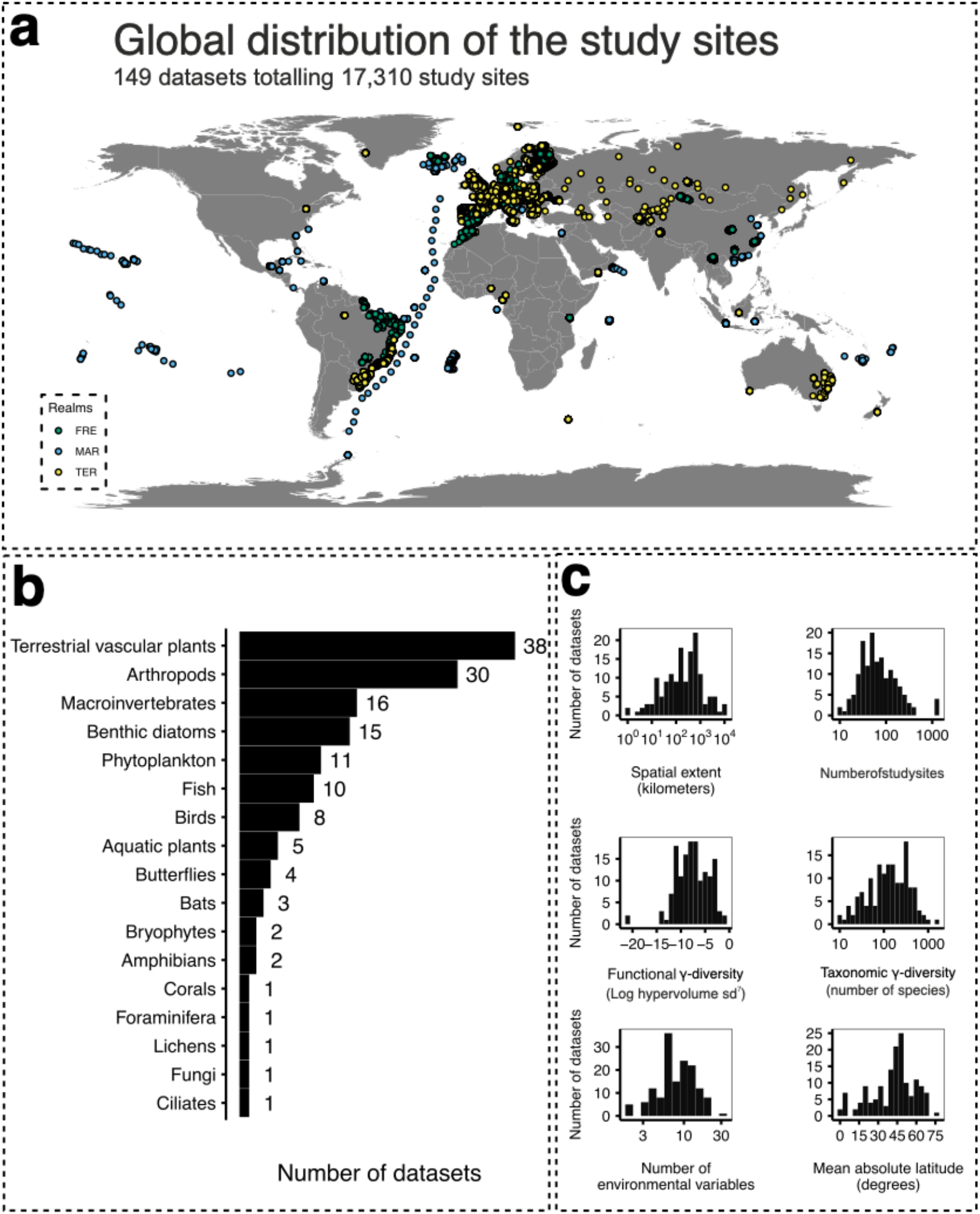
Study design highlighting (a) map of the study sites coloured according to the realms (FRE = Freshwater, TER = Terrestrial, MAR = Marine); (b) the number of data sets for major biotic groups; and (c) the distribution of the datasets with respect to spatial extent, number of study sites, functional γ-diversity (log hypervolume sd^7^), taxonomic γ-diversity (number of species), number of environmental variables, and latitude.

## Material and methods

### Data collection

We gathered our data by directly contacting data owners or using the existing data sources, such as sPlot^69^ and CESTES^70^. We included datasets that provided raw data of species abundances, functional traits, environmental variables and spatial coordinates of the study sites. A few datasets (n = 6) provided only species occurrence rather than abundance information (Appendix S1). The traits included in the datasets were chosen by data owners from a suite of traits that should respond well to environmental variation. For plant datasets compiled from the sPlot database, trait information was commonly derived from the TRY database^71^. Regarding the CESTES database, we compiled 48 datasets, specifically from: fish communities^22,72–74^, terrestrial vascular plants^75–86^, aquatic macroinvertebrates^87–89^, terrestrial arthropods^86,90–98^, birds^83,90, 99–102^, bats^102,103^, bryophytes^85^, butterflies^98,104^, corals^105^, and foraminifera^106^. We only included datasets with at least ten sites, two environmental variables and three traits or trait categories. In some cases, more than one dataset representing different taxonomic groups with different responses to environment and dispersal abilities (e.g., stream macroinvertebrates and diatoms) were collected in the same study area. In total, 149 datasets representing 17 major biotic groups from terrestrial (n = 87), freshwater (n = 41) and marine (n = 21) environments were assembled amounting to over 17,000 study sites around the globe (Fig. 2). From the 149 datasets, 118 were published in peer reviewed journals (Appendix S1).Taxa were mostly identified to species or morphospecies level but, in a few cases, we used data at genus level if existing taxonomic knowledge did not allow distinguishing individual species. Finally, each dataset included (i) a sites-by-species abundances matrix, (ii) a species-by-traits table, (iii) a sites-by- spatial coordinates table, and (iv) a sites-by-environmental variables table (Fig. 3a). Detailed information about collected datasets can be found in Appendix S1.

**Figure 3.**
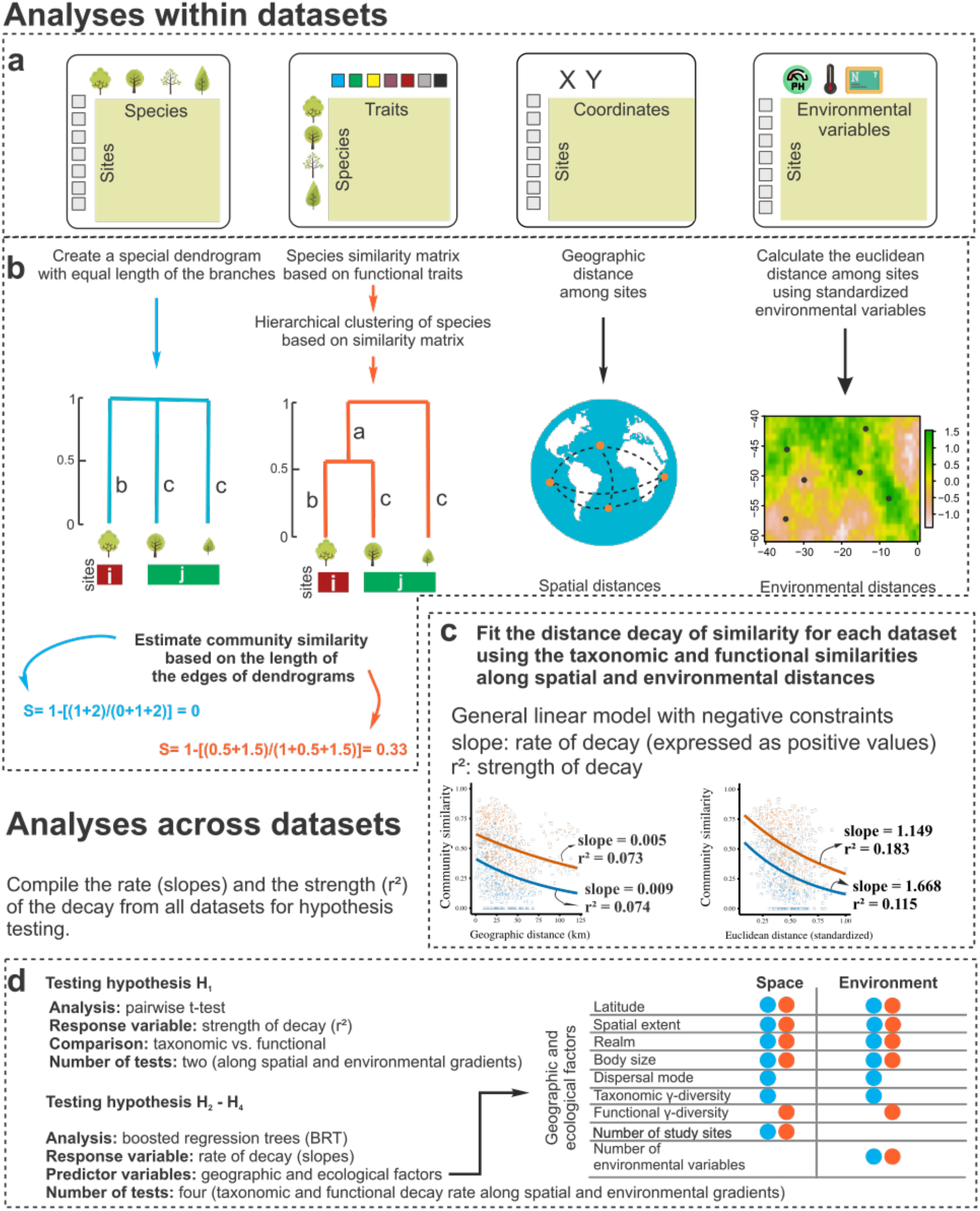
The analytical framework described step-wisely. The blocks a-c hierarchically describe the methods performed at dataset level, including the estimation of similarities and distances as well as the distance decay models of each dataset. The block d describes the tests performed after the compilation of the metrics from all datasets. The first block (a) shows the four objects used in the analyses: a species-by-traits table, a sites-by-species matrix, a sites-by-coordinates table and a sites- by-environment table. The second block (b) illustrates the calculation of taxonomic and functional similarities, and spatial and environmental distances. In the first example, only species identities are taken into account and as sites *i* and *j* do not share any species, community similarity (blue) equals zero. In the second example, sites *i* and *j* do not share any species, but as two species have same body size, community similarity (orange) is higher than zero. Similarity is estimated using the length of the edge of the dendrograms as S = 1-[(b+c)/(2a+b+c)]. The third example shows how spatial distances were calculated as the geographic distances among sites using spatial coordinates. The fourth example illustrates how sites far from each other may show similar environmental conditions and therefore small environmental distance. Environmental distances were calculated as the Euclidean distances of standardized environmental variables. The third block (c) illustrates the metrics extracted to study the distance decay across datasets. The strength (r^2^) and rate (slope) of decay were extracted from each dataset using log-binomial generalized linear models (GLM). The models were built separately for each response variable (taxonomic or functional similarity) and explanatory variables (spatial or environmental distance), totalling four r^2^ values and four slopes. Also, the data of marine fish from the Mediterranean Sea is shown as an example where the distance decay of similarity along environmental distance is stronger (higher r^2^) for functional similarity than for taxonomic similarity, irrespectively of the rate of decay (slope). The fourth block (d) describes the analyses used to test the hypotheses and which metrics were considered for each analysis. The strength (r^2^) of decay was used to test hypothesis H_1_ while the rate of decay (slope) was used to hypotheses H_2_-H_4_.

### Data curation

For each dataset, we removed the sites with less than two observed species, and the species with lower than three traits considered. Trait data included ordered, categorical and continuous traits, the latter of which were log transformed (Log10) when needed. Environmental variables were log-transformed (Log10) to approximate normality (except for e.g., temperature, pH and variables given as eigenvectors), and the environmental variables showing strong inter- correlations (pairwise rp < 0.7) were excluded from further analyses^107^. Spatial coordinates were converted to the World Geodetic System 1984 (WGS84) datum and geographic coordinate system and expressed in decimal degrees with an accuracy up to five decimals. All the data curation and further analyses were performed in the software R v.4.0.2 (ref.^108^) using the appropriate R packages. We will consistently refer to the functions used and their respective packages from here on.

### Taxonomic and functional similarities

Pairwise between-site taxonomic and functional similarities were obtained for each dataset following the tree-based approach implemented in the function beta in the package ‘BAT’ v.2.1.0 (ref.^109^). We used the tree-based approach because it provides an unequivocal comparison of taxonomic and functional similarities^110^. Community similarity (S) ranges between zero and one and is commonly calculated for the pairs of communities as the sum of the unique features of each community over the sum of the shared features between communities and the unique features of each community. In the tree-based approach, these features are edges, which may have different lengths and be shared by different species that may be present in different communities^110^. Taxonomic and functional similarities were calculated for species occurrences and abundances based on a Podani family of Sørensen-based indices^111^. Here, we estimated S between communities *j* and *k* as 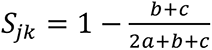 (1), where *a* is the sum of the length of the edges shared between the communities *j* and *k, b* is the sum of the length of the edges unique to the community *j,* and *c* is the sum of the length of the edges unique to the community *k*.

When estimating taxonomic similarities, each species is a unique entity that share no edges with others and, therefore, all the edges of the tree have same length (Fig. 3b). Thus, the sum of the length of the edges equals the sum of the number of the observed species. For functional similarities, the length of the edge shared between two species depends on how similar species are with respect to their traits. To estimate the length of the edges shared by species, we first construct a global (i.e. considering all the species within the dataset) matrix of species similarities by applying the Gower similarity index^112^ to the species-by-traits table using the function gowdis of the package ‘FD’ v.1.0 (ref.^113,114^). We used a modified version of the Gower index extended to accommodate variables in ordinal scales^115^. Using the species similarity matrix, we built a global tree of species similarities based on an unweighted pair group method with arithmetic mean (UPGMA) hierarchical cluster using function hclust of package ‘stats’ v.4.0.2 (ref.^108^). The length of the edge shared by two species was estimated as the distance between the intersection of two species in the global tree to the root of the tree (Fig. 3b). Based on the length of those edges, functional similarities between the pairs of communities were estimated using the equation 1. Therefore, even if two communities do not share any species, taxonomic similarity would be lower than functional similarity in case of the comparison of a continuous functional trait (e.g., body size; Fig. 3b). Note that the calculation of similarities was carried out within each dataset separately. Details of the calculation of similarities using the Sørensen-based indices for occurrence and abundance (i.e., percentage differences index) data can be found in the Appendix S2. We used both occurrence and abundance data because occurrences should be very informative about the drivers and patterns of communities along geographic gradients while abundances should inform well patterns along environmental gradients^116^. Main results are given for occurrence data in the main text, and abundance-based results can be found in Appendix S3.

### Spatial and environmental distances

We estimated the spatial and environmental distances between all the pairs of sites separately for each dataset. Spatial distances within each dataset were calculated as the geographic distance in kilometres between the pairs of sites using the function earth.dist of the package ‘fossil’ v.0.4.0 (ref.^117^; Fig. 3b). To estimate environmental distances, we first standardized the environmental variables to µ = 0 and σ = 1. Then, we calculated the environmental distance between sites as the Euclidean distance using the measured and standardized environmental variables for all the pairs of sites within each dataset (Fig. 3b) using the function vegdist of the package ‘vegan’ v.2.5-6 (ref.^118^). Because the datasets comprised different number and types of environmental variables, the values of environmental distance were context-dependent and not very informative for comparison across datasets. We therefore assumed that the environmental gradient scaled positively with spatial extent and rescaled the actual environmental distance to range between zero and one in each dataset by dividing actual values by the average environmental distance of the dataset.

### Distance decay of similarity

We modelled the distance decay of similarity following a negative exponential curve between the community similarity and distance^12^. This is because maximum spatial distances within our datasets were on average 795.5 kilometres; 95% CI [506.08, 1084.95], and therefore, it is highly likely that many of the datasets encompassed multiple species pools. One of the main assumptions of the distance decay is that Sij > Sjk if the distance between the sites *i* and *j* is shorter than the distance between *j* and *k*^12^. That is, the slope of the relationship should be negative, and positive slopes suggest either periodicity in the environmental gradient or a mismatch between the communities and the measured environmental variables^11^. Here, we calculated distance decay separately for taxonomic and functional similarities along spatial and environmental distance using a generalized linear model (GLM) following a binomial distribution of errors with a log link^119^ (Fig. 3c). Following Latombe et al.^120^, we included a negative constraint in GLMs such that the slopes are forced to be negative (i.e., slope <= 0). Besides, we included a negative constraint to the intercept of the model such that intercept <= 0. Therefore, because *e*^0^ = 1, we avoided intercept values that fall outside the range of taxonomic and functional similarities. We forced the negative coefficients via a non-positive least-square regression^121,122^ within the iterative re-weighted least-square algorithm^123^ implemented in the function glm.cons of the package ‘zetadiv’ v.1.2.0 (ref.^120,124^). We estimated a pseudo-R² (hereafter r^2^) as 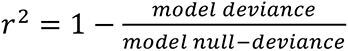 (2). Because of the pairwise structure of the data, similarities are non-independent, so we performed a leave-one-out Jack knife procedure to obtain the mean and confidence interval of the intercepts and slopes for each model^119^. Within such framework, the slope represents the *rate of decay*, that is, the proportion of similarity loss per unit distance, and the r^2^ represents the *strength of the relationship* between similarity and distance. Although it can be argued that slopes and r^2^ are highly correlated, the correlation between slopes and r^2^ in this study was small (Pearson’s r = 0.10; p-value = 0.240).

### Statistical analysis

We tested our hypothesis using two different approaches. Firstly, we investigated whether taxonomic or functional distance decay is stronger along spatial and environmental distances (H_1_) by performing a pairwise t-test to compare r^2^ drawn from GLMs using taxonomic similarity and the GLMs using functional similarity for each dataset (Fig. 3d). Totally, we carried out two pairwise t-tests, one considering the r^2^ from the models using spatial distances, and a second considering the r^2^ from the models using environmental distances.

We also investigated the ecological and geographical factors driving the rate of the distance decay across datasets. Each dataset was characterized with respect to (i) latitude, recorded as the absolute mean value of all the sites of the dataset; (ii) spatial extent, expressed as the largest pairwise distance (in km) between study sites; (iii) realm, classified into freshwater, marine and terrestrial environments; (iv) body size, estimated at organism-level as the log transformed fresh weight (g) drawn from literature^47,125^; (v) dispersal mode, classified as active and passive modes and organisms dispersed by seeds; (vi) taxonomic γ-diversity expressed as the total number of species in the dataset; (vii) functional γ-diversity, measured as the total volume of the union of the n-dimensional hypervolumes estimated within the dataset; (viii) total number of study sites in the dataset and (ix) the number of environmental variables in the dataset. For body sizes, we note that although the size range within the biotic group may be large (up to five orders of magnitude), it is small compared to the overall variation obtained across organism groups (twelve orders of magnitude). For more details on body size approximations, see refs.^47,49^. The taxonomic γ-diversity was included to study if there is a typical positive relationship between γ-diversity (taxonomic and functional) and β-diversity^7,52^. Functional γ-diversity was estimated based on geometrical n-dimensional hypervolumes^126,127^. We used the species functional similarity matrix based on Gower’s index (see the ‘taxonomic and functional similarities’ section) to extract orthogonal synthetic trait axes through a principal coordinate analysis^128^. Then, the hypervolume of each site within the dataset was calculated using a gaussian kernel density estimate via the function kernel.alpha of the package ‘BAT’^129^. The hypervolume of all sites were sequentially merged using the function hypervolume_set of the package ‘hypervolume’ v.2.0.12 (ref.^130^), and the united-hypervolume was used to estimate the total amount of functional space occupied by all the species within the dataset using the function get_volume of the package ‘hypervolume’. Because trait dimensionality affects the accuracy of the functional separation of species^131,132^, we standardized the number of dimensions to seven synthetic traits axes for all datasets. Hypervolumes are expressed in units of SDs to the power of the number of trait dimensions used (i.e., seven). The number of study sites and the number of environmental variables for each dataset were included to explore their potential effect on distance decay.

Finally, we used boosted regression trees (BRT) to test the effects of latitude (**H_2_**), spatial extent (**H_3_**) and realm (**H_4_**) on the rate of taxonomic and functional distance decay along spatial and environmental distance across the datasets. In addition, we included dispersal mode, body size, taxonomic and functional γ-diversity, number of sites, and number of environmental variables in the dataset as predictors in the BRTs (Fig. 3d). BRT is a regression modelling technique able to fits nonlinear relationships between predictor and response variables, including interaction among variables by using a boosting strategy to combine results from a large number (usually thousands) of simple regression tree models^133^. Our BRT outputs included graphs of the shapes of relationships between predictors and the response variable (e.g., linear, curvilinear and sigmoidal response shapes) and a relative importance of predictor variables. We also plotted a LOESS line on these plots to allow for easy visualization of the central tendency of the predicted values. Relative importance is constructed by counting the number of times a variable is selected for splitting in each tree, weighted by the squared improvement of the model as a result of each split, and averaged over all trees (see ref.^133–135^ for more details). BRT parameters were selected to amplify the deviance explained by the model. We tested interaction depth between 2 and 5, and the learning rates of 0.1, 0.01, and 0.001. The best models were the ones with learning rate of 5 and interaction depth of 0.001.We performed a 50–50 cross-validation procedure and estimated the model performance 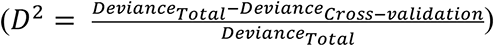 following Leathwick et al.^107^. As the datasets in this study have not always followed the same sampling methodology, and show different functional traits and environmental variables, we fitted the BRT models following a Laplace distribution of the errors to reduce the absolute error loss from the variation among datasets. BRT models were fitted using the function gbm.step of the package ‘dismo’ v.1.1-4 (ref.^136^).

Main results show the distance decay results based on total similarities (equation 1), but we also partitioned the similarities into replacement and richness difference components following the methodology described in the Appendix S2. Replacement gives the variation as a result of the substitution of species (turnover) or functional traits (functional replacement), and richness differences accounts for the variation as a result of net differences induced by the loss/gain of species or traits^137^. We only show the results of the partitioned components using occurrence data for simplicity. The final figures were prepared using the tools from the tidyverse environment^138^ in the R software v.4.0.2 (ref.^108^).

## Results

### Strength of the distance decay

The taxonomic and functional similarities had a mean correlation of 0.74 (sd ± 0.20) within datasets. The distance decays showed a wide range of shapes, from very steep decays to almost flat relationships (Fig. 4). The average r^2^ using occurrence data for taxonomic similarities was 0.099 (sd ± 0.129) and 0.061 (sd ± 0.091) for functional similarities. Spatial distance decays of taxonomic similarities were significantly stronger than the distance decays of functional similarities when considering both occurrence (Fig 4a; t = 6.330, p < 0.001, df = 148) and abundance data (Appendix S3, Fig. S1), supporting **H_1a_ –** spatial distance decay is stronger for taxonomic than functional similarities (Fig. 4a).

**Figure 4.**
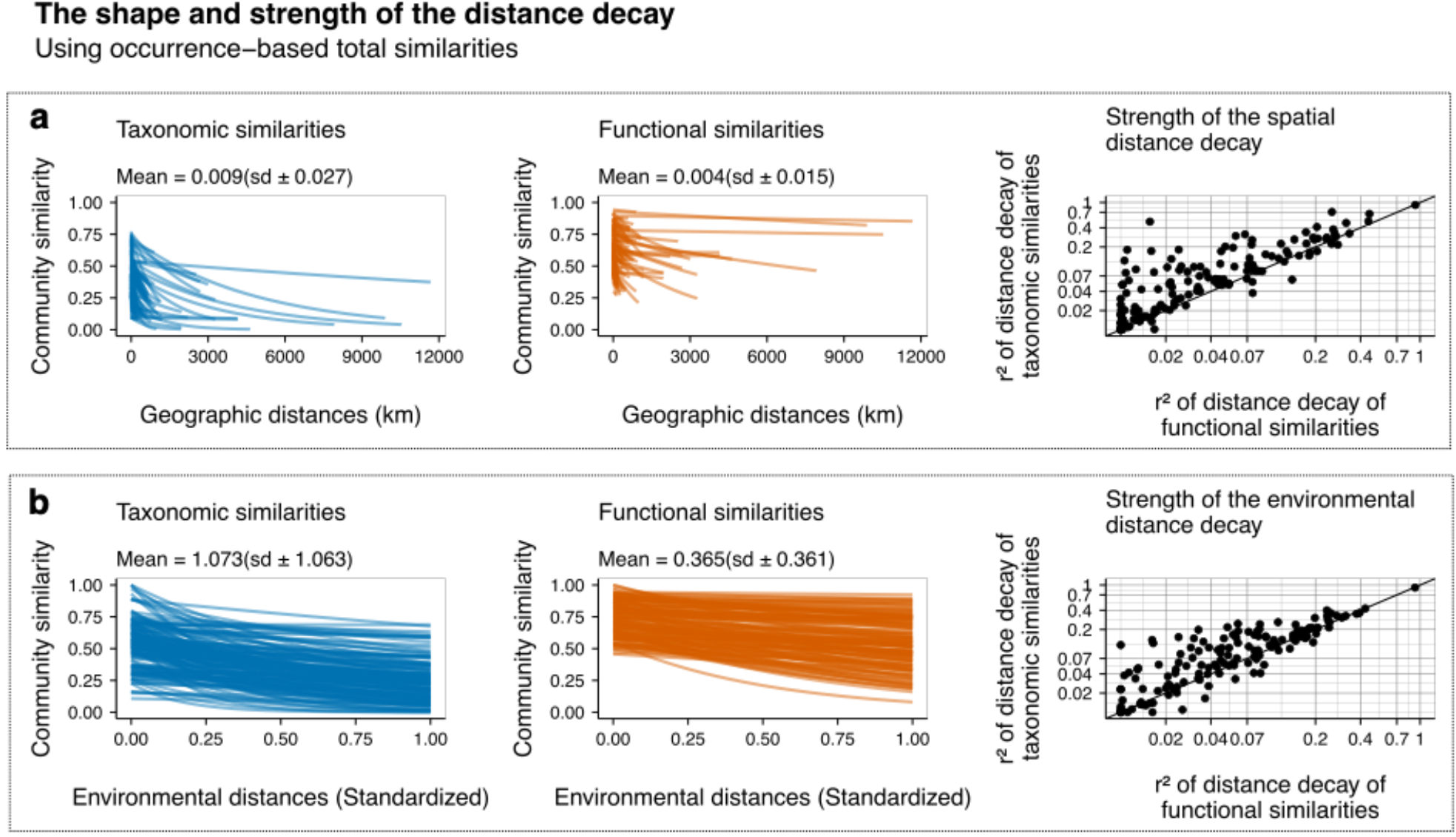
The distance decay along (a) spatial distance, and (b) environmental distance. Each line in the panels of left and middle columns shows the shape of the distance decay of an individual dataset. The mean and standard deviation of slopes are given in the plots. The blue lines show the distance decay of taxonomic similarity while the orange lines show the distance decay of functional similarity. The panels on the right column show the strength of the distance decay of taxonomic (y- axis) and functional (x-axis) similarity. The 1:1 line marks the equivalence of r^2^ between taxonomic and functional similarities. The dots below the line indicate a dataset with stronger decay of functional than taxonomic similarity, whereas circles above the line indicates stronger decay of taxonomic than functional similarities.

However, our results did not support **H_1b_** as the distance decay for taxonomic similarities (mean r^2^ = 0.103, sd ± 0.095) were also, on average, stronger than for functional similarities (mean r^2^ = 0.076, sd ± 0.086) along environmental distances (Fig 4b; t = 6.935, p < 0.001, df = 148). Note, however, that 41 out of 149 datasets had stronger distance decay of functional similarities than taxonomic similarities along environmental gradients. Most of the biotic groups had at least one dataset with a stronger relationship for functional similarities than for taxonomic similarities, except for corals, foraminifera, lichens, amphibians and fungi each of which comprised only one dataset.

### Rate of the distance decay

The mean slope of the spatial distance decay was 0.009 (sd ± 0.027) for taxonomic similarities, and 0.004 (sd ± 0.015) for functional similarities (Fig 4a). For environmental distances, the mean slope of the distance decay was 1.073 (sd ± 1.063) for taxonomic similarities and 0.365 (sd ± 0.361) for functional similarities (Fig 4b). Regarding the biotic groups, terrestrial plants had the steepest slopes along spatial distance both for taxonomic and functional similarities (Fig. 5). Along environmental distance, corals had the steepest slopes (Fig. 5). Similar patterns were found for abundance-based similarities, except for the biotic groups, where aquatic plants had the steepest slopes along spatial distances (Appendix S3).

**Figure 5.**
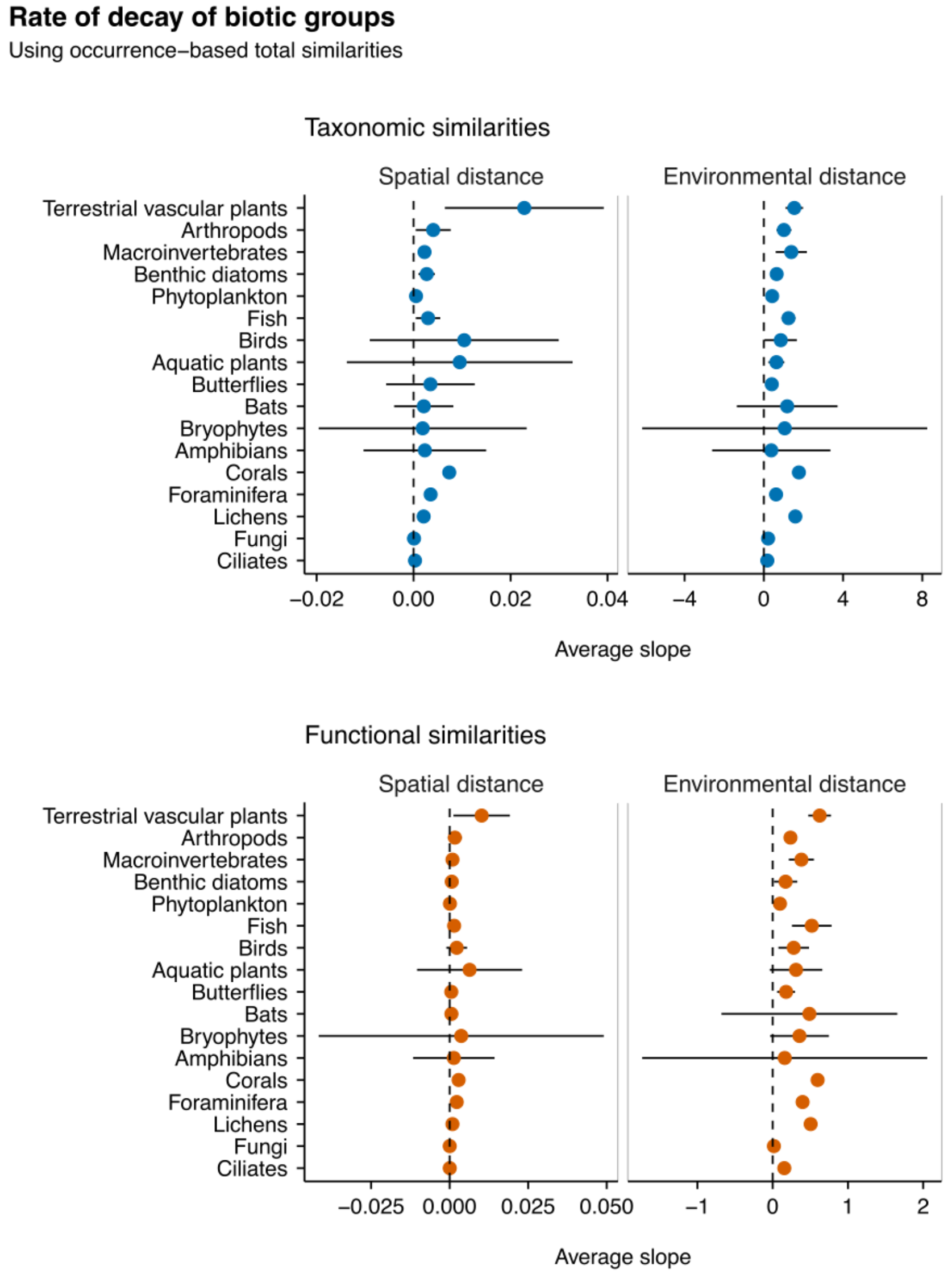
The average rate of decay of biotic groups using occurrence data along spatial and environmental distance. The vertical dotted lines highlight the zero rate (absence of decay) and the horizontal lines indicate the standard deviation of the mean. The blue circles show the rate of decay of taxonomic similarities while the orange circles show the rate of decay of functional similarities.

Across datasets, BRT explained 36.51% of the deviance of the slopes of the spatial distance decay for taxonomic similarities, and 36.86% for functional similarities using occurrence data. For the distance decay along environmental distances, BRT explained 14.43% of the deviance of the slopes of the decay of taxonomic similarities and 20.40% for functional similarities. Spatial extent and γ- diversity contributed most to the variation in slopes along either spatial or environmental distance using both occurrence and abundance-based similarities (Fig. 6 – 7a, Appendix S3).

**Figure 6.**
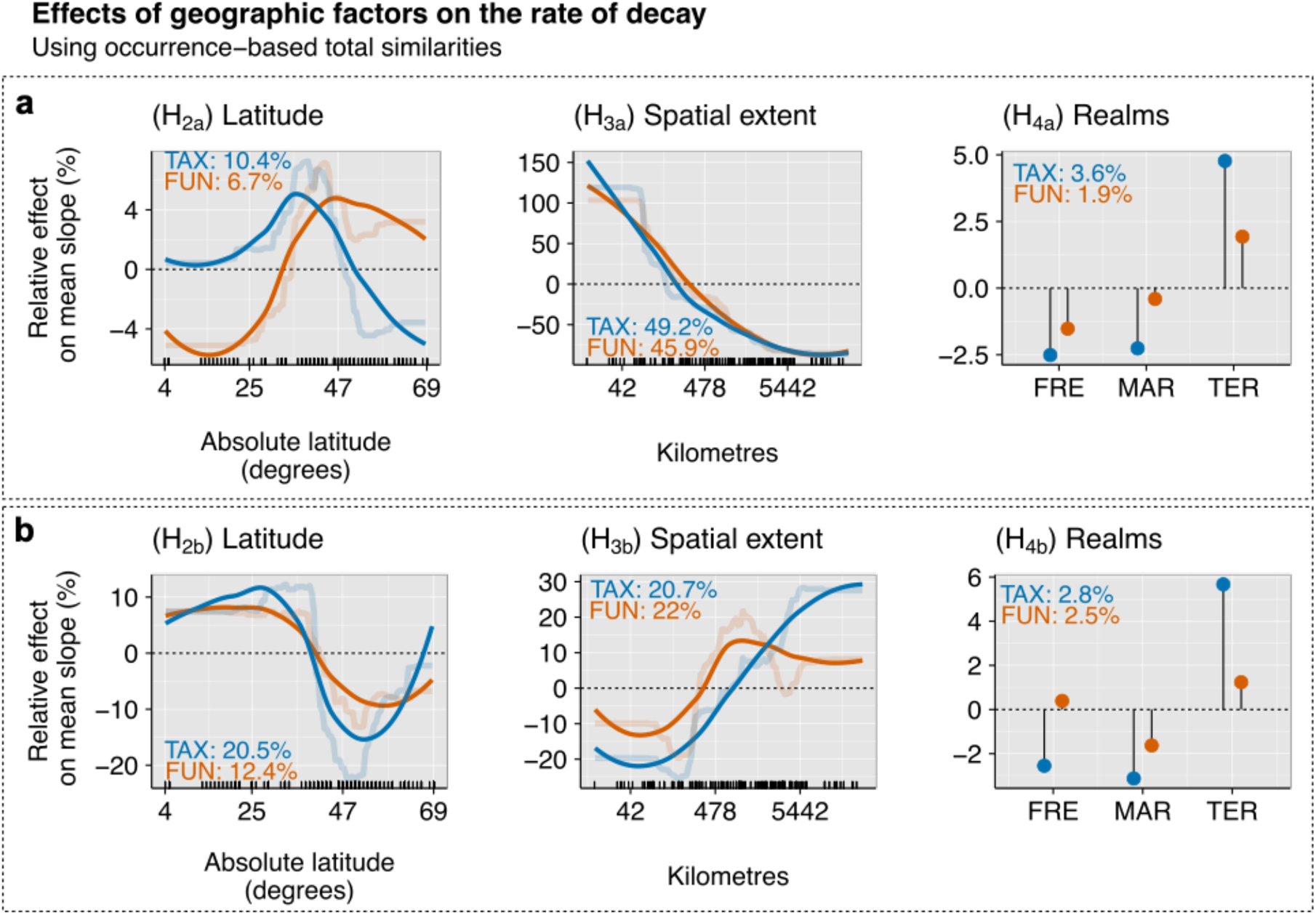
Relative effects (%) of geographic factors on the rate of decay along spatial (a) and environmental (b) distance decay of the total component of taxonomic (TAX - blue) and functional (FUN - orange) similarities using occurrence data across datasets. Partial dependence plots show the effects of a predictor variable on the response variable after accounting for the average effects of all other variables in the model. Semi-transparent lines represent the actual predicted effects; solid lines represent LOESS fits to predicted values from BRT. We show here only the variables related to the specific hypotheses, i.e., latitude, spatial extent, and realms (FRE = Freshwater, TER = Terrestrial, MAR = Marine).

**Figure 7.**
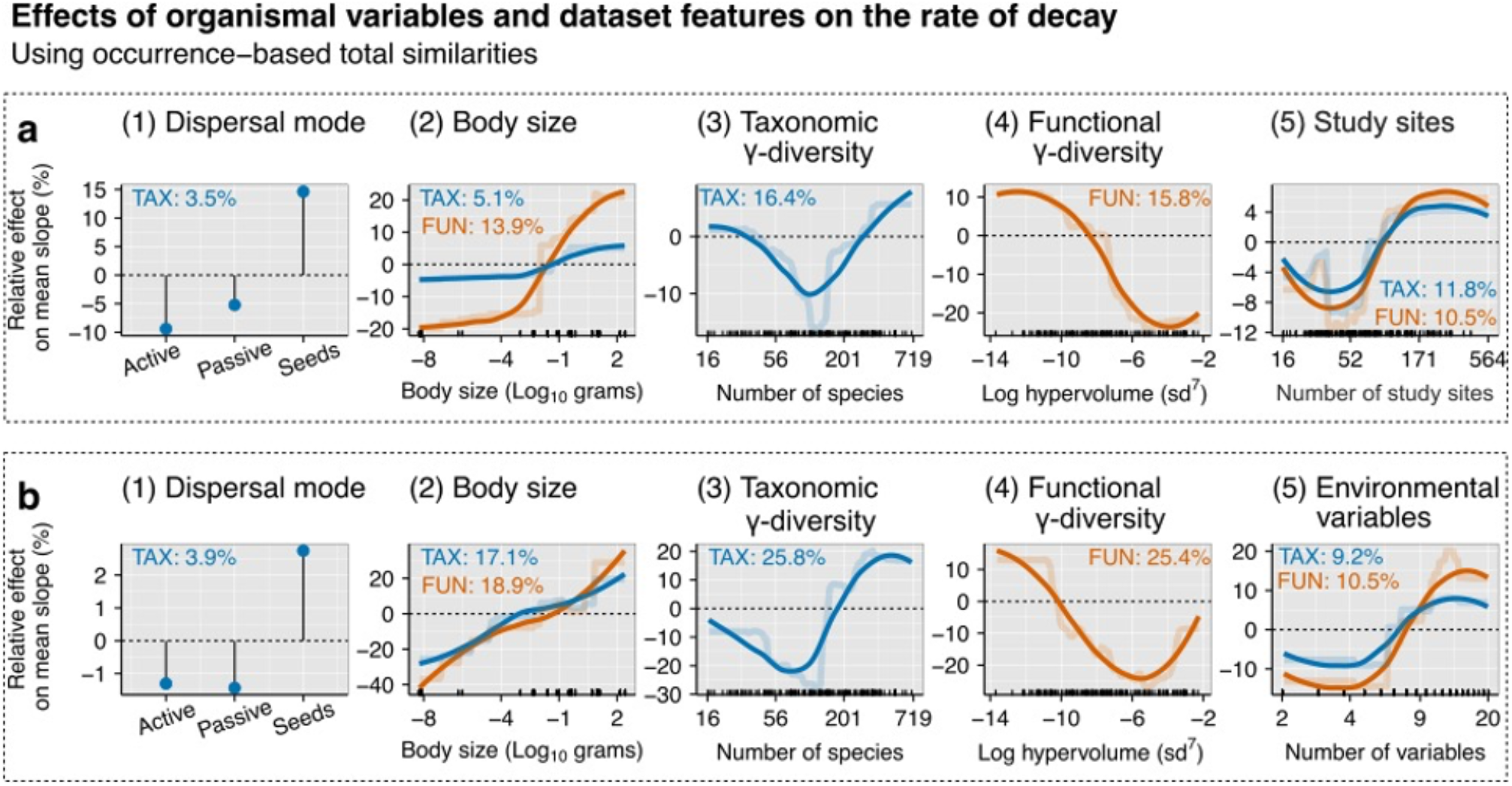
Relative effects (%) of organismal variables and dataset features on the rate of decay along spatial (a) and environmental (b) distance considering the total component of taxonomic (blue lines) and functional (orange lines) similarities using occurrence data across datasets. Partial dependence plots show the effects of a predictor variable on the response variable after accounting for the average effects of all other variables in the model. Semi-transparent lines represent the actual predicted effects; solid lines represent LOESS fits to predicted values from BRT. We show here the organismal variables and the variables related to the dataset features.

#### Latitudinal patterns

The slopes of spatial distance decay of both taxonomic and functional similarities were the steepest in datasets centred at ca. 35–45°, partly supporting **H_2a_** that distance decay was flatter at high latitudes (Fig. 6a). However, note that taxonomic spatial distance decay sharply decreased towards the poles. The slopes of environmental distance decay were flatter in the datasets from high latitudes (Fig. 6b), providing no support to hypothesis **H_2b_**.

#### Spatial extent

The distance decay of taxonomic and functional similarities was flatter in the datasets that covered larger spatial extent both for occurrence (Fig. 6a) and abundance data (Appendix S3, Fig. S3a), supporting hypothesis **H_3a_ –** distance decay becomes flatter with increasing spatial extent. For environmental distances, distance decay was steeper in the datasets that covered larger spatial extents for both taxonomic and functional similarities, agreeing thus with **H_3b_** that distance decay would become steeper with larger spatial extent.

#### Realms

Marine ecosystems had flatter slopes compared to freshwater or terrestrial ecosystems considering environmental distances, but not for spatial distances, thus partly agreeing with **H_4_** (Fig. 6). However, the importance of the realms in BRTs was overall low. A similar pattern emerged for abundance- based similarities (Appendix S3, Fig. S3).

#### Organismal variables and dataset features

The slopes of both spatial and environmental distance decays were steeper for larger-bodied organisms in taxonomic and functional similarity (Fig. 7a–b). Organisms relying on seed dispersal had steeper slopes along spatial and environmental distances than other dispersal types, but the overall importance of dispersal mode was low (Fig. 7b). Taxonomic γ-diversity had a U-shaped relationship with slopes for distance decay along spatial and environmental distances (Fig. 7b). Slopes of distance decay had an overall decreasing trend for functional γ-diversity for both spatial and environmental distances (Fig. 7a–b). Generally, slopes were steeper in the datasets where the number of study sites was higher (Fig. 7a), and flatter when datasets comprised only a few environmental variables (Fig 7b).

#### Replacement and richness differences

The slopes of taxonomic replacement along spatial distance decreased rapidly in the datasets above 35° while the functional replacement peaked at ca. 45° (Appendix S4, Fig. S1a). Along environmental distance, the taxonomic replacement increased towards higher latitudes while the functional replacement did not vary notably along latitude (Appendix S4, Fig. S1b). For the richness differences component, the slopes of both taxonomic and functional similarities were the steepest in the datasets at ca. 45° degrees for the spatial distance decay (Appendix S4, Fig. S2a). For environmental distances, slopes became flatter from low to high latitudes up to ca. 50° degrees for taxonomic similarities while for functional similarities, slopes did not vary along latitude (Appendix S4, Fig. S2b). Both replacement and richness differences showed flatter spatial slopes with increasing spatial extent (Appendix S4, Fig S1-S2). In contrast, environmental slopes increased with spatial extent only replacement (Appendix S4, Fig. S1b) while the effects of spatial extent for the slopes of richness differences along environment was very low (Appendix S4, Fig. S2b). Furthermore, marine ecosystems showed the flattest slopes of replacement along environmental gradients (Appendix S4, Fig. S1b) while freshwater ecosystems had the flattest slopes of richness differences (Appendix S4, Fig. S2b). Details about the organismal variables and datasets features can be found in the Appendix S4.

## Discussion

Community ecology and biogeography have lacked a comprehensive evaluation of functional β- diversity across different taxa and ecosystems globally. Earlier studies suggest that functional β- diversity better reflects environmental variability compared with taxonomic β-diversity, and that focusing on functional β-diversity may help, for example, understand how humans impact ecosystems by modifying the local environment^33,39–41^. This is because functional traits should reflect best the ecological requirements of species. Using a comparative analysis across biotic groups, ecosystem types and realms, we show here that (i) taxonomic distance decay is generally stronger along spatial gradients than functional distance decay, and that (ii) the decay of functional similarities along environmental gradients is typically not stronger than the decay of taxonomic similarities, unlike previously suggested.

### The strength of the distance decay of taxonomic and functional similarities

The stronger taxonomic than functional distance decay along space provides empirical evidence for the idea that the taxonomic distance decay is a robust approach for ecological and biogeographical studies, supporting **H_1a_**. Compositional differences effectively summarize dispersal-related factors as well as species responses to climatic and other spatially structured environmental variables. However, spatial distance decay of functional similarities may not reflect well geographic differences in biotic communities. This probably stems from the different roles played by deterministic and stochastic drivers when shaping taxonomic and functional composition: functional composition mirrors mostly local environmental filtering and typically does not strongly reflect dispersal limitations or species pool effects that influence stronger taxonomic composition^42^. Yet, the specific outcomes of any analysis of functional diversity depends on the functional traits included in the analysis^139^ and how researchers handle individual trait variability^140^. Also, some morphological or size-related traits with no clear functional meaning may turn out informative when exploring geographic patterns in functional composition^42^. For example, functional traits rather than species identities explained more variability of tree communities along broad spatial gradients^141^ or the variation of phytoplankton communities along a large South America gradient^142^. Such findings point to the fact that the decisions about which functional traits to include in the analysis is critical.

Our analysis suggests that, overall, functional distance decay is also somewhat weaker than taxonomic distance decay along environmental gradients. However, this result is likely context- dependent, and the stronger functional than taxonomic distance decay depends on whether the species replaced from one community to another are a random subsample of functionally redundant species from the regional pool or not^34^. In fact, in 40 datasets, distance decay of functional similarities was stronger than taxonomic similarities along environmental gradients. The datasets with stronger distance decay of functional than taxonomic similarities spanned a broad range of latitudes, number of study sites and environmental variables. Therefore, for using such heterogeneous datasets, we are not able to provide any strict guidance on the choice of functional traits or environmental variables to be measured in future studies. For example, the dataset on grassland arthropods from the Biodiversity exploratories project had standardized traits and environmental variables, but only Homoptera out of four different taxa showed stronger functional than taxonomic distance decay along environmental gradients. One explanation is that the whole organisms are susceptible to environmental filtering, and each species comprises a set of traits that cannot be physically filtered as a response to the environment. Therefore, environmental filtering on a given trait of a species may also filter other traits simultaneously, or a given species may comprise a trait not filtered by the environment, which tends to increase the community similarity among sites. Yet, we emphasize that the variation in the rate of distance decay of functional similarities along environmental gradients across datasets was better explained in BRT than the variation in the rate of the distance decay of taxonomic similarities. This suggests that the taxonomic metrics may be more context dependent than the functional metrics along environmental gradients and that functional features may be more useful to generalize across taxa and ecosystems^24^. Furthermore, functional distance decay should not be much affected by dispersal effects and regional species pools as compared to taxonomic distance decay.

### The effects of latitude on the rate of distance decay

In addition to our master hypothesis, we investigated whether the rate of distance decay showed consistent variation across ecosystems, along geographic gradients and among major taxonomic groups. We did not find slower rates of decay in the datasets at higher latitudes, but rather, concurring with the recent meta-analysis of species turnover^143^, we found that taxonomic similarities decayed the fastest at mid latitudes, above which the rate lowered down. Traditionally, this pattern has been explained with the Rapoport’s rule, whereby there is an increase in species range size at higher latitudes^144^ and hence lower taxonomic turnover. Yet, such finding may also stem from landscape fragmentation that increases β-diversity^145^, especially at mid latitudes prone to strong human impact and at local spatial scales^50^. We also observed a faster rate of functional spatial distance decay towards poles, agreeing with our hypothesis. This may reflect the fact that the high species diversity of the tropics is mainly due to niche overlap^51^, which increases the functional redundancy and reduces the functional turnover^52^. Furthermore, the latitudinal decrease in the rate of abundance-based functional distance decay (Appendix S3, Fig. S1) suggests an optimal utilization of the functional space, as have been observed earlier exclusively for marine organisms^146^.

Taxonomic and functional distance decay along environmental gradients exhibited a clear minimum in the datasets near 50° while increasing notably from 60° towards the poles especially for taxonomic similarities. This result points to a breakpoint in total similarities that stems from richness differences, as the replacement component did not have similar breakpoints but, rather, had similar replacement levels in the tropics with decreasing trend at mid- and high latitudes. Latitudinal breakpoints in turnover have been found earlier^147^ in terrestrial vertebrates at ca. 30°, where turnover decreased substantially, while nestedness component increased. Soininen et al.^143^ found a breakpoint for turnover component at 41°, whereas there was no breakpoint in nestedness component. Present results suggest that the rate of distance decay is relatively similar through the extensive tropical region, whereas it either increases or decreases rapidly at mid latitudes, depending on β-diversity metric or whether this phenomenon is examined along spatial or environmental gradients.

### The effect of spatial extent on the rate of distance decay

The rate of spatial distance decay was slower in the datasets covering larger spatial extent as we hypothesized, perhaps suggesting that regional species pools are limited, and new species are not found constantly at the same frequency when extent is larger. Lower decay rates in larger study areas could also result from repeated patterns in environmental variation, that is, environmental patchiness or natural periodicity in the environment^11^. Agreeing with our hypothesis, we also found that the rate of decay along environmental distance was higher in the datasets covering larger spatial extent. These findings indicate that spatial distance decay is more affected by species pool effects and dispersal processes than environmental distance decay, possibly because the latter reflects more strongly the level of local deterministic environmental filtering processes. Similar evidence has accumulated from case studies conducted in various ecosystems^33,39,41,148^. The finding that the rate of distance decay along environmental distance was higher in the datasets covering larger extents indicates the stronger environmental filtering at larger study areas. We also note that, in our BRT models, extent and γ- diversity had by far the largest relative importance, suggesting that their interplay plays a key role in shaping distance decay.

### The effect of realm on the distance decay

We found evidence for a lower rate of distance decay in marine versus terrestrial or freshwater ecosystems. Moreover, we found very comparable distance decay slopes for terrestrial and freshwaters, and the factor ‘realm’ showed low relative importance in the BRT models. Overall, this finding agrees with earlier meta-review on β-diversity^19^, suggesting that large-scale diversity patterns are generally weaker in marine ecosystems^149^. However, marine ecosystems would have lower species turnover than freshwater or terrestrial systems^49^. As connectivity, energy flows, dispersal modes, body size structure and trophic dynamics differ substantially between dry and wet ecosystems^150^, it would be vital to investigate possible differences in turnover among the realms more closely.

### Organismal variables and dataset features

Organism size did seem to affect taxonomic or functional distance decay along spatial and environmental gradients as the slopes typically increased with organism body size. This may be because β-diversity should be low among the small microbial taxa with efficient passive dispersal^19^. The rationale behind such idea is that efficient dispersal homogenizes communities among sites resulting in lower β-diversity^151^. Body size is also a key driver of organisms’ biological complexity^152^, and it may be that smaller organisms show a much more limited set of trait combinations than macroorganisms, leading to a lower functional redundancy among larger species. Furthermore, our knowledge about the taxonomy and functional traits of organisms is typically size-dependent. For example, the identification of larger species is much easier than that of microorganisms, which also applies to the identification and measurement of soft functional traits^153,154^. Therefore, the values of β-diversity of small organisms may be typically underestimated.

Patterns in environmental distance decay were relatively congruent with spatial distance decay regarding dispersal mode, suggesting that taxa which disperse passively do not seem to track environmental gradients more efficiently compared with less dispersive taxa. It may also be that small-sized taxa were filtered along some unmeasured spatially-structured environmental gradients, and the pattern was thus detected as spatial turnover even if caused by some underlying unmeasured environmental factors. Forthcoming studies would greatly benefit from disentangling the signal of unmeasured environmental variables from true dispersal limitation^155^.

### Study design

There are also some possibly influential aspects in our study design that should be discussed. Although the study is global in its extent, the availability of datasets was not evenly distributed geographically. This is a well-known problem in biodiversity research^156^ that calls for complementary studies to verify that these trends hold true in poorly sampled regions.

Also, we relied on the suite of traits and environmental variables included in the original datasets and, thus, the collection of traits and environmental variables used differed somewhat among datasets even for the same focal taxonomic groups. This increases the uncertainty on how environmental variables filter the functional structure of communities in different contexts and how strong the taxonomic community-environment relationships are. An alignment of key traits and environmental variables is therefore desirable, but requires a suite of sister studies following the same protocol, which is unfortunately not yet available. Moreover, the fact that some of the biotic groups (e.g., corals, foraminifera) were underrepresented in our analysis with only one dataset included (Fig. 2), or the total lack of some taxa (e.g. aquatic and terrestrial mammals, bacteria), makes it more difficult to generalize distance decay across taxa.

### Concluding remarks

In summary, we believe our analysis is an important step towards a more comprehensive understanding of patterns and drivers of functional β-diversity, particularly in comparison with the patterns and drivers of taxonomic β-diversity that have so far attracted much more research interest compared with functional β-diversity. Here, we found that functional distance decay is scale- dependent and a product of large-scale geographic factors (latitude) and taxonomic and functional γ-diversity, but is also driven by organisms’ biology to some degree. In general, taxonomic distance decay provides a better tool for many aspects of biogeographical research, because it reflects dispersal-related factors as well as species responses to climatic and other typically spatially- structured environmental variables. However, functional distance decay may be a cost-effective option for investigating how humans impact ecosystems via modifying the environment. Overall, the present findings and data shed light into the congruence between the functional and taxonomic diversity patterns and provide useful new information to the field of functional biogeography.

## Author contributions

Caio Graco-Roza and Janne Soininen contributed equally to the original idea, data analysis, and the writing of the first draft. Jani Heino and Otso Ovaskainen advised on the main idea, analysis and commented on the first draft. Francesco Maria Sabatini coordinated the data compilation from the sPlot database and commented on the first draft, Martin Gossner coordinated the compilation of the data from the Biodiversity exploratories project and commented on the first draft. All other authors shown in alphabetical order contributed data and commented on the draft.

## Acknowledgements

This research was funded by the Coordination for the Improvement of Higher Education Personnel (CAPES), the Carlos Chagas Filho Research Support Foundation (FAPERJ), and the Ella and Georg Erhnrooth foundation. The sPlot project was initiated by sDiv, the Synthesis Centre of the German Centre for Integrative Biodiversity Research (iDiv) Halle-Jena-Leipzig, funded by the German Research Foundation (DFG FZT 118) and is now a platform of iDiv. The study was supported by the TRY initiative on plant traits (http://www.try-db.org). We are also grateful to Jens Kattge and TRY database. TRY is hosted, developed and maintained at the Max Planck Institute for Biogeochemistry (MPI-BGC) in Jena, Germany, in collaboration with the German Centre for Integrative Biodiversity Research (iDiv) Halle-Jena-Leipzig. The CESTES database of metacommunities is also an initiative of iDiv led by Alienor Jeliazkov. We thank sDiv for supporting the open science initiative.

## References

1. IPBES. Global assessment report on biodiversity and ecosystem services of the Intergovernmental Science-Policy Platform on Biodiversity and Ecosystem Services. (IPBES secretariat, 2019).

2. Hooper, D. U. et al. Effects of biodiversity on ecosystem functioning: a consensus of current knowledge. Ecological Monographs 75, 3–35 (2005).

3. Cardinale, B. J. et al. Biodiversity loss and its impact on humanity. Nature 486, 59–67 (2012).

4. Anderson, M. J. et al. Navigating the multiple meanings of β diversity: A roadmap for the practicing ecologist. Ecology Letters 14, 19–28 (2011).

5. Mori, A. S., Isbell, F. & Seidl, R. β-Diversity, Community Assembly, and Ecosystem Functioning. Trends in Ecology and Evolution vol. 33 549–564 (2018).

6. Qian, H., Ricklefs, R. E. & White, P. S. Beta diversity of angiosperms in temperate floras of eastern Asia and eastern North America. Ecology Letters 8, 15–22 (2004).

7. Kraft, N. J. B. et al. Disentangling the drivers of β diversity along latitudinal and elevational gradients. Science 333, 1755–1758 (2011).

8. Blowes, S. A. et al. The geography of biodiversity change in marine and terrestrial assemblages. Science 366, 339–345 (2019).

9. Gossner, M. M. et al. Land-use intensification causes multitrophic homogenization of grassland communities. Nature 540, 266–269 (2016).

10. Socolar, J. B., Gilroy, J. J., Kunin, W. E. & Edwards, D. P. How Should Beta-Diversity Inform Biodiversity Conservation? Trends in Ecology and Evolution 31, 67–80 (2016).

11. Nekola, J. C. & White, P. S. The distance decay of similarity in biogeography and ecology. Journal of Biogeography 26, 867–878 (1999).

12. Nekola, J. C. & McGill, B. J. Scale dependency in the functional form of the distance decay relationship. Ecography 37, 309–320 (2014).

13. Hubbell, S. P. The Unified Neutral Theory of Biodiversity and Biogeography. Monographs in Population Biology vol. 32 (Princeton University Press, 2001).

14. Cottenie, K. Integrating environmental and spatial processes in ecological community dynamics. Ecology Letters 8, 1175–1182 (2005).

15. Astorga, A. et al. Distance decay of similarity in freshwater communities: Do macro- and microorganisms follow the same rules? Global Ecology and Biogeography 21, 365–375 (2012).

16. Daws, S. C. et al. Do shared traits create the same fates? Examining the link between morphological type and the biogeography of fungal and bacterial communities. Fungal Ecology 46, 100948 (2020).

17. Mammola, S. et al. Local- versus broad-scale environmental drivers of continental *β* - diversity patterns in subterranean spider communities across Europe. Proceedings of the Royal Society B: Biological Sciences 286, 20191579 (2019).

18. Morlon, H. et al. A general framework for the distance-decay of similarity in ecological communities. Ecology Letters 11, 904–917 (2008).

19. Soininen, J., McDonald, R. & Hillebrand, H. The distance decay of similarity in ecological communities. Ecography 30, 3–12 (2007).

20. Petchey, O. L. On the statistical significance of functional diversity effects. Functional Ecology 18, 297–303 (2004).

21. Petchey, O. L. & Gaston, K. J. Functional diversity: Back to basics and looking forward. Ecology Letters 9, 741–758 (2006).

22. Villéger, S., Ramos Miranda, J., Flores Hernandez, D. & Mouillot, D. Low functional β- diversity despite high taxonomic β-diversity among tropical estuarine fish communities. PloS one 7, e40679–e40679 (2012).

23. Violle, C., Reich, P. B., Pacala, S. W., Enquist, B. J. & Kattge, J. The emergence and promise of functional biogeography. Proceedings of the National Academy of Sciences of the United States of America 111, 13690–13696 (2014).

24. McGill, B. J., Enquist, B. J., Weiher, E. & Westoby, M. Rebuilding community ecology from functional traits. Trends in Ecology and Evolution 21, 178–185 (2006).

25. Mouillot, D. et al. A functional approach reveals community responses to disturbances. Trends in Ecology and Evolution 28, 167–177 (2013).

26. Villéger, S., Mason, N. W. H. H. & Mouillot, D. New multidimensional functional diversity indices for a multifaceted framework in functional ecology. Ecology 89, 2290–2301 (2008).

27. Buisson, L., Grenouillet, G., Villéger, S., Canal, J. & Laffaille, P. Toward a loss of functional diversity in stream fish assemblages under climate change. Global Change Biology 19, 387–400 (2013).

28. Pool, T. K. & Olden, J. D. Taxonomic and functional homogenization of an endemic desert fish fauna. Diversity and Distributions 18, 366–376 (2011).

29. Villéger, S., Grenouillet, G. & Brosse, S. Decomposing functional β-diversity reveals that low functional β-diversity is driven by low functional turnover in European fish assemblages. Global Ecology and Biogeography 22, 671–681 (2013).

30. Villéger, S., Grenouillet, G. & Brosse, S. Functional homogenization exceeds taxonomic homogenization among European fish assemblages. Global Ecology and Biogeography 23, 1450–1460 (2014).

31. Penone, C et al. Global mammal beta diversity shows parallel assemblage structure in similar but isolated environments. Proceedings of the Royal Society B: Biological Sciences 283, 1–9 (2016).

32. Heino, J. & Tolonen, K. T. Ecological drivers of multiple facets of beta diversity in a lentic macroinvertebrate metacommunity. Limnology and Oceanography 62, 2431–2444 (2017).

33. Sokol, E. R., Benfield, E. F., Belden, L. K. & Maurice Valett, H. The Assembly of Ecological Communities Inferred from Taxonomic and Functional Composition. The American Naturalist 177, 630–644 (2011).

34. Swenson, N. G., Anglada-Cordero, P. & Barone, J. A. Deterministic tropical tree community turnover: Evidence from patterns of functional beta diversity along an elevational gradient. Proceedings of the Royal Society B: Biological Sciences 278, 877–884 (2011).

35. Leibold, M. A. & McPeek, M. A. Coexistence of the niche and neutral perspectives in community ecology. Ecology 87, 1399–1410 (2006).

36. Baiser, B. & Lockwood, J. L. The relationship between functional and taxonomic homogenization. Global Ecology and Biogeography 20, 134–144 (2011).

37. Mouton, T. L., Tonkin, J. D., Stephenson, F., Verburg, P. & Floury, M. Increasing climate- driven taxonomic homogenization but functional differentiation among river macroinvertebrate assemblages. Global Change Biology gcb.15389 (2020) doi:10.1111/gcb.15389.

38. Ricotta, C., Laroche, F., Szeidl, L. & Pavoine, S. From alpha to beta functional and phylogenetic redundancy. Methods in Ecology and Evolution 11, 487–493 (2020).

39. Meynard, C. N. et al. Beyond taxonomic diversity patterns: how do α, β and γ components of bird functional and phylogenetic diversity respond to environmental gradients across France? Global Ecology and Biogeography 20, 893–903 (2011).

40. Spasojevic, M. J., Yablon, E. A., Oberle, B. & Myers, J. A. Ontogenetic trait variation influences tree community assembly across environmental gradients. Ecosphere 5, 1–20 (2014).

41. Weinstein, B. G. et al. Taxonomic, Phylogenetic, and Trait Beta Diversity in South American Hummingbirds. The American Naturalist 184, 211–224 (2014).

42. Soininen, J., Jamoneau, A., Rosebery, J. & Passy, S. I. Global patterns and drivers of species and trait composition in diatoms. Global Ecology and Biogeography 25, 940–950 (2016).

43. Graco-Roza, C. et al. Functional rather than taxonomic diversity reveals changes in the phytoplankton community of a large dammed river. Ecological Indicators 107048 (2020) doi:10.1016/j.ecolind.2020.107048.

44. Soininen, J., Lennon, J. J. & Hillebrand, H. A multivariate analysis of beta diversity across organisms and environments. Ecology 88, 2830–2838 (2007).

45. Qian, H. Global comparisons of beta diversity among mammals, birds, reptiles, and amphibians across spatial scales and taxonomic ranks. Journal of Systematics and Evolution 47, 509–514 (2009).

46. Qian, H., Badgley, C. & Fox, D. L. The latitudinal gradient of beta diversity in relation to climate and topography for mammals in North America. Global Ecology and Biogeography 18, 111–122 (2009).

47. Hillebrand, H. On the generality of the latitudinal diversity gradient. American Naturalist 163, 192–211 (2004).

48. Soininen, J. Species Turnover along Abiotic and Biotic Gradients: Patterns in Space Equal Patterns in Time? BioScience 60, 433–439 (2010).

49. Drakare, S., Lennon, J. J. & Hillebrand, H. The imprint of the geographical, evolutionary and ecological context on species-area relationships. Ecology Letters 9, 215–227 (2005).

50. Rohde, K. Rapoport’s Rule is a local phenomenon and cannot explain latitudinal gradients in species diversity. Biodiversity Letters vol. 3 10–13 (1996).

51. May, R. M. & MacArthur, R. H. Niche overlap as a function of environmental variability. Proceedings of the National Academy of Sciences of the United States of America 69, 1109– 1113 (1972).

52. Lamanna, C. et al. Functional trait space and the latitudinal diversity gradient. Proceedings of the National Academy of Sciences of the United States of America 111, 13745–13750 (2014).

53. Terborgh, J. On the Notion of Favorableness in Plant Ecology. The American Naturalist 107, 481–501 (1973).

54. Soininen, J. & Hillebrand, H. Disentangling distance decay of similarity from richness gradients: response to Baselga (2007). Ecography 30, 842–844 (2007).

55. Cornwell, W. K. & Ackerly, D. D. Community assembly and shifts in plant trait distributions across an environmental gradient in coastal California. Ecological Monographs 79, 109–126 (2009).

56. Cornwell, W. K., Schwilk, D. W. & Ackerly, D. D. A trait-based test for habitat filtering: Convex hull volume. Ecology 87, 1465–1471 (2006).

57. Steinbauer, M. J., Dolos, K., Reineking, B. & Beierkuhnlein, C. Current measures for distance decay in similarity of species composition are influenced by study extent and grain size. Global Ecology and Biogeography 21, 1203–1212 (2012).

58. Triantis, K. A., Guilhaumon, F. & Whittaker, R. J. The island species-area relationship: biology and statistics. Journal of Biogeography 39, 215–231 (2011).

59. Pianka, E. R. Latitudinal Gradients in Species Diversity: A Review of Concepts. The American Naturalist 100, 33–46 (1966).

60. Questad, E. J. & Foster, B. L. Coexistence through spatio-temporal heterogeneity and species sorting in grassland plant communities. Ecology Letters 11, 717–726 (2008).

61. Bergholz, K. et al. Environmental heterogeneity drives fine-scale species assembly and functional diversity of annual plants in a semi-arid environment. Perspectives in Plant Ecology, Evolution and Systematics 24, 138–146 (2017).

62. Kadmon, R. & Allouche, O. Integrating the effects of area, isolation, and habitat heterogeneity on species diversity: A unification of island biogeography and niche theory. American Naturalist 170, 443–454 (2007).

63. Gazol, A. et al. A negative heterogeneity–diversity relationship found in experimental grassland communities. Oecologia 173, 545–555 (2013).

64. Laanisto, L. et al. Microfragmentation concept explains non-positive environmental heterogeneity–diversity relationships. Oecologia 171, 217–226 (2012).

65. Clarke, A. Is there a latitudinal diversity cline in the sea? Trends in Ecology & Evolution 7, 286–287 (1992).

66. Cornell, H. v & Harrison, S. P. What Are Species Pools and When Are They Important? Annual Review of Ecology, Evolution, and Systematics 45, 45–67 (2014).

67. Jenkins, D. G. et al. Does size matter for dispersal distance? Global Ecology and Biogeography 16, 415–425 (2007).

68. Bie, T. et al. Body size and dispersal mode as key traits determining metacommunity structure of aquatic organisms. Ecology Letters 15, 740–747 (2012).

69. Bruelheide, H. et al. sPlot – A new tool for global vegetation analyses. Journal of Vegetation Science 30, 161–186 (2019).

70. Jeliazkov, A. et al. A global database for metacommunity ecology, integrating species, traits, environment and space. Scientific data 7, 6 (2020).

71. Kattge, J. et al. TRY - a global database of plant traits. Global Change Biology 17, 2905– 2935 (2011).

72. Brind’Amour, A. et al. Relationships between species feeding traits and environmental conditions in fish communities: A three-matrix approach. Ecological Applications 21, 363– 377 (2011).

73. Chong-Seng, K. M., Mannering, T. D., Pratchett, M. S., Bellwood, D. R. & Graham, N. A. J. J. The influence of coral reef benthic condition on associated fish assemblages. PLoS ONE 7, 1–10 (2012).

74. Cleary, D. F. R. R. et al. Variation in the composition of corals, fishes, sponges, echinoderms, ascidians, molluscs, foraminifera and macroalgae across a pronounced in-to- offshore environmental gradient in the Jakarta Bay–Thousand Islands coral reef complex. Marine Pollution Bulletin 110, 701–717 (2016).

75. Ribera, I., Dolédec, S., Downie, I. S. & Foster, G. N. Effect of land disturbance and stress on species traits of ground beetle assemblages. Ecology 82, 1112–1129 (2001).

76. Pakeman, R. J. Multivariate identification of plant functional response and effect traits in an agricultural landscape. Ecology 92, 1353–1365 (2011).

77. Bagaria, G., Pino, J., Rodà, F. & Guardiola, M. Species traits weakly involved in plant responses to landscape properties in Mediterranean grasslands. Journal of Vegetation Science 23, 432–442 (2012).

78. Frenette-Dussault, C., Shipley, B., Léger, J.-F., Meziane, D. & Hingrat, Y. Functional structure of an arid steppe plant community reveals similarities with Grime’s C-S-R theory. Journal of Vegetation Science 23, 208–222 (2011).

79. Fried, G., Kazakou, E. & Gaba, S. Trajectories of weed communities explained by traits associated with species’ response to management practices. Agriculture, Ecosystems and Environment 158, 147–155 (2012).

80. Raevel, V., Violle, C. & Munoz, F. Mechanisms of ecological succession: Insights from plant functional strategies. Oikos 121, 1761–1770 (2012).

81. Eallonardo, A. S., Leopold, D. J., Fridley, J. D. & Stella, J. C. Salinity tolerance and the decoupling of resource axis plant traits. Journal of Vegetation Science 24, 365–374 (2013).

82. Jamil, T., Ozinga, W. A., Kleyer, M. & ter Braak, C. J. F. F. Selecting traits that explain species-environment relationships: A generalized linear mixed model approach. Journal of Vegetation Science 24, 988–1000 (2013).

83. Meffert, P. J. & Dziock, F. The influence of urbanisation on diversity and trait composition of birds. Landscape Ecology 28, 943–957 (2013).

84. Chmura, D., Żarnowiec, J. & Staniaszek-Kik, M. Interactions between plant traits and environmental factors within and among montane forest belts: A study of vascular species colonising decaying logs. Forest Ecology and Management 379, 216–225 (2016).

85. Robroek, B. J. M. M. et al. Taxonomic and functional turnover are decoupled in European peat bogs. Nature Communications 8, 1161 (2017).

86. van Klink, R. et al. No detrimental effects of delayed mowing or uncut grass refuges on plant and bryophyte community structure and phytomass production in low-intensity hay meadows. Basic and Applied Ecology 20, 1–9 (2017).

87. Díaz, A. M., Alonso, M. L. S. & Gutiérrez, M. R. V.-A. Biological traits of stream macroinvertebrates from a semi-arid catchment: patterns along complex environmental gradients. Freshwater Biology 53, 1–21 (2007).

88. Gallardo, B., Gascón, S., García, M. & Comín, F. A. Testing the response of macroinvertebrate functional structure and biodiversity to flooding and confinement. Journal of Limnology 68, 315–326 (2009).

89. Jeliazkov, A. et al. Level-dependence of the relationships between amphibian biodiversity and environment in pond systems within an intensive agricultural landscape. Hydrobiologia 723, 7–23 (2014).

90. Barbaro, L. & van Halder, I. Linking bird, carabid beetle and butterfly life-history traits to habitat fragmentation in mosaic landscapes. Ecography 32, 321–333 (2009).

91. Dziock, F. et al. Reproducing or dispersing? Using trait based habitat templet models to analyse Orthoptera response to flooding and land use. Agriculture, Ecosystems and Environment 145, 85–94 (2011).

92. Gibb, H. et al. Responses of foliage-living spider assemblage composition and traits to a climatic gradient in Themeda grasslands. Austral Ecology 40, 225–237 (2015).

93. Barbosa, E. P. et al. Fine-scale Beta-diversity Patterns Across Multiple Arthropod Taxa Over a Neotropical Latitudinal Gradient. Biotropica 47, 588–594 (2015).

94. Krasnov, B. R. et al. Assembly rules of ectoparasite communities across scales: Combining patterns of abiotic factors, host composition, geographic space, phylogeny and traits. Ecography 38, 184–197 (2015).

95. Spake, R., Barsoum, N., Newton, A. C. & Doncaster, C. P. Drivers of the composition and diversity of carabid functional traits in UK coniferous plantations. Forest Ecology and Management 359, 300–308 (2016).

96. Lowe, E. C., Threlfall, C. G., Wilder, S. M. & Hochuli, D. F. Environmental drivers of spider community composition at multiple scales along an urban gradient. Biodiversity and Conservation 27, 829–852 (2018).

97. Gonçalves-Souza, T., Romero, G. Q. & Cottenie, K. Metacommunity versus Biogeography: A Case Study of Two Groups of Neotropical Vegetation-Dwelling Arthropods. PLoS ONE 9, e115137 (2014).

98. Bartonova, A., Benes, J., Fric, Z. F., Chobot, K. & Konvicka, M. How universal are reserve design rules? A test using butterflies and their life history traits. Ecography 39, 456–464 (2016).

99. Cleary, D. F. R. R. et al. Bird species and traits associated with logged and unlogged forest in Borneo. Ecological Applications 17, 1184–1197 (2007).

100. Barbaro, L., Brockerhoff, E. G., Giffard, B. & van Halder, I. Edge and area effects on avian assemblages and insectivory in fragmented native forests. Landscape Ecology 27, 1451–1463 (2012).

101. Barbaro, L. et al. Avian pest control in vineyards is driven by interactions between bird functional diversity and landscape heterogeneity. Journal of Applied Ecology 54, 500–508 (2017).

102. Charbonnier, Y. M. et al. Bat and bird diversity along independent gradients of latitude and tree composition in European forests. Oecologia 182, 529–537 (2016).

103. Farneda, F. Z. et al. Trait-related responses to habitat fragmentation in Amazonian bats. Journal of Applied Ecology 52, 1381–1391 (2015).

104. Robinson, N., Kadlec, T., Bowers, M. D. & Guralnick, R. P. Integrating species traits and habitat characteristics into models of butterfly diversity in a fragmented ecosystem. Ecological Modelling 281, 15–25 (2014).

105. Rachello-Dolmen, P. G. & Cleary, D. F. R. R. Relating coral species traits to environmental conditions in the Jakarta Bay/Pulau Seribu reef system, Indonesia. Estuarine, Coastal and Shelf Science 73, 816–826 (2007).

106. Cleary, D. F. R. R. & Renema, W. Relating species traits of foraminifera to environmental variables in the Spermonde Archipelago, Indonesia. Marine Ecology Progress Series 334, 73–82 (2007).

107. Leathwick, J. R., Elith, J. & Hastie, T. Comparative performance of generalized additive models and multivariate adaptive regression splines for statistical modelling of species distributions. Ecological Modelling 199, 188–196 (2006).

108. R Core Team. R: A Language and Environment for Statistical Computing. (2020).

109. Cardoso, P., Mammola, S., Rigal, F. & Carvalho, J. C. BAT: Biodiversity Assessment Tools. (2020).

110. Cardoso, P. et al. Partitioning taxon, phylogenetic and functional beta diversity into replacement and richness difference components. Journal of Biogeography 41, 749–761 (2014).

111. Podani, J., Ricotta, C. & Schmera, D. A general framework for analyzing beta diversity, nestedness and related community-level phenomena based on abundance data. Ecological Complexity 15, 52–61 (2013).

112. Gower, J. C. A General Coefficient of Similarity and Some of Its Properties. Biometrics 27, 857 (1971).

113. Laliberté, E. & Legendre, P. A distance-based framework for measuring functional diversity from multiple traits. Ecology 91, 299–305 (2010).

114. Laliberté, E., Legendre, P. & Shipley, B. FD: measuring functional diversity from multiple traits, and other tools for functional ecology. (2014).

115. Podani, J. Extending Gower’s general coefficient of similarity to ordinal characters. Taxon 48, 331–340 (1999).

116. Declerck, S. A. J. J., Coronel, J. S., Legendre, P. & Brendonck, L. Scale dependency of processes structuring metacommunities of cladocerans in temporary pools of High-Andes wetlands. Ecography 34, 296–305 (2011).

117. Vavrek, M. J. fossil: palaeoecological and palaeogeographical analysis tools. Palaeontologia Electronica 14, 1T (2011).

118. Oksanen, J., et al. vegan: Community Ecology Package. (2019).

119. Millar, R. B., Anderson, M. J. & Tolimieri, N. Much ado about nothings: using zero similarity points in distance-decay curves. Ecology vol. 92 (2011).

120. Latombe, G., Hui, C. & McGeoch, M. A. Multi-site generalised dissimilarity modelling: using zeta diversity to differentiate drivers of turnover in rare and widespread species. Methods in Ecology and Evolution 8, 431–442 (2017).

121. Mullen, K. M. & van Stokkum, I. H. M. nnls: The Lawson-Hanson algorithm for non- negative least squares (NNLS). (2012).

122. Lawson, C. L. & Hanson, R. J. Solving Least Squares Problems. Solving Least Squares Problems (1995). doi:10.1137/1.9781611971217.

123. Marschner, I. C. glm2: Fitting generalized linear models with convergence problems. The R Journal 3, 12–15 (2011).

124. Latombe, G., McGeoch, M. A., Nipperess, D. A. & Hui, C. zetadiv: Functions to Compute Compositional Turnover Using Zeta Diversity. (2020).

125. Peters, R. H. The Ecological Implications of Body Size. (Cambridge University Press, 1983). doi:10.1017/cbo9780511608551.

126. Blonder, B., Lamanna, C., Violle, C. & Enquist, B. J. The n-dimensional hypervolume. Global Ecology and Biogeography 23, (2014).

127. Blonder, B. Hypervolume concepts in niche- and trait-based ecology. Ecography 41, 1441– 1455 (2018).

128. Gower, J. C. Some Distance Properties of Latent Root and Vector Methods Used in Multivariate Analysis. Biometrika 53, (1966).

129. Mammola, S. & Cardoso, P. Functional diversity metrics using kernel density n- dimensional hypervolumes. Methods in Ecology and Evolution 11, (2020).

130. Blonder, B. & Harris, D. J. hypervolume: High Dimensional Geometry and Set Operations Using Kernel Density Estimation, Support Vector Machines, and Convex Hulls. (2019).

131. Podani, J. & Schmera, D. On dendrogram-based measures of functional diversity. Oikos 115, 179–185 (2006).

132. Maire, E., Grenouillet, G., Brosse, S. & Villéger, S. How many dimensions are needed to accurately assess functional diversity? A pragmatic approach for assessing the quality of functional spaces. Global Ecology and Biogeography 24, 728–740 (2015).

133. Friedman, J. H. Greedy function approximation: a gradient boosting machine. Annals of statistics 1189–1232 (2001).

134. Elith, J., Leathwick, J. R. & Hastie, T. A working guide to boosted regression trees. Journal of Animal Ecology 77, 802–813 (2008).

135. Hastie, T., Tibshirani, R. & Friedman, J. The elements of statistical learning: data mining, inference, and prediction. (Springer Science & Business Media, 2009).

136. Hijmans, R. J., Phillips, S., Leathwick, J. & Elith, J. dismo: Species Distribution Modeling. (2017).

137. Carvalho, J. C., Cardoso, P. & Gomes, P. Determining the relative roles of species replacement and species richness differences in generating beta-diversity patterns. Global Ecology and Biogeography 21, 760–771 (2012).

138. Wickham, H. et al. Welcome to the Tidyverse. Journal of Open Source Software 4, 1686 (2019).

139. Zhu, L. et al. Trait choice profoundly affected the ecological conclusions drawn from functional diversity measures. Scientific reports 7, 3643 (2017).

140. Wong, M. K. L. & Carmona, C. P. Including intraspecific trait variability to avoid distortion of functional diversity and ecological inference: lessons from natural assemblages. bioRxiv 2020.09.17.302349 (2020) doi:10.1101/2020.09.17.302349.

141. Siefert, A., Ravenscroft, C., Weiser, M. D. & Swenson, N. G. Functional beta-diversity patterns reveal deterministic community assembly processes in eastern North American trees. Global Ecology and Biogeography 22, 682–691 (2013).

142. Kruk, C. et al. Phytoplankton community composition can be predicted best in terms of morphological groups. Limnology and Oceanography 56, 110–118 (2011).

143. Soininen, J., Heino, J. & Wang, J. A meta-analysis of nestedness and turnover components of beta diversity across organisms and ecosystems. Global Ecology and Biogeography 27, 96–109 (2018).

144. Stevens, G. C. The Latitudinal Gradient in Geographical Range: How so Many Species Coexist in the Tropics. Source: The American Naturalist vol. 133 (1989).

145. Jamoneau, A., Chabrerie, O., Closset-Kopp, D. & Decocq, G. Fragmentation alters beta- diversity patterns of habitat specialists within forest metacommunities. Ecography 35, 124– 133 (2012).

146. Edie, S. M., Jablonski, D. & Valentine, J. W. Contrasting responses of functional diversity to major losses in taxonomic diversity. Proceedings of the National Academy of Sciences of the United States of America 115, 732–737 (2018).

147. Castro-Insua, A., Gómez-Rodríguez, C. & Baselga, A. Break the pattern: breakpoints in beta diversity of vertebrates are general across clades and suggest common historical causes. Global Ecology and Biogeography 25, 1279–1283 (2016).

148. Zagmajster, M. et al. Geographic variation in range size and beta diversity of groundwater crustaceans: Insights from habitats with low thermal seasonality. Global Ecology and Biogeography 23, 1135–1145 (2014).

149. Bierne, N., Bonhomme, F. & David, P. Habitat preference and the marine-speciation paradox. Proceedings of the Royal Society of London. Series B: Biological Sciences 270, 1399–1406 (2003).

150. Shurin, J. B., Gruner, D. S. & Hillebrand, H. Review All wet or dried up? Real differences between aquatic and terrestrial food webs. Proceedings of the Royal Society B: Biological Sciences 273, 1–9 (2006).

151. Mouquet, N. & Loreau, M. Community Patterns in Source-Sink Metacommunities. The American Naturalist 162, 544–557 (2003).

152. Heim, N. A. et al. Hierarchical complexity and the size limits of life. *Proceedings*. Biological sciences 284, 20171039 (2017).

153. Hodgson, J. G., Wilson, P. J., Hunt, R., Grime, J. P. & Thompson, K. Allocating C-S-R Plant Functional Types: A Soft Approach to a Hard Problem. Oikos 85, 282 (1999).

154. Martínez, A. et al. Habitat differences filter functional diversity of low dispersive microscopic animals 1 2. bioRxiv 2020.09.22.308353 (2020) doi:10.1101/2020.09.22.308353.

155. Stegen, J. C. et al. Quantifying community assembly processes and identifying features that impose them. The ISME journal 7, 2069–2079 (2013).

156. Titley, M. A., Snaddon, J. L. & Turner, E. C. Scientific research on animal biodiversity is systematically biased towards vertebrates and temperate regions. PLOS ONE 12, e0189577 (2017).

