## Supplemental Information 1 for "Distance decay 2.0 – a global synthesis of taxonomic and functional turnover in ecological communities"

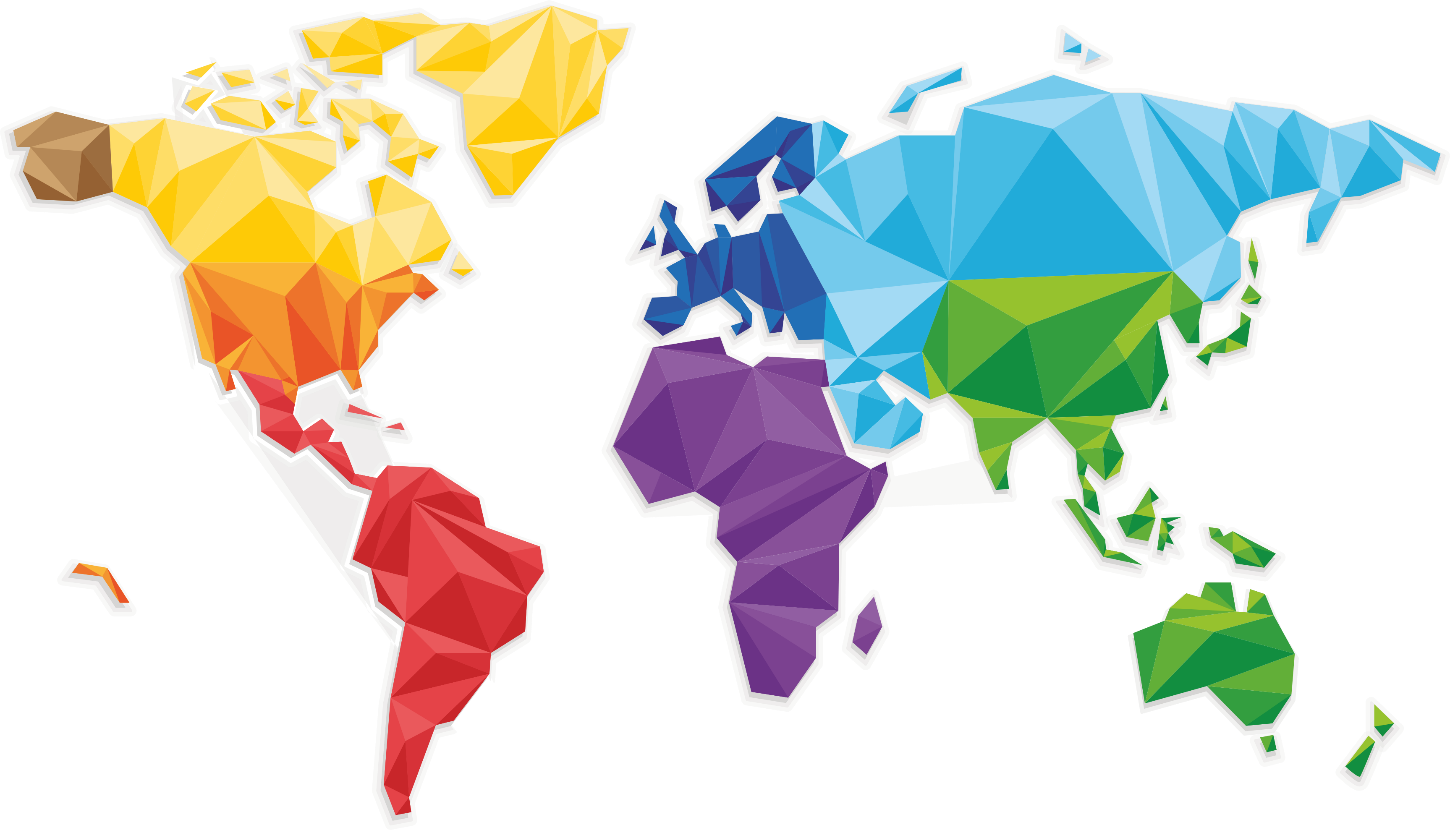

**Distance decay 2.0**

a global synthesis of taxonomic and functional turnover in ecological communities

**Appendix S1:**

Datasets information

I. Table of Contents

The datasets were coded based on their hemisphere (i.e., **N**orth or **S**outh), mean latitude of the sites, realm (i.e., **T**errestrial, **M**arine, and **F**reshwaters), and biotic groups (Table 1). Therefore, a dataset of benthic diatoms from Hawaiian streams would be coded as N21FBD.

Table 1. Abreviations used for each biotic group

| **Biotic group** | **Abbreviation** |
| --- | --- |
| Aquatic Macroinvertebrates | MI |
| Aquatic Vascular Plants | AP |
| Amphibian | AB |
| Arthropods | AT |
| Bats | BT |
| Benthic diatoms | BD |
| Birds | BI |
| Bryophytes | BP |
| Butterflies | BF |
| Fishes | FI |
| Foraminifera | FR |
| Fungi | FG |
| Lichens | LC |
| Marine Ciliates | CI |
| Phytoplankton | PP |
| Reef Corals | CO |
| Spiders | SP |
| Terrestrial vascular plants | TP |

### II. Data compiled directly from data owners

#### 1) N21FBD: Benthic diatoms from Hawaiian streams – *Jenny Jyrkänkallio-Mikkola et al.*

**Data owner(s):** Jenny Jyrkänkallio-Mikkola^1, 2^, Virpi Pajunen^2^, Anette Teittinen^2^ and Janne Soininen^2^.

**Affiliation(s):** ^1^ WWF Finland; ^2^Department of Geosciences and Geography, University of Helsinki, Helsinki, Finland

**e-mail**:.

**Description:** Benthic diatom data and supporting environmental dataset

**Geographical scope:** Hawaiian Islands: Hawaii, Oahu and Kauai

**Taxonomical scope:** Benthic diatoms

**Ecological scope:** Freshwater (Streams)

**Number of sampling units:** 49 sites

**Minimum distance between sampling units:** < 0.01 kilometres

**Maximum distance between sampling units:** 541 kilometres

**Environmental variables:** elevation (m), water temperature (°C), conductivity (mS m^-1^), pH, total phosphorus (µg L^-1^), total nitrogen (µg L^-1^), water colour (Pt mg L^-1^), stream width (m), stream depth (m), current velocity (ms^-1^), shading of the surrounding vegetation (%), percentages of moss, sand, gravel, pebble, cobble, boulders and bedrock.

**Functional traits:** Life‐forms (mobile, adnate, pedunculate, pad-attached, stalk-attached, colonial; Rimet & Bouchez 2012), ecological guilds (high-profile, low-profile, motile; Passy 2007; Rimet & Bouchez 2012)

**Reference:** Unpublished.

#### 2) N21FBD_2: Diatoms from Xishuangbanna, China – Siwen He

**Data owner(s)**: Beixin Wang and Siwen He

**Affiliation(s):** Department of Entomology, College of Plant Protection, Nanjing Agricultural University, Nanjing, China

**Description:** A total of 57 stream and river sites were sampled in March and April 2013.

**Geographical scope:** China

**Taxonomical scope:** Diatoms

**Ecological scope:** Freshwater

**Number of sampling units:** 57 sites in different-sized streams and rivers.

**Minimum distance between sampling units:** 1.3 kilometres

**Maximum distance between sampling units:** 192 kilometres

**Environmental variables:** Water temperature, pH, total dissolved solids, conductivity, calcium concentrations, magnesium concentrations, total nitrogen, total phosphorus, ammonia nitrogen, phosphate contents, potassium permanganate index, elevation, Strahler order.

**Functional traits:** size (biovolume classes: large > 1000µm³ / small < 1000µm³), mobility (mobile / non-mobile), type of attachment (adnate / pedunculate [which was further divided to pad-attached / stalk-attached] / non-attached), colonization (colonial / non-colonial), growth form (low-profile / high-profile / motile / planktonic), and nitrogen-fixing abilities (nitrogen-fixer / non-nitrogen-fixer)

**Reference:** Unpublished

#### 3) N21FMI: Macroinvertebrates from Xishuangbanna, China – Siwen He

**Data owner(s)**: Beixin Wang and Siwen He

**Affiliation(s):** Department of Entomology, College of Plant Protection, Nanjing Agricultural University, Nanjing, China

**Description:** A total of 60 stream and river sites were sampled in March and April 2013

**Geographical scope:** China

**Taxonomical scope:** Macroinvertebrates

**Ecological scope:** Freshwater

**Number of sampling units:** 60 sites in different-sized streams and rivers.

**Minimum distance between sampling units:** 1.30 kilometres

**Maximum distance between sampling units:** 192 kilometres

**Environmental variables:** Water temperature, pH, total dissolved solids, conductivity, calcium concentrations, magnesium concentrations, total nitrogen, total phosphorus, ammonia nitrogen, phosphate contents, potassium permanganate index, elevation, Strahler order

**Functional traits:** 1. Refuge: None, Fixed networks and retreats, Shelters of sand, debris, and/or shells, Shelters of leaf parts or wood

2. Exoskeleton or external protection: Soft body, Lightly sclerotized, Well protected

3. Respiration: Integumentary, Branchial, Air (spiracles, tracheae, plastrons)

4. Body size: Small (<9 mm), Mid (9–16 mm), Large (>16 mm)

5. Body shape: Hydrodynamic (flat, fusiform), Not hydrodynamic (cylindrical, round)

6. Rheophily: Only depositional, Depositional and erosional, Erosional

7. Habit: Burrowers, Climbers, Sprawlers, Clingers, Swimmers, Skaters

8. Functional feeding groups: Collector-gatherers, Collector-filterers, Herbivores (scrapers, leaf miners), Predators (piercers and engulfers), Shredders

**Reference:** Ding, N., Yang, W., Zhou, Y., González-Bergonzoni, I., Zhang, J., Chen, K., *et al.* (2017). Different responses of functional traits and diversity of stream macroinvertebrates to environmental and spatial factors in the Xishuangbanna watershed of the upper Mekong River Basin, China. *Sci. Total Environ.*, 574, 288–299.

#### 4) N25FBD: Diatoms from Li River Basin, China – Siwen He

**Data owner(s)**: Beixin Wang and Siwen He

**Affiliation(s):** Department of Entomology, College of Plant Protection, Nanjing Agricultural University, Nanjing, China

**Description:** A total of 26 stream and river sites were sampled in March 2008

**Geographical scope:** China

**Taxonomical scope:** Diatoms

**Ecological scope:** Freshwater

**Number of sampling units:** 26 sites in different-sized streams and rivers

**Minimum distance between sampling units:** 0.58 kilometres

**Maximum distance between sampling units:** 70 kilometres

**Environmental variables:** Water temperature, pH, total dissolved solids, conductivity, calcium concentrations, magnesium concentrations, total nitrogen, total phosphorus, ammonia nitrogen, phosphate contents, potassium permanganate index, elevation, Strahler order

**Functional traits:** size (biovolume classes: large > 1000µm³ / small < 1000µm³), mobility (mobile / non-mobile), type of attachment (adnate / pedunculate [which was further divided to pad-attached / stalk-attached] / non-attached), colonization (colonial / non-colonial), growth form (low-profile / high-profile / motile / planktonic), and nitrogen-fixing abilities (nitrogen-fixer / non-nitrogen-fixer)

**Reference:** Unpublished

#### 5) N25FMI: Macroinvertebrates from Li River Basi, China – Siwen He

**Data owner(s)**: Beixin Wang and Siwen He

**Affiliation(s):** Department of Entomology, College of Plant Protection, Nanjing Agricultural University, Nanjing, China

**Description:** A total of 34 stream and river sites were sampled in March 2008

**Geographical scope:** China

**Taxonomical scope:** Macroinvertebrates

**Ecological scope:** Freshwater

**Number of sampling units:** 34 sites in different-sized streams and rivers

**Minimum distance between sampling units:** 0.58 kilometres

**Maximum distance between sampling units:** 121 kilometres

**Environmental variables:** Water temperature, pH, total dissolved solids, conductivity, calcium concentrations, magnesium concentrations, total nitrogen, total phosphorus, ammonia nitrogen, phosphate contents, potassium permanganate index, elevation, Strahler order

**Functional traits:** 1. Refuge: None, Fixed networks and retreats, Shelters of sand, debris, and/or shells, Shelters of leaf parts or wood

2. Exoskeleton or external protection: Soft body, Lightly sclerotized, Well protected

3. Respiration: Integumentary, Branchial, Air (spiracles, tracheae, plastrons)

4. Body size: Small (<9 mm), Mid (9–16 mm), Large (>16 mm)

5. Body shape: Hydrodynamic (flat, fusiform), Not hydrodynamic (cylindrical, round)

6. Rheophily: Only depositional, Depositional and erosional, Erosional

7. Habit: Burrowers, Climbers, Sprawlers, Clingers, Swimmers, Skaters

8. Functional feeding groups: Collector-gatherers, Collector-filterers, Herbivores (scrapers, leaf miners), Predators (piercers and engulfers), Shredders

**Reference:** Chen, K., Hughes, R.M., Xu, S., Zhang, J., Cai, D. & Wang, B. (2014). Evaluating performance of macroinvertebrate-based adjusted and unadjusted multi-metric indices (MMI) using multi-season and multi-year samples. *Ecol. Indic.*, 36, 142–151.

#### 6) N26MCI: Coastal benthic ciliates of China – *Yuan XU*

**Data owner(s)**: Yuan XU

**Affiliation(s):** State Key Laboratory of Estuarine and Coastal Research, East China Normal University, 200241, Shanghai, China.

**Description:** The dataset contains 220 ciliate genera collected from 31 sites along the Chinese coast: the data from the coastlines of the Chinese Bohai Sea, Yellow Sea and South China Sea were obtained from the marine ciliate biodiversity survey conducted by the Laboratory of Protozoology, Ocean University of China, from 1991 to 2018 and the habitat types included intertidal sandy beach and mangrove; the data from the Yangtze River Estuary originated from Xu et al. (2018) and the habitat type was salt marsh. Five traits reflecting morphological characteristics (body size, degree of flexibility and body form) and behaviour (feeding and movement type) were sub-divided into 13 categories. Data on traits were mainly obtained from the original sources in which the species were described, as well as from expert opinions and literature. The dataset was published in Xu & Soininen (2019).

**Geographical scope:** China

**Taxonomical scope:** Ciliated protozoa

**Ecological scope:** Marine

**Number of sampling units:** 31 sites

**Minimum distance between sampling units:** 0.70 kilometres

**Maximum distance between sampling units:** 2200 kilometres

**Environmental variables:** Salinity and pH were averaged for each site. Two climatic variables (annual mean temperature and annual precipitation) were obtained from WorldClim at a 30 s resolution. Net primary productivity was extracted from SEDAC at 0.25 decimal degrees.

**Functional traits:**

Five traits were selected and sub-divided into 13 categories according to Xu et al. (2018). These traits reflect morphological characteristics (body size, degree of flexibility and body form) and behavior (feeding and mobility). Data on traits were mainly obtained from the original source in which species were described, as well as from expert’s opinions and literature (Lynn 2008).

| **Trait** | **Category** | **Description** |
| --- | --- | --- |
| Feeding type | Bactivores | Feeding on bacteria |
|  | Algivores | Feeding on algae |
|  | Predators | Feeding on flagellates and ciliates |
| Body size | Small | Cell length < 50 μm |
|  | Medium | 50 μm < Cell length < 200 μm |
|  | Large | 200 μm < Cell length |
| Mobility | Attached to substrate | Self–explanatory |
|  | Swimming | Locomotion by swimming |
|  | Crawling | Locomotion by crawling on substrate |
| Body form | Dorso-ventrally flattened | Ratio of thickness: width < 1: 4 |
|  | Cylindrical | Ratio of thickness: width > 1: 4 |
| Cell flexibility | Cell non-flexible | Cell non-flexible and non–contractile |
|  | Cell flexible | Cell either flexible or contractile or both |

**Reference:** Lynn, D.H. (2010). *The Ciliated Protozoa*. *Ciliated Protozoa Charact. Classif. Guid. to Lit. Third Ed.* Springer Netherlands, Dordrecht.

Xu, Y. & Soininen, J. (2019). Spatial patterns of functional diversity and composition in marine benthic ciliates along the coast of China. *Mar. Ecol. Prog. Ser.*, 627, 49–60.

Xu, Y., Fan, X., Warren, A., Zhang, L. & Xu, H. (2018). Functional diversity of benthic ciliate communities in response to environmental gradients in a wetland of Yangtze Estuary, China. *Mar. Pollut. Bull.*, 127, 726–732.

#### 7) N29FBD: Diatoms from Qiantang River Basin, China – Siwen He

**Data owner(s)**: Beixin Wang and Siwen He

**Affiliation(s):** Department of Entomology, College of Plant Protection, Nanjing Agricultural University, Nanjing, China

**Description:** A total of 82 stream and river sites were sampled in April 2010

**Geographical scope:** China

**Taxonomical scope:** Diatoms

**Ecological scope:** Freshwater

**Number of sampling units:** 82 sites in different-sized streams and rivers

**Minimum distance between sampling units:** 0.55 kilometres

**Maximum distance between sampling units:** 205 kilometres

**Environmental variables:** Water temperature, pH, total dissolved solids, conductivity, calcium concentrations, magnesium concentrations, total nitrogen, total phosphorus, ammonia nitrogen, phosphate contents, potassium permanganate index, elevation, Strahler order

**Functional traits:** size (biovolume classes: large > 1000µm³ / small < 1000µm³), mobility (mobile / non-mobile), type of attachment (adnate / pedunculate [which was further divided to pad-attached / stalk-attached] / non-attached), colonization (colonial / non-colonial), growth form (low-profile / high-profile / motile / planktonic), and nitrogen-fixing abilities (nitrogen-fixer / non-nitrogen-fixer)

**Reference:** Liu, S., Xie, G., Wang, L., Cottenie, K., Liu, D. & Wang, B. (2016). Different roles of environmental variables and spatial factors in structuring stream benthic diatom and macroinvertebrate in Yangtze River Delta, China. *Ecol. Indic.*, 61, 602–611.

#### 8) N29FMI: Macroinvertebrates from Qiantang River Basin, China – Siwen He

**Data owner(s)**: Beixin Wang and Siwen He

**Affiliation(s):** Department of Entomology, College of Plant Protection, Nanjing Agricultural University, Nanjing, China

**Description:** A total of 90 stream and river sites were sampled in April 2010

**Geographical scope:** China

**Taxonomical scope:** Macroinvertebrates

**Ecological scope:** Freshwater

**Number of sampling units:** 90 sites in different-sized streams and rivers

**Minimum distance between sampling units:** 0.55 kilometres

**Maximum distance between sampling units:** 205 kilometres

**Environmental variables:** Water temperature, pH, total dissolved solids, conductivity, calcium concentrations, magnesium concentrations, total nitrogen, total phosphorus, ammonia nitrogen, phosphate contents, potassium permanganate index, elevation, Strahler order

**Functional traits:** 1. Refuge: None, Fixed networks and retreats, Shelters of sand, debris, and/or shells, Shelters of leaf parts or wood

2. Exoskeleton or external protection: Soft body, Lightly sclerotized, Well protected

3. Respiration: Integumentary, Branchial, Air (spiracles, tracheae, plastrons)

4. Body size: Small (<9 mm), Mid (9–16 mm), Large (>16 mm)

5. Body shape: Hydrodynamic (flat, fusiform), Not hydrodynamic (cylindrical, round)

6. Rheophily: Only depositional, Depositional and erosional, Erosional

7. Habit: Burrowers, Climbers, Sprawlers, Clingers, Swimmers, Skaters

8. Functional feeding groups: Collector-gatherers, Collector-filterers, Herbivores (scrapers, leaf miners), Predators (piercers and engulfers), Shredders

**Reference:** Liu, S., Xie, G., Wang, L., Cottenie, K., Liu, D. & Wang, B. (2016). Different roles of environmental variables and spatial factors in structuring stream benthic diatom and macroinvertebrate in Yangtze River Delta, China. *Ecol. Indic.*, 61, 602–611.

#### 9) N34FBD: Diatoms from Wei River Basin, China – Siwen He

**Data owner(s)**: Beixin Wang and Siwen He

**Affiliation(s):** Department of Entomology, College of Plant Protection, Nanjing Agricultural University, Nanjing, China

**Description:** A total of 57 stream and river sites were sampled in June 2013

**Geographical scope:** China

**Taxonomical scope:** Diatoms

**Ecological scope:** Freshwater

**Number of sampling units:** 57 sites in different-sized streams and rivers

**Minimum distance between sampling units:** < 0.01 kilometres

**Maximum distance between sampling units:** 164 kilometres

**Environmental variables:** Water temperature, pH, total dissolved solids, conductivity, calcium concentrations, magnesium concentrations, total nitrogen, total phosphorus, ammonia nitrogen, phosphate contents, potassium permanganate index, elevation, Strahler order

**Functional traits:** size (biovolume classes: large > 1000µm³ / small < 1000µm³), mobility (mobile / non-mobile), type of attachment (adnate / pedunculate [which was further divided to pad-attached / stalk-attached] / non-attached), colonization (colonial / non-colonial), growth form (low-profile / high-profile / motile / planktonic), and nitrogen-fixing abilities (nitrogen-fixer / non-nitrogen-fixer)

**Reference:** Unpublished

#### 10) N34FMI: Macroinvertebrates from Wei River Basin, China – Siwen He

**Data owner(s)**: Beixin Wang and Siwen He

**Affiliation(s):** Department of Entomology, College of Plant Protection, Nanjing Agricultural University, Nanjing, China

**Description:** A total of 57 stream and river sites were sampled in June 2013

**Geographical scope:** China

**Taxonomical scope:** Macroinvertebrates

**Ecological scope:** Freshwater

**Number of sampling units:** 57 sites in different-sized streams and rivers

**Minimum distance between sampling units:** < 0.01 kilometres

**Maximum distance between sampling units:** 164 kilometres

**Environmental variables:** Water temperature, pH, total dissolved solids, conductivity, calcium concentrations, magnesium concentrations, total nitrogen, total phosphorus, ammonia nitrogen, phosphate contents, potassium permanganate index, elevation, Strahler order

**Functional traits:** 1. Refuge: None, Fixed networks and retreats, Shelters of sand, debris, and/or shells, Shelters of leaf parts or wood

2. Exoskeleton or external protection: Soft body, Lightly sclerotized, Well protected

3. Respiration: Integumentary, Branchial, Air (spiracles, tracheae, plastrons)

4. Body size: Small (<9 mm), Mid (9–16 mm), Large (>16 mm)

5. Body shape: Hydrodynamic (flat, fusiform), Not hydrodynamic (cylindrical, round)

6. Rheophily: Only depositional, Depositional and erosional, Erosional

7. Habit: Burrowers, Climbers, Sprawlers, Clingers, Swimmers, Skaters

8. Functional feeding groups: Collector-gatherers, Collector-filterers, Herbivores (scrapers, leaf miners), Predators (piercers and engulfers), Shredders

**Reference:** Li, S., Yang, W., Wang, L., Chen, K., Xu, S. & Wang, B. (2018). Influences of environmental factors on macroinvertebrate assemblages: differences between mountain and lowland ecoregions, Wei River, China. *Environ. Monit. Assess.*, 190, 152.

#### 11) N38MFI: Mediterranean Fish – Johnatan Belmaker and Shira Salingré

**Data owner(s)**: Jonathan Belmaker and Shira Salingré

**Affiliation(s):** School of Zoology, George S. Wise Faculty of Life Sciences, Tel Aviv University, Tel Aviv, Israel

**Description:** Underwater fish census surveys in the Mediterranean Sea. Environmental data extracted from Bio-ORACLE.

**Geographical scope:** Mediterranean Sea

**Taxonomical scope:** Fish

**Ecological scope:** Marine (Sea)

**Number of sampling units:** 51 (125 m^2^)

**Minimum distance between sampling units:** < 0.01 kilometres

**Maximum distance between sampling units:** 2997 kilometres

**Environmental variables:** Mean annual temperature, Temperature range, Mean salinity, Mean primary productivity, Primary productivity range

**Functional traits:** R Dataframe *sp_traits* with 6 traits: Size, Mobility, Activity, Schooling, Position, Diet.

- *Size* coded as an ordered categorical variable with 6 levels:

1= 0-7cm (S1)

2= 7.1-15cm (S2)

3= 15.1- 30cm (S3)

4= 30.1-50cm (S4)

5= 50.1-80cm (S5)

6= >80cm (S6)

- *Mobility* (=Home range) coded as an ordered categorical variable, 3 levels:

1= sedentary (Sed)

2= mobile within a reef (Mob)

3= highly mobile i.e. between reefs (VMob)

- Period of *Activity* coded as an ordered categorical variable, 3 levels:

1= diurnal (Day)

2= diurnal & nocturnal (Both)

3= nocturnal (Night)

- *Schooling* coded as a categorical variable, 5 levels:

1= solitary (Sol)

2= pairing (Pair)

3= small group (SmallG)

4= medium group (MedG)

5= large group (LargeG)

- *Position* in the water column (=Level-water) coded as an ordered categorical variable, 3 levels:

1= bottom (Bottom)

2= above bottom (Low)

3= pelagic (High)

*- Diet* coded as a categorical variable, 7 levels:

HD= herbivorous-detritivorous (undefined organic material, often grouped by many authors under the name detritus and/or undefined vegetal material, turf or filamentous algae)

HM= herbivorous macro-algal (macro algae (large fleshy algae) and sea grass)

IS= invertivorous sessile (sessile invertebrates: coral, sponge, ascidians and so on)

IM= invertivorous mobile (large benthic invertebrates + small benthic invertebrates + undefined invertebrates)

PK= planktonivorous (plankton and small organisms which migrate in the water column, such as many benthic copepods, amphipods, crustacean larvaes etc which migrate in the water column at night)

FC= Pelagic macro-organisms (large organisms living in the water column, usually fish and cephalopods) and benthic fish

OM= omnivorous (herbivorous and/or detritivorous AND carnivorous)

**Reference:** Unpublished.

#### 12) N39FMI: Aquatic beetles from lentic systems across mainland Spain – *Félix Picazo, Andrés Millán and José Luis Moreno*

**Data owner(s)**: Félix Picazo^1,2^ and Andrés Millán^3^ and José Luis Moreno^4^

**Affiliation(s):** ^1^Pyrenean Institute of Ecology-CSIC, ^2^Nanjing Institute of Geography and Limnology-CAS, ^3^University of Murcia, ^4^University of Castilla-La Mancha

**Description:** Dataset comprising limnological variables, abundance classes (0, absence; 1, 1 individual; 2, 2-4 individuals; 3, 5-10 individuals; 4, >10 individuals) and traits for 155 species of water beetles inhabiting lentic systems from Spain.

**Geographical scope:** Mainland Spain

**Taxonomical scope:** Aquatic Coleoptera

**Ecological scope:** Freshwater (lentic systems).

**Number of sampling units:** 102

**Minimum distance between sampling units:** 0.18 kilometres

**Maximum distance between sampling units:** Latitudinal extent, 750 km; Longitudinal extent, 600 km.

**Environmental variables:** Mean annual temperature for period 1960-1990, mean annual precipitation for period 1960-1990, conductivity.

**Functional traits:**

1. The minimal and maximal size of the species
2. The main and secondary type of locomotion or relationship with the substrate/environment: crawler, flier, interstitial, surface swimmer, diving swimmer.
3. The main and secondary source of food: fine sediment, microorganisms and detritus < 1 mm; plant detritus > 1 mm; living microphytes; living microinvertebrates; living macrophytes; living macroinvertebrates; vertebrates.
4. Main and secondary feed habit: piercer, predator, scraper, shredder.

**Reference:** Unpublished as a dataset itself, partially included in:

Picazo, F., Bilton, D.T., Moreno, J.L., Sánchez-Fernández, D. & Millán, A. (2012). Water beetle biodiversity in Mediterranean standing waters: assemblage composition, environmental drivers and nestedness patterns. *Insect Conserv. Divers.*, 5, 146–158.

Sánchez-Fernández, D., Millán, A., Abellán, P., Picazo, F., Carbonell, J.A. & Ribera, I. (2015). Atlas of Iberian water beetles (ESACIB database). *Zookeys*, 520, 147–154.

#### 13) N39FMI_2: Trichoptera Iberian Mediterranean Rivers – Núria Bonada

**Data owner(s)**: Núria Bonada

**Affiliation(s):** University of Barcelona

**Description:** Dataset comprising 36 lotic sites; 62 species; abundance class data (0, absence; 1, 1-3 individuals; 2, 4-10 individuals; 3, 10-100 individuals; 4, >100 individuals); traits: maximal size, locomotion, food source, feeding habit.

**Geographical scope:** Mediterranean Iberia.

**Taxonomical scope:** Trichoptera (larvae)

**Ecological scope:** Freshwater (lotic systems)

**Number of sampling units:** 36

**Minimum distance between sampling units:** 1.21 kilometres

**Maximum distance between sampling units:** Latitudinal extent, 610 km; Longitudinal extent, 450 km.

**Environmental variables:** Mean annual temperature for period 1960-1990, mean annual precipitation for period 1960-1990, conductivity.

**Functional traits:**

1. Maximal size of the the species
2. The main and secondary type of locomotion or relationship with the substrate/environment (i.e., crawler, temporarily attached, burrower, diving swimmer).
3. The main and secondary source of food (i.e., detritus < 1 mm; dead plants > 1mm; dead animals > 1mm; living microphytes; living microinvertebrates; living macrophytes; living macroinvertebrates;)
4. Main and secondary feed habit (i.e., filter-feeder, piercer, predator, scraper, shredder)

**Reference:** Bonada, N., Zamora-Muñoz, C., Rieradevall, M. & Prat, N. (2004). Trichoptera (Insecta) collected in Mediterranean river basins of the Iberian Peninsula: taxonomic remarks and notes on ecology. *Graellsia*, 60, 41–69.

#### 14) N40FMI: Aquatic beetles from lotic systems across central Iberia – *Félix Picazo, Andrés Millán and José Luis Moreno*

**Data owner(s)**: Félix Picazo^1,2^ and Andrés Millán^3^ and José Luis Moreno^4^

**Affiliation(s):** ^1^Pyrenean Institute of Ecology-CSIC, ^2^Nanjing Institute of Geography and Limnology-CAS, ^3^University of Murcia, ^4^University of Castilla-La Mancha

**Description:** Dataset comprising limnological variables, abundances and traits for 157 species of water beetles from lotic systems across central Spain

**Geographical scope:** Central Iberia

**Taxonomical scope:** Aquatic Coleoptera

**Ecological scope:** Freshwater (Lotic systems)

**Number of sampling units:** 64

**Minimum distance between sampling units:** 6.11 kilometres

**Maximum distance between sampling units:** 353 kilometres

**Environmental variables:** Mean annual temperature for period 1960-1990, mean annual precipitation for period 1960-1990, oxygen concentration, oxygen percentage, water temperature, conductivity, alkalinity, flow volume, water hardness, ionic concentration, total suspended solids, total organic carbon, turbidity.

**Functional traits:**

1. The minimal and maximal size of the species
2. The main and secondary type of locomotion or relationship with the substrate/environment (i.e., crawler, flier, interstitial, diving swimmer, surface swimmer)
3. The main and secondary source of food (i.e., fine sediment, microorganisms and detritus < 1 mm; plant detritus > 1 mm; living microphytes; living microinvertebrates; living macrophytes; living macroinvertebrates; vertebrates)
4. Main and secondary feed habit (i.e., piercer, predator, scraper, shredder)

**Reference:** Unpublished as a dataset itself, partially included in:

Millán, A., Sánchez-Fernández, D., Abellán, P., Picazo, F., Carbonell, J.A., Lobo, J.M., *et al.* (2014). Atlas de los coleópteros acuáticos de España peninsular.

Sánchez-Fernández, D., Millán, A., Abellán, P., Picazo, F., Carbonell, J.A. & Ribera, I. (2015). Atlas of Iberian water beetles (ESACIB database). *Zookeys*, 520, 147–154.

#### 15) N43MMI: Benthic macrofauna from Spanish sandy beaches – *Ivan F. Rodil and Alf Norkko*

**Data owner(s)**: Ivan F. Rodil and Alf Norkko

**Affiliation(s):** Tvärminne Zoological Station, University of Helsinki, Helsinki, Finland.

**Description:** Benthic macrofauna data from **39** exposed sandy beaches along the entire north coast of Spain (more than 2000 km of uninterrupted coastline). At each beach three replicate-transects including 10 intertidal levels (from the supratidal to the swash level) were sampled with benthic cores.

**Geographical scope:** Spain

**Taxonomical scope:** Benthic invertebrate species from terrestrial origin (e.g. insect larvae), semiterrestrial (e.g. sandhoppers) and truly marine species (e.g. bivalve *Donax trunculus*).

**Ecological scope:** Marine (Sandy beaches)

**Number of sampling units:** 39 cores (0.05 m^2^)

**Minimum distance between sampling units:** 4.90 kilometres

**Maximum distance between sampling units:** 610 kilometres

**Environmental variables:**

1. Chlorophyll a: chla in the nearshore water column [mg m^-3^]
2. Sstmax: maximum sea surface temperature [ ºC]
3. Sstmean: mean sea surface temperature [ ºC]
4. Length: Beach distance along-shore. [meters]
5. Width: Beach distance across-shore (from above the drift line to the low swash zone) [meters]
6. Slope: This was determined by Emery’s profiling technique and calculated dividing the beach height by the distance between two points. [dimensionless]
7. Mean grain size: Sediment grain size using a Coulter LS 200 laser diffraction particle size analyser and the coarser fraction (> 2 mm) by dry sieving [μm]
8. Water: Sediment water content (% of sediment water at each beach level). [%]
9. Shear strength: Measure sediment compacting force using a hand held shear vane tester (Pilcon) [Kilopascal]
10. Wave height [meters]
11. Wave period [seconds]
12. Dean: Dean’s index was used to characterize the beach morphodynamic type (Ω = (Hb/Ws)* Tb, where Hb is breaker height (m), Ws is sand fall velocity (m s-1) and Tb is wave period (s).
13. Beach State Index: BSI is Ω multiplied by tide range and indicates the ability of waves and tides to move sand. < 0.5 reflective, 0.5-1 low to medium energy intermediate, 1.0-1.5 high-energy intermediate dissipative, 1.5-2.0 dissipative, and > 2.0 ultra-dissipative macrotidal beaches.
14. Relative tide range: RTR = TR/Hb, where TR is spring tide range and Hb is breaker height, both in m
15. Exposure rate: The 20-point rating system is used to estimate the wave exposure rate, which takes into account observations of wave action, median particle diameter, slope, depth of the redox potential layer and the presence of macrofaunal structures (0 is the less exposed situation).

**Functional traits:** feeding type, degree of mobility, developmental mechanism (larvae) and environmental position

**Reference:** Rodil, I.F., Lucena-Moya, P. & Lastra, M. (2018). The Importance of Environmental and Spatial Factors in the Metacommunity Dynamics of Exposed Sandy Beach Benthic Invertebrates. *Estuaries and Coasts*, 41, 206–217.

Rodil, I.F., Compton, T.J. & Lastra, M. (2014). Geographic variation in sandy beach macrofauna community and functional traits. *Estuar. Coast. Shelf Sci.*, 150, 102–110.

#### 16) N45FFI: Fish fauna of Lapland, Finland – Kimmo Kahilainen and Brian Hayden

**Data owner(s)**: Kimmo Kahilainen and Brian Hayden

**Affiliation(s):** ^1^University of Helsinki, Lammi Biological Station

^2^ University of New Brunswick, Rivers Institute

**Description: F**reshwater fish data contain 33 subarctic lakes with environmental variables, relative fish abundance from multimesh gill nets and ecological traits.

**Geographical scope:** Finland (Subartic)

**Taxonomical scope:** Fish

**Ecological scope:** Freshwater (Lakes)

**Number of sampling units:** 33

**Minimum distance between sampling units:** 0.67 kilometres

**Maximum distance between sampling units:** 370 kilometres

**Environmental variables:** Altitude (m a.s.l), Total nitrogen (µg/L), Total phosphorus (µg/L), Compensation depth (m), Maximum depth (m), Mean depth (m), Littoral proportion (%), Lake area (km^2^)

**Functional traits:**

Origin (NativeSpecies, IntroducedSpecies), Major habitat (Lentic, Lotic, Anadromous, Catadromous), thermal-guild (Warm-waterSpecies, Cool-waterSpecies, Cold-waterSpecies), foraging habitat (LittoralSpecies, PelagicSpecies, ProfundalSpecies), foraging guild (Benthivore, Planktivore, Piscivore, Omnivore), spawning site (LakeSpawningSpecies, RiverSpawningSpecies, SeaSpawningSpecies), spawning time (Winter spawning species, Spring Spawning species, AutumnSpawningSpecies), diel activity time (DayActiveSpecies, NightActiveSpecies).

**Reference:** Hayden, B., Myllykangas, J.-P., Rolls, R.J. & Kahilainen, K.K. (2017). Climate and productivity shape fish and invertebrate community structure in subarctic lakes. *Freshw. Biol.*, 62, 990–1003.

Hayden, B., Harrod, C., Thomas, S.M., Eloranta, A.P., Myllykangas, J. ‐P., Siwertsson, A., *et al.* (2019). From clear lakes to murky waters – tracing the functional response of high‐latitude lake communities to concurrent ‘greening’ and ‘browning.’ *Ecol. Lett.*, 22, 807–816.

#### 17) N46FMI: Water beetles from a latitudinal transect across the Western Palaearctic – *Félix Picazo, Andrés Millán and Ignacio Ribera*

**Data owner(s)**: Félix Picazo^1,2^ and Andrés Millán^3^ and Ignacio Ribera^4^

**Affiliation(s):** ^1^Pyrenean Institute of Ecology-CSIC, ^2^Nanjing Institute of Geography and Limnology-CAS, ^3^University of Murcia, ^4^Institute of Evolutionary Biology-CSIC.

**Description:** Dataset comprising climatological and limnological variables, incidence and traits for 512 species of water beetles from both lotic and lentic systems.

**Geographical scope:** Latitudinal transect from Morocco to Sweden.

**Taxonomical scope:** Aquatic Coleoptera

**Ecological scope:** Freshwater (Lotic and Lentic systems)

**Number of sampling units:** 133 lotic sites and 117 lentic sites

**Minimum distance between sampling units:** < 0.01 kilometres

**Maximum distance between sampling units:** 3950 kilometres

**Environmental variables:** Mean annual temperature for period 1960-1990, mean annual precipitation for period 1960-1990

**Functional traits:**

1. Maximal size of the species.
2. The main and secondary type of locomotion or relationship with the substrate/environment (i.e., crawler, flier, interstitial, diving swimmer, surface swimmer)
3. The main and secondary source of food (i.e., fine sediment, microorganisms and detritus < 1 mm; plant detritus > 1 mm; living microphytes; living microinvertebrates; living macrophytes; living macroinvertebrates; vertebrates)
4. Main and secondary feed habit (i.e., piercer, predator, scraper, shredder)

**Reference:** Unpublished.

#### 18) N47FAP: Macrophytes from Hungarian lakes – *Balázs A. Lukács and Janne Alahuhta.*

**Data owner(s)**: Balázs A. Lukács^1^ and Janne Alahuhta^2^

**Affiliation(s):** ^1^ Wetland Ecology Research Group, Centre for Ecological Research, Bem tér 18/C, Debrecen 4026, Hungary.

^2^ Geography Research Unit, University of Oulu, P.O. Box 3000, 90014 Oulu, Finland.

**Description:** Lakes surveyed between June and September over the period 2004-2012 (Lukács et al. 2015).

**Geographical scope:** Hungary

**Taxonomical scope:** Plant (Macrophyte)

**Ecological scope:** Freshwater (Lakes)

**Number of sampling units:** 26

**Minimum distance between sampling units:** 2.08 kilometres

**Maximum distance between sampling units:** 301 kilometres

**Environmental variables:** Multiple samples of pH, TN (μg/L) and TP (μg/L) collected between 2006-2012, complemented with lake area (km^2^), lake depth (m) and altitude (m.a.s.l.).

**Functional traits:** Life form (submerged, helophyte, floating-leaved, free-floating), normal method of propagation (by seed/spore, mostly by seed/spore but also vegetatively, by seed/spore and vegetatively, mostly vegetatively and also by seed/spore and vegetatively; Klotz et al. 2002), maximum height (based on Mossberg and Stenberg 2012) and perennation (annual or perennial; Willby et al. 2000).

**Reference:** Lukács, B.A., Tóthmérész, B., Borics, G., Várbíró, G., Juhász, P., Kiss, B., *et al.* (2015). Macrophyte diversity of lakes in the Pannon Ecoregion (Hungary). *Limnologica*, 53, 74–83.

Klotz, S., Kühn, I. & Durka, W. (2002). BIOLFLOR - Eine Datenbank mit biologisch-ökologischen Merkmalen zur Flora von Deutschland. *Schriftenr. für Veg.*

Lindholm, M., Alahuhta, J., Heino, J. & Toivonen, H. (2020). No biotic homogenisation across decades but consistent effects of landscape position and pH on macrophyte communities in boreal lakes. *Ecography (Cop.).*, 43, 294–305.

Mossberg, B., Stenberg, L., Vuokko, S. & Väre, H. (2005). *Suuri pohjolan kasvio*. Tammi.

Willby, N. J. et al. 2000. Attribute-based classification of European hydrophytes and its relationship to habitat utilization. *Freshwater Biology* 43: 43–74.

#### 19) N47FFI: Hungarian Fish fauna – *Tibor Erös*

**Data owner(s)**: Tibor Erös

**Affiliation(s):** Centre for Ecological Research, Balaton Limnological Institute

**Description:** Fish assemblage, trait and environmental data for 51 stream and river sites

**Geographical scope:** Hungary, Pannon Biogeographic Region

**Taxonomical scope:** Fish

**Ecological scope:** Freshwater

**Number of sampling units:** 51 sites - At each site 150 and 500 m long sections were examined in streams and rivers, respectively.

**Minimum distance between sampling units:** 1.84 kilometres

**Maximum distance between sampling units:** 516 kilometres

**Environmental variables:** Basic environmental physical, hydromorphological and chemical variables as follows: altitude (m), bank vegetation (%), wetted width (0.1 m), mean depth (cm), mean current velocity (cm/sec), substrate composition (%), macrophyte composition (%), temperature (C), pH, oxygen content (mg/l), conductivity (microS/sec), nitrite (micro g/l), nitrate (mg/l), ammónium (mg/l), calcium (mg/l), phosphor (micro g/l), phosphate (mg/l), For details, see the cited article and the data set.

**Functional traits:** feeding group (HER: herbivore, OMN: omnivore, PLA: planktivore, INV_PIS: invertivore-piscivore, INV_BEN: benthic invertivore, PIS: piscivore), habitat (PEL: pelagic, MET: metaphytic, BEN: benthic), velocity preference (RHE: rheophil, EU: eurytop, STA: stagnophil), spawning guild (LIT: lithophil, PHY: phytophil, PHY_LIT: phyto-litophyl, PSA: psammophil, OST: ostracophil, LIT_PEL: litho-pelagophil, SPE: speleophil), native/nonnative status (N: native, NN: non-native), life history (agemat: age at maturity, maxage: maximum age, maxlength: maximum length, lengthmat: length at maturity, maxfecund: maximum fecundity, meanfecun: mean fecundity, meaneggsize: mean egg size, maxeggsize: maximum egg size, larval_development, larvasize: length of the larvae at hatching, number_of_ spawnings, parecare: parental care)

**Reference:** Erős, T., Takács, P., Specziár, A., Schmera, D. & Sály, P. (2017). Effect of landscape context on fish metacommunity structuring in stream networks. *Freshw. Biol.*, 62, 215–228.

Erős, T., Sály, P., Takács, P., Specziár, A. & Bíró, P. (2012). Temporal variability in the spatial and environmental determinants of functional metacommunity organization - stream fish in a human-modified landscape. *Freshw. Biol.*, 57, 1914–1928.

Eros, T. (2005). Life-history diversification in the Middle Danubian fish fauna - a conservation perspective. *River Syst.*, 16, 289–304.

#### 20) N48FBD: Diatoms from Irtysh River Basin, China – Siwen He

**Data owner(s)**: Beixin Wang and Siwen He

**Affiliation(s):** Department of Entomology, College of Plant Protection, Nanjing Agricultural University, Nanjing, China

**Description:** A total of 35 stream and river sites were sampled in June and July 2013

**Geographical scope:** China

**Taxonomical scope:** Diatoms

**Ecological scope:** Freshwater

**Number of sampling units:** 35 sites in different-sized streams and rivers

**Minimum distance between sampling units:** 0.17 kilometres

**Maximum distance between sampling units:** 393 kilometres

**Environmental variables:** Water temperature, pH, total dissolved solids, conductivity, calcium concentrations, magnesium concentrations, total nitrogen, total phosphorus, ammonia nitrogen, phosphate contents, potassium permanganate index, elevation, Strahler order

**Functional traits:** size (biovolume classes: large > 1000µm³ / small < 1000µm³), mobility (mobile / non-mobile), type of attachment (adnate / pedunculate [which was further divided to pad-attached / stalk-attached] / non-attached), colonization (colonial / non-colonial), growth form (low-profile / high-profile / motile / planktonic), and nitrogen-fixing abilities (nitrogen-fixer / non-nitrogen-fixer)

**Reference:** Chen, K., Sun, D., Rajper, A.R., Mulatibieke, M., Hughes, R.M., Pan, Y., *et al.* (2019). Concordance in biological condition and biodiversity between diatom and macroinvertebrate assemblages in Chinese arid-zone streams. *Hydrobiologia*, 829, 245–263.

#### 21) N48FMI: Macroinvertebrates from Irtysh River Basin, China – Siwen He

**Data owner(s)**: Beixin Wang and Siwen He

**Affiliation(s):** Department of Entomology, College of Plant Protection, Nanjing Agricultural University, Nanjing, China

**Description:** A total of 41 stream and river sites were sampled in June and July 2013

**Geographical scope:** China

**Taxonomical scope:** Macroinvertebrates

**Ecological scope:** Freshwater

**Number of sampling units:** 41 sites in different-sized streams and rivers

**Minimum distance between sampling units:** 0.17 kilometres

**Maximum distance between sampling units:** 393 kilometres

**Environmental variables:** Water temperature, pH, total dissolved solids, conductivity, calcium concentrations, magnesium concentrations, total nitrogen, total phosphorus, ammonia nitrogen, phosphate contents, potassium permanganate index, elevation, Strahler order

**Functional traits:** 1. Refuge: None, Fixed networks and retreats, Shelters of sand, debris, and/or shells, Shelters of leaf parts or wood

2. Exoskeleton or external protection: Soft body, Lightly sclerotized, Well protected

3. Respiration: Integumentary, Branchial, Air (spiracles, tracheae, plastrons)

4. Body size: Small (<9 mm), Mid (9–16 mm), Large (>16 mm)

5. Body shape: Hydrodynamic (flat, fusiform), Not hydrodynamic (cylindrical, round)

6. Rheophily: Only depositional, Depositional and erosional, Erosional

7. Habit: Burrowers, Climbers, Sprawlers, Clingers, Swimmers, Skaters

8. Functional feeding groups: Collector-gatherers, Collector-filterers, Herbivores (scrapers, leaf miners), Predators (piercers and engulfers), Shredders

**Reference:** Chen, K., Sun, D., Rajper, A.R., Mulatibieke, M., Hughes, R.M., Pan, Y., *et al.* (2019). Concordance in biological condition and biodiversity between diatom and macroinvertebrate assemblages in Chinese arid-zone streams. *Hydrobiologia*, 829, 245–263.

#### 22) N51TAT (6-8): Arthropods from the Forest sites of the Biodiversity Exploratories, Germany – *Martin M. Gossner and Wolfgang W. Weisser*

**Data owner(s)**: Martin M. Gossner^1^ and Wolfgang W. Weisser2

**Affiliation(s):** ^1^ Forest Entomology, Swiss Federal Research Institute WSL, Birmensdorf CH-8903, Switzerland; ^2^ Terrestrial Ecology Research Group, Department of Ecology and Ecosystem Management, Technical University of Munich, Hans-Carl-von-Carlowitz-Platz 2, 85354 Freising, Germany

**Description:** Activity densities of arthropods sampled by pitfall traps and flight-interception traps in the understorey and canopy on 150 sites within the long-term and large-scale functional biodiversity project “Biodiversity Exploratories” in 2008

**Geographical scope:** Germany, Biosphere Reserves Schwäbische Alb & Schorfheide-Chorin, National Park Hainich and surrounding Hainich-Dün”

**Taxonomical scope:** Arthropods

6. Araneae
7. Coleoptera
8. Hemiptera

**Ecological scope:** Terrestrial (Forest)

**Number of sampling units:** 150 plots (100m x 100m)

**Minimum distance between sampling units:** 0.2 kilometres

**Maximum distance between sampling units:** 630 kilometres

**Environmental variables:** Elevation, main tree species of forest stand, forest management intensity (Schall & Amer 2013), deadwood volume [m3/ha], Shannon Diversity of Plants, Percentage of conifer tree in the tree layer, canopy closure.

**Functional traits:** Based on Gossner *et al.* 2015

1. Feeding guild: herbivore, carnivore, detritivore, fungivore, omnivore
2. Stratum use: ground dwelling, herb layer, tree and shrub layer, soil dwelling, water, indifferent.
3. Dispersal ability (based on wing development across sexes in insects and based on ballooning and migration behaviour of spiders): 0 – low; 0.25 – low to intermediate; 0.5 – intermediate; 0.75 – intermediate to high; 1 – high

**Reference:** Gossner, M.M., Simons, N.K., Achtziger, R., Blick, T., Dorow, W.H.O., Dziock, F., *et al.* (2015). A summary of eight traits of Coleoptera, Hemiptera, Orthoptera and Araneae, occurring in grasslands in Germany. *Sci. Data*, 2, 150013.

Schall, P. & Ammer, C. (2013). How to quantify forest management intensity in Central European forests. *Eur. J. For. Res.*, 132, 379–396.

Lange, M., Türke, M., Pašalić, E., Boch, S., Hessenmöller, D., Müller, J., *et al.* (2014). Effects of forest management on ground-dwelling beetles (Coleoptera; Carabidae, Staphylinidae) in Central Europe are mainly mediated by changes in forest structure. *For. Ecol. Manage.*, 329, 166–176.

Schall, P., Gossner, M.M., Heinrichs, S., Fischer, M., Boch, S., Prati, D., *et al.* (2018). The impact of even-aged and uneven-aged forest management on regional biodiversity of multiple taxa in European beech forests. *J. Appl. Ecol.*, 55, 267–278.

Penone, C., Allan, E., Soliveres, S., Felipe‐Lucia, M.R., Gossner, M.M., Seibold, S., *et al.* (2019). Specialisation and diversity of multiple trophic groups are promoted by different forest features. *Ecol. Lett.*, 22, 170–180.

#### 23) N50TAT (1-2): Ground-dwelling arthropods from the Grassland sites of the Biodiversity Exploratories, Germany – *Martin M. Gossner and Wolfgang W. Weisser*

**Data owner(s)**: Martin M. Gossner^1^ and Wolfgang W. Weisser2

**Affiliation(s):** ^1^ Forest Entomology, Swiss Federal Research Institute WSL, Birmensdorf CH-8903, Switzerland 2Terrestrial Ecology Research Group, Department of Ecology and Ecosystem Management, Technical University of Munich, Hans-Carl-von-Carlowitz-Platz 2, 85354 Freising, Germany

**Description:** Activity densities of arthropods sampled by pitfall traps on 43 sites within the long-term and large-scale functional biodiversity project “Biodiversity Exploratories” in 2008

**Geographical scope:** Germany, National Park Hainich and surrounding Hainich-Dün region

**Taxonomical scope:** Arthropods:

1. Aranea
2. Coleoptera

**Ecological scope:** Terrestrial (Grasslands)

**Number of sampling units:** 43 plots (50m x 50m)

**Minimum distance between sampling units:** 0.2 kilometres

**Maximum distance between sampling units:** 37 kilometres

**Environmental variables:** Perimeter-area ratio (mean) of land cover types in a 500m radius around plots, Shannon diversity of land cover types in a 500m radius around plots, Proportion of grasslands in a 500m radius around plots, Elevation, Soil depth [cm], Soil Bulk Density [g/cm3] , Soil pH, Soil Texture (PCA axis), Soil Nutrients (PCA-axis), Land-use intensity combining mowing, grazing and fertilization between 2006 and 2008 following Blüthgen et al 2012

**Functional traits:** Based on the literature:

1. Ecological type: according to Platen et al. (1991, 1999), Malten (1999, 2001), Malten & Blick (2007), Blick (2009)
2. Activity type: according to Platen *et al.* (1991), Malten (1999, 2001), Malten & Blick (2007), Blick (2009), adapted by authors
3. Guild: according to Cardoso *et al.* 2011
4. Feeding guild: herbivore, carnivore, detritivore, fungivore, omnivore
5. Stratum use: ground dwelling, herb layer, tree and shrub layer, soil dwelling, water, indifferent. (Gossner *et al.* 2015)
6. Dispersal ability (based on wing development across sexes in Carabidae and based on ballooning and migration behaviour of spiders): 0 – low; 0.25 – low to intermediate; 0.5 – intermediate; 0.75 – intermediate to high; 1 – high; (Gossner *et al.* 2015)

**Reference:** Blick, T. (2009) Die Spinnen (Araneae) des Naturwaldreservats Goldbachs- und Ziebachsrück (Hessen). Untersuchungszeitraum 1994–1996. *Mitt. Hess. Landesforstverw*. 45, 57–138 (2009).

Blüthgen, N., Dormann, C.F., Prati, D., Klaus, V.H., Kleinebecker, T., Hölzel, N., *et al.* (2012). A quantitative index of land-use intensity in grasslands: Integrating mowing, grazing and fertilization. *Basic Appl. Ecol.*, 13, 207–220.

Cardoso, P., Pekár, S., Jocqué, R. & Coddington, J.A. (2011). Global Patterns of Guild Composition and Functional Diversity of Spiders. *PLoS One*, 6, e21710.

Gossner, M.M., Simons, N.K., Achtziger, R., Blick, T., Dorow, W.H.O., Dziock, F., *et al.* (2015). A summary of eight traits of Coleoptera, Hemiptera, Orthoptera and Araneae, occurring in grasslands in Germany. *Sci. Data*, 2, 150013.

Malten, A. & Blick, T. (2007) Araneae (Spinnen). in Naturwaldreservate in Hessen. Band 7/2.2. Hohestein. Zoologische Untersuchungen 1994-1996 (eds Dorow W. H. O., Kopelke J.-P.). *Mitt. Hess. Landesforstverw*. 42, 7–93.

Malten, A. & Blick, T. (2007). 3.5 Araneae (Spinnen). *Hohestein–Zoologische Untersuchungen*, 2, 341.

Malten, A. (2004). Araneae (Spinnen) & Opiliones (Weberknechte). *Schön buche*, 30.

Platen, R., Broen, B., Herrmann, A., Ratschker, U. M. & Sacher, P. (1999) Gesamtartenliste und Rote Liste der Webspinnen, Weberknechte und Pseudoskorpione (Araneae, Opiliones, Pseudoscorpiones) mit Angaben zur Häufigkeit und Ökologie. *Natursch. Landschaftspfl. Brandenburg 8*(Suppl.) 1–79.

Platen, R., Moritz, M., Broen, B. von, Bothmann, I., Bruhn, K. & Simon, U. (1991). Liste der Webspinnen-und Weberknechtarten (Arach.: Araneida, Opilionida) des Berliner Raumes und ihre Auswertung für Naturschutzzwecke (Rote Liste). *A. Auhagen, R. Platen, H. Sukopp Rote List. der gefährdeten Pflanz. und Tiere Berlin Schwerpkt. Berlin (West). -Landschaftsentwicklung und Umweltforschung, Sonderh. S*, 6, 169–206.

#### 24) N50TAT_3: Saproxylic Beetles of the Steigerwald forest – *Martin M. Gossner and Joerg Mueller*

**Data owner(s)**: Martin M. Gossner^1^ and Joerg Mueller^2^

**Affiliation(s):** ^1^ Forest Entomology, Swiss Federal Research Institute WSL, Birmensdorf CH-8903, Switzerland.

^2^ Department of Animal Ecology and Tropical Biology, University of Würzburg, 97074 Würzburg, Germany.

**Description:** Activity densities of saproxylic beetles sampled by flight-interception traps in the understorey on 69 plots in the Steigerwald in 2004.

**Geographical scope:** Germany, forest sites in beech-dominated forests of the Steigerwald

**Taxonomical scope:** Saproxylic beetles (Coleoptera)

**Ecological scope:** Terrestrial (Forest)

**Number of sampling units:** 69 plots (500 m^2^)

**Minimum distance between sampling units:** 0.1 kilometre

**Maximum distance between sampling units:** 16 kilometres

**Environmental variables:** Protection, Amount of dead wood (m^3^ha^-1^), Number of tree genera, Age of the oldest trees, Percentage of Conifers.

**Functional traits: Mean:** Body size (mm), dead wood niche position regarding diameter, decay, and canopy cover preferences (for details see Gossner *et al*. 2013)

**Reference:** Gossner, M.M., Lachat, T., Brunet, J., Isacsson, G., Bouget, C., Brustel, H., *et al.* (2013). Current Near-to-Nature Forest Management Effects on Functional Trait Composition of Saproxylic Beetles in Beech Forests. *Conserv. Biol.*, 27, 605–614.

#### 25) N50TAT_4: Saproxylic Beetles across European beech forests– *Martin M. Gossner and Joerg Mueller*

**Data owner(s)**: Martin M. Gossner^1^ and Joerg Müller^2^

**Affiliation(s):** ^1^ Forest Entomology, Swiss Federal Research Institute WSL, Birmensdorf CH-8903, Switzerland

^2^ Department of Animal Ecology and Tropical Biology, University of Würzburg, 97074 Würzburg, Germany

**Description:** Activity densities of saproxylic beetles sampled by flight-interception traps in the understorey on 1156 forest sites across Europe between 1990 and 2010.

**Geographical scope:** Europe, 1156 forest sites in beech-dominated forests across Europe.

**Taxonomical scope:** Saproxylic beetles (Coleoptera)

**Ecological scope:** Terrestrial (Forest)

**Number of sampling units:** 241 plots (500 m^2^)

**Minimum distance between sampling units:** 0.1 kilometres

**Maximum distance between sampling units:** 1966 kilometres

**Environmental variables:** Time since Protection, climate data of the warmest month from WorldClim, Deadwood amount in 3 categories: low (0–29 m3 ha−1), medium (30–70 m3 ha−1), high (>70 m3 ha−1), Number of tree genera, proportion of broad-leaved trees in the surrounding relative to total forest area 3000m^2^, proportion of urban area in the surrounding 3000m^2^.

**Functional traits:** Mean Body size [mm], dead wood niche position regarding diameter, decay, and canopy cover preferences (for details see Gossner et al. 2013).

**Reference:** Gossner, M.M., Lachat, T., Brunet, J., Isacsson, G., Bouget, C., Brustel, H., *et al.* (2013). Current Near-to-Nature Forest Management Effects on Functional Trait Composition of Saproxylic Beetles in Beech Forests. *Conserv. Biol.*, 27, 605–614

#### 26) N50TFG: Saproxylic fungi data from Europe – Jacob Heilmann-Clausen, Morten Christensen, and Nerea Abrego.

**Data owner(s):** Jacob Heilmann-Clausen^1^, Morten Christensen and Nerea Abrego.

**Affiliation(s):** ^1^ Forest Entomology, Swiss Federal Research Institute WSL, Birmensdorf CH-8903, Switzerland

**Description:** Fungal fruit-body data collected across 53 beech (*Fagus sylvatica)* forest reserves across Europe. Within each of the reserves, logs of at least 10 cm in diameter and representing all decay classes (from decay class 1 to decay class 5) were visually inspected for fungal fruit bodies. The data were collected from 2001 to 20xx, each log was sampled once to three times. Altogether, the data comprises data on xxxx fungal species found in 1618 logs.

**Geographical scope:** Europe

**Taxonomical scope:** Wood-inhabiting fungi

**Ecological scope**: Beech (*Fagus sylvatica*) forest

**Number of sampling units:** 53 reserves

**Minimum distance between sampling units:** < 0.01 kilometres

**Maximum distance between sampling units:** 2532 kilometres

**Environmental variables:**

1. Area: are of the reserve (ha)

2. Altitude: mean elevation of the reserve (m.a.s.l.)

3. Temperature range: difference between the mean temperatures of the

warmest and coldest months of the year (°C).

4. Annual precipitation: Sum of the monthly precipitation values within a year (mm).

5. Connectivity 1: Connectivity of the reserve to other beech forests in a 1 km buffer area (index)

6. Connectivity 10: Connectivity of the reserve to other beech forests in a 1 km buffer area (index)

7. Connectivity 100: Connectivity of the reserve to other beech forests in a 100 km buffer area (index).

8. DBH: Diameter of the log at the breast height (cm)

9. Decay stage: Average decay stage of the log, in a scale ranging from 1 (early decay stage) to 5 (late decay stage).

**Functional traits:**

1. Fruit body type: Categorical trait classifying the fungal species according to their fruit body morphology. As agaricoids are classified species with a pileous and stipe, as corticioids are classified species developing smooth crust-like fruit bodies, as polyporoid resupinates are classified species developing poroid crust-like fruit bodies, as polyporoid pileates are classified species developing poroid fruit bodies with a pileous, as stromatoids are classified species developing crust-like fruit bodies with embedded sexual structures, as as “others” are classified those species for which their fruit body could not clearly be classified in either of these categories.
2. Fruit body longevity: Categorical trait indicating whether the species develop long-lived perennial fruit bodies (perennial) or annual (annual) fruit bodies
3. Presence of asexual structures in the hymenial layer: Categorical trait indicating whether fungi develop cystidia or paraphyses (yes/no)
4. Thickness of the spore wall: Categorical trait indicating whether the spore wall is thin or thick (thick/thin).
5. Spore ornamentation: Categorical trait indicating whether the spor develop ornamentations (yes/no).
6. Spore length: Continuous trait indicating the average length of the spores (µm).
7. Spore width: Continuous trait indicating the average spore width (µm).
8. Spore volume: Continuous trait indicating the average spore volume (µm^3^).
9. Standardized spore volume: Spore volume standardized to mean 0 and variance 1.
10. Spore shape: Categorical trait describing the spore shape (allantoid, angulose, cylindrical, ellipsoid, globose, ovoid).
11. Type of hyphal system: Categorical trait indicating the thickness of the cells in the hyphal system of the fruit bodies (monomitic, dimitic and trimitic in te case of basidiomycetes, and thick or thin in the case of ascomycetes).
12. Preferred host tree species: Categorical trait indicating what is the preferred host tree species (coniferous hosts, broadleaved hosts, both kinds, or only beech).
13. Preferred size class: Categorical trait indicating what is the preferred log size class (1 refers to species preferring logs smaller than 10 cm in diameter, 2 logs between 10 and 20 cm in diameter, and 3 logs larger than 20 in diameter).
14. Hymenium shape: Categorical trait indicating the shape of the reproductive layer (Angular pores, Branched, Circular pores, Gilled, Gleba, Hydnoid, Jelly, Laberintic pores, Meruloid, Perithecia, Reticulate, Smooth, Volva).
15. Presence of rhizomorphs: Categorical trait indicating whether fungi develop rhizomorphs (yes/no).
16. Rot type: Categorical trait indicating what kind of rot fungi exert (white rot, soft rot or brown rot).
17. Ecology: Categorical trait indicating the guild of (mycorrhizal, wood saprotrophic, litter saprotrophic, mycoparasitic…)

**Reference:** Hagge, J., Abrego, N., Bässler, C., Bouget, C., Brin, A., Brustel, H., *et al.* (2019). Congruent patterns of functional diversity in saproxylic beetles and fungi across European beech forests. *J. Biogeogr.*, 46, 1054–1065.

#### 27) N51TAT (1-5): Herb-dwelling arthropods from the Grassland sites of the Biodiversity Exploratories, Germany – *Martin M. Gossner and Wolfgang W. Weisser*

**Data owner(s)**: Martin M. Gossner^1^ and Wolfgang W. Weisser^2^

**Affiliation(s):** ^1^ Forest Entomology, Swiss Federal Research Institute WSL, Birmensdorf CH-8903, Switzerland

^2^ Terrestrial Ecology Research Group, Department of Ecology and Ecosystem Management, Technical University of Munich, Hans-Carl-von-Carlowitz-Platz 2, 85354 Freising, Germany

**Description:** Activity densities of arthropods sampled by sweepnet samples on 139 sites within the long-term and large-scale functional biodiversity project “Biodiversity Exploratories” in 2008.

**Geographical scope:** Germany, Biosphere Reserve Schwäbische Alb, “Hainich-Dün region, Biosphere Reserve, Schorfheide-Chorin”

**Taxonomical scope:** Arthropods:

1. Araneae
2. Coleoptera
3. Heteroptera
4. Homoptera
5. Orthoptera

**Ecological scope:** Terrestrial (Grasslands)

**Number of sampling units:** 139 plots (50m x 50m)

**Minimum distance between sampling units:** 0.2 kilometres

**Maximum distance between sampling units:** 625 kilometres

**Environmental variables:** Perimeter-area ratio (mean) of land cover types in a 500m radius around plots, Shannon diversity of land cover types in a 500m radius around plots, Proportion of grasslands in a 500m radius around plots, Elevation, Soil depth [cm], Soil Bulk Density [g/cm3] , Soil pH, Soil Texture (PCA axis), Soil Nutrients (PCA-axis), Land-use intensity combining mowing, grazing and fertilization between 2006 and 2008 following Blüthgen et al 2012

**Functional traits:** Based on Gossner *et al.* 2015

1. Feeding guild: herbivore, carnivore, detritivore, fungivore, omnivore
2. Stratum use: ground dwelling, herb layer, tree and shrub layer, soil dwelling, water, indifferent.
3. Dispersal ability (based on wing development across sexes in insects and based on ballooning and migration behaviour of spiders): 0 – low; 0.25 – low to intermediate; 0.5 – intermediate; 0.75 – intermediate to high; 1 – high

**References:** Blüthgen, N., Dormann, C.F., Prati, D., Klaus, V.H., Kleinebecker, T., Hölzel, N., *et al.* (2012). A quantitative index of land-use intensity in grasslands: Integrating mowing, grazing and fertilization. *Basic Appl. Ecol.*, 13, 207–220.

Gossner, M.M., Lewinsohn, T.M., Kahl, T., Grassein, F., Boch, S., Prati, D., *et al.* (2016). Land-use intensification causes multitrophic homogenization of grassland communities. *Nature*, 540, 266–269.

Gossner, M.M., Simons, N.K., Achtziger, R., Blick, T., Dorow, W.H.O., Dziock, F., *et al.* (2015). A summary of eight traits of Coleoptera, Hemiptera, Orthoptera and Araneae, occurring in grasslands in Germany. *Sci. Data*, 2, 150013.

Simons, N.K., Weisser, W.W. & Gossner, M.M. (2015). Multi-taxa approach shows consistent shifts in arthropod functional traits along grassland land-use intensity gradient. *Ecology*, 15-0616.1.

#### 28) N52FPP: Phytoplankton from Poland – Janne Soininen

**Data owner(s)**: Janne Soininen

**Affiliation(s):** University of Helsinki

**Description:** Data comprises phytoplankton samples from 46 Polish lakes. Sampled in 2010.

**Geographical scope:** Poland

**Taxonomical scope:** Phytoplankton

**Ecological scope:** Freshwater (Lakes)

**Environmental variables:** Lake area (hectar), mean depth (m), maximum depth (m), lake volume (1000 m^3^), pH, conductivity (µS/cm), water temperature (ºC), visibility (m), chlorophyll a (mg/l), NO3-N (mg/l), NH4-N (mg/l), total N (mg/l), PO4 (mg/l), total P (mg/l).

**Functional traits:** We classified the species according to its life form, i.e., colony, unicellular, and filament. We also observed the presence or absence of flagella, mucilage and aerotopes.

**Reference:** Kokocinski, M., Mankiewicz Boczek, J., Jurczak, T., Spoof, L., Meriluoto, J., Rajmonczyk, E., Hautala, H., Vehniäinen, M., Pawelczyk, J. & Soininen, J. 2013. Aphanizomenon gracile (Nostocales), a cylindrospermopsin-producing cyanobacterium in Polish lakes. Environmental Sciences and Pollution Research 20: 5243-5264.

#### 29) N56FAP: Macrophytes from Denmark lakes – *Lars Baastrup-Spohr and Janne Alahuhta.*

**Data owner(s)**: Lars Baastrup-Spohr^1^ and Janne Alahuhta^2^

**Affiliation(s):** ^1^ Freshwater Biological Laboratory, Department of Biology, University of Copenhagen, Universitetsparken 4 2100 København Ø, Denmark.

^2^ Geography Research Unit, University of Oulu, P.O. Box 3000, 90014 Oulu, Finland.

**Description:** 208 lake samples taken during a growing season between 2010 and 2015.

**Geographical scope:** Denmark

**Taxonomical scope:** Plant (Macrophyte)

**Ecological scope:** Freshwater (Lakes)

**Number of sampling units:** 208 lakes with different plots (75-200) depending on lake size.

**Minimum distance between sampling units:** 0.50 kilometres

**Maximum distance between sampling units:** 454 kilometres

**Environmental variables:** Average values from monthly samplings between May and September of alkalinity (meq/l), chlorophyl a (µg/l), TN (mg/L), TP (mg/L) and Secchi depth (m) collected between 2010-2015, complemented with lake area (km^2^) and maximum lake depth (m).

**Functional traits:** Life form (submerged, helophyte, floating-leaved, free-floating), normal method of propagation (by seed/spore, mostly by seed/spore but also vegetatively, by seed/spore and vegetatively, mostly vegetatively and also by seed/spore and vegetatively; Klotz et al. 2002), maximum height (based on Mossberg and Stenberg 2012) and perennation (annual or perennial; Willby et al. 2000).

**Reference:** Klotz, S. et al. 2002. BIOLFLOR – Eine Datenbank zu biologischökologischen Merkmalen der Gefäßpflanzen in Deutschland. *Schr. Reihe Vegetationskde* 38: 1–334.

Lindholm, M., Alahuhta, J., Heino, J. & Toivonen, H. (2020). No biotic homogenisation across decades but consistent effects of landscape position and pH on macrophyte communities in boreal lakes. *Ecography (Cop.).*, 43, 294–305.

Mossberg, B. and Stenberg, L. 2012. Suuri Pohjolan kasvio. Tammi.

Willby, N. J. et al. 2000. Attribute-based classification of European hydrophytes and its relationship to habitat utilization. *Freshwater Biology* 43: 43–74.

#### 30) N59MAP: Macrophytes from Finland – *Camilla Gustafsson, Alf Norkko*

**Data owner(s)**: Camilla Gustafsson ^1^ and Alf Norkko^1,2^

**Affiliation(s):** ^1^ University of Helsinki, Tvärminne Zoological Station, J.A. Palménin tie 260, FI-10900 Hanko, Finland.

^2^ Baltic Sea Centre, Stockholm University, SE-106 91, Stockholm, Sweden

**Description:** 17 morphological and chemical (continuous) traits for marine and limnic aquatic plants from sites along a wave exposure gradient

**Geographical scope:** Finland, Gulf of Finland (Baltic Sea)

**Taxonomical scope:** Vascular plant

**Ecological scope:** Marine (brackish)

**Number of sampling units:** 30 (plots of 1 m²)

**Minimum distance between sampling units:** 0.3 kilometres

**Maximum distance between sampling units:** 48 kilometres

**Environmental variables:** Depth (m), salinity, temperature (°C), exposure index (unitless, integrating depth and LG10-transformed) NH4^+^-N (µM) water column, NH4^+^-N (µM) porewater, sediment organic matter (%), Sediment grain size mean (according to Folk & Warde 1957), grain size fraction 2 mm, grain size fraction 1 mm, grain size fraction 0.5 mm, grain size fraction 0.250 mm, grain size fraction 0.125 mm, grain size fraction >0.063 mm, grain size fraction <0.063 mm

**Functional traits:** In the laboratory, nine different morphological and chemical traits were measured on each plant species within a plot following standardized trait measurement protocols by Pérez‐Harguindeguy et al. ([2013](https://besjournals.onlinelibrary.wiley.com/doi/full/10.1111/1365-2745.13011#jec13011-bib-0051)). The maximum vegetative height (Hmax, cm), specific leaf area (SLA, mm^2^/mg), the leaf and root nitrogen (Leaf and Root N, mg g^-1^), elemental concentrations (Leaf and Root CN) and leaf and root δ^15^N and δ^13^C, the maximum root length (Root, cm), and the root: shoot ratio (R: S‐ratio).

**Reference:** Folk, R.L. & Ward, W.C. (1957). Brazos River bar [Texas]; a study in the significance of grain size parameters. *J. Sediment. Res.*, 27, 3–26.

Gustafsson, C. & Norkko, A. (2019). Quantifying the importance of functional traits for primary production in aquatic plant communities. *J. Ecol.*, 107, 154–166.

Pérez-Harguindeguy, N., Díaz, S., Garnier, E., Lavorel, S., Poorter, H., Jaureguiberry, P., *et al.* (2013). New handbook for standardised measurement of plant functional traits worldwide. *Aust. J. Bot.*, 61, 167.

#### 31) N59MMI: Bentic macrofauna across habitats in Finland – Johanna Gammal and Alf Norkko

**Data owner(s)**: Johanna Gammal and Alf Norkko

**Affiliation(s):** Tvärminne Zoological Station, University of Helsinki. J.A. Palméns väg 260, FI-10900 Hangö, Finland.

**Description:** The dataset includes information on benthic macrofauna, functional traits, and environmental variables from 18 shallow (3-4 m deep) coastal sites on a sedimentary gradient in the Gulf of Finland, Northern Baltic Sea. At each site 5 replicate samples of macrofauna and environmental variables were collected by SCUBA diving.

**Geographical scope:** Finland \ Hanko (Gulf of Finland, Baltic Sea)

**Taxonomical scope:** Benthic macrofauna

**Ecological scope:** Marine (brackish)

**Number of sampling units:** 18 plots (1m²)

**Minimum distance between sampling units:** 0.15 kilometres

**Maximum distance between sampling units:** 19 kilometres

**Environmental variables:** Depth (m), temperature (°C), salinity, sediment organic content (LOI, %), C/N, sediment chlorophyll a content (µg/g sediment), grain size (% per fraction, median grain size µm), oxygen (mg/l) and nutrient concentrations (µmol/l) in the bottom water and nutrient concentrations (µmol/l) in the porewater.

**Functional traits:**

1. Feeding group: suspension feeder, surface detrivore, borrowing detrivore, carnivore, herbivore
2. Mobility: stationary, swim, crawl
3. Body size: XS, S, M, L
4. Longevity: <1 year, 1-2 years, 2-3 years, 3-5 years, 5-10 years,
5. Development: larval, brooding/direct
6. Living habit: tube-dweller, gallery diffuser, bio diffuser
7. Position: infaunal, deeper infaunal, pelagic, epibenthic.

**Reference:** Gammal, J., Järnström, M., Bernard, G., Norkko, J. & Norkko, A. (2019). Environmental Context Mediates Biodiversity–Ecosystem Functioning Relationships in Coastal Soft-sediment Habitats. *Ecosystems*, 22, 137–151.

Bernard, G., Gammal, J., Järnström, M., Norkko, J. & Norkko, A. (2019). Quantifying bioturbation across coastal seascapes: Habitat characteristics modify effects of macrofaunal communities. *J. Sea Res.*, 152, 101766.

#### 32) N61FBD: East_FIN stream diatoms – *Virpi Pajunen*

**Data owner(s)**: Virpi Pajunen

**Affiliation(s):** Department of Geosciences and Geography, University of Helsinki

**Description:** Dataset comprised 51 streams located in South-Eastern Finland. Each stream was sampled once in one riffle site during Summer 2017.

**Geographical scope:** South-eastern Finland

**Taxonomical scope:** Diatoms

**Ecological scope:** Freshwater

**Number of sampling units:** 51 sites

**Minimum distance between sampling units:** 1.25 kilometres

**Maximum distance between sampling units:** 489 kilometres

**Environmental variables:** Water colour, Total nitrogen, Total phosphorus, Isotopes (hydrogen-2 and oxygen-18), anions, cations and metals (Li, Be, Al, Si, V, Cr, Mn, Fe, Co, Ni, Cu, Zn, As, Se, Sr, Mo, Cd, Sn, Sb, Ba, Tl, Pb, Bi, U, F, Cl, NO3, SO4, Na, NH4, K, Ca, Mg), Alkalinity, Sediment temperature, Water temperature, Pressure, DO%, DO concentration, Conductivity, pH, Moss cover (%), Substrate % (Sand, Gravel, Pebble, Cobble, Boulders, and Bedrock), Stream width, Stream depth, Shading by the canopy, Current velocity

**Functional traits:** Life‐forms (mobile, adnate, pedunculate, pad-attached, stalk-attached, colonial; Rimet & Bouchez 2012), ecological guilds (high-profile, low-profile, motile; Passy 2007; Rimet & Bouchez 2012), acid tolerance, nitrogen-fixing ability (Soininen et al. 2016).

**Reference:**

#### 33) N62FMI: Macroinvertebrates from Western Finnish streams and rivers – *Jani Heino and Kimmo T. Tolonen*

**Data owner(s)**: Jani Heino and Kimmo T. Tolonen

**Affiliation(s):** Finnish Environment Institute, Freshwater Centre, P.O. Box 413, FIN-90014 Oulu, Finland.

**Description:** A total of 105 stream and river sites were sampled in September 2014.

**Geographical scope:** Finland

**Taxonomical scope:** Aquatic macroinvertebrates

**Ecological scope:** Freshwater (Lotic systems)

**Number of sampling units:** 105 sites from different sized rivers

**Minimum distance between sampling units**: 0.82 kilometres

**Maximum distance between sampling units:** 569 kilometres

**Environmental variables:** bedrock, rocks, moraine, clay, total N, total P, colour, conductivity, pH, deciduous trees, shading, velocity, depth, discharge, moss, sand, gravel, pebble, cobble, boulder.

**Functional traits:**

1. Functional feeding groups: filterers, gatherers, shredders, scrapers, piercers, and predators.
2. Habit trait groups: burrowers, climbers, crawlers, sprawlers, semi‐sessiles, swimmers‐divers
3. Body mass: ranging from 0.2 to 26130

**Reference:** Perez Rocha, M., Bini, L.M., Domisch, S., Tolonen, K.T., Jyrkänkallio-Mikkola, J., Soininen, J., *et al.* (2018). Local environment and space drive multiple facets of stream macroinvertebrate beta diversity. *J. Biogeogr.*, 45, 2744–2754.

#### 34) N62FBD: West_FIN stream diatoms – *Virpi Pajunen*

**Data owner(s)**: Virpi Pajunen^1^ and Jenny Jyrkänkallio-Mikkola^2^

**Affiliation(s):** ^1^Department of Geosciences and Geography, University of Helsinki; ^2^WWF Finland

**Description:** Dataset comprised 105 streams located in Western coast of Finland. Each stream was sampled once in one riffle site in September 2014.

The data were originally used in doi: 10.1111/jbi.13002

**Geographical scope:** Western Finland

**Taxonomical scope:** Diatoms

**Ecological scope:** Freshwater

**Number of sampling units:** 105

**Minimum distance between sampling units:** 0.82 kilometres

**Maximum distance between sampling units:** 569 kilometres

**Environmental variables:** Total nitrogen,	Total phosphorus, Water colour, Conductivity, pH, deciduous trees(%), stream width, canopy shading, current velocity, stream depth, moss cover, substrate (sand%, gravel%, pebble%, cobble%, boulders%), catchment are size, slope, land use and cover (Artificial areas%, Agricultural areas%, Broad leaved forest%, Coniferous forest%, Mixed forest%, Shrubs%, Open spaces%, Wetlands%, Water bodies%), soil type (Bedrock%, Rocks%, Coarse grained soil%, Moraine%, Fine grained soil%, Clay%, Muddy soil%, Peat%,)

**Functional traits:** Life‐forms (mobile, adnate, pedunculate, pad-attached, stalk-attached, colonial; Rimet & Bouchez 2012), ecological guilds (high-profile, low-profile, motile; Passy 2007; Rimet & Bouchez 2012), acid tolerance, nitrogen-fixing ability (Soininen et al. 2016).

**Reference:** Jyrkänkallio-Mikkola, J., Meier, S., Heino, J., Laamanen, T., Pajunen, V., Tolonen, K.T., *et al.* (2017). Disentangling multi-scale environmental effects on stream microbial communities. *J. Biogeogr.*, 44, 1512–1523.

#### 35) N62MBD: Diatoms from the Baltic Sea – Leena Virta

**Data owner(s)**: Leena Virta¹²

**Affiliation(s):** ¹Department of Geosciences and Geography, University of Helsinki, Helsinki, Finland

²Tvärminne Zoological Station, University of Helsinki, Hanko, Finland;

**Description:** Benthic epilithic diatoms collected from the Gulf of Finland and Gulf of Bothnia

**Geographical scope:** Finland / Baltic Sea

**Taxonomical scope:** Diatoms

**Ecological scope:** Marine (Brackish)

**Number of sampling units:** 37 plots (0.0025m²)

**Minimum distance between sampling units:** 13.03 kilometres

**Maximum distance between sampling units:** 661 kilometres

**Environmental variables:** Water temperature (°C), pH, salinity, total P (µg/l), total N (ml/l), Si, oxygen (mg/l)

**Functional traits:** size (biovolume classes: large > 1000µm³ / small < 1000µm³), mobility (mobile / non-mobile), type of attachment (adnate / pedunculate [which was further divided to pad-attached / stalk-attached] / non-attached), colonization (colonial / non-colonial), growth form (low-profile / high-profile / motile / planktonic), and nitrogen-fixing abilities (nitrogen-fixer / non-nitrogen-fixer)

**Reference:** Virta, L. & Soininen, J. (2017). Distribution patterns of epilithic diatoms along climatic, spatial and physicochemical variables in the Baltic Sea. *Helgol. Mar. Res.*, 71, 16.

#### 36) N63FAP: Macrophytes from Finnish lakes – *Janne Alahuhta.*

**Data owner(s)**: Janne Alahuhta^1,2^

**Affiliation(s):** ^1^ Geography Research Unit, University of Oulu, P.O. Box 3000, 90014 Oulu, Finland.

^2^ Finland & Finnish Environment Institute, Freshwater Centre, P.O. Box 413, 90014 Oulu, Finland.

**Description:** 150 surveyed lakes between 2006-2012 (June-September) covering most of Finland. In addition to true aquatic plants (i.e. hydrophytes), emergent plant species were also recorded (Gillard et al. 2020). Samples were taken using main belt transect method, where each 5-m wide transect extended, perpendicular to the shoreline, from the upper eulittoral to the outer depth limit of vegetation (Gillard et al. 2020).

**Geographical scope:** Finland

**Taxonomical scope:** Plant (Macrophyte)

**Ecological scope:** Freshwater (Lakes)

**Number of sampling units:** 150

**Minimum distance between sampling units:** 2.35 kilometres

**Maximum distance between sampling units:** 890 kilometres

**Environmental variables:** Multiple samples of alkalinity (mM), colour (mg PT/L), conductivity (mS/m), pH, TN (μg/L), TP (μg/L), Secchi depth (m) and turbidity (PTU) collected between 2006-2012, complemented with lake area (km^2^).

**Functional traits:** Life form (submerged, helophyte, floating-leaved, free-floating), normal method of propagation (by seed/spore, mostly by seed/spore but also vegetatively, by seed/spore and vegetatively, mostly vegetatively and also by seed/spore and vegetatively; Klotz et al. 2002), maximum height (based on Mossberg and Stenberg 2012) and perennation (annual or perennial; Willby et al. 2000).

**Reference:** Gillard, M.B., Aroviita, J. & Alahuhta, J. (2020). Same species, same habitat preferences? The distribution of aquatic plants is not explained by the same predictors in lakes and streams. *Freshw. Biol.*, 65, 878–892.

Klotz, S. et al. 2002. BIOLFLOR – Eine Datenbank zu biologischökologischen Merkmalen der Gefäßpflanzen in Deutschland. *Schr. Reihe Vegetationskde* 38: 1–334.

Lindholm, M., Alahuhta, J., Heino, J. & Toivonen, H. (2020). No biotic homogenisation across decades but consistent effects of landscape position and pH on macrophyte communities in boreal lakes. *Ecography (Cop.).*

Mossberg, B. and Stenberg, L. 2012. Suuri Pohjolan kasvio. Tammi.

Willby, N.J., Abernethy, V.J. & Demars, B.O.L. (2000). Attribute-based classification of European hydrophytes and its relationship to habitat utilization. *Freshw. Biol.*, 43, 43–74.

#### 37) N63FAP_2: Macrophytes from Finnish rivers – *Jukka Aroviita and Janne Alahuhta*

**Data owner(s)**: Jukka Aroviita^1^ and Janne Alahuhta^1,2^

**Affiliation(s):** ^1^ Finland & Finnish Environment Institute, Freshwater Centre, P.O. Box 413, 90014 Oulu, Finland.

^2^ Geography Research Unit, University of Oulu, P.O. Box 3000, 90014 Oulu, Finland.

**Description:** 150 surveyed rivers covering most of Finland in July–August between 2009 and 2016 (Gillard et al. 2020). Samples were surveyed from two different sections (divided into five 20-m long subsections) of 100 m in each stream: a riffle and adjacent pool section (Gillard et al. 2020).

**Geographical scope:** Finland

**Taxonomical scope:** Plants (Macrophyte)

**Ecological scope: Freshwater (Rivers)**

**Number of sampling units:** 150

**Minimum distance between sampling units:** 1.43 kilometres

**Maximum distance between sampling units:** 944 kilometres

**Environmental variables:** Multiple samples of alkalinity (mM), colour (mg PT/L), conductivity (mS/m), pH, TN (μg/L), TP (μg/L), Secchi depth (m) and turbidity (PTU) collected between 2006-2012, complemented with stream wide (m).

**Functional traits:** Life form (submerged, helophyte, floating-leaved, free-floating), normal method of propagation (by seed/spore, mostly by seed/spore but also vegetatively, by seed/spore and vegetatively, mostly vegetatively and also by seed/spore and vegetatively; Klotz *et al.* 2002), maximum height (based on Mossberg and Stenberg 2012) and perennation (annual or perennial; Willby et al. 2000).

**Reference:** Gillard, M.B., Aroviita, J. & Alahuhta, J. (2020). Same species, same habitat preferences? The distribution of aquatic plants is not explained by the same predictors in lakes and streams. *Freshw. Biol.*, 65, 878–892.

Kühn, I., Durka, W. & Klotz, S. (2004). BiolFlor - a new plant-trait database as a tool for plant invasion ecology. *Divers. Distrib.*, 10, 363–365.

Lindholm, M., Alahuhta, J., Heino, J. & Toivonen, H. (2020). No biotic homogenisation across decades but consistent effects of landscape position and pH on macrophyte communities in boreal lakes*. Ecography (Cop.).,* 43, 294–305.

Mossberg, B., Stenberg, L., Vuokko, S. & Väre, H. (2012). *Suuri pohjolan kasvio*. Tammi.

Willby, N.J., Abernethy, V.J. & Demars, B.O.L. (2000). Attribute-based classification of European hydrophytes and its relationship to habitat utilization. *Freshw. Biol.*, 43, 43–74.

#### 38) N63TBI: Birds from Finland – *Aleksi Lehikoinen*

**Data owner(s)**: Aleksi Lehikoinen

**Affiliation(s):** Finnish Museum of Natural History, PO Box 17, FI-00014 University of Helsinki, Finland.

**Description:** Includes (1) bird surveys across the country using systematic sampling design (Lindström et al. 2015), (2) species-trait matrix (migratory behaviour and body mass; Tikhonov et al. 2020), (3) local environmental variable matrix for the plots (Tikhonov et al. 2020) and (4) plot coordinates.

**Geographical scope:** Finland

**Taxonomical scope:** Birds

**Ecological scope:** Terrestrial and aquatic ecosystems

**Number of sampling units:** 241

**Minimum distance between sampling units:** 2.6 km

**Maximum distance between sampling units:** 1157 kilometres

**Environmental variables:** Totally, 10 environmental variables were measured. Habitat type (i.e., the relative amount of Broadleaved forests, coniferous forests, open habitats, urban habitats and wetlands), Temperature (ºC) for 2013, winter temperature, and sampling effort in meters (usually 6000).

**Functional traits:** migratory behaviour (i.e. resident, short-distance migrant or tropical long-distance migrant) and body mass (i.e. weight in grams).

**References:** Lindström, Å., Green, M., Husby, M., Kålås, J.A. & Lehikoinen, A. (2015). Large-Scale Monitoring of Waders on Their Boreal and Arctic Breeding Grounds in Northern Europe. *Ardea*, 103, 3–15.

Tikhonov, G., Opedal, Ø.H., Abrego, N., Lehikoinen, A., Jonge, M.M.J., Oksanen, J., *et al.* (2020). Joint species distribution modelling with the package Hmsc. *Methods Ecol. Evol.*, 11, 442–447.]

#### 39) N64FBD: Benthic diatoms from Iceland – Jianjun Wang, Vilja Tupola, and Janne Soininen

**Data owner(s)**: Jianjun Wang^1^, Janne Soininen^2^, Vilja Tupola^2^

**Affiliation(s):** Department of Geosciences and Geography, University of Helsinki

**Description:** Data comprises benthic stream diatoms from 45 sites in Iceland. Sampled in 2013.

**Geographical scope:** Iceland

**Taxonomical scope:** Diatoms

**Ecological scope:** Freshwater

**Number of sampling units:** 45 sites

**Minimum distance between sampling units:** 0.13 kilometres

**Maximum distance between sampling units:** 318 kilometres

**Environmental variables:** Conductivity (µS/cm), water temperature (ºC), stone size (cm^3^), depth (cm), width (m), current velocity (m/s), total organic carbon (ppm), NO_x_-N (mg/l), NO2-N (mg/l), PO4-P (µg/l), NH4-N (mg/l), total N (mg/l), total P (mg/l).

**Functional traits:** Life‐forms (mobile, adnate, pedunculate, pad-attached, stalk-attached, colonial; Rimet & Bouchez 2012), ecological guilds (high-profile, low-profile, motile; Passy 2007; Rimet & Bouchez 2012)

**Reference:** Tupola, V. 2019. Islannin virtavesien piileväyhteisöt – vertailukohteena Pohjois-Fennoskandia. MSc thesis (in Finnish), University of Helsinki.

#### 40) N65FBD: Benthic diatoms from Finland – Janne Soininen

**Data owner(s)**: Janne Soininen

**Affiliation(s):** Department of Geosciences and Geography, University of Helsinki

**Description:**

**Geographical scope:** Finland

**Taxonomical scope:** Benthic diatoms

**Ecological scope:** Freshwaters

**Number of sampling units:** 105 sites

**Minimum distance between sampling units:** 0.99 kilometres

**Maximum distance between sampling units:** 1093 kilometres

**Environmental variables:**

**Functional traits:** Life‐forms (mobile, adnate, pedunculate, pad-attached, stalk-attached, colonial; Rimet & Bouchez 2012), ecological guilds (high-profile, low-profile, motile; Passy 2007; Rimet & Bouchez 2012)

**Reference:**

#### 41) N65TBF: Butterflies from Finland – *Miska Luoto*

**Data owner(s)**: Miska Luoto

**Affiliation(s):** University of Helsinki

**Description:** Day-active butterfly data covering years 2002-2010

**Geographical scope:** Norway

**Taxonomical scope:** Day active butterflies (Lepidoptera)

**Ecological scope:** Terrestrial

**Number of sampling units:** 1466

**Minimum distance between sampling units:** 10 kilometres

**Maximum distance between sampling units:** 1200 kilometres

**Environmental variables:** cover of rocky soils, cover of sand/gravel soils, cover of clay soils, cover of built up area, cover of arable land, cover of sparse vegetation, cover of wetland

**Functional traits:**

1. Width of larval host plant use
2. Growth form of host plant
3. Number of adult foraging habitats
4. Mobility index
5. Body size (wing span in mm)

**Reference:** Eskildsen, A., le Roux, P.C., Heikkinen, R.K., Høye, T.T., Kissling, W.D., Pöyry, J., *et al.* (2013). Testing species distribution models across space and time: High latitude butterflies and recent warming. *Glob. Ecol. Biogeogr.*

#### 42) N66FBD: North_FIN stream diatoms – *Virpi Pajunen*

**Data owner(s)**: Virpi Pajunen

**Affiliation(s):** Department of Geosciences and Geography, University of Helsinki

**Description:** Dataset comprised 28 streams located in Northern Finland. Each stream was sampled once in one riffle site in September 2015.

The data were used as a part of larger data set in:

https: //doi.org/10.1002/eap.1917

**Geographical scope:** Northern Finland

**Taxonomical scope:** Diatoms

**Ecological scope:** Freshwater

**Number of sampling units:** 28 sites

**Minimum distance between sampling units:** 1.96 kilometres

**Maximum distance between sampling units:** 650 kilometres

**Environmental variables:** Total phosphorus, total nitrogen, pH, conductivity, water colour, water temperature, current velocity, stream depth, stream width, canopy shading, particle size.

**Functional traits:** Life‐forms (mobile, adnate, pedunculate, pad-attached, stalk-attached, colonial; Rimet & Bouchez 2012), ecological guilds (high-profile, low-profile, motile; Passy 2007; Rimet & Bouchez 2012), acid tolerance, nitrogen-fixing ability (Soininen et al. 2016).

**Reference:** Pajunen, V., Jyrkänkallio‐Mikkola, J., Luoto, M. & Soininen, J. (2019). Are drivers of microbial diatom distributions context dependent in human‐impacted and pristine environments? *Ecol. Appl.*, 29.

#### 43) N66FMI: Macroinvertebrates from Koutajoki lakes, Finland – *Jani Heino and Kimmo T. Tolonen*

**Data owner(s)**: Jani Heino and Kimmo T. Tolonen

**Affiliation(s):** Finnish Environment Institute, Freshwater Centre, P.O. Box 413, FIN-90014 Oulu, Finland

**Description:** A total of 48 lakes were sampled in the Koutajoki drainage basin in September 2005.

**Geographical scope:** Finland

**Taxonomical scope:** Macroinvertebrates

**Ecological scope:** Freshwater

**Number of sampling units:** 48 lakes differing in surface area

**Minimum distance between sampling units:** 0.26 kilometres

**Maximum distance between sampling units:** 18 kilometres

**Environmental variables:** Conductivity, colour, total phosphorus, macrophytes, shore depth, shoreline, shore develop, PC1 particles, PC2 particles (see the reference below for details).

**Functional traits:**

1. Feeding habits: scrapers, piercers, collector-gatherers,

filterers, commensals, parasites, and predators

2. Locomotion and substrate relation: Burrower, crawler, semi-sessile, sessile, swimmer
3. Body mass: ranging from 0.02 to 3051

**Reference:** Heino, J. & Tolonen, K.T. (2017). Ecological drivers of multiple facets of beta diversity in a lentic macroinvertebrate metacommunity. *Limnol. Oceanogr.*

#### 44) N66FMI_2: Macroinvertebrates from Koutajoki streams and rivers, Finland – *Jani Heino*

**Data owner(s)**: Jani Heino

**Affiliation(s):** Finnish Environment Institute, Research Programme for Biodiversity, P.O. Box 413, FIN-90014 Oulu, Finland

**Description:** A total of 77 stream and river sites were sampled in the autumn of 2001.

**Geographical scope:** Finland

**Taxonomical scope:** Macroinvertebrates

**Ecological scope:** Freshwater (Lotic systems)

**Number of sampling units:** 77 sites from rivers or streams differing in size

**Minimum distance between sampling units:** 0.18 kilometres

**Maximum distance between sampling units:** 60 kilometres

**Environmental variables:** Catchment area, distance to upstream lake, area of upstream lake, stream gradient (slope), site altitude, and % peatlands (wetland), % deciduous forest, % coniferous forest, and % mixed forests in the catchment.

**Functional traits:**

1. Functional feeding group: Collector-gatherer, commensal, filterer, piercer, predator, scraper, shredder
2. Habit trait groups: Burrowers, climbers, clingers, divers, sprawlers, swimmers
3. Body size classes: 0.25-0.5cm, 0.5-1.0cm, 1-2cm, 2-4cm, 4-8cm.

**Reference:** Heino, J., Mykrä, H. & Kotanen, J. (2008). Weak relationships between landscape characteristics and multiple facets of stream macroinvertebrate biodiversity in a boreal drainage basin. *Landsc. Ecol.*, 23, 417–426.

#### 45) N68FBD: Diatoms from Finland and Norway – *Anette Teittinen and Janne Soininen*

**Data owner(s)**: Anette Teittinen and Janne Soininen

**Affiliation(s):** Department of Geosciences and Geography, University of Helsinki, Helsinki, Finland

**Description:** Benthic diatom dataset collected from 102 freshwater ponds in subarctic Finland and Norway in August 2015.

**Geographical scope:** Finland and Norway

**Taxonomical scope:** Benthic diatoms

**Ecological scope:** Freshwater (ponds)

**Number of sampling units:** 102

**Minimum distance between sampling units:** 0.01 kilometres

**Maximum distance between sampling units:** 58 kilometres

**Environmental variables:** Water pH, conductivity (µS/cm), temperature (°C), total nitrogen (mg/l), silica (mg/l), calcium (mg/l), magnesium (mg/l), potassium (mg/l).

**Functional traits:** Life‐forms (mobile, adnate, pedunculate, pad-attached, stalk-attached, colonial; Rimet & Bouchez 2012), ecological guilds (high-profile, low-profile, motile; Passy 2007; Rimet & Bouchez 2012), acid tolerance, nitrogen-fixing ability (Soininen et al. 2016).

**References**:

Passy, S.I. (2007). Diatom ecological guilds display distinct and predictable behavior along nutrient and disturbance gradients in running waters. *Aquat. Bot.*, 86, 171–178.

Rimet, F. & Bouchez, A. (2012). Life-forms, cell-sizes and ecological guilds of diatoms in European rivers. *Knowl. Manag. Aquat. Ecosyst.*, 01.

Soininen, J., Jamoneau, A., Rosebery, J. & Passy, S.I. (2016). Global patterns and drivers of species and trait composition in diatoms. *Glob. Ecol. Biogeogr.*, 25, 940–950.

Teittinen, A., Weckström, J. & Soininen, J. (2018). Cell size and acid tolerance constrain pond diatom distributions in the subarctic. *Freshw. Biol.*, 63, 1569–1578.

#### 46) N67TTP: Plants from Greenland - *Julia Kemppinen*

**Data owner(s)**: Pekka Niittynen, Julia Kemppinen and Miska Luoto

**Affiliation(s):** Department of Geosciences and Geography, University of Helsinki

**Description:** Data comprising 58 plant species

**Geographical scope:** Greenland

**Taxonomical scope:** Terrestrial vascular plants

**Ecological scope:** Terrestrial

**Number of sampling units:** 72 plots

**Minimum distance between sampling units: <** 0.01 kilometres

**Maximum distance between sampling units:** 3.77 kilometres

**Environmental variables:** Within each plot, several abiotic and biotic variables were collected. Rock cover (%; i.e. the cover of rocks or boulders large enough to inhibit plant colonization and growth) was visually estimated. Soil depth (cm) was measured at four different points in each cell, and an average soil depth then calculated for the cell. Mesotopography was scored a value between 1 and 10, where 1 represents a valley bottom, 5 the middle of a slope and 10 the slope crest. Soil moisture (% volumetric water content) measurements were taken in each plot. Soil samples were taken from a minimum of 16 cells per grid and used to determine soil pH in the laboratory using the CaCl_2_ method (with pH linearly interpolated for all other cells within a grid). Slope angle (°) was measured in the field for each cell. Slope angle, aspect and latitude was subsequently used to calculate potential incident solar radiation (MJ · cm–2 · yr–1) for each cell (following McCune & Keon, 2002).

**Functional traits:**

| *Functional traits* | *Unit* |
| --- | --- |
| Specific leaf area | m2kg-1 |
| Leaf N | mg g-1 |
| Leaf P | mg g-1 |
| Plant height | m |
| Leaf dry matter content | g g-1 |
| Seed mass | mg |
| Leaf area | mm2 |

#### 47) N45FFI_2: Fish fauna from Italy – *Anna Gavioli et al.*

**Data owner(s)**: Anna Gavioli ^1^, Giuseppe Castaldelli ^1^ and Marco Milardi ^2^

**Affiliation(s):** ^1^ University of Ferrara, Department of Life Sciences and Biotechnology, Ferrara, V. Luigi Borsari 46, 44121, Italy.

^2^ Fisheries New Zealand, Tini a Tangaroa, Ministry for Primary Industries, Manatū Ahu Matua, 34-38 Bowen Street, Wellington, New Zealand

**Description:** Data were collected within a monitoring program for the compilation of official Fish Inventories in northern Italy. The dataset included data from the Emilia-Romagna region, the Padova province, the Po River and the Oglio River. Fish species abundance is expressed using Moyle classes (Moyle & Nichols, 1973) ranging from 1 (lower abundance, 1-2 individuals per site) to 5 (higher abundance, more than 50 individuals per site).

**Geographical scope:** Italy

**Taxonomical scope:** Fish fauna

**Ecological scope:** Freshwaters (waterways, i.e., streams, rivers, canals)

**Number of sampling units:** 337

**Minimum distance between sampling units:** < 0.01 kilometres

**Maximum distance between sampling units:** 454 kilometres

**Environmental variables:** Water quality data were given by the Environmental Protection Agencies of Emilia-Romagna, Veneto, Lombardy and Piedmont Regions. Totally, 8 water physico-chemical variables were present: water temperature (°C), electrical conductivity (μS cm-1), chemical oxygen demand (COD (O2 mg l-1)), biological oxygen demand (BOD5 (O2 mg l-1)), total suspended solids (mg l-1), total phosphorus (P mg l-1), ammonia (N mg l-1) and nitrate nitrogen (N mg l-1).

**Functional traits:** Fish ecological traits in the dataset were based on an article by Milardi and Castaldelli (2018):

| *Ecological function* | *Traits* | *Description* |
| --- | --- | --- |
| Feeding | Planktivores | Plankton feeders |
|  | Herbivores | Vegetation feeders |
|  | Benthivores | Bottom feeders |
|  | Invertivores | Invertebrate feeders |
|  | Piscivores | Fish feeders |
|  | Parasite | Ematophages |
|  | Generalists | Unspecialized feeding |
| Reproduction | Lithophils | Spawning on stones or gravel |
|  | Phytophils | Spawning on submerged vegetation |
|  | Phytolithophils | Spawning on stones or vegetation |
|  | Psammophils | Spawning on sand or mud |
|  | Ostracophils | Spawning in molluscs |
|  | Pelagophils or live breeding | Pelagic spawners or live spawners |
|  | Polyphils | Generalist spawners |
|  | Sea spawning | Saltwater spawners |
| Migration | Short | Within or close to the site |
|  | Intermediate | Up and downstream or into flooded areas |
|  | Long | Anadromous and catadromous species |
| Tolerance | Low oxygen tolerants | Tolerance/intolerance to low oxygen  (indicatively below 3 ppm) |
|  | Low oxygen intolerants |  |
|  | High temperature tolerants | Tolerance/intolerance to high temperature (indicatively above 20 °C) |
|  | High temperature intolerants |  |
| Habitat use | Rheophiles | Preferring fast flowing water |
|  | Limnophiles | Preferring slow or no current |
|  | Eurytopics | Having no preference on current velocity |
|  | Clear water | Clear water adapted |
|  | Turbid water | Turbid water adapted |
|  | Wide range of conditions | Adapted to a wide range of water turbidity |

**References**: Gavioli, A., Milardi, M., Castaldelli, G., Fano, E.A. & Soininen, J. (2019). Diversity patterns of native and exotic fish species suggest homogenization processes, but partly fail to highlight extinction threats. *Divers. Distrib.*, 25, 983–994.

Milardi, M. & Castaldelli, G. (2018). A novel approach to an ecofunctional fish index for Mediterranean countries. *Ecol. Indic.*, 89, 376–385.

Milardi, M., Gavioli, A., Castaldelli, G. & Soininen, J. (2020). Partial decoupling between exotic fish and habitat constraints remains evident in late invasion stages. *Aquat. Sci.*, 82, 14.

Milardi, M., Gavioli, A., Soininen, J. & Castaldelli, G. (2019). Exotic species invasions undermine regional functional diversity of freshwater fish. *Sci. Rep.*, 9, 17921.

#### 48) N69FMI: Macroinvertebrates from the Tenojoki drainage basin, Finland – *Jani Heino*

**Data owner(s)**: Jani Heino

**Affiliation(s):** Finnish Environment Institute, Research Programme for Biodiversity, P.O. Box 413, FIN-90014 Oulu, Finland

**Description:** A total of 55 stream and river sites were sampled in June 2012.

**Geographical scope:** Finland

**Taxonomical scope:** Macroinvertebrates

**Ecological scope:** Freshwater (Lotic systems)

**Number of sampling units:** 55 sites in different-sized streams and rivers.

**Minimum distance between sampling units:** 0.14 kilometres

**Maximum distance between sampling units:** 142 kilometres

**Environmental variables:** Total nitrogen, color, manganese, pH, conductivity, sand, gravel, pebble, cobble, moss, velocity, depth, width.

**Functional traits:**

1. Functional feeding group: Collector-gatherer, filterer, predator, scraper, shredder
2. Habit trait groups: Burrowers, climbers, clingers, sprawlers, swimmers
3. Body size classes: 0-0.25cm, 0.25-0.5cm, 0.5-1.0cm, 1-2cm, 2-4cm, 4-8cm.

**Reference:** Kärnä, O.-M., Heino, J., Grönroos, M. & Hjort, J. (2018). The added value of geodiversity indices in explaining variation of stream macroinvertebrate diversity. *Ecol. Indic.*, 94, 420–429.

#### 49) N70TBI: Birds from Norway – *Miska Luoto*

**Data owner(s)**: Miska Luoto

**Affiliation(s):** University of Helsinki

**Description:** 420 sites, bird abundance information per sites, 50 m radius, year 2019

**Geographical scope:** Norway

**Taxonomical scope:** Birds

**Ecological scope:** Terrestrial (Tundra plants)

**Number of sampling units:** 420 sites (50 m radius)

**Minimum distance between sampling units:**

**Maximum distance between sampling units:** About 20 kilometres

**Environmental variables:** local solar radiation, slope angle, topographic wetness, topographic position, edaphic quality, fluvial disturbance, frost disturbance, snow cover duration

**Functional traits:** Body size, Wing span, Weight, Diet breadth

**Reference:**

#### 50) N70TBP: Bryophytes from Norway – *Pekka Niitynen and Miska Luoto*

**Data owner(s)**: Pekka Niittynen, Julia Kemppinen and Miska Luoto

**Affiliation(s):** Department of Geosciences and Geography, University of Helsinki

**Description:** Data comprising 145 bryophytes species sampled between 2014 and 2018 in sub-Arctic forests and mountain tundra along wide environmental gradients.

**Geographical scope:** Norway, Rastigaisa

**Taxonomical scope:** Bryophytes (mosses, growing on soil surfaces)

**Ecological scope:** Terrestrial

**Number of sampling units:** 1325 plots (4 x 1m^2^ each).

**Minimum distance between sampling units:** 0.02 kilometres

**Maximum distance between sampling units:** 19 kilometres

**Environmental variables:** local solar radiation, slope angle, topographic wetness index, topographic position, soil edaphic quality, fluvial disturbance, frost disturbance, snow cover duration.

**Functional traits:** General growth form (categorical), hairy leaf points (binary), branches (binary), shoot length (based on multiple floras; continuous).

**Reference:**

Niittynen, P., Heikkinen, R.K. & Luoto, M. (2018). Snow cover is a neglected driver of Arctic biodiversity loss. *Nat. Clim. Chang.*, 8, 997–1001.

Niittynen, P. & Luoto, M. (2018). The importance of snow in species distribution models of arctic vegetation. *Ecography (Cop.).*, 41, 1024–1037.

#### 51) N70TLC: Lichens from Norway – *Pekka Niitynen and Miska Luoto*

**Data owner(s)**: Pekka Niittynen, Julia Kemppinen and Miska Luoto

**Affiliation(s):** Department of Geosciences and Geography, University of Helsinki

**Description:** Data comprising 111 lichen species sampled between 2014 and 2018 in sub-Arctic forests and mountain tundra along wide environmental gradients.

**Geographical scope:** Norway, Rastigaisa

**Taxonomical scope:** Lichens (Ascomycota; soil dwelling species)

**Ecological scope:** Terrestrial

**Number of sampling units:** 1325 plots (4 x 1m^2^ each).

**Minimum distance between sampling units:** 0.02 kilometres

**Maximum distance between sampling units:** 19 kilometres

**Environmental variables:** local solar radiation, slope angle, topographic wetness index, topographic position, soil edaphic quality, fluvial disturbance, frost disturbance, snow cover duration.

#### 52) N70TTP: Tundra plants – Miska Luoto

**Data owner(s)**: Miska Luoto

**Affiliation(s):** University of Helsinki

**Description:** 1325 sites, plant cover information per 4 x 1-m2 square in mountain tundra

**Geographical scope:** Norway

**Taxonomical scope:** Vascular plant

**Ecological scope:** Terrestrial

**Number of sampling units:** 1325 plots (4m²)

**Minimum distance between sampling units:** 0.02 kilometres

**Maximum distance between sampling units:** 44.27 kilometres

**Environmental variables:** local solar radiation, slope angle, topographic wetness, topographic position, edaphic quality, fluvial disturbance, frost disturbance, snow cover duration.

**Functional traits:** The following traits were measured:

| *Functional traits* | *Unit* |
| --- | --- |
| Specific leaf area | m2kg-1 |
| Leaf N | mg g-1 |
| Leaf P | mg g-1 |
| Plant height | m |
| Leaf dry matter content | g g-1 |
| Seed mass | mg |
| Leaf area | mm2 |

**Reference:** Niittynen, P., Heikkinen, R.K. & Luoto, M. (2018). Snow cover is a neglected driver of Arctic biodiversity loss. *Nat. Clim. Chang.*, 8, 997–1001.

Niittynen, P. & Luoto, M. (2018). The importance of snow in species distribution models of arctic vegetation. *Ecography (Cop.).*, 41, 1024–1037.

Norberg, A., Abrego, N., Blanchet, F.G., Adler, F.R., Anderson, B.J., Anttila, J., *et al.* (2019). A comprehensive evaluation of predictive performance of 33 species distribution models at species and community levels. *Ecol. Monogr.*, 89.

#### 53) N78TTP: Plants from Svalbard – *Pekka Niitynen*

**Data owner(s)**: Pekka Niittynen, Julia Kemppinen and Miska Luoto

**Affiliation(s):** Department of Geosciences and Geography, University of Helsinki

**Description:** Data comprising 54 plant species

**Geographical scope:** Norway, Svalbard

**Taxonomical scope:** Terrestrial vascular plants

**Ecological scope:** Terrestrial

**Number of sampling units:** 72 plots

**Minimum distance between sampling units: <** 0.01 kilometres

**Maximum distance between sampling units:** 3.77 kilometres

**Environmental variables:** Within each plot, several abiotic and biotic variables were collected. Rock cover (%; i.e. the cover of rocks or boulders large enough to inhibit plant colonization and growth) was visually estimated. Soil depth (cm) was measured at four different points in each cell, and an average soil depth then calculated for the cell. Mesotopography was scored a value between 1 and 10, where 1 represents a valley bottom, 5 the middle of a slope and 10 the slope crest. Soil moisture (% volumetric water content) measurements were taken in each plot. Soil samples were taken from a minimum of 16 cells per grid and used to determine soil pH in the laboratory using the CaCl_2_ method (with pH linearly interpolated for all other cells within a grid). Slope angle (°) was measured in the field for each cell. Slope angle, aspect and latitude was subsequently used to calculate potential incident solar radiation (MJ · cm–2 · yr–1) for each cell (following McCune & Keon, 2002).

**Functional traits:** The following traits were measured

| *Functional traits* | *Unit* |
| --- | --- |
| Specific leaf area | m2kg-1 |
| Leaf N | mg g-1 |
| Leaf P | mg g-1 |
| Plant height | m |
| Leaf dry matter content | g g-1 |
| Seed mass | mg |
| Leaf area | mm2 |
| **References:** Unpublished. |  |

#### 54) S01MFI: Pacific fish fauna – *Michel Kulbiki, Alan Friedlander and Itai Granot.*

**Data owner(s)**: Michel Kulbiki^1^, Alan Friedlander^2^ and Itai Granot^3^.

**Affiliation(s):** ^1^IRD (Institut de Recherche pour le Développement), Laboratoire d’Excellence Labex Corail, UMR IRD-UR-CNRS ENTROPIE, Université de Perpignan, 66860 Perpignan Cedex 9, France; ^2^Department of Biology, University of Hawaii, Honolulu, HI 96822, USA; ^3^School of Zoology, George S. Wise Faculty of Life Sciences, Tel Aviv University, Tel Aviv, Israel.

**Description:** Reef fish abundance data gathered from nearly 6000 UVCs (Underwater Visual Census Surveys) from 85 sites across the Pacific Ocean. Environmental data extracted from Bio-ORACLE.

**Geographical scope:** Pacific Ocean

**Taxonomical scope:** Fish

**Ecological scope:** Marine (Ocean)

**Number of sampling units:** 85 plots (250 m²)

**Minimum distance between sampling units:** < 0.01 kilometres

**Maximum distance between sampling units:** 17434 kilometres

**Environmental variables** - all variables where averaged across a 15 km buffer area around the sampled site: (1) mean sea surface temperature (SST), (2) temperature (SST) range, (3) salinity - mean sea water salinity at the bottom at mean bottom depth, (4) mean Chlorophyll - mean mass concentration of chlorophyll in sea water at minimum bottom depth	, (5) Chlorophyll range - range of the mass concentration of chlorophyll in sea water at minimum bottom depth

**Functional traits:** R Dataframe *sp_traits* with 6 traits: Size, Mobility, Activity, Schooling, Position, Diet.

- *Size* coded as an ordered categorical variable with 6 levels:

1= 0-7cm (S1)

2= 7.1-15cm (S2)

3= 15.1- 30cm (S3)

4= 30.1-50cm (S4)

5= 50.1-80cm (S5)

6= >80cm (S6)

- *Mobility* (=Home range) coded as an ordered categorical variable, 3 levels:

1= sedentary (Sed)

2= mobile within a reef (Mob)

3= highly mobile i.e. between reefs (VMob)

- Period of *Activity* coded as an ordered categorical variable, 3 levels:

1= diurnal (Day)

2= diurnal & nocturnal (Both)

3= nocturnal (Night)

- *Schooling* coded as a categorical variable, 5 levels:

1= solitary (Sol)

2= pairing (Pair)

3= small group (SmallG)

4= medium group (MedG)

5= large group (LargeG)

- *Position* in the water column (=Level-water) coded as an ordered categorical variable, 3 levels:

1= bottom (Bottom)

2= above bottom (Low)

3= pelagic (High)

*- Diet* coded as a categorical variable, 7 levels:

HD= herbivorous-detritivorous (undefined organic material, often grouped by many authors under the name detritus and/or undefined vegetal material, turf or filamentous algae)

HM= herbivorous macro-algal (macro algae (large fleshy algae) and sea grass)

IS= invertivorous sessile (sessile invertebrates: coral, sponge, ascidians and so on)

IM= invertivorous mobile (large benthic invertebrates + small benthic invertebrates + undefined invertebrates)

PK= planktonivorous (plankton and small organisms which migrate in the water column, such as many benthic copepods, amphipods, crustacean larvaes etc which migrate in the water column at night)

FC= Pelagic macro-organisms (large organisms living in the water column, usually fish and cephalopods) and benthic fish

OM= omnivorous (herbivorous and/or detritivorous AND carnivorous)

**Reference:** Unpublished

#### 55) S03FBD Diatoms from Kenya – Jenny Jyrkänkallio-Mikkola et al.

**Data owner(s):** Jenny Jyrkänkallio-Mikkola^1^ and Janne Soininen^2^.

**Affiliation(s):** ^1^ WWF Finland; ^2^Department of Geosciences and Geography, University of Helsinki, Helsinki, Finland

**Description:** Benthic diatom data and supporting environmental dataset

**Geographical scope:** Kenya - Taita Hills

**Taxonomical scope:** Benthic diatoms

**Ecological scope:** Freshwater (rivers)

**Number of sampling units:** 67

**Minimum distance between sampling units:** 0.54 kilometres

**Maximum distance between sampling units:** 63 kilometres

**Environmental variables:** coordinates, elevation (m), water temperature (, conductivity (mSm^-1^), pH, total nitrogen (mgl^-1^), water colour (Pt mgl^-1^), stream width (cm), stream depth (cm), current velocity (ms^-1^), shading of the surrounding vegetation (%), percentages of moss, sand, gravel, pebble, cobble, boulders and bedrock.

**Functional traits:** Life‐forms (mobile, adnate, pedunculate, pad-attached, stalk-attached, colonial; Rimet & Bouchez 2012), ecological guilds (high-profile, low-profile, motile; Passy 2007; Rimet & Bouchez 2012)

**Reference**: Jyrkänkallio-Mikkola, J., Siljander, M., Heikinheimo, V., Pellikka, P. & Soininen, J. (2018). Tropical stream diatom communities – The importance of headwater streams for regional diversity. *Ecol. Indic.*, 95, 183–193.

#### 56) S07MFI: Atlantic fish fauna – Sergio R. Floeter, Carlos Eduardo Leite Ferreira and Itai Granot

**Data owner(s)**: Sergio R. Floeter^1^ and Carlos Eduardo Leite Ferreira^2^ and Itai Granot

**Affiliation(s):** ^1^Marine Macroecology and Biogeography Lab, Depto. de Ecologia e Zoologia, CCB, Universidade Federal de Santa Catarina, Florianopolis, SC, 88040-900, Brazil; ^2^ Universidade Federal Fluminense (UFF), Departamento de Biologia Marinha, Niterói, Brazil; ^3^ School of Zoology, George S. Wise Faculty of Life Sciences, Tel Aviv University, Tel Aviv, Israel.

**Description:** Reef fish abundance data gathered from nearly 5,500 UVCs (Underwater Visual Census Surveys) from 202 sites across the Atlantic Ocean.

Environmental data extracted from Bio-ORACLE.

**Geographical scope:** Atlantic fish

**Taxonomical scope:** Fish

**Ecological scope:** Marine (Ocean)

**Number of sampling units:** 202 plots (40m²)

**Minimum distance between sampling units:** < 0.01 kilometres

**Maximum distance between sampling units:** 10538 kilometres

**Environmental variables** - all variables where averaged across a 15 km buffer area around the sampled site**:** (1) mean sea surface temperature (SST), (2) temperature (SST) range, (3) salinity - mean sea water salinity at the bottom at mean bottom depth, (4) mean Chlorophyll - mean mass concentration of chlorophyll in sea water at minimum bottom depth	, (5) Chlorophyll range - range of the mass concentration of chlorophyll in sea water at minimum bottom depth

F**unctional traits:** R Dataframe *sp_traits* with 6 traits: Size, Mobility, Activity, Schooling, Position, Diet.

- *Size* coded as an ordered categorical variable with 6 levels:

1= 0-7cm (S1)

2= 7.1-15cm (S2)

3= 15.1- 30cm (S3)

4= 30.1-50cm (S4)

5= 50.1-80cm (S5)

6= >80cm (S6)

- *Mobility* (=Home range) coded as an ordered categorical variable, 3 levels:

1= sedentary (Sed)

2= mobile within a reef (Mob)

3= highly mobile i.e. between reefs (VMob)

- Period of *Activity* coded as an ordered categorical variable, 3 levels:

1= diurnal (Day)

2= diurnal & nocturnal (Both)

3= nocturnal (Night)

- *Schooling* coded as a categorical variable, 5 levels:

1= solitary (Sol)

2= pairing (Pair)

3= small group (SmallG)

4= medium group (MedG)

5= large group (LargeG)

- *Position* in the water column (=Level-water) coded as an ordered categorical variable, 3 levels:

1= bottom (Bottom)

2= above bottom (Low)

3= pelagic (High)

*- Diet* coded as a categorical variable, 7 levels:

HD= herbivorous-detritivorous (undefined organic material, often grouped by many authors under the name detritus and/or undefined vegetal material, turf or filamentous algae)

HM= herbivorous macro-algal (macro algae (large fleshy algae) and sea grass)

IS= invertivorous sessile (sessile invertebrates: coral, sponge, ascidians and so on)

IM= invertivorous mobile (large benthic invertebrates + small benthic invertebrates + undefined invertebrates)

PK= planktivorous (plankton and small organisms which migrate in the water column, such as many benthic copepods, amphipods, crustacean larvaes etc which migrate in the water column at night)

FC= Pelagic macro-organisms (large organisms living in the water column, usually fish and cephalopods) and benthic fish

OM= omnivorous (herbivorous and/or detritivorous AND carnivorous)

**References:** Barneche, D.R., Rezende, E.L., Parravicini, V., Maire, E., Edgar, G.J., Stuart-Smith, R.D., *et al.* (2019). Body size, reef area and temperature predict global reef-fish species richness across spatial scales. *Glob. Ecol. Biogeogr.*, 28, 315–327.

Morais, R.A., Ferreira, C.E.L. & Floeter, S.R. (2017). Spatial patterns of fish standing biomass across Brazilian reefs. *J. Fish Biol.*, 91, 1642–1667.

Quimbayo, J.P., Dias, M.S., Kulbicki, M., Mendes, T.C., Lamb, R.W., Johnson, A.F., *et al.* (2019). Determinants of reef fish assemblages in tropical Oceanic islands. *Ecography (Cop.).*, 42, 77–87.

#### 57) S14FPP (1-2): Phytoplankton from Brazil das Águas Project –*Vera Lucia de Moraes Huszar*

**Data owner(s)**: Vera Lucia de Moraes Huszar

**Affiliation(s):** Phycology Laboratory, Botany Department, National Museum, Federal University of Rio de Janeiro, Brasil.

**Description:** Includes phytoplankton species (morphospecies) in abundance (individuals/mL and cels/mL), richness and diversity

**Geographical scope:** Brazil

**Taxonomical scope:** Phytoplankton

**Ecological scope:** Freshwaters and brakish systems.

1. Lentic
2. Lotic

**Number of sampling units:** 281 georeferenced samples from river (183) and lakes (98) in the Brazilian hydrographic coastal basins in a latitudinal gradient from a 03^o^47`51¨N and 33^o^29`53¨S

**Minimum distance between sampling units:** 1.10 kilometres

**Maximum distance between sampling units:** ca. 4100 km overland straightline

**Environmental variables:** Water temperature, pH, turbidity, dissolved oxygen, dissolved inorganic nitrogen, soluble reactive phosphorus, total phosphorus.

**Functional traits:** In the laboratory we measured the maximum linear dimension (µm) of 20 individuals from each species whenever possible. We also observed the presence or absence of flagella, mucilage and aerotopes.

**Reference:** Unplished data

#### 58) S22FPP: Phytoplankton from Paraiba do Sul river, Brazil – *Felipe pacheco et al.*

**Data owner(s)**: Felipe Siqueira Pacheco^1^, Marcela Miranda^1^ and Jean Ometto^1^.

**Affiliation(s):** ^2^ National Institute for Space Research.

**Description:** Phytoplankton community and Limnological data from sites each 5km along the full length of the Paraíba do Sul River – RJ, Brazil.

**Geographical scope:** Brazil

**Taxonomical scope:** Phytoplankton

**Ecological scope:** Freshwaters (River)

**Number of sampling units:** 206

**Minimum distance between sampling units:** 0.25 kilometres

**Maximum distance between sampling units:** 1136 kilometres

**Environmental variables:** Water: Temperature (Cº), Conductivity, Dissolved Oxygen (mg L^-1^), pH, Dissolved Inorganic Carbon (mg L^-1^), Total Organic Carbon (mg L^-1^), Nitrogen (mg L^-1^), Phosphorus (mg L^-1^), Heavy metals (mg L^-1^).

**Functional traits:** In the laboratory we measured the area (µm²), volume (µm³), the maximum linear dimension (µm) of 20 individuals from each species whenever possible. We also observed the presence or absence of flagella, mucilage and aerotopes. Lastly, we calculated the ratio between the area and volume of each species and used as a trait.

**Reference:** Pacheco, F.S., Miranda, M., Pezzi, L.P., Assireu, A., Marinho, M.M., Malafaia, M., *et al.* (2017). Water quality longitudinal profile of the Paraíba do Sul River, Brazil during an extreme drought event. *Limnol. Oceanogr.*, 62, S131–S146.

#### 59) S47TTP: Marion Island fine-scale grids – *Peter le Roux and Mia Momberg*

**Data owner(s)**: Peter le Roux and Mia Momberg

**Affiliation(s):** Department of Plant and Soil Sciences, University of Pretoria

**Description:** Vascular plant cover and identity and several abiotic environmental variables were recorded in nine grids of 8 x 20 m each were sampled across two adjacent landforms on the eastern section of sub-Antarctic Marion Island, each comprising 160 contiguous cells of 1 m^2^ size. For these analyses, cells at 4 m intervals were used in analyses.

Grids were located to sample the full range of conditions within the site, covering the range of vegetation types and topography present. Individual grids, in turn, were orientated to encompass as much environmental variability as possible locally. This sampling replicates protocols applied in the Arctic-alpine tundra of northern Finland (le Roux *et al.,* 2013)

**Geographical scope:** Sub-Antarctic Marion Island (Southern Ocean)

**Taxonomical scope:** Vascular plants

**Ecological scope:** Terrestrial

**Number of sampling units:** 180 (1 m^2^ each)

**Minimum distance between sampling units:** 4 meters

**Maximum distance between sampling units:** 940 m

**Environmental variables:** Within each of the 1 m^2^ cells several abiotic and biotic variables were collected. Rock cover (%; i.e. the cover of rocks or boulders large enough to inhibit plant colonization and growth) was visually estimated. Soil depth (cm) was measured at four different points in each cell, and an average soil depth then calculated for the cell (where soil depth exceeded 0.6 m, a value of 0.7 m was assigned). Mesotopography was scored a value between 1 and 10, where 1 represents a valley bottom, 5 the middle of a slope and 10 the slope crest. Soil moisture (% volumetric water content) measurements were taken in each cell in April, June, July and October 2016 and January 2017. These instantaneous readings were only taken on days where there had been at least 24 hours since the last rainfall event. Soil samples were taken from a minimum of 16 cells per grid and used to determine soil pH in the laboratory using the CaCl_2_ method (with pH linearly interpolated for all other cells within a grid). Slope angle (°) was measured in the field for each cell. Slope angle, aspect and latitude was subsequently used to calculate potential incident solar radiation (MJ · cm–2 · yr–1) for each cell (following McCune & Keon, 2002).

**Functional traits:** Plant height (m), leaf dry matter content (g/g), leaf area (mm²), leaf nitrogen concentration (mg/g), leaf phosphorus concentration (mg/g), seed mass (mg), specific leaf area (mm²/mg) (these traits are all continuous variables; measured following the protocols of Pérez-Harguindeguy et al., 2013).

**Reference:** McCune, B. & Keon, D. (2002). Equations for potential annual direct incident radiation and heat load. *J. Veg. Sci.*, 13, 603–606.

Pérez-Harguindeguy, N., Díaz, S., Garnier, E., Lavorel, S., Poorter, H., Jaureguiberry, P., *et al.* (2013). New handbook for standardised measurement of plant functional traits worldwide. *Aust. J. Bot.*

le Roux, P.C., Lenoir, J., Pellissier, L., Wisz, M.S. & Luoto, M. (2013). Horizontal, but not vertical, biotic interactions affect fine-scale plant distribution patterns in a low-energy system. *Ecology*, 94, 671–682.

#### 60) S62MPP: Phytoplankton from Antarctica – *Gleyci A. O. Moser, Domênica T. Lima and José J. Barrera-Alba*

**Data owner(s)**: Gleyci Aparecida Oliveira Moser^1^, Domênica Texeira de Lima^1^, José Juan Barrera-Alba^2^

**Affiliation(s):** ^1^ Rio de Janeiro State University; ^2^ Federal University of Sao Paulo.

**Description:** Dataset comprising oceanographic variables, and abundances and traits of 109 species of marine phytoplankton of coastal marine waters from Antarctica.

**Geographical scope:** Admiralty Bay, King George Island, South Shetland Islands, Western Antarctic Peninsula.

**Taxonomical scope:** Diatoms, dinoflagellates, cyanobacteria and chlorophytes.

**Ecological scope:** Marine, coastal waters (Polar region)

**Number of sampling units:** 30

**Minimum distance between sampling units:** 0.01 kilometres

**Maximum distance between sampling units:** 0.10 kilometres

**Environmental variables:** salinity, temperature, dissolved inorganic nutrients (nitrate, phosphate, silicate), % of Melting Water (MW)

**Functional traits:** GALD, greatest axial linear dimension; S.V, surface to volume ratio; VAC, presence of large vacuoles; SIL, presence of silica structure; COL, coloniality or chain formation; RAF, presence of raphe; HABs, Harmful algal bloom formation capacity; AUT, autotrophy; HET, heterotrophy; MIX, mixotrophy; CYST, resting stages; FLAG, presence of flagella.

**Reference:**

Lima, D.T. de, Moser, G.A.O., Piedras, F.R., Cunha, L.C. da, Tenenbaum, D.R., Tenório, M.M.B., *et al.* (2019). Abiotic Changes Driving Microphytoplankton Functional Diversity in Admiralty Bay, King George Island (Antarctica). *Front. Mar. Sci.*, 6.

Litchman, E. & Klausmeier, C.A. (2008). Trait-Based Community Ecology of Phytoplankton. *Annu. Rev. Ecol. Evol. Syst.*, 39, 615–639.

### III. Database mobilized in collaboration with the sPlot database

The datasets from sPlot were compiled by Francesco Maria Sabatini. Specifically, for these datasets, trait information was drawn from the TRY database (v 3.0), following the following steps:

First, species names were benchmaked in original datasets sPlot 2.1 backbone. Hence, we used the names in the backbone to match the corresponding entry in the gap-filled trait matrix from TRY. Finally, we derived a standard set of traits (Table 1.) for all dataset from sPlot.

For more information about data curation and standardization please click [here](https://www.try-db.org/TryWeb/Database.php).

Table 1. Set of traits drawn for TRY database, with respective abbreviations used in the dataset, the code in the thesaurus of plants (TOP), and units of measurements.

| Trait | Abbreviation in dataset | Top | Unit |
| --- | --- | --- | --- |
| *Stem specific density* | SSD | 286 | g cm^-3^ |
| *Specific leaf area* | SLA | 50 | m^2^ kg^-1^ |
| *Leaf n* | LeafN | 462 | mg g-1 |
| *Plant height* | Plant.Height | 68 | m |
| *Seed mass* | Seed.Mass | 103 | mg |
| *Seed number per reproductive unit* | Seed.Num.Rep.Unit | - |  |

#### 61) N05TTP: Cameroon Tropical Forest Vegetation – *Jiri Dolezal and Jan Altman*

**Data owner(s)**: Jiri Dolezal¹² and Jan Altman¹

**Affiliation(s):** ^1^Institute of Botany of The Czech Academy of Sciences, Zámek 1, 25243 Průhonice, Czech Republic;

^2^Faculty of Science, Department of Botany, University of South Bohemia, Na Zlaté Stoce 1, 37005 České Budějovice, Czech Republic

**Description:** 10628 tree individuals measured of more than 250 tree species, covering all major forest types from sea level to 3000 masl.

**Geographical scope:** Cameroon/Africa

**Taxonomical scope:** Primary forest

**Ecological scope:** Terrestrial

**Number of sampling units:** 167 plots (1000m²)

**Minimum distance between sampling units:**

**Maximum distance between sampling units:** 50 kilometres

**Environmental variables:** Soil chemistry (pH, texture, N and P nutrients, Mg and Ca ions (ppm)), Potential Annual Direct Incident Radiation (MJ cm-2 yr-1), gap area (m2), temperature (°C), precipitation (mm)

**Functional traits:** Wood density (kg/m3), DBH - stem diameter (cm), tree height (m).

**Reference:** Hořák, D., Ferenc, M., Sedláček, O., Motombi, F.N., Svoboda, M., Altman, J., *et al.* (2019). Forest structure determines spatial changes in avian communities along an elevational gradient in tropical Africa. *J. Biogeogr.*, 46, 2466–2478.

#### 62) N33TTP_2: Himalayan Alpine Vegetation – *Jiri Dolezal and Jan Altman*

**Data owner(s)**: Jiri Dolezal¹² and Jan Altman¹

**Affiliation(s):** ^1^Institute of Botany of The Czech Academy of Sciences, Zámek 1, 25243 Průhonice, Czech Republic;

^2^Faculty of Science, Department of Botany, University of South Bohemia, Na Zlaté Stoce 1, 37005 České Budějovice, Czech Republic;

**Description:** 370 plots covering main vegetation types (semi-deserts, steppes, shrublands, alpine meadows, saline wetlands, subnival) from 3800 to 6000 m elevation, with more than 270 plant species.

**Geographical scope:** Ladakh/NW Himalaya, India

**Taxonomical scope:** Primary mountain vegetation

**Ecological scope:** Terrestrial

**Number of sampling units:** 370 plots (100 m²)

**Minimum distance between sampling units:**

**Maximum distance between sampling units:** 100 kilometres

**Environmental variables:** Soil conditions, mean annual temperature and precipitation.

**Functional traits:**

**Plant clonality**: 22 clonal growth forms based on Klimešova et al. 2011.

Ecological Indicator Values: stability (1-3) nutrients (1-3), light (1-3), soil moisture (1-4).

**Plant size**: CanopyHeight [m].

**Plant traits**: LDMC - leaf dry matter content (mg/g), LNC - leaf N concentration (%), LCC - leaf carbon (%), LPC - leaf phosphorus (%), RNC - root nitrogen (%), δ^15^N - nitrogen stable isotope (‰), δ^13^C - carbon stable isotope (‰), RPC - root phosphorus (%), Suma_sugar_root (%), Fructan_root (%), Starch_root (%), Free_sugar_root (%), Vessel.diameter.no (µm) , Lignification - lignified tissue proportion on root collar cross-sections (%).

LDMC (mg/g), Seed.mass (mg), SLA (mm^2^/mg), non-woody (%), woody (%), terminal velocity (m/s^2^)

**Reference:** Dolezal, J., Dvorsky, M., Kopecky, M., Altman, J., Mudrak, O., Capkova, K., *et al.* (2019). Functionally distinct assembly of vascular plants colonizing alpine cushions suggests their vulnerability to climate change. *Ann. Bot.*, 123, 569–578.

Dolezal, J., Dvorsky, M., Kopecky, M., Liancourt, P., Hiiesalu, I., Macek, M., *et al.* (2016). Vegetation dynamics at the upper elevational limit of vascular plants in Himalaya. *Sci. Rep.*, 6, 24881.

Klimešová, J., Doležal, J., Dvorský, M., de Bello, F. & Klimeš, L. (2011). Clonal Growth Forms in Eastern Ladakh, Western Himalayas: Classification and Habitat Preferences. *Folia Geobot.*, 46, 191–217.

#### 63) N38TTP: Dry grassland vegetation of Sicily, Italy – *EDGG*

**Data owner(s)**:

Eurasian Dry Grassland Group (EDGG; [www.edgg.org](http://www.edgg.org)), represented by the

- GrassPlot database through Idoia Biurrun^1^ and Jürgen Dengler^2^
- Local Organising Committee of the 4th EDGG Field Workshop through Riccardo Guarino^3^

**Affiliation(s):** ^1^Department of Plant Biology and Ecology, University of the Basque Country UPV/EHU, P.O. Box 644, 48080 Bilbao, Spain; ^2^Vegetation Ecology, Institute of Natural Resource Management (IUNR), Zurich University of Applied Sciences (ZHAW), Grüentalstr. 14, 8820 Wädenswil, Switzerland; ^3^Department STEBICEF – Botanical Unit, University of Palermo, via Archirafi, 38, 90123 Palermo, Italy

**E-mail**:

**Description:** Data from the 4th EDGG Field Workshop on sampling multi-scale plant diversity data in Palaearctic grasslands, conducted 2012 in Sicily, Italy (GrassPlot ID IT_A; Guarino et al. 2012). The EDGG Field Workshops use a standardized sampling approach (Dengler et al. 2016), and their data are stored in the collaborative GrassPlot database (Dengler et al. 2018, 2020; Biurrun et al. 2019). In this paper only the vascular plant data from the 10-m² plots were used.

**Geographical scope:** Sicily, Italy

**Taxonomical scope:** Vascular plants

**Ecological scope:** Terrestrial (Dry grasslands s.l.)

**Number of sampling units:** 67 (10 m²)

**Minimum distance between sampling units:** 0.01 km

**Maximum distance between sampling units:** 226 km

**Environmental variables:** Elevation (m a.s.l.), aspect (°), inclination (°), heat load index, maximum microrelief (cm), soil depth mean (cm), soil depth standard deviation (cm), skeleton content (mass %; for uppermost 15 cm of soil), pH (measured in H2O; for uppermost 15 cm of soil), electrical conductivity (μS/cm; for uppermost 15 cm of soil); sand content (%; for uppermost 15 cm of soil, approximative from soil texture class); silt content (%; for uppermost 15 cm of soil, approximative from soil texture class); clay content (%; for uppermost 15 cm of soil, approximative from soil texture class); carbonate content (%; for uppermost 15 cm of soil, approximative from ordinal scale); organic matter (%; for uppermost 15 cm of soil)

**Functional traits:**

**References:**

Biurrun, I., Burrascano, S., Dembicz, I., Guarino, R., Kapfer, J., Pielech, R., Garcia-Mijangos, I., Wagner, V., Palpurina, S., (…) & Dengler, J. 2019. GrassPlot v. 2.00 – first update on the database of multi-scale plant diversity in Palaearctic grasslands. *Palaearctic Grasslands* 44: 26–47.

Dengler, J., Boch, S., Filibeck, G., Chiarucci, A., Dembicz, I., Guarino, R., Henneberg, B., Janišová, M., Marcenò, C., (…) & Biurrun, I. 2016. Assessing plant diversity and composition in grasslands across spatial scales: the standardised EDGG sampling methodology. *Bulletin of the Eurasian Grassland Group* 32: 13−30.

Dengler, J., Wagner, V., Dembicz, I., García-Mijangos, I., Naqinezhad, A., Boch, S., Chiarucci, A., Conradi, T., Filibeck, G., (…) & Biurrun, I. 2018. GrassPlot – a database of multi-scale plant diversity in Palaearctic grasslands. *Phytocoenologia* 48: 331–347.

Dengler, J., Matthews, T.J., Steinbauer, M.J., Wolfrum, S., Boch, S., Chiarucci, A., Conradi, T., Dembicz, I., Marcenò, C., (…) & Biurrun, I. 2020. Species-area relationships in continuous vegetation: Evidence from Palaearctic grasslands. *Journal of Biogeography* 60: 72–86.

Guarino, R., Becker, T., Dembicz, I., Dolnik, C., Kącki, Z., Kozub, Ł., Rejžek, M. & Dengler, J. 2012. Impressions from the 4th EDGG Research Expedition to Sicily: community composition and diversity of Mediterranean grasslands. *Bulletin of the European Dry Grassland Group* 15: 12–22.

#### 64) N39TTP: Vegetation of Middle Asia – *Arkadiusz Nowak*

**Data owner(s):** Arkadiusz Nowak

**Affiliation(s):** ^1,^ Polish Academy of Sciences Botanical Garden - Center for Biological Diversity Conservation in Powsin, Prawdziwka 2, 02-973 Warsaw, Poland;

^2^University of Opole, Institute of Biology, Oleska 22, 45-052 Opole, Poland;

**Description:** 20 10m^2^ pairs of plots (40 plots) in Tajikistan from ca. 500 to 4000 m, and 7 pairs of 10m2 pairs of plots (14 plots) in Kyrgyzstan with soil data, coordinates and typical header data for releves.

**Geographical scope:** Middle Asia (Tajikistan, Kyrgyzstan)

**Taxonomical scope:** grasslands (steppes, meadows)

**Ecological scope:** Terrestrial

**Number of sampling units:** 54 plots (10 m^2^)

**Minimum distance between sampling units:**

**Maximum distance between sampling units:** approx. 900 km

**Environmental variables:** Elevation [m a.s.l.], Longitude original [°], Latitude original [°], Relief position, [verbal], Parental geological substrate, Aspect [°], Inclination [°], Heat load index, Max. microrelief [cm], Soil depth in 5 points [cm], Mean height herb layer [m], Land use [verbal], Land use category, Cover tree layer [%], Cover shrub layer [%], Cover herb layer [%], Cover cryptogam layer [%], Total vegetation cover [%], Maximum height tree layer [m], Maximum height shrub layer [m], Maximum height herb layer [m], Cover litter [%], Cover dead wood [%], Cover stones and rocks [%], Cover gravel [%], Cover fine soil [%], Bare ground [%],pH H2O (only uppermost layer), pH CaCl2 (only uppermost layer), pH KCl (only uppermost layer), EC (µS/cm), Soil N content (%), Soil C content (%), C/N ratio, Soil texture class, Sand: % from soil texture class, Silt: % from soil texture class, Clay: % from soil texture class, Carbonate %, K: 1/0, P: 1/0, Organic matter, Soil water content (%) = (fresh soil weight (g) - dry soil weight (g)/ fresh soil weight (g) x 100, soil organic C (%), Available P (mg/100g), Available K (mg/100g)

**Functional traits:** Please see the section III

**References:**

#### 65) N41TTP: Dry grassland vegetation of Northern Greece – *EDGG*

**Data owner(s)**:

Eurasian Dry Grassland Group (EDGG; [www.edgg.org](http://www.edgg.org)), represented by the

- GrassPlot database through Idoia Biurrun^1^ and Jürgen Dengler^2^
- Local Organising Committee of the 5th EDGG Field Workshop through Ioannis Tsiripidis^3^

**Affiliation(s):** ^1^Department of Plant Biology and Ecology, University of the Basque Country UPV/EHU, P.O. Box 644, 48080 Bilbao, Spain; ^2^Vegetation Ecology, Institute of Natural Resource Management (IUNR), Zurich University of Applied Sciences (ZHAW), Grüentalstr. 14, 8820 Wädenswil, Switzerland; ^3^Department of Botany, School of Biology, Aristotle University of Thessaloniki, 54124 Thessaloniki, Greece

**Description:** Data from the 5th EDGG Field Workshop on sampling multi-scale plant diversity data in Palaearctic grasslands, conducted 2012 in Northern Greece (GrassPlot ID GR_A; Dengler & Demina 2012). The EDGG Field Workshops use a standardized sampling approach (Dengler et al. 2016), and their data are stored in the collaborative GrassPlot database (Dengler et al. 2018, 2020; Biurrun et al. 2019). In this paper only the vascular plant data from the 10-m² plots were used.

**Geographical scope:** Northern Greece

**Taxonomical scope:** Vascular plants

**Ecological scope:** Terrestrial (Dry grasslands s.l.)

**Number of sampling units:** 31 (10 m²)

**Minimum distance between sampling units:** 0.01 km

**Maximum distance between sampling units:** 268 km

**Environmental variables:** Elevation (m a.s.l.), aspect (°), inclination (°), maximum microrelief (cm), soil depth mean (cm), soil depth standard deviation (cm), skeleton content (mass %; for uppermost 15 cm of soil), pH (measured in H2O; for uppermost 15 cm of soil), pH (measured in CaCl2; for uppermost 15 cm of soil), electrical conductivity (μS/cm; for uppermost 15 cm of soil); organic matter (%; for uppermost 15 cm of soil)

**Functional traits:** Please see the section III

**References:** Biurrun, I., Burrascano, S., Dembicz, I., Guarino, R., Kapfer, J., Pielech, R., *et al.* (2019). GrassPlot v. 2.00: first update on the database of multi-scale plant diversity in Palaearctic grasslands. *Palaearct. Grasslands*, 44, 26–47.

Dengler, J. & Demina, O. (2012). 5th EDGG Research Expedition to Northern Greece, May 2012. *Bull. Eur. Dry Grassl. Gr.*, 16, 18–20.

Dengler, J., Boch, S., Filibeck, G., Chiarucci, A., Dembicz, I., Guarino, R., *et al.* (2016). Assessing plant diversity and composition in grass ‐ lands across spatial scales: the standardised EDGG sampling methodology. *Bullettin Eurasian Dry Grassl. Gr.*

Dengler, J., Wagner, V., Dembicz, I., García-Mijangos, I., Naqinezhad, A., Boch, S., *et al.* (2018). GrassPlot – a database of multi-scale plant diversity in Palaearctic grasslands. *Phytocoenologia*, 48, 331–347.

Dengler, J., Matthews, T.J., Steinbauer, M.J., Wolfrum, S., Boch, S., Chiarucci, A., *et al.* (2020). Species–area relationships in continuous vegetation: Evidence from Palaearctic grasslands. *J. Biogeogr.*

#### 66) N42TTP: Plant data from the Lazio region, Italy – *Marta Carboni and Alicia T. R. Acosta.*

**Data owner(s)**: Marta Carboni and Alicia T. R. Acosta

**Affiliation(s):** Department of Science, University of Roma Tre - Viale Guglielmo Marconi 446, I-00146, Rome, Italy

**Description:** Includes plot-species matrix with visually estimated cover, local environmental variable matrix for the plots and plot coordinates (on 6 main sites along the coast of Lazio).

**Geographical scope:** Italy, Lazio region

**Taxonomical scope:** Coastal dune vegetation (mainly herbaceous, including beach, embryo- and mobile dunes and interdune grasslands, but also some backdune woody vegetation)

**Ecological scope:** Terrestrial

**Number of sampling units:** 62 plots (4 m²)

**Minimum distance between sampling units:** 500 m

**Maximum distance between sampling units:** 205 kilometres

**Environmental variables:** Soil: granulometry (micron), content of organic matter (%), soil moisture (g), pH and conductivity (salinity, µS/cm)
Wind: Erosion of tatter flags (cm2), aerosol (µS/cm), sand burial (cm)

**Functional traits:** Please see the section III

**Reference:** Carboni M., Santoro R. & Acosta A.T. R. 2011. Dealing with scarce data to understand how environmental gradients and propagule pressure shape fine-scale alien distribution patterns on coastal dunes. Journal of Vegetation Science 22: 751–765

#### 67) N42TTP_2: Quercus suber forest of Italy – *Michele de Sanctis and Fabio Attorre*

**Data owner(s):** Michele de Sanctis^1^, Fabio Attorre^1^

**Affiliation(s):** Department of Environmental Biology, University Sapienza of Rome, P. le Aldo Moro 5, 00185, Rome, Italy.

**Description:** Soil analysis, phytosociological relevés, presence of disturb assessment and renovation index have been carried out in 2007-2008 in Quercus suber forest of Central Italy.

**Geographical scope:** Central Italy (Lazio and Tuscany region)

**Taxonomical scope:** Archeophyta (Quercus suber forests)

**Ecological scope:** Terrestrial (Forest)

**Number of sampling units:** 43 (314m2, i.e., round plot with 10m radius)

**Minimum distance between sampling units:**

**Maximum distance between sampling units:** 387 kilometres

**Environmental variables:** Ph, Organic carbon content (gr7100gr), C, N, C/N, Ca, Mg, K, Na, Base saturation, assimilable P, Phosphate, Silt %, Clay %, Sand (Fine, Medium, Coars) %, Skeleton

**Reference**:

#### 68) N42TTP_3: Dry grassland vegetation of the Central Apennines, Italy – *EDGG*

**Data owner(s)**:

Eurasian Dry Grassland Group (EDGG; [www.edgg.org](http://www.edgg.org)), represented by the

- GrassPlot database through Idoia Biurrun^1^ and Jürgen Dengler^2^
- Local Organising Committee of the 10th EDGG Field Workshop through Goffredo Filibeck^3^

**Affiliation(s):** ^1^Department of Plant Biology and Ecology, University of the Basque Country UPV/EHU, P.O. Box 644, 48080 Bilbao, Spain; ^2^Vegetation Ecology, Institute of Natural Resource Management (IUNR), Zurich University of Applied Sciences (ZHAW), Grüentalstr. 14, 8820 Wädenswil, Switzerland; ^3^Department of Agricultural and Forest Sciences, University of Tuscia, 01100 Viterbo, Italy

**Description:** Data from the 10th EDGG Field Workshop on sampling multi-scale plant diversity data in Palaearctic grasslands, conducted 2012 in Sicily, Italy (GrassPlot ID IT_L; Filibeck et al. 2018). The EDGG Field Workshops use a standardized sampling approach (Dengler et al. 2016), and their data are stored in the collaborative GrassPlot database (Dengler et al. 2018, 2020; Biurrun et al. 2019). In this paper only the vascular plant data from the 10-m² subplots within the nested-plot series were used.

**Geographical scope:** Central Apennines, Italy

**Taxonomical scope:** Vascular plants

**Ecological scope:** Terrestrial (Dry grasslands s.l.)

**Number of sampling units:** 40 (10 m²)

**Minimum distance between sampling units:** 0.01 km

**Maximum distance between sampling units:** 70 km

**Environmental variables:** Elevation (m a.s.l.), aspect (°), inclination (°), maximum microrelief (cm), soil depth mean (cm), soil depth standard deviation (cm), skeleton content (mass %; for uppermost 15 cm of soil), pH (measured in H2O; for uppermost 15 cm of soil), N content (%; for uppermost 15 cm of soil), C/N ratio (for uppermost 15 cm of soil), sand content (%; for uppermost 15 cm of soil); silt content (%; for uppermost 15 cm of soil); clay content (%; for uppermost 15 cm of soil); cation exchange capacity (cmol/kg; for uppermost 15 cm of soil), organic matter (%; for uppermost 15 cm of soil), organic C content (%; for uppermost 15 cm of soil); plant available P content (mg/100 g; for uppermost 15 cm of soil); plant available K content (mg/100 g; for uppermost 15 cm of soil)

**Functional traits:** Please see the section III

**References:**

Biurrun, I., Burrascano, S., Dembicz, I., Guarino, R., Kapfer, J., Pielech, R., Garcia-Mijangos, I., Wagner, V., Palpurina, S., (…) & Dengler, J. 2019. GrassPlot v. 2.00 – first update on the database of multi-scale plant diversity in Palaearctic grasslands. *Palaearctic Grasslands* 44: 26–47.

Dengler, J., Boch, S., Filibeck, G., Chiarucci, A., Dembicz, I., Guarino, R., Henneberg, B., Janišová, M., Marcenò, C., (…) & Biurrun, I. 2016. Assessing plant diversity and composition in grasslands across spatial scales: the standardised EDGG sampling methodology. *Bulletin of the Eurasian Grassland Group* 32: 13−30.

Dengler, J., Wagner, V., Dembicz, I., García-Mijangos, I., Naqinezhad, A., Boch, S., Chiarucci, A., Conradi, T., Filibeck, G., (…) & Biurrun, I. 2018. GrassPlot – a database of multi-scale plant diversity in Palaearctic grasslands. *Phytocoenologia* 48: 331–347.

Dengler, J., Matthews, T.J., Steinbauer, M.J., Wolfrum, S., Boch, S., Chiarucci, A., Conradi, T., Dembicz, I., Marcenò, C., (…) & Biurrun, I. 2020. Species-area relationships in continuous vegetation: Evidence from Palaearctic grasslands. *Journal of Biogeography* 60: 72–86.

Filibeck, G., Cancellieri, L., Sperandii, M.G., Belonovskaya, E., Sobolev, N., Tsarevskaya, N., Becker, T., Berastegi, A., Bückle, C., (…) & Biurrun, I. 2018. Biodiversity patterns of dry grasslands in the Central Apennines (Italy) along a precipitation gradient: experiences from the 10th EDGG Field Workshop. *Bulletin of the Eurasian Grassland Group* 36: 25–41

#### 69) N43TTP: Dry grassland vegetation of Northwest Bulgaria – *EDGG*

**Data owner(s)**:

Eurasian Dry Grassland Group (EDGG; [www.edgg.org](http://www.edgg.org)), represented by the

- GrassPlot database through Idoia Biurrun^1^ and Jürgen Dengler^2^
- Local Organising Committee of the 11th EDGG Field Workshop through Kiril Vassilev^3^

**Affiliation(s):** ^1^Department of Plant Biology and Ecology, University of the Basque Country UPV/EHU, P.O. Box 644, 48080 Bilbao, Spain; ^2^Vegetation Ecology, Institute of Natural Resource Management (IUNR), Zurich University of Applied Sciences (ZHAW), Grüentalstr. 14, 8820 Wädenswil, Switzerland; ^3^Department of Plant and Fungal Diversity and Resources, Institute of Biodiversity and Ecosystem Research, Bulgarian Academy of Sciences, Acad. G. Bonchev, bl. 23, 1113 Sofia, Bulgaria

**Description:** Data from the 3rd EDGG Field Workshop on sampling multi-scale plant diversity data in Palaearctic grasslands, conducted 2011 in the Northwest Bulgaria (GrassPlot ID BG_A; Pedashenko et al. 2013). The EDGG Field Workshops use a standardized sampling approach (Dengler et al. 2016), and their data are stored in the collaborative GrassPlot database (Dengler et al. 2018, 2020; Biurrun et al. 2019). In this paper only the vascular plant data from the 10-m² plots were used.

**Geographical scope:** Northwest Bulgaria

**Taxonomical scope:** Vascular plants

**Ecological scope:** Terrestrial (Dry grasslands s.l.)

**Number of sampling units:** 98 (10 m²)

**Minimum distance between sampling units:** 0.01 km

**Maximum distance between sampling units:** 110 km

**Environmental variables:** *Elevation* (m a.s.l.), *aspect* (°), *inclination* (°), *maximum microrelief* (cm), *pH* (measured in H_2_O; for uppermost 15 cm of soil), *electical conductivity* (μS/cm; for uppermost 15 cm of soil); *sand content* (%; for uppermost 15 cm of soil, approximative from soil texture class); *silt content* (%; for uppermost 15 cm of soil, approximative from soil texture class); *clay content* (%; for uppermost 15 cm of soil, approximative from soil texture class); *organic C content* (%; for uppermost 15 cm of soil)

**Functional traits:** Please see the section III

**References:**

Biurrun, I., Burrascano, S., Dembicz, I., Guarino, R., Kapfer, J., Pielech, R., *et al.* (2019). GrassPlot v. 2.00: first update on the database of multi-scale plant diversity in Palaearctic grasslands. *Palaearct. Grasslands*, 44, 26–47.

Dengler, J., Boch, S., Filibeck, G., Chiarucci, A., Dembicz, I., Guarino, R., *et al.* (2016). Assessing plant diversity and composition in grass ‐ lands across spatial scales: the standardised EDGG sampling methodology. *Bullettin Eurasian Dry Grassl. Gr.*

Dengler, J., Wagner, V., Dembicz, I., García-Mijangos, I., Naqinezhad, A., Boch, S., *et al.* (2018). GrassPlot – a database of multi-scale plant diversity in Palaearctic grasslands. *Phytocoenologia*, 48, 331–347.

Dengler, J., Matthews, T.J., Steinbauer, M.J., Wolfrum, S., Boch, S., Chiarucci, A., *et al.* (2020). Species–area relationships in continuous vegetation: Evidence from Palaearctic grasslands. *J. Biogeogr.*, 47, 72–86.

Magnes, M., Kirschner, P., Janišová, M., Mayrhofer, H., Berg, C., Mora, A., *et al.* (2020). On the trails of Josias Braun-Blanquet–changes in the grasslands of the inneralpine dry valleys during the last 70 years. First results from the 11 th EDGG Field Workshop in Austria. *Palaearct. Grasslands*, 45, 34–58.

Pedashenko, H., Apostolova, I., Boch, S., Ganeva, A., Janišová, M., Sopotlieva, D., *et al.* (2013). Dry grasslands of NW bulgarian mountains: First insights into diversity, ecology and syntaxonomy. *Tuexenia*. 33: 309–346.

#### 70) N43TTP_1: Bay of Biscay dunes – *Juan Antonio Campos Prieto*

**Data owner(s)**: Juan Antonio Campos Prieto

**Affiliation(s):** Department of Plant Biology and Ecology, University of the Basque Country UPV/EHU, P.O. Box 644, 48080 Bilbao, Spain.

**Description:** Vegetation database (288 plots x 110 vascular plants) about the Atlantic dune ecosystem: Embryonic shifting dunes” (habitat code 2110), “Shifting dunes along the shoreline with Ammophila arenaria” (habitat code 2120) and “Fixed coastal dunes with herbaceous vegetation” (habitat code 2130). Vascular plants (and bryophytes and lichens too) were sampled during the months of june and july of 2014 and 2015, in physically delimited squared plots of 10 m2 (96 plots by each dune habitat).

The extended Braun-Blanquet scale (Westhoff and Van Der Maarel, 1978) was used, in which value r = rare + = sparse, 1 =<4%, 2 is divided in 2 m = 5%, 2a = 5–12% and 2b = 12–25%, 3 = 25–49%, 4 = 50–74%, 5 = 75–100%.

Partial data are available in: https: //doi.org/10.1016/j.dib.2018.12.005

**Geographical scope:** Cantabrian Coast: from Northwest of Iberian Peninsula to Southern of France

**Taxonomical scope:** Atlantic dune vegetation

**Ecological scope:** Terrestrial

**Number of sampling units:** 288 plots (10 m²)

**Minimum distance between sampling units:**

**Maximum distance between sampling units:** Aprox. 750 km

**Environmental variables:** pH, organic matter (%), Kjeldhal Nitrogen (%), P (mg/l), Na (mg/l), K (mg/l), Mg (mg/l), Ca (mg/l) and soil texture (%)

**Functional traits:** Please see the section III

**Reference:** Torca, M., Campos, J.A. & Herrera, M. (2019). Species composition and plant traits of south Atlantic European coastal dunes and other comparative data. *Data in Brief*, 22: 207-213.

Westhoff, V. & E. Van der Maarel, E. (1978). *The Braun-Blanquet approach*. R.H. Whittaker (Ed.), Classification of Plant Communities, Dr. W. Junk Publishers, The Hague, pp. 287-399.

#### 71) N43TTP_2: Pinus pinea afforestations – Gianmaria Bonari and Claudia Angiolini

**Data owner(s)**: Gianmaria Bonari¹ (60 plots) & Claudia Angiolini² (20 plots)

**Affiliation(s):** ¹Faculty of Science and Technology, Free University of Bozen-Bolzano, Piazza Università, 5, 39100, Bozen-Bolzano, Italy.

²Department of Life Sciences, University of Siena, Via P.A. Mattioli, 4, 53100, Siena, Italy.

**Description:** The dataset includes original data recently collected at several sites along the Italian Peninsula, basically covering the whole distribution of Stone pine in Italy (excluding islands). At all sites, *Pinus pinea* was planted approximately 100 years ago. The peculiarity of the data is the semi-natural character of these forests and the vegetation therein.

**Geographical scope:** Italy

**Taxonomical scope:** *Pinus pinea* forests on sandy soils (EU Habitat 2270)

**Ecological scope:** Terrestrial

**Number of sampling units:** 80 plots (100 m²)

**Minimum distance between sampling units:** 0.1 km

**Maximum distance between sampling units:** 400 km (N-S gradient)

**Environmental variables:** Tree cover, Shrub cover, Herb cover, plant chorology, life form, pH, CaCO3, Soil organic matter.

**Functional traits:** Please see the section III

**Reference**: These data have been used for the following publications:

Angiolini, C., Landi, M., Pieroni, G., Frignani, F., Finoia, M.G. & Gaggi, C. (2013). Soil chemical features as key predictors of plant community occurrence in a Mediterranean coastal ecosystem. *Estuar. Coast. Shelf Sci.*, 119, 91–100.

Bonari, G., Acosta, A.T.R. & Angiolini, C. (2017). Mediterranean coastal pine forest stands: Understorey distinctiveness or not? *For. Ecol. Manage.*, 391, 19–28.

Bonari, G., Migliorini, M., Landi, M., Protano, G., Fanciulli, P.P. & Angiolini, C. (2017). Concordance between plant species, oribatid mites and soil in a Mediterranean stone pine forest. *Arthropod. Plant. Interact.*, 11, 61–69.

Bonari, G., Acosta, A.T.R. & Angiolini, C. (2018). EU priority habitats: rethinking Mediterranean coastal pine forests. *Rend. Lincei. Sci. Fis. e Nat.*, 29, 295–307.

Bonari, G., Těšitel, J., Migliorini, M., Angiolini, C., Protano, G., Nannoni, F., *et al.* (2019). Conservation of the Mediterranean coastal pine woodlands: How can management support biodiversity? *For. Ecol. Manage.*, 443, 28–35.

And they are currently stored in this database:

Gianmaria, B., Knollová, I., Vlčková, P., Xystrakis, F., Çoban, S., Sağlam, C., *et al.* (2019). CircumMed Pine Forest Database: an electronic archive for Mediterranean and Submediterranean pine forest vegetation data. *Phytocoenologia*, 49, 311–318.

#### 72) N43TTP_3: Dry grassland vegetation of Navarre, Spain – *EDGG*

**Data owner(s)**:

Eurasian Dry Grassland Group (EDGG; [www.edgg.org](http://www.edgg.org)), represented by the

- GrassPlot database through Idoia Biurrun^1^ and Jürgen Dengler^2^
- Local Organising Committee of the 7th EDGG Field Workshop through Idoia Biurrun^1^

**Affiliation(s):** ^1^Department of Plant Biology and Ecology, University of the Basque Country UPV/EHU, P.O. Box 644, 48080 Bilbao, Spain; ^2^Vegetation Ecology, Institute of Natural Resource Management (IUNR), Zurich University of Applied Sciences (ZHAW), Grüentalstr. 14, 8820 Wädenswil, Switzerland

**Description:** Data from the 7th EDGG Field Workshop on sampling multi-scale plant diversity data in Palaearctic grasslands, conducted 2014 in Navarre, Spain (GrassPlot ID ES_A; Biurrun et al. 2014). The EDGG Field Workshops use a standardized sampling approach (Dengler et al. 2016), and their data are stored in the collaborative GrassPlot database (Dengler et al. 2018, 2020; Biurrun et al. 2019). In this paper only the vascular plant data from the 10-m² plots were used.

**Geographical scope:** Navarre, Spain

**Taxonomical scope:** Vascular plants

**Ecological scope:** Terrestrial (Dry grasslands s.l.)

**Number of sampling units:** 119 (10 m²)

**Minimum distance between sampling units:** 0.01 km

**Maximum distance between sampling units:** 120 km

**Environmental variables:** Elevation (m a.s.l.), aspect (°), inclination (°), maximum microrelief (cm), soil depth mean (cm), soil depth standard deviation (cm), skeleton content (mass %; for uppermost 15 cm of soil), pH (measured in H2O; for uppermost 15 cm of soil), electrical conductivity (μS/cm; for uppermost 15 cm of soil); carbonate content (%; for uppermost 15 cm of soil), organic C content (%; for uppermost 15 cm of soil)

**Functional traits:**

**References:** Biurrun, I., García-Mijangos, I., Berastegi, A., Ambarli, D., Dembicz, I., Filibeck, G., *et al.* (2014). Diversity of dry grasslands in Navarre (Spain). *Bull. Eur. Dry Grassl. Gr.*, 24, 4.

Biurrun, I., Burrascano, S., Dembicz, I., Guarino, R., Kapfer, J., Pielech, R., *et al.* (2019). GrassPlot v. 2.00: first update on the database of multi-scale plant diversity in Palaearctic grasslands. *Palaearct. Grasslands*, 44, 26–47.

Dengler, J., Boch, S., Filibeck, G., Chiarucci, A., Dembicz, I., Guarino, R., *et al.* (2016). Assessing plant diversity and composition in grass ‐ lands across spatial scales: the standardised EDGG sampling methodology. *Bullettin Eurasian Dry Grassl. Gr.*

Dengler, J., Wagner, V., Dembicz, I., García-Mijangos, I., Naqinezhad, A., Boch, S., *et al.* (2018). GrassPlot - A database of multi-scale plant diversity in Palaearctic grasslands. *Phytocoenologia*.

Dengler, J., Matthews, T.J., Steinbauer, M.J., Wolfrum, S., Boch, S., Chiarucci, A., *et al.* (2020). Species–area relationships in continuous vegetation: Evidence from Palaearctic grasslands. *J. Biogeogr.*, 47, 72–86.

#### 73) N44TTP: Dry grassland vegetation of Serbia – *EDGG*

**Data owner(s)**:

Eurasian Dry Grassland Group (EDGG; [www.edgg.org](http://www.edgg.org)), represented by the

- GrassPlot database through Idoia Biurrun^1^ and Jürgen Dengler^2^
- Local Organising Committee of the 9th EDGG Field Workshop through Zora Dajić Stevanović,^3^

**Affiliation(s):** ^1^Department of Plant Biology and Ecology, University of the Basque Country UPV/EHU, P.O. Box 644, 48080 Bilbao, Spain; ^2^Vegetation Ecology, Institute of Natural Resource Management (IUNR), Zurich University of Applied Sciences (ZHAW), Grüentalstr. 14, 8820 Wädenswil, Switzerland; ^3^Department of Agrobotany, Faculty of Agriculture, University of Belgrade, Nemanjina 6, 11080 Belgrade-Zemun, Serbia

**Description:** Data from the 9th EDGG Field Workshop on sampling multi-scale plant diversity data in Palaearctic grasslands, conducted 2016 in Serbia (GrassPlot ID RS_A; Aćić et al. 2017). The EDGG Field Workshops use a standardized sampling approach (Dengler et al. 2016), and their data are stored in the collaborative GrassPlot database (Dengler et al. 2018, 2020; Biurrun et al. 2019). In this paper only the vascular plant data from the 10-m² plots were used.

**Geographical scope:** Serbia

**Taxonomical scope:** Vascular plants

**Ecological scope:** Terrestrial (Dry grasslands s.l.)

**Number of sampling units:** 141 (10 m²)

**Minimum distance between sampling units:** 0.01 km

**Maximum distance between sampling units:** 212 km

**Environmental variables: *Elevation*** (m a.s.l.), ***aspect*** (°), ***inclination*** (°), ***maximum microrelief*** (cm), ***soil depth mean*** (cm), ***soil depth standard deviation*** (cm)

**Functional traits:** Please see the section III

**References:**

Aćić, S., Dengler, J., Biurrun, I., Becker, T., Becker, U., Berastegi, A., et al. (2017). Biodiversity patterns of dry grasslands at the meeting point of Central Europe and the Balkans: Impressions and first results from the 9th EDGG Field Workshop in Serbia*. Bull. Eurasian Dry Grassl. Gr*., 34, 19–31.

Biurrun, I., Burrascano, S., Dembicz, I., Guarino, R., Kapfer, J., Pielech, R., *et al.* (2019). GrassPlot v. 2.00: first update on the database of multi-scale plant diversity in Palaearctic grasslands. *Palaearct. Grasslands*, 44, 26–47.

Dengler, J., Boch, S., Filibeck, G., Chiarucci, A., Dembicz, I., Guarino, R., et al. (2016). Assessing plant diversity and composition in grass ‐ lands across spatial scales: the standardised EDGG sampling methodology. *Bullettin Eurasian Dry Grassl*. Gr.

Dengler, J., Wagner, V., Dembicz, I., García-Mijangos, I., Naqinezhad, A., Boch, S., et al. (2018). GrassPlot – a database of multi-scale plant diversity in Palaearctic grasslands. *Phytocoenologia*, 48, 331–347.

Dengler, J., Matthews, T.J., Steinbauer, M.J., Wolfrum, S., Boch, S., Chiarucci, A., et al. (2020). Species–area relationships in continuous vegetation: Evidence from Palaearctic grasslands*. J. Biogeogr*., 47, 72–86.

#### 74) N47TTP: Dry grassland vegetation of the inneralpine dry valleys, Austria – *EDGG*

**Data owner(s)**:

Eurasian Dry Grassland Group (EDGG; [www.edgg.org](http://www.edgg.org)), represented by the

- GrassPlot database through Idoia Biurrun^1^ and Jürgen Dengler^2^
- Local Organising Committee of the 11th EDGG Field Workshop through Martin Magnes^3^

**Affiliation(s):** ^1^Department of Plant Biology and Ecology, University of the Basque Country UPV/EHU, P.O. Box 644, 48080 Bilbao, Spain; ^2^Vegetation Ecology, Institute of Natural Resource Management (IUNR), Zurich University of Applied Sciences (ZHAW), Grüentalstr. 14, 8820 Wädenswil, Switzerland; ^3^Division of Plant Sciences, Institute of Biology, Karl-Franzens-University of Graz, Holteigasse 6, 8010 Graz, Austria

**E-mail**:

**Description:** Data from the 11th EDGG Field Workshop on sampling multi-scale plant diversity data in Palaearctic grasslands, conducted 2018 in the inneralpine dry valleys of Austria (GrassPlot ID AT_E; Magnes et al. 2020). The EDGG Field Workshops use a standardized sampling approach (Dengler et al. 2016), and their data are stored in the collaborative GrassPlot database (Dengler et al. 2018, 2020; Biurrun et al. 2019). In this paper only the vascular plant data from the 10-m² plots were used.

**Geographical scope:** Inneralpine dry valleys, Austria

**Taxonomical scope:** Vascular plants

**Ecological scope:** Terrestrial (Dry grasslands s.l.)

**Number of sampling units:** 67 (10 m²)

**Minimum distance between sampling units:** 0.01 km

**Maximum distance between sampling units:** 330 km

**Environmental variables:** Elevation (m a.s.l.), aspect (°), inclination (°), maximum microrelief (cm), soil depth mean (cm), soil depth standard deviation (cm), skeleton content (mass %; for uppermost 15 cm of soil), pH (measured in H2O; for uppermost 15 cm of soil), electrical conductivity (μS/cm; for uppermost 15 cm of soil); N content (%; for uppermost 15 cm of soil), C/N ratio (for uppermost 15 cm of soil), organic matter (%; for uppermost 15 cm of soil), organic C content (%; for uppermost 15 cm of soil); plant available P content (mg/100 g; for uppermost 15 cm of soil)

**Functional traits:** Please see the section III

**References:**

Biurrun, I., Burrascano, S., Dembicz, I., Guarino, R., Kapfer, J., Pielech, R., *et al.* (2019). GrassPlot v. 2.00: first update on the database of multi-scale plant diversity in Palaearctic grasslands. *Palaearct. Grasslands*, 44, 26–47.

Dengler, J., Boch, S., Filibeck, G., Chiarucci, A., Dembicz, I., Guarino, R., *et al.* (2016). Assessing plant diversity and composition in grass ‐ lands across spatial scales: the standardised EDGG sampling methodology. *Bullettin Eurasian Dry Grassl. Gr.*

Dengler, J., Wagner, V., Dembicz, I., García-Mijangos, I., Naqinezhad, A., Boch, S., *et al.* (2018). GrassPlot – a database of multi-scale plant diversity in Palaearctic grasslands. *Phytocoenologia*, 48, 331–347.

Dengler, J., Matthews, T.J., Steinbauer, M.J., Wolfrum, S., Boch, S., Chiarucci, A., *et al.* (2020). Species–area relationships in continuous vegetation: Evidence from Palaearctic grasslands. *J. Biogeogr.*, 47, 72–86.

Magnes, M., Kirschner, P., Janišová, M., Mayrhofer, H., Berg, C., Mora, A., *et al.* (2020). On the trails of Josias Braun-Blanquet–changes in the grasslands of the inneralpine dry valleys during the last 70 years. First results from the 11 th EDGG Field Workshop in Austria. *Palaearct. Grasslands*, 45, 34–58.

#### 75) N47TTP_3: Grassland vegetation from Ukraine – *Iwona Dembicz and Łukasz Kozub*

**Data owner(s)**: Iwona Dembicz and Łukasz Kozub

**Affiliation(s):** Department of Plant Ecology and Environmental Conservation, Faculty of Biology, University of Warsaw.

**Description:** Series of plots from different refuges of natural grassland vegetation of the Kherson Region

**Geographical scope:** Ukraine\Kherson Region

**Taxonomical scope:** steppe (Festucetalia valesiaceae) - potential natural vegetation of the area

**Ecological scope:** Terrestrial

**Number of sampling units:** 64 (10m²)

**Minimum distance between sampling units:**

**Maximum distance between sampling units:** 100 km

**Environmental variables:** soil: pH, EC (μS/cm, dillution 1: 2.5), loss on ignition (1.5 h, 550۫C), granulometry (percentage of sand, silt and clay); microclimate: inclination, aspect; aboveground standing crop [g/m^2^]

**Functional traits:** Please see the section III

**Reference:**

#### 76) N47TTP_2: Dry grassland vegetation of Transylvania, Romania – *EDGG*

**Data owner(s)**:

Eurasian Dry Grassland Group (EDGG; [www.edgg.org](http://www.edgg.org)), represented by the

- GrassPlot database through Idoia Biurrun^1^ and Jürgen Dengler^2^
- Local Organising Committee of the 1st EDGG Field Workshop through Eszter Ruprecht^3^

**Affiliation(s):** ^1^Department of Plant Biology and Ecology, University of the Basque Country UPV/EHU, P.O. Box 644, 48080 Bilbao, Spain; ^2^Vegetation Ecology, Institute of Natural Resource Management (IUNR), Zurich University of Applied Sciences (ZHAW), Grüentalstr. 14, 8820 Wädenswil, Switzerland; ^3^Hungarian Department of Biology and Ecology, Faculty of Biology and Geology, Babeș-Bolyai University, Republicii street 42, 400015 Cluj-Napoca, Romania

**E-mail**:

**Description:** Data from the 1st EDGG Field Workshop on sampling multi-scale plant diversity data in Palaearctic grasslands, conducted 2009 in Transylania, Romania (GrassPlot ID RO_A; Dengler et al. 2009, 2012; Tutureanu et al. 2014). The EDGG Field Workshops use a standardized sampling approach (Dengler et al. 2016), and their data are stored in the collaborative GrassPlot database (Dengler et al. 2018, 2020; Biurrun et al. 2019). In this paper only the vascular plant data from the 10-m² plots were used.

**Geographical scope:** Transylvania, Romania

**Taxonomical scope:** Vascular plants

**Ecological scope:** Terrestrial (Dry grasslands s.l.)

**Number of sampling units:** 82 (10 m²)

**Minimum distance between sampling units:** 0.01 km

**Maximum distance between sampling units:** 180 km

**Environmental variables:** *Elevation* (m a.s.l.), *aspect* (°), *inclination* (°), *heat load index*, *maximum microrelief* (cm), *pH* (measured in H_2_O; for uppermost 15 cm of soil), *pH* (measured in CaCl_2_; for uppermost 15 cm of soil), *sand content* (%; for uppermost 15 cm of soil, approximative from soil texture class); *silt content* (%; for uppermost 15 cm of soil, approximative from soil texture class); *clay content* (%; for uppermost 15 cm of soil, approximative from soil texture class); *organic matter* (%; for uppermost 15 cm of soil)

**Functional traits:** Please see the section III

**References:**

Biurrun, I., Burrascano, S., Dembicz, I., Guarino, R., Kapfer, J., Pielech, R., et al. (2019). GrassPlot v. 2.00: first update on the database of multi-scale plant diversity in Palaearctic grasslands. *Palaearct. Grasslands*, 44, 26–47.

Rusina, S. & Kuzemko, A. (2009). EDGG cooperation on syntaxonomy and biodiversity of *Festuco-Brometea* communities in Transylvania (Romania): report and pre-liminary results. *Bull. Eur. Dry Grassl. Gr.*, 4, 13–19.

Dengler, J., Becker, T., Ruprecht, E., Szabó, A., Becker, U., Beldean, M., *et al.* (2012). Festuco-Brometea communities of the Transylvanian Plateau (Romania) - A preliminary overview on syntaxonomy, ecology, and biodiversity. *Tuexenia*, 32, 319–359.

Dengler, J., Boch, S., Filibeck, G., Chiarucci, A., Dembicz, I., Guarino, R., *et al.* (2016). Assessing plant diversity and composition in grass ‐ lands across spatial scales: the standardised EDGG sampling methodology. *Bullettin Eurasian Dry Grassl. Gr.*

Dengler, J., Wagner, V., Dembicz, I., García-Mijangos, I., Naqinezhad, A., Boch, S., *et al.* (2018). GrassPlot – a database of multi-scale plant diversity in Palaearctic grasslands. *Phytocoenologia*, 48, 331–347.

Dengler, J., Matthews, T.J., Steinbauer, M.J., Wolfrum, S., Boch, S., Chiarucci, A., *et al.* (2020). Species–area relationships in continuous vegetation: Evidence from Palaearctic grasslands. *J. Biogeogr.*, 47, 72–86.

Kącki, Z., Dembicz, I., Kozub, Ł., Swacha, G. & Dengler, J. 2014. Invitation to the 8th EDGG Field Workshop, Poland, June 2015. *Bulletin of the European Dry Grassland Group* 24/25: 26–34.

Turtureanu, P.D., Palpurina, S., Becker, T., Dolnik, C., Ruprecht, E., Sutcliffe, L.M.E., *et al.* (2014). Scale- and taxon-dependent biodiversity patterns of dry grassland vegetation in Transylvania. *Agric. Ecosyst. Environ.*, 182, 15–24.

#### 77) N48TTP: Dry grassland vegetation of Central Podolia, Ukraine – *EDGG*

**Data owner(s)**:

Eurasian Dry Grassland Group (EDGG; [www.edgg.org](http://www.edgg.org)), represented by the

- GrassPlot database through Idoia Biurrun^1^ and Jürgen Dengler^2^
- Local Organising Committee of the 2nd EDGG Field Workshop through Anna Kuzemko^3^

**Affiliation(s):** ^1^Department of Plant Biology and Ecology, University of the Basque Country UPV/EHU, P.O. Box 644, 48080 Bilbao, Spain; ^2^Vegetation Ecology, Institute of Natural Resource Management (IUNR), Zurich University of Applied Sciences (ZHAW), Grüentalstr. 14, 8820 Wädenswil, Switzerland; ^3^M.G. Kholodny Institute of Botany of the NAS of Ukraine, 2, Tereshchenkivska str., 01601 Kyiv, Ukraine

**Description:** Data from the 2nd EDGG Field Workshop on sampling multi-scale plant diversity data in Palaearctic grasslands, conducted 2010 in Central Podolia, Ukraine (GrassPlot ID UA_A; Kuzemko et al. 2014, 2016). The EDGG Field Workshops use a standardized sampling approach (Dengler et al. 2016), and their data are stored in the collaborative GrassPlot database (Dengler et al. 2018, 2020; Biurrun et al. 2019). In this paper only the vascular plant data from the 10-m² plots were used.

**Geographical scope:** Central Podolia, Ukraine

**Taxonomical scope:** Vascular plants

**Ecological scope:** Terrestrial (Dry grasslands s.l.)

**Number of sampling units:** 226 (10 m²)

**Minimum distance between sampling units:** 0.01 km

**Maximum distance between sampling units:** 155 km

**Environmental variables:** *Elevation* (m a.s.l.), *aspect* (°), *inclination* (°), *maximum microrelief* (cm), *skeleton content* (mass %; for uppermost 15 cm of soil), *pH* (measured in H_2_O; for uppermost 15 cm of soil), *electrical conductivity* (μS/cm; for uppermost 15 cm of soil); *N content* (%; for uppermost 15 cm of soil), *C/N ratio* (for uppermost 15 cm of soil), *organic C content* (%; for uppermost 15 cm of soil); *sand content* (%; for uppermost 15 cm of soil, approximative from soil texture class); *silt content* (%; for uppermost 15 cm of soil, approximative from soil texture class); *clay content* (%; for uppermost 15 cm of soil, approximative from soil texture class); *carbonate content* (%; for uppermost 15 cm of soil, approximative from ordinal scale)

**Functional traits:** Please see the section III

**References:**

Biurrun, I., Burrascano, S., Dembicz, I., Guarino, R., Kapfer, J., Pielech, R., *et al.* (2019). GrassPlot v. 2.00: first update on the database of multi-scale plant diversity in Palaearctic grasslands. *Palaearct. Grasslands*, 44, 26–47.

Dengler, J., Boch, S., Filibeck, G., Chiarucci, A., Dembicz, I., Guarino, R., *et al.* (2016). Assessing plant diversity and composition in grass ‐ lands across spatial scales : the standardised EDGG sampling methodology. *Bullettin Eurasian Dry Grassl. Gr.*, 32, 13–30.

Dengler, J., Wagner, V., Dembicz, I., García-Mijangos, I., Naqinezhad, A., Boch, S., *et al.* (2018). GrassPlot – a database of multi-scale plant diversity in Palaearctic grasslands. *Phytocoenologia*, 48, 331–347.

Dengler, J., Matthews, T.J., Steinbauer, M.J., Wolfrum, S., Boch, S., Chiarucci, A., *et al.* (2020). Species–area relationships in continuous vegetation: Evidence from Palaearctic grasslands. *J. Biogeogr.*, 47, 72–86.

Kuzemko, A.A., Becker, T., Didukh, Y.P., Ardelean, I.V., Becker, U., Beldean, M., *et al.* (2014). Dry grassland vegetation of Central Podolia (Ukraine)-a preliminary overview of its syntaxonomy, ecology and biodiversity. *Tuexenia*, 391–430.

Kuzemko, A.A., Steinbauer, M.J., Becker, T., Didukh, Y.P., Dolnik, C., Jeschke, M., *et al.* (2016). Patterns and drivers of phytodiversity in steppe grasslands of Central Podolia (Ukraine). *Biodivers. Conserv.*, 25, 2233–2250.

Dengler, J., Wagner, V., Dembicz, I., García-Mijangos, I., Naqinezhad, A., Boch, S., *et al.* (2018). GrassPlot – a database of multi-scale plant diversity in Palaearctic grasslands. *Phytocoenologia*, 48, 331–347.

Dengler, J., Boch, S., Filibeck, G., Chiarucci, A., Dembicz, I., Guarino, R., *et al.* (2016). Assessing plant diversity and composition in grass ‐ lands across spatial scales : the standardised EDGG sampling methodology. *Bullettin Eurasian Dry Grassl. Gr.*, 32, 13–30.

#### 78) N49TTP: Central European Lowland Forests – Jiri Dolezal and Jan Altman

**Data owner(s)**: Jiri Dolezal¹² and Jan Altman¹

**Affiliation(s):** ^1^Institute of Botany of The Czech Academy of Sciences, Zámek 1, 25243 Průhonice, Czech Republic;

^2^Faculty of Science, Department of Botany, University of South Bohemia, Na Zlaté Stoce 1, 37005 České Budějovice, Czech Republic;

**Description:** Plots covering 387 plant species of floodplain forests along Morava river

**Geographical scope:** Czech Republic

**Taxonomical scope:** Forest vegetation

**Ecological scope:** Terrestrial

**Number of sampling units:** 294 plots (100 m^2^)

**Minimum distance between sampling units:**

**Maximum distance between sampling units:** 30 kilometres

**Environmental variables:** Spatial coordinates, elevation (m), bioclimate variables.

**Functional traits:** Plant clonality: CLOPLA database.

Ellenberg Ecological Indicator Values: nutrients (1-9), light (1-9), soil moisture (1-5).

Plant size: Canopy height (m), DBH - stem diameter (cm).

Life forms: annuals/biennials/perennials.

Plant traits: LDMC (mg/g), Seed.mass (mg), SLA (mm^2^/mg), non-woody (%), woody (%), terminal velocity (m/s^2^).

Life forms: Chamaephyte, Phanerophyte, Liana, Hemicryptophyte, Hydrophyte, Therophyte, Geophyte.

**Reference:**

Klimešová, J. & de Bello, F. (2009). CLO-PLA: the database of clonal and bud bank traits of Central European flora. *J. Veg. Sci.*, 20, 511–516.

Karger, D.N., Conrad, O., Böhner, J., Kawohl, T., Kreft, H., Soria-Auza, R.W., *et al.* (2016). CHELSA climatologies at high resolution for the earth’s land surface areas (Version 1.1). *World Data Cent. Clim.*

Kleyer, M., Bekker, R.M., Knevel, I.C., Bakker, J.P., Thompson, K., Sonnenschein, M., *et al.* (2008). The LEDA Traitbase: A database of life-history traits of the Northwest European flora. *J. Ecol.*, 96, 1266–1274.

Chytrý, M., Tichý, L., Dřevojan, P., Sádlo, J. & Zelený, D. (2018). Ellenberg-type indicator values for the Czech flora. *Preslia*, 90, 83–103.

#### 79) N49TTP_2: Floodplain forests along the Thaya river*– Jiri Dolezal and Jan Altman*

**Data owner(s)**: Jiri Dolezal¹² and Jan Altman¹

**Affiliation(s):** ^1^Institute of Botany of The Czech Academy of Sciences, Zámek 1, 25243 Průhonice, Czech Republic;

^2^Faculty of Science, Department of Botany, University of South Bohemia, Na Zlaté Stoce 1, 37005 České Budějovice, Czech Republic;

**Description:** 417 plots covering floodplain forests along Thaya river

**Geographical scope:** Czech Republic

**Taxonomical scope:** Forest vegetation

**Ecological scope:** Terrestrial

**Number of sampling units:** 417 plots (4m²)

**Minimum distance between sampling units:**

**Maximum distance between sampling units:** 30 kilometres

**Environmental variables:**

Habitat and Land Use Categories: Forest - closed forests, Open - relict open oak forests, Edge - forest edges, CLconn - 40 m x 40 m forests clearings connected to alluvial meadows, CLisol - 40 m x 40 m clearings isolated within closed forests, Mead - alluvial meadows.

**Functional traits**

Plant clonality: CLOPLA database.

Ellenberg Ecological Indicator Values: nutrients (1-9), light (1-9), soil moisture (1-5).

Plant size: CanopyHeight (m).

Life forms: annuals/biennials/perennials.

Plant traits: LDMC (mg/g), Seed.mass (mg), SLA (mm^2^/mg), non-woody (%), woody (%), terminal velocity (m/s^2^).

Life forms: Chamaephyte, Phanerophyte, Liana, Hemicryptophyte, Hydrophyte, Therophyte, Geophyte.

**Reference:** Lanta, V., Mudrák, O., Liancourt, P., Bartoš, M., Chlumská, Z., Dvorský, M., *et al.* (2019). Active management promotes plant diversity in lowland forests: A landscape-scale experiment with two types of clearings. *For. Ecol. Manage.*, 448, 94–103.

Sebek, P., Bace, R., Bartos, M., Benes, J., Chlumska, Z., Dolezal, J., *et al.* (2015). Does a minimal intervention approach threaten the biodiversity of protected areas? A multi-taxa short-term response to intervention in temperate oak-dominated forests. *For. Ecol. Manage.*

#### 80) N51TTP: Dry grassland vegetation of Southern Poland – *EDGG*

**Data owner(s)**:

Eurasian Dry Grassland Group (EDGG; [www.edgg.org](http://www.edgg.org)), represented by the

- GrassPlot database through Idoia Biurrun^1^ and Jürgen Dengler^2^
- Local Organising Committee of the 8th EDGG Field Workshop through Zygmunt Kącki^3^ and Grzegorz Swacha^3^

**Affiliation(s):** ^1^Department of Plant Biology and Ecology, University of the Basque Country UPV/EHU, P.O. Box 644, 48080 Bilbao, Spain; ^2^Vegetation Ecology, Institute of Natural Resource Management (IUNR), Zurich University of Applied Sciences (ZHAW), Grüentalstr. 14, 8820 Wädenswil, Switzerland; ^3^Botanical Garden, University of Wrocław, ul. Henryka Sienkiewicza 23, 50-335 Wrocław, Poland.

**Description:** Data from the 4th EDGG Field Workshop on sampling multi-scale plant diversity data in Palaearctic grasslands, conducted 2015 in Southern Poland (GrassPlot ID PL_A; Kącki et al. 2014). The EDGG Field Workshops use a standardized sampling approach (Dengler et al. 2016), and their data are stored in the collaborative GrassPlot database (Dengler et al. 2018, 2020; Biurrun et al. 2019). In this paper only the vascular plant data from the 10-m² plots were used.

**Geographical scope:** Southern Poland

**Taxonomical scope:** Vascular plants

**Ecological scope:** Terrestrial (Dry grasslands s.l.)

**Number of sampling units:** 117 (10 m²)

**Minimum distance between sampling units:** 0.01 km

**Maximum distance between sampling units:** 495 km

**Environmental variables:** Elevation (m a.s.l.), inclination (°), maximum microrelief (cm), soil depth mean (cm), soil depth standard deviation (cm), skeleton content (mass %; for uppermost 15 cm of soil), pH (measured in H2O; for uppermost 15 cm of soil), pH (measured in CaCl2; for uppermost 15 cm of soil), electrical conductivity (μS/cm; for uppermost 15 cm of soil); organic matter (%; for uppermost 15 cm of soil), organic C content (%; for uppermost 15 cm of soil)

**Functional traits:** Please see the section III

**References:**

Biurrun, I., Burrascano, S., Dembicz, I., Guarino, R., Kapfer, J., Pielech, R., *et al.* (2019). GrassPlot v. 2.00: first update on the database of multi-scale plant diversity in Palaearctic grasslands. *Palaearct. Grasslands*, 44, 26–47.

Dengler, J., Boch, S., Filibeck, G., Chiarucci, A., Dembicz, I., Guarino, R., *et al.* (2016). Assessing plant diversity and composition in grass ‐ lands across spatial scales: the standardised EDGG sampling methodology. *Bullettin Eurasian Dry Grassl. Gr.*, 32, 13–30.

Dengler, J., Wagner, V., Dembicz, I., García-Mijangos, I., Naqinezhad, A., Boch, S., *et al.,* (2018). GrassPlot – a database of multi-scale plant diversity in Palaearctic grasslands. *Phytocoenologia,* 48, 331–347.

Dengler, J., Matthews, T.J., Steinbauer, M.J., Wolfrum, S., Boch, S., Chiarucci, A., *et al.* (2020). Species–area relationships in continuous vegetation: Evidence from Palaearctic grasslands. *J. Biogeogr.*, 47, 72–86.

Kącki, Z., Dembicz, I., Kozub, Ł., Swacha, G. & Dengler, J. (2014). Invitation to the 8th EDGG Field Workshop, Poland, June 2015. *Bulletin of the European Dry Grassland Group* 24/25, 26–34.

#### 81) N55TTP: Dry grassland vegetation of Khakassia, Russia – *EDGG*

**Data owner(s)**: Eurasian Dry Grassland Group (EDGG; [www.edgg.org](http://www.edgg.org)), represented by the

- GrassPlot database through Idoia Biurrun^1^ and Jürgen Dengler^2^
- Local Organising Committee of the 6th EDGG Field Workshop through Nikolai Ermakov^3^

**Affiliation(s):** ^1^Department of Plant Biology and Ecology, University of the Basque Country UPV/EHU, P.O. Box 644, 48080 Bilbao, Spain; ^2^Vegetation Ecology, Institute of Natural Resource Management (IUNR), Zurich University of Applied Sciences (ZHAW), Grüentalstr. 14, 8820 Wädenswil, Switzerland; ^3^Laboratory of Ecology and Geobotany, Central Siberian Botanical Garden, Russian Academy of Science, Siberian Branch, Zolotodolinskaya Str., 101, Novosibirsk, 630090, Russia

**Description:** Data from the 6th EDGG Field Workshop on sampling multi-scale plant diversity data in Palaearctic grasslands, conducted 2013 in Khakassia, Russia (GrassPlot ID RU_A; Janišová et al. 2013; Polyakova et al. 2016). The EDGG Field Workshops use a standardized sampling approach (Dengler et al. 2016), and their data are stored in the collaborative GrassPlot database (Dengler et al. 2018, 2020; Biurrun et al. 2019). In this paper only the vascular plant data from the 10-m² plots were used.

**Geographical scope:** Khakassia, Russia

**Taxonomical scope:** Vascular plants

**Ecological scope:** Terrestrial (Dry grasslands s.l.)

**Number of sampling units:** 132 (10 m²)

**Minimum distance between sampling units:** 0.01 km

**Maximum distance between sampling units:** 78 km

**Environmental variables:** Elevation (m a.s.l.), inclination (°), maximum microrelief (cm), soil depth mean (cm), soil depth standard deviation (cm), skeleton content (mass %; for uppermost 15 cm of soil), pH (measured in H2O; for uppermost 15 cm of soil), carbonate content (%; for uppermost 15 cm of soil), organic C content (%; for uppermost 15 cm of soil)

**Functional traits:** Please see the section III

**References:**

Biurrun, I., Burrascano, S., Dembicz, I., Guarino, R., Kapfer, J., Pielech, R., *et al.,* (2019). GrassPlot v. 2.00 – first update on the database of multi-scale plant diversity in Palaearctic grasslands. *Palaearctic Grasslands,* 44, 26–47.

Dengler, J., Boch, S., Filibeck, G., Chiarucci, A., Dembicz, I., Guarino, R., *et al.* (2016). Assessing plant diversity and composition in grass ‐ lands across spatial scales: the standardised EDGG sampling methodology. *Bullettin Eurasian Dry Grassl. Gr.*, 32, 13–30.

Dengler, J., Wagner, V., Dembicz, I., García-Mijangos, I., Naqinezhad, A., Boch, S., *et al.,* (2018). GrassPlot – a database of multi-scale plant diversity in Palaearctic grasslands. *Phytocoenologia,* 48, 331–347.

Dengler, J., Matthews, T.J., Steinbauer, M.J., Wolfrum, S., Boch, S., Chiarucci, A., *et al.* (2020). Species–area relationships in continuous vegetation: Evidence from Palaearctic grasslands. *J. Biogeogr.*, 47, 72–86.

Janišová, M., Becker, T., Becker, U., Demina, O., Dembicz, I., Ermakov, N., *et al.,* (2013). Steppes of Southern Siberia – Experiences from the 6th EDGG Research Expedition to Khakassia, Russia (22 July – 1 August 2013). *Bulletin of the European Dry Group,* 19/20, 31–48.

Polyakova, M.A., Dembicz, I., Becker, T., Becker, U., Demina, O.N., Ermakov, N., *et al.* (2016). Scale- and taxon-dependent patterns of plant diversity in steppes of Khakassia, South Siberia (Russia). *Biodivers. Conserv.*, 25, 2251–2273.

#### 82) S29TTP: Vegetation Monitoring of Naree and Yantabulla Stations – *John T. Hunter*

**Data owner(s)**: John T. Hunter

**Affiliation(s):** School of Rural and Environmental Sciences, University of New England, Armidale, NSW

**Description:** Full floristic plot data monitored yearly since 2015.

**Geographical scope:** Australia, New South Wales, Western Plains Botanical District, Mulga Bioregion

**Taxonomical scope:** Semi-arid woodland, shrublands and ephemeral wetlands.

**Ecological scope:** Terrestrial (Semi-arid)

**Number of sampling units:** 33 plots (20x20 m)

**Minimum distance between sampling units:**

**Maximum distance between sampling units:** 30 kilometres

**Environmental variables:** EC, pH, Mass, Nitrogen, Carbon.

Also dung weight to type, litter weight, percent count (80 points around plot) of life form type, dung, rock, litter. Cryptogam, herb, shrub, tree. Foliage projective cover of overstorey. Count of shrubs, trees.

**Functional traits:** Please see the section III

**Reference:** Hunter, J.T. (2016). Vegetation of Naree and Yantabulla stations on the Cuttaburra Creek, Far North Western Plains, New South Wales. *Cunninghamia*.

#### 83) S30TTP: Plant data from Brazil – Luciana da Silva Menezes and Gerhard E. Overbeck.

**Data owner(s)**: Luciana da Silva Menezes and Gerhard E. Overbeck*.*

**Affiliation(s):** Laboratory of Grassland Vegetation Studies, Universidade Federal do Rio Grande do Sul, Av. Bento Gonçalves 9500, Porto Alegre, RS 91501-970, Brazil

**Description:** Plant data from 1080 sampling units (1x1 m) nested within 108 plots (250x1 m) distributed in groups of nine, inside 12 grids (5x5 km) in the Campos Sulinos grasslands.

**Geographical scope:** Brazil/Campos Sulinos

**Taxonomical scope:** Terrestrial vascular plants

**Ecological scope:** Terrestrial (Grasslands)

**Number of sampling units:** 1080 (1m²)– 108 plots (250 m²)

**Minimum distance between sampling units:** 25m (sampling units of 1m²) - 1 km (plots of 250m²)

**Maximum distance between sampling units:** 660 km

**Environmental variables:**

| *Variable (measurement unit)* | *Method in brief* |
| --- | --- |
| pH | measured with soil dissolved in water |
| SMP index | amount of lime needed to reach pH 6 |
| P (mg dm^-3^) | phosphorus content, extracted with Mehlich I method |
| K (mg dm^-3^) | potassium content, extracted with Mehlich I method |
| O.M. (%) | organic matter content, measured through wet digestion |
| Al (cmol_c_ dm^-3^) | aluminium content, extracted with KCl 1 mol L-1 |
| Ca (cmol_c_ dm^-3^) | calcium content, extracted with KCl 1 mol L^-1^ |
| Mg (cmol_c_ dm^-3^) | magnesium content, extracted with KCl 1 mol L^-1^ |
| N (%) | percentage of nitrogen, extracted with Kjeldahl method |
| CTC (cmol_c_ dm^-3^) | cation exchange capacity, extracted with ammonium acetate |
| base saturation (%) | percentage of bases in CTC |
| Al saturation (%) | percentage of Al in CTC |
| clay (%) | percentage of particles smaller than <0.002 mm (sieved) |
| coarse sand (%) | percentage of particles with size between 0.2 and 2 mm (sieved) |
| fine sand (%) | percentage of particles with size between 0.054 and 0.2 mm (sieved) |
| silt (%) | percentage of particles with size between 0.002 and 0.053 mm (sieved) |

**Functional traits:** Most traits values were obtained from the TRY database through imputation by the gap-filling method described in Schrodt et al. (2015, Global Ecol. Biogeogr.). Traits obtained from local database (unpublished) were measured following the Pérez-Harguindeguy (2013, Australian Journal of Botany) protocol.

| *Trait (measurement unit)* | *Description* |
| --- | --- |
| LA (mm²) | Leaf area: one-sided or projected  area of an individual leaf. |
| LDMC (mg/g) | Leaf dry matter content: oven-dry mass (mg) of a leaf, divided by its water-saturated fresh mass (g). |
| SLA (mm²/mg) | Specific leaf area: one-sided area of a fresh leaf,  divided by its oven-dry mass. |

**Reference:** Unpublished data.

### IV. Datasets compiled from the cestes database

Datasets compiled from the CESTES database are publicy available [here](https://idata.idiv.de/ddm/Data/ShowData/286?version=22).

The "CESTES" database is composed of 80 datasets each of which includes four matrices: species community abundances or presences/absences across multiple sites, species trait information, environmental variables, and spatial coordinates of the sampling sites. All the information contained in this section can be found in the original publication and associated papers. The analysis of the distance decay 2.0 study used 49 datasets from CESTES, namely:

#### 84) N19MFI Fish data from Mexico – Villegér *et al.,* 2012

**Data owner(s)**: Domingo Flores Hernandez, David Mouillot, Julia ramos Miranda, Sébastién Villéger

**e-mail**:

**Description:** The survey focused on a 150-km long transect (18°37′16N–92°42′28W to 18°30′20N–91°28′03W) of 37 stations distributed in the south-western part of the Terminos Lagoon and along the adjacent coast (Villegér *et al.,* 2008).

**Geographical scope:** Mexico

**Taxonomical scope:** Fishes

**Ecological scope:** Marine

**Number of sampling units:** 35 sites

**Minimum distance between sampling units:** 3.00 kilometres

**Maximum distance between sampling units:** 124.85 kilometres

**Environmental variables:**

| *Variable* | *Unit or factor levels* |
| --- | --- |
| Depth | meters |
| Transparency | percentage |
| Salinity | Psu (practical salinity unit) |
| Dissolved oxygen | mg L^-1^ |

**Functional traits:**

| *Code* | *Variable* | *Description* |
| --- | --- | --- |
| logM | Mass | Size, metabolism |
| OgSh | Oral gape shape | Method to capture food items |
| OgPo | Oral gape position | Feeding method in the water column |
| GrLg | Gill raker length | Filtering ability or gill protection |
| GtLg | Gut length | Processing of energy poor resources such as vegetation and detritus |
| EySz | Eye size | Prey detection |
| EyPo | Eye position | Vertical position in the water column |
| BdSh | Body transversal shape | Vertical position in the water column and hydrodynamism |
| BdSf | Body transversal surface | Mass distribution along the body for hydrodynamism |
| PfPo | Pectoral fin position | Pectoral fin use for maneuverability |
| PfSh | Aspect ratio of the pectoral fin | Pectoral fin use for propulsion |
| CpHt | Caudal peduncle throttling | Caudal propulsion efficiency through reduction of drag |
| CfSh | Aspect ratio of the caudal fin | Caudal fin use for propulsion and/or direction |
| FsRt | Fins surface ratio | Main type of propulsion between caudal and pectoral fins |
| FsSf | Fins surface to body size ratio | Acceleration and/or manoeuvrability efficiency |

**Reference:** Villéger, S., Miranda, J.R., Hernandez, D.F. & Mouillot, D. (2012). Low Functional β-Diversity Despite High Taxonomic β-Diversity among Tropical Estuarine Fish Communities. *PLoS One*, 7, e40679.

#### 85) N33TTP (3-4): Plants from Morocco - Frenette *et al.,* 2012

**Data owner(s)**: Cedric Frenette-Dussault

**Description:** Five study sites in eastern Morocco where plant communities were assessed and functional traits measured inside and outside exclosures (50 plots).

**Geographical scope:** Eastern Morocco.

**Taxonomical scope:** Terrestiral vascular plants

**Ecological scope:** Terrestrial (steppe vegetation)

**Number of sampling units:** 50 plots

**Minimum distance between sampling units:** 0.11 kilometres

**Maximum distance between sampling units:** 177 kilometres

**Environmental variables:**

| *Code* | *Environmentalvariable* | *Units* | *Description* |
| --- | --- | --- | --- |
| Aridity | Aridity index |  | (Potential Evapotranspiration/Precipitation) computed from the FAO "Climate Information Tool" data |
| Exclosure | Duration of exclosures (up to 2009) |  |  |
| Bare ground 2009 | % of ground that had no plant cover in 2009 |  |  |
| Altitude | Elevation | m.a.s.l. | in meters above sea level measured by GPS |

**Functional traits:**

| *Code* | *Variable* | *Unit or factor levels* | *Description* |
| --- | --- | --- | --- |
| LA | Leaf Area | mm^2^ |  |
| SLA | Specific Leaf Area | mm^2^ mg^-1^ |  |
| LDMC | Leaf Dry Matter Content | mg g^-1^ |  |
| LC13 | Leaf Carbon 13 Isotope Content | ‰ |  |
| LN15 | Leaf Nitrogen 15 Isotope Content | ‰ |  |
| LCC | Leaf Carbon Content | μg mg^-1^ |  |
| LNC | Leaf Nitrogen Content | μg mg^-1^ |  |
| LCNR | Leaf Carbon: Nitrogen Ratio | unitless |  |
| VH | Vegetative Height | cm |  |
| MH | Maximum Height | cm |  |
| SM | Seed Mass | g |  |
| Clon | Clonality | Binary |  |
| Spin | Spinescence on stem or leaves | Binary |  |
| PosInflor | Position of Inflorescence | Binary |  |
| GrowthForm | Growth Form | Nominal |  |
| FlowerOnset | Onset of Flowering |  | Month when flowering starts for a majority of individuals |
| PastoralVal | Pastoral Value | 0-10 scale (0: poorest pastoral value, 10: best pastoral value) | Index combining nutrient content, productivity and palatability on a 0-10 scale (0: poorest pastoral value, 10: best pastoral value) |
| Succ | Succulence of Leaves or Stem | Binary |  |
| PhotoPath | Photosynthetic Pathway | Binary |  |

**References:** Frenette-Dussault, C., Shipley, B., Léger, J.-F., Meziane, D. & Hingrat, Y. (2012). Functional structure of an arid steppe plant community reveals similarities with Grime’s C-S-R theory. *J. Veg. Sci.*, 23, 208–222.

#### 86) N38FMI: Macroinvertebrates from a semi-arid catchment in Spain – Diaz *et al.,* 2009

**Data owner(s)**: Andres Mellado-Díaz

**Description:** 105 macroinvertebrate samples taken from 16 streams on seven occasions from 1999 to 2001: April 1999, July 1999, November 1999, February 2000, April–May 2000, July 2000 and December 2000–February 2001.

**Geographical scope:** Spain

**Taxonomical scope:** Macroinvertebrates

**Ecological scope:** Frehwaters

**Number of sampling units:** 51

**Minimum distance between sampling units:** < 0.01 kilometres

**Maximum distance between sampling units:** 2997 kilometres

**Environmental variables:**

| *Code* | *Environmentalvariable* | *Units* | *Description* |
| --- | --- | --- | --- |
| ENV1 | total suspended solids | mg/L |  |
| ENV2 | Ammonium | mg/L |  |
| ENV4 | Nitrate | mg/L |  |
| ENV5 | Phosphate | mg/L |  |
| ENV6 | Alkalinity | mEq/L |  |
| ENV7 | Discharge | L/s |  |
| ENV8 | Dissolved oxygen | mg/L |  |
| ENV10 | pH | none |  |
| ENV11 | Water temperature | Celsius degrees |  |
| ENV17 | Elevation | m |  |
| ENV19 | Substrate | categorical | substrate size (1=silt; 2= sand; 3= gravel; 4=  pebble-cobble; 5= boulder-rock) |

**Functional traits:**

| *Trait* | *Categories* |
| --- | --- |
| Maximal size | < 0.25 cm |
|  | > 0.25–0.5 cm |
|  | > 0.5–1 cm |
|  | > 1–2 cm |
|  | > 2–4 cm |
|  | > 4–8 cm |
|  | > 8 cm |
| Life cycle duration | <= 1 year |
|  | > 1 year |
| Potential No. reproductive cycles per year | < 1 |
|  | 1 |
|  | > 1 |
| Aquatic stages | egg |
|  | larva |
|  | pupa |
|  | adult |
| Reproduction | ovoviviparity |
|  | isolated eggs, free |
|  | isolated eggs, cemented |
|  | clutches, cemented or fixed |
|  | clutches, free |
|  | clutches, in vegetation |
|  | clutches, terrestrial |
|  | asexual reproduction |
| Dissemination | aquatic passive |
|  | aquatic active |
|  | aerial passive |
|  | aerial active |
| Resistance form | eggs, statoblasts |
|  | cocoons |
|  | cells against desiccation |
|  | diapause or dormancy |
|  | none |
| Food | fine sediment + microorganisms |
|  | detritus < 1mm |
|  | plant detritus > 1mm |
|  | living microphytes |
|  | living macrophytes |
|  | dead animal > 1mm |
|  | living microinvertebrates |
|  | living macroinvertebrates |
|  | vertebrates |
| Feeding habits | absorber |
|  | deposit feeder |
|  | shredder |
|  | scraper |
|  | filter feeder |
|  | piercer (plants or animals) |
|  | predator (carver/engulfer/swallower) |
|  | parasite |
| Respiration | tegument |
|  | gill |
|  | plastron |
|  | spiracle (aerial) |
| Locomotion and substrate relation | flier |
|  | surface swimmer |
|  | full water swimmer |
|  | crawler |
|  | burrower (epibenthic) |
|  | interstitial (endobenthic) |
|  | temporarily attached |
|  | permanently attached |

**References:** Díaz, A.M., Alonso, M.L.S. & Gutiérrez, M.R.V.A. (2008). Biological traits of stream macroinvertebrates from a semi-arid catchment: Patterns along complex environmental gradients. *Freshw. Biol.*, 53, 1–21.

#### 87) N41FMI: Macroinvertebrates from the Ebro River – Gallardo *et al.* 2009

**Data owner(s)**: Belinda Gallardo

**Description:** The authors selected 17 wetlands in the Ebro River and its floodplain along a 100-km length of the river. To account for spatial variability, the authors collected two to three samples at the upstream, mid- stream and downstream ends within each wetland (see Fig. 1) during two sampling surveys (September 2006 and August 2007).

**Geographical scope:**	Spain

**Taxonomical scope:** Macroinvertebrates

**Ecological scope:** Freshwaters

**Number of sampling units:** 76

**Minimum distance between sampling units:** < 0.01 kilometres

**Maximum distance between sampling units:** 95.58 kilometres

**Environmental variables:**

| *Code* | *Variable* | *Unit* |
| --- | --- | --- |
| TSS | Total suspended solid |  |
| TDS | Total dissolved solis |  |
| AFDM | (Ash free dry mass) |  |
| HCO3 | Bicarbonate |  |
| Ph |  |  |
| T | Temperature | ºC |
| O2 | Oxygen |  |
| ClorA |  |  |
| ClorC |  |  |
| DOP | Dissolved Organic Phosphorus | µg/L |
| Fl |  | µg/L |
| Cl |  | µg/L |
| NO2 |  | µg/L |
| Br |  | µg/L |
| NO3 |  | µg/L |
| DIP | Dissolved Inorganic Phosphorus | µg/L |
| SO4 |  | µg/L |
| K |  | µg/L |
| DIN | Dissolved inorganic nitrogen | µg/L |

**Functional traits:**

| *Functional trait* | *Category* |
| --- | --- |
| Maximal size | 2 |
|  | 5 |
|  | 10 |
|  | 20 |
|  | 40 |
|  | 80 |
|  | 100 |
| Respiration | Teg |
|  | Gill |
|  | Plast |
|  | Spir |
|  | Hydros |
| Life cycle duration | < 1 year |
|  | > 1 year |
| Potential number of reproduction cycles per year | < 1 |
|  | = 1 |
|  | > 1 |
| Reproduction | Ovovivi |
|  | Isoleggs |
|  | Cemeggs |
|  | Cemclutche |
|  | Freeclutche |
|  | Vegclutche |
|  | Terclutche |
|  | Asexual |
| Dissemination | AquaticP |
|  | AquaticA |
|  | AerialP |
|  | AerialA |
| Resistance form | Egg |
|  | Coccon |
|  | Diap |
|  | None |
| Locomotion and substrate relation | Flier |
|  | Surface |
|  | Swim |
|  | Crawler |
|  | Burr |
|  | Interst |
|  | Tattach |
|  | Pattach |
|  | Deposit |
| Feeding habits | Shred |
|  | Scrap |
|  | Filt |
|  | Piercer |
|  | Pred |
|  | Paras |
| Microhabitat | Pebb |
|  | gray |
|  | Sand |
|  | Silt |
|  | Macroph |
|  | Microph |
|  | Twigs |
|  | Org_det |
|  | Mud |
| Food | Sed_mic |
|  | Det < 1 |
|  | Det > 1 mm |
|  | Micro |
|  | Macroph |
|  | Dead_an |
|  | Microinv |
|  | Macroinv |
|  | Vert |
| Trophic level | Oligotrophic |
|  | Mesotrophic |
|  | Eutrophic |
|  | Cold |
|  | Warm |
|  | euryth |
|  | xenoS |
| Saprobity | oligoS |
|  | betaS |
|  | alphaS |
|  | polyS |
| pH | < 4 |
|  | 4.5 |
|  | 5 |
|  | 5.5 |
|  | 6 |
|  | > 6 |
| Aquatic stages | egg |
|  | larva |
|  | nymph |
|  | imago |

**References:** Gallardo, B., Gascón, S., García, M. & Comín, F.A. (2009). Testing the response of macroinvertebrate functional structure and biodiversity to flooding and confinement. *J. Limnol.*, 68, 315–326.

#### 88) N41TTP_2: Swiss meadows - van Klink *et al.* 2017

**Data owner(s)**: Roel van Klink

**Description:** 129 species from meadows located in the Swiss lowlands sampled from 2010 onwards

**Geographical scope:**	Switzerland

**Taxonomical scope:** Terrestrial vascular plants

**Ecological scope:** Terrestrial (Swiss lowlands)

**Number of sampling units:** 35

**Minimum distance between sampling units:** 0.04 kilometres

**Maximum distance between sampling units:** 0.73 kilometres

**Environmental variables:**

| *Code* | *Variable* | *Unit* | *Description* | Site identifier |
| --- | --- | --- | --- | --- |
| Precipitation | Total annual precipitation | mm | Millimeter | Mean precipitation per year |
| Forest | Forest | % | Degree | Percentage forest in 500m circle |
| Slope | Slope | % | Degree | Slope of meadow |
| TEMP4_9 | Temperature (April-September) | ºC | Degree celsius | Mean temperature April - September |
| Plant_biomass.m2 | Plant biomass m² | g/m² | Grams per square meter | Plant biomass in June 2015 per 1 m2 |
| Plant_sp_richness | Plant species richness | n/16 m² | Number of species per sixteen square meters | Number of plant species per 16m2 in June 2015 |

**Functional traits:**

| *Sp* | *Species identifier* | *none* | *none* | *Species identifier* |
| --- | --- | --- | --- | --- |
| Canopy_mean | Canopy mean size | m | Meter | Mean plant height in LEDA |
| Canopy_max | Canopy maximal size | m | Meter | Maximum plant height in LEDA |
| flowering.start | Flowering start | month number | Month | Starting month of flowering period |
| flowering.end | Flowering end | month number | Month | Latest month of flowering |
| flowering.mean | Flowering mean | month number | Month | Mean between start and end of flowering |

**References:** van Klink, R., Boch, S., Buri, P., Rieder, N.S., Humbert, J.-Y. & Arlettaz, R. (2017). No detrimental effects of delayed mowing or uncut grass refuges on plant and bryophyte community structure and phytomass production in low-intensity hay meadows. *Basic Appl. Ecol.*, 20, 1–9.

#### 89) N41TTP_3: Mediterranean grasslands – Bagaria *et al.,* 2012

**Data owner(s)**: Joan Pino

**Description:** 49 species from the Mediterranean mountain grasslands of southern Catalonia (NE Iberian Peninsula), over an area of 100 × 20 km

**Geographical scope:**	Mountain ranges of southern Catalonia (NE Iberian Peninsula) aligned along a NE–SW axis between Prades and Ports massifs (40°39′– 41°23′N, 0°10′– 1°10′E).

**Taxonomical scope:** Semi-natural Mediterranean mountain grasslands

**Ecological scope:** Terrestrial (Mountains)

**Number of sampling units:** 29

**Minimum distance between sampling units:** 0.98 kilometres

**Maximum distance between sampling units:** 101 kilometres

**Environmental variables:**

| Environmental variable | Units | Source |
| --- | --- | --- |
| Habitat amount and dynamics |  |  |
| Past patch area | ha | Photo-interpretation on ortho-corrected 1956 aerial photographs |
| Current patch area | ha | Photo-interpretation on 2004 ICC ortho-images |
| Percentage of patch area reduction | % ((past area-current area/past area)*100) | Calculation |
| Percentage of past grassland area in the landscape | % (grassland area in patch+buffer/patch+buffer area) | Photo-interpretation on ortho-corrected 1956 aerial photographs |
| Percentage of current grassland area in the landscape | % (grassland area in patch+buffer/patch+buffer area) | Photo-interpretation on 2004 ICC ortho-images |
| Percentage of grassland area reduction in the landscape | % ((past percentage-current percentage/past percentage)*100) | Calculation |
| Geographical location and climate |  |  |
| Longitude (UTMx) | m | GIS |
| Latitude (UTMy) | m | GIS |
| Mean annual temperature | °C | Digital Climatic Atlas of Catalonia (Pons 1996; Ninyerola et al. 2000) |
| Mean annual precipitation | mm | Digital Climatic Atlas of Catalonia (Pons 1996; Ninyerola et al. 2000) |

**Functional traits:**

| Trait | Description | Sources |
| --- | --- | --- |
| Reproductive traits |  |  |
| Seed size^1,2,3^ | Maximum length of a seed (mm) | Bolòs and Vigo (1984), Grime et al. (1988), Bishop and Davy (1994), García-Fayos and Cerdà (1997), Klotz et al. (2002), Bolòs et al. (2005), J.M. Ninot (pers. comm.), Flora Iberica (www.floraiberica.es), personal observation on fresh and herbarium material |
| Dispersal type^1^ | Anemochorous; Barochorous (or without any specific dispersal mechanism); Zoochorous | Julve (1998), Paula et al. (2009), Paula and Pausas (2009), personal observation on fresh and herbarium material |
| Corolla type^3^ | Based on flower morphology: Anemophilous corolla; Open entomophilous; Tubular; Zygomorphic | Personal observation on fresh and herbarium material |
| Flower or pseudanthium size^3^ | Maximum length of a flower or inflorescence when it acts as a unit of attraction (mm) | Bolòs and Vigo (1984), Dixon (1991), Bolòs et al. (2005), Flora Iberica (www.floraiberica.es), personal observation on fresh and herbarium material |
| Vegetative traits |  |  |
| Resprouting ability after fire^3^ | Yes; No | Papió (1994), Guerrero-Campo (1998), Arnan et al. (2007), Paula and Pausas (2008, 2009), Saura (2008), Paula et al. (2009), A. Rodrigo (pers. comm.), X. Arnan (pers. comm.), personal observation on fresh and herbarium material |
| Life-form^3^ | Annual; Herbaceous perennial; Woody | Based on Bolòs and Vigo (1984) |
| Mean plant height^3^ | Small (<25cm); Intermediate (25-50cm); Tall (>50cm) | Bolòs et al. (2005), J.M. Ninot (pers. comm.) |
| Vegetative spread^3^ | Yes; No | Fitter and Peat (1994), Klotz et al. (2002), Hill et al. (2004), Klimešová and Klimeš (2006), Klimešová and de Bello (2009), personal observation on fresh and herbarium material |
| Leaf anatomy^3^ | Aphyllous; Succulent; Mesomorphic; Scleromorphic | Klotz et al. (2002), J.M. Ninot (pers. comm.), personal observation on fresh and herbarium material |
| Leaf area^3^ | Area of a leaf or leaflet in the case of compound leaves, without considering lobulation (cm^2^) | Bolòs and Vigo (1984), Dixon (1991), Bishop and Davy (1994), Bolòs et al. (2005), J.M. Ninot (pers. comm.), Flora Iberica (www.floraiberica.es), personal observation on fresh and herbarium material |
| Spinescence^3^ | Yes; No | Personal observation on fresh and herbarium material |
| Ecological traits |  |  |
| Phytogeography^2,3^ | Strictly Mediterranean; Broadly Mediterranean; Other | Based on Bolòs et al. (2005) |
| Mammal herbivore preference^3^ | Preferred; Indifferent; Rejected | J. Bartolomé (pers. comm.), P. Casals (pers. comm.) |
| ^1^Dispersal, ^2^Establishment, ^3^Persistence |  |  |

**References:** Bagaria, G., Pino, J., Rodà, F. & Guardiola, M. (2012). Species traits weakly involved in plant responses to landscape properties in Mediterranean grasslands. *J. Veg. Sci.*, 23, 432–442.

#### 90) N43TTP_4: Vegetation from the New York coastal areas – Eallonardo *et al*., 2013

**Data owner(s)**: Anthony Eallonardo

**Description:** Vegetation of three inland salt/freshwater marsh sites near Montezuma, NY: Carncross (43.08° N, 76.71° W), Howland Island (43.07° N, 76.70° W) and Fox Ridge (43.05° N, 76.70° W).

**Geographical scope:** NewYork State, USA**.**

**Taxonomical scope:** Terrestrial vascular plants

**Ecological scope:** Terrestrial

**Number of sampling units:** 76

**Minimum distance between sampling units:** < 0.01 kilometres

**Maximum distance between sampling units:** 2.12 kilometres

**Environmental variables:**

| *Entry* | *Variable* | *Unit or factor levels* | *Description* |
| --- | --- | --- | --- |
| EC | electrical conductivity | dS/m | in soil samples |
| Ca | extractible calcium concentration | cmol c/kg | in soil samples |
| Fe | extractible iron concentration | cmol c/kg | in soil samples |
| K | extractible potassium concentration | cmol c/kg | in soil samples |
| Mg | extractible magnesium concentration | cmol c/kg | in soil samples |
| Na | extractible sodium concentration | cmol c/kg | in soil samples |
| S | extractible sulfur concentration | cmol c/kg | in soil samples |
| Na_K | ratio sodium-potassium | cmol: cmol | in soil samples |
| pH | pH | none | in soil samples |
| TotalN | total nitrogen | % | in soil samples |
| TotalC | total carbon | % | in soil samples |
| NO3 | nitrates | mg/kg | in soil samples |
| NH4 | ammonium | mg/kg | in soil samples |
| Flooding_dur | flooding duration | % of sampling events |  |

**Functional traits:**

| *Entry* | *Variable* | *Unit or factor levels* | *Description* |
| --- | --- | --- | --- |
| Area_mm2 | leaf size | mm^2 | not used as raw in the original study but log transformed (see below) |
| SLA_mm2_mg | specific leaf area | mm^2/mg | not used as raw in the original study but log transformed (see below) |
| mature_ht_cm | mature height | cm | not used as raw in the original study but log transformed (see below) |
| Perennial | perennial life span | binary, 1=yes, 0=no |  |
| Rhizomatous | rhizomatous growth | binary, 1=yes, 0=no |  |
| Graminoid | graminoid growth form | binary, 1=yes, 0=no |  |
| C4 | C4 photosynthetic pathway | binary, C4=1, C3=0 |  |
| Succulence | Succulence | binary, 1=yes, 0=no |  |
| arenchyma |  |  | not used in the original study |
| nmass | leaf N concentration | continuous (%) |  |
| narea | leaf N concentration | continuous (gN/m^2 leaf area) |  |
| cn | leaf C: N | continuous |  |
| logla | log transformed leaf size | continuous |  |
| logsla | log transformed specific leaf area | continuous |  |
| lognmass | log transformed leaf N concentration | continuous |  |
| lognarea | log transformed leaf N concentration | continuous |  |
| logcn | log transformed leaf C: N | continuous |  |
| loght | log transformed mature height | continuous |  |
| EC_95th | 95th percentile of EC | continuous | not used as trait in the original study |
| EC_max | max EC | continuous | not used as trait in the original study |

**References:** Eallonardo, A.S., Leopold, D.J., Fridley, J.D. & Stella, J.C. (2013). Salinity tolerance and the decoupling of resource axis plant traits. *J. Veg. Sci.*, 24, 365–374.

#### 91) N44TTP_2: Plants from the South of France – Raevel *et al.,* 2012

**Data owner(s)**: Valerie Raevel

**Description:** 97 plant species from the southern France

**Geographical scope:**	Mediterranean region of southern France (48°37′–43°58′N, 3°28′–4°E)

**Taxonomical scope:** Terrestrial vascular plants

**Ecological scope:** Terrestrial vascular plants

**Number of sampling units:** 52

**Minimum distance between sampling units:** < 0.01 kilometres

**Maximum distance between sampling units:** 50.10 kilometres

**Environmental variables:**

| *Entry* | *Variable* | *Unit or factor levels* | *Description* |
| --- | --- | --- | --- |
| Height | height | Meters | Height of the outcrop face |
| Slope | slope | Degrees | Slope of the outcrop face |

**Functional traits:**

| *Entry* | *Variable* | *Unit or factor levels* |
| --- | --- | --- |
| vegetative height | Cm | Maximal vegetative height |
| specific leaf area | mm² mg-1 | Leaf weight per unit area |
| seed mass | Mg | Mean seed mass From Navas et al. (2010) and Seed Information Database (Liu et al. 2008) |
| start of flowering | Early flowering, January-March (JM); Spring flowering, April-May (AM); Summer flowering, June-August (JA) | From Tison et al. (2014) |
| seed dispersal mode | Anemochory (Ano); Barochory and autochory (Baro) ; Endozoochory (End) ; Epizoochory and myrmecochory (Epi) | From LEDA (Kleyer et al. 2008); Biolflor (Klotz et al. 2002) |
| life span | Herbaceous perennial (HP); Woody perennial (WP); Short-lived annuals and biennials (SL) | From Tison et al. (2014) |
| life form | therophytes (T); chamaephytes (C); hemicryptophytes and geophytes grouped as herbaceous species (H); Phanerophytes (P) | Life forms according to Raunkiaer (1934) were taken from Tison et al. (2014) |

**References:**

Raevel, V., Violle, C. & Munoz, F. (2012). Mechanisms of ecological succession: Insights from plant functional strategies. *Oikos*, 121, 1761–1770.

Navas, M.-L., Roumet, C., Bellmann, A., Laurent, G. & Garnier, E. (2010). Suites of plant traits in species from different stages of a Mediterranean secondary succession. *Plant Biol.*, 12, 183–196.

Liu, K., Eastwood, R. J., Flynn, S., Turner, R. M., & Stuppy, W. H. (2008). Seed information database (release 7.1, May 2008). Available on-line at: http: //http: www. kew. org/data/sid (http: //http: www. kew. org/data/sid).

Tison, J. M., Jauzein, P., & Michaud, H. (2014). Flore de la France méditerranéenne continentale. *Conservatoire botanique national méditerranéen de Porquerolles* (CBNMed).

Spake, R., Barsoum, N., Newton, A.C. & Doncaster, C.P. (2016). Drivers of the composition and diversity of carabid functional traits in UK coniferous plantations. *For. Ecol. Manage.*, 359, 300–308.

Tison, J. M., Jauzein, P., & Michaud, H. (2014). Flore de la France méditerranéenne continentale. *Conservatoire botanique national méditerranéen de Porquerolles* (CBNMed).

Gargominy, O., Tercerie, S., Régnier, C., Ramage, T., Dupont, P., Daszkiewicz, P. & Poncet, L. (2017). *TAXREF v11, référentiel taxonomique pour la France: méthodologie, mise en oeuvre et diffusion. Muséum national d’Histoire naturelle, Paris. Rapport Patrinat 2017-116*. 152 pp.

#### 92) N44TAT : Carabid beetles from French forests – Barbaro and Halder 2009

**Data owner(s)**: Luc Barbaro

**Description:** The data comprises carabid beetles from plantation forests that cover around 1 million ha in the Landes de Gascogne region, south‐western France.

**Geographical scope:**	France

**Taxonomical scope:** Carabid beetles

**Ecological scope:** Terrestrial (Forests)

**Number of sampling units:** 195

**Minimum distance between sampling units:** 0.07 kilometres

**Maximum distance between sampling units:** 7.76 kilometres

**Environmental variables:**

| *Variable* | *Unit* | *Unit or factor levels* |
| --- | --- | --- |
| Distance to the nearest deciduous forest patch | m | Meter |
| Edge density | m ha¯¹ | Meter per hectare |
| Mean patch area | ha | Hectare |
| Mean shape index | none | Mean of all patch ratio of area on perimeter shape indices |
| Shannon diversity index | none | Shannon diversity of patch types |
| Firebreak cover | % | Percentage |
| Meadow cover | % | Percentage |
| Heatland cover | % | Percentage |
| Young pine cover | % | Percentage |
| Mature pine cover | % | Percentage |
| Deciduous wood cover | % | Percentage |

**Functional traits:**

| Variable | Unit | Unit or factor levels | Description |
| --- | --- | --- | --- |
| European trend | none | (1) Increasing, (2) Stable, (3) Declining | Overall population number trend. Defined as the percentage of national or regional range where the species is present. |
| European rarity | none | (1) Non-threatened, (2) Threatened |  |
| Regional rarity | none | (1) >15 regional data, (2) 10-15 regional data, (3) 4-9 regional data, (4) <3 regional data |  |
| Biogeographical position | none | (1) Mediterranean or atlantic, (2) Widespread, (3) Northern or central |  |
| Daily activity | none | (1) Diurnal, (2) Both diurnal/nocturnal, (3) Nocturnal |  |
| Diet type | none | (1) Collembola, (2) Generalist predators, (3) Phytophagous or mixed |  |
| Overwintering | none | (1) Imago only, (2) Imago and larvae |  |
| Body color | none | (1) Black or pale brown, (2) Metallic |  |
| Breeding season | none | (1) Spring breeder, (2) Summer breeder, (3) Autumn breeder |  |
| Body size | mm | (1) <6 mm (2) 6-7.9 mm, (3) 8-9.9 mm, (4) 10-11.9 mm |  |
| Wing development | none | (1) Brachypterous, (2) Dimorphic, (3) Macropterous |  |
| Adult activity period | none | (1) Early spring, (2) Late spring, (3) Summer/autumn |  |

**References:** Barbaro, L. & Van Halder, I. (2009). Linking bird, carabid beetle and butterfly life-history traits to habitat fragmentation in mosaic landscapes. *Ecography (Cop.).*, 32, 321–333.

#### 93) N44TBI: Birds from French forests – Barbaro and Halder 2009

**Data owner(s)**: Luc Barbaro

**Description:** The data comprises birds from plantation forests that cover around 1 million ha in the Landes de Gascogne region, south‐western France.

**Geographical scope:**	France

**Taxonomical scope:** Birds

**Ecological scope:** Terrestrial (Forests)

**Number of sampling units:** 195

**Minimum distance between sampling units:** 0.07 kilometres

**Maximum distance between sampling units:** 7.76 kilometres

**Environmental variables:**

| *Entry* | *Variable* | *Unit* | *Unit or factor levels* |
| --- | --- | --- | --- |
| DISDEC | Distance to the nearest deciduous forest patch | m | Meter |
| ED 400 | Edge density | m ha¯¹ | Meter per hectare |
| AREA 400 | Mean patch area | ha | Hectare |
| MEASH 400 | Mean shape index | none | Mean of all patch ratio of area on perimeter shape indices |
| SHDI 400 | Shannon diversity index | none | Shannon diversity of patch types |
| FIRE 400 | Firebreak cover | % | Percentage |
| MEAD 400 | Meadow cover | % | Percentage |
| HEAT 400 | Heatland cover | % | Percentage |
| YPIN 400 | Young pine cover | % | Percentage |
| MPIN 400 | Mature pine cover | % | Percentage |
| DECI 400 | Deciduous wood cover | % | Percentage |

**Functional traits:**

| *Entry* | *Variable* | *Unit* | *Unit or factor levels* |
| --- | --- | --- | --- |
| trend | National trend | none | (1) Increasing or stable, (2) Recently declining, (3) Long-term declining |
| forag | Foraging technique | none | (1) Ground prober, (2) Ground gleaner, (3) Understory gleaner |
| diet | Diet | none | (1) Insectivore, (2) Mixed diet, (3) Granivore |
| nest | Nest location | none | (1) Cavity (tree or others), (2) Open in tree, (3) Open in shrub, (4) Open on ground |
| eggs | Clutch size | none | (1) ≤ 3 eggs, (2) 4 eggs, (3) 5-6 eggs, (4) ≥ 7 eggs |
| mass | Body mass | g | (1) ≤ 14 g, (2) 15-24 g, (3) 25-49 g, (4) ≥ 50 g |
| migr | Migration status | none | (1) Resident, (2) Temperate migrant, (3) Tropical migrant |
| natio | National rarity | % | (1) > 95 %, (2) 75-95 %, (3) <75 % |
| regio | Regional rarity | % | (1) > 95 %, (2) 75-95 %, (3) <75 % |
| biog | Biogeographical position | none | (1) Mediterranean or atlantic, (2) Widespread, (3) Northern or central |
| date | Average laying date | month | (1) March, (2) Early April, (3) Late April, (4) Early May, (5) Late May and June |
| range | Home-range size | ha | (1) Small (<1 ha), (2) Medium (1-4 ha), (3) Large (>4 ha) |

**References:** Barbaro, L. & Van Halder, I. (2009). Linking bird, carabid beetle and butterfly life-history traits to habitat fragmentation in mosaic landscapes. *Ecography (Cop.).*, 32, 321–333.

#### 94) N44TBI_2: Birds from French vineyards – Barbaro *et al*., 2017

**Data owner(s)**: Luc Barbaro

**Description:** Bird communities were sampled using transect counts, where all birds heard and seen were recorded except flyovers, within a width of 100 m, that is 50 m from the observer on each transect side.

**Geographical scope:**	Aquitaine, south‐western France

**Taxonomical scope:** Birds

**Ecological scope:** Terrestrial (Vineyeards)

**Number of sampling units:** 20

**Minimum distance between sampling units:** 1.62 kilometres

**Maximum distance between sampling units:** 39.89 kilometres

**Environmental variables:**

| *Entry* | *Variable* | *Unit* | *Unit or factor levels* |
| --- | --- | --- | --- |
| GrassCover | Grass cover | none | Full, half |
| SNH100 | SNH in 100 meters buffer | % | Proportion |
| SNH250 | SNH in 250 meters buffer | % | Proportion |
| SNH500 | SNH in 500 meters buffer | % | Proportion |
| SNH750 | SNH in 750 meters buffer | % | Proportion |
| SNH1000 | SNH in 1000 meters buffer | % | Proportion |

**Functional traits:**

| *Entry* | *Variable* | *Unit* | *Unit or factor levels* | *Description* |
| --- | --- | --- | --- | --- |
| DietBreed | Diet in a breeding season | none | Insectivore, omnivore, granivore, functional insectivore |  |
| ForagGuild | Foraging guild | none | Canopy gleaner, bark forager, ground gleaner, understorey gleaner, hawker flycatcher, ground prober |  |
| ClutchSize | Clutch size | none | None | Number of eggs produced by birds, particularly those laid in a nest |
| BM | Body mass | gm | Gram |  |
| Nest | Nesting site | none | (1) cavity (building), (2) cavity (tree), (3) open in tree, (4) open in shrub, (5) open on ground |  |
| Migr | Migration strategy | none | (1) resident, (2) temperate migrant, (3) early tropical migrant, (4) late tropical migrant | Migration habits |
| Date | Lying date | none | (1) March, (2) early April, (3) late April, (4) early May, (5) late May and June | Timing in which bird lies the eggs |
| Home | Home range size | none | (1) small (< 1 ha), (2) medium (1-4 ha), (3) large (> 4 ha) | Mean home range size |

**References:** Barbaro, L., Rusch, A., Muiruri, E.W., Gravellier, B., Thiery, D. & Castagneyrol, B. (2017). Avian pest control in vineyards is driven by interactions between bird functional diversity and landscape heterogeneity. *J. Appl. Ecol.*, 54, 500–508.

#### 95) N44TBT: Butterflies from the Czech Republic – Bartonova *et al.* 2016

**Data owner(s)**: Alena Bartonova

**Description:** Butterfly data for this study originated from a survey of 125 National Nature Reserves and National Natural Monuments, commissioned by the Nature Conservation Agency of the Czech Republic, and carried out in 2004–2006

**Geographical scope:**	nature reserves in the Czech Republic

**Taxonomical scope:** Butterflies

**Ecological scope:** Terrestrial

**Number of sampling units:** 122

**Minimum distance between sampling units:** 0.26 kilometres

**Maximum distance between sampling units:** 476 kilometres

**Environmental variables:**

| *Code* | *Variable* |
| --- | --- |
| Perim | Perimeter |
| RelPerim | Relative Perimeter (Perimeter/Area) |
| MeanAlt | Average altitude [m a.s.l.] |
| RangeAlt | range of altitudes [m] |
| Biotope | prevailing biotope type |
| ReserveArea | Reserve area [m2] |
| Npatch_inside | N patches inside (in reserve) |
| Nbiotop_inside | N biotopes inside (in reserve) |
| BufArea | 2000 m Buffer area [m2] |
| Npatch_outside | N of patches outside (2000 m buffer minus reserve) |
| Nbiotop_outside | N of biotopes outside (2000 m buffer minus reserve) |

**Functional traits:**

| *Trait* | *Unit or category* | *Description* |
| --- | --- | --- |
| body size | Min = 11.0, max = 37.5 [mm] | approximated as forewing length (from Higgins and Riley 1970) |
| mobility | 1 – extremely sedentary, 2 – very sedentary, 3 – sedentary, 4 – rather sedentary, 5 – less sedentary, 6 – willingly dispersing, 7 – mobile, 8 – very mobile, 9 – extremely mobile | the propensity to disperse (ranked 1–9 scale, from extremely sedentary to extremely mobile; Reinhardt et al. 2007) |
| density | 1 – 2/km^2^, 2 – 6/km^2^_,_ 3 – 25/km^2^, 4 – 1/ha, 5 – 4/ha, 6 – 16/ha, 7 – 64/ha, 8 – 260/ha, 9 – 1000/ha | the number of individuals which can occur per unit area of habitat (ranked 1–9 scale, sparse to dense, adapted from area demand in Reinhardt et al. 2007) |
| voltinism | Min = 1, max = 4 [average number of generations] | the average number of generations per year in the Czech Republic (Benes et al. 2002) |
| flight period lenght | Min = 1.5, max = 12 [months] | number of months of summed adult flight, excluding months of hibernation (Benes et al. 2002) |
| range size | 1 – smaller or maximal size of Europe, 2 – as large as Western Palaearctic, 3 – Palaearctic, 4 – larger than Palaearctic realm | the total geographical range (ranked 1–4 scale, from the size of Europe to larger than Palaearctic realm, adopted from Tolman and Lewington 2008) |
| fertility | 1 –19-27, 2 –28-39, 3 – 40-57, 4 – 58-82, 5 – 83-119, 6 – 120-173, 7 – 174-250, 8 – 251-363, 9 – 364-527 | defined as number of eggs for a female at eclosion (ranked 1–9 scale, from few to many, Reinhardt et al. 2007) |
| overwintering stage | 1 – egg, 2 – larva, 3 – pupa, 4 – hibernating adult, 5 – migrating adult | a proxy for development speed (ranked 1–5 scale, from egg to migrating adult, from Tolman and Lewington 2008) |
| trophic range | Min = 1.0, max = 5.7 | expressed as an index that weights the number of consumed host plant families (F) and genera (G): I = (G×F^a^)^½^, where a = F/2G (data obtained from Ebert and Rennwald 1991; index modified after Garcia-Barros 2000) |
| host plant form | 1 – ephemerals and small forbs, 2 – large forbs and grasses, 3 –bushes, creepers and small trees, 4 – large trees | expressing prevailing host plant apparency (ranked 1–4, from small forbs to trees) |
| distribution | Min = 3, max = 647 [grid squares of the Czech Republic] | the number of Czech Republic atlas grid squares with positive records for a given species in 2002–2013 (obtained from Czech Butterflies and Moths Recording [CBMR]) |
| Redlist | 1 – least concern, 2 – near threatened, 3 – vulnerable, 4 – endangered, 5 – critically endangered | the species status in the national Red list (Benes et al. 2005). |
| S-G continuum | Min = –1.5, max = 3.8 [position on PCA axis 1] | first ordination axis of a principal component analysis (PCA), relating all the butterflies to their traits |

**References:**

Bartonova, A., Benes, J., Fric, Z.F., Chobot, K. & Konvicka, M. (2016). How universal are reserve design rules? A test using butterflies and their life history traits. *Ecography (Cop.).*, 39, 456–464.

Benes, J., Konvicka, M., Dvorak, J., Fric, Z., Havelka, Z., Pavlicko, A., *et al.* (2002). Motyli Ceske republiky: Rozsireni a ochrana I. *Spol. pro Ochr. motylu, Praha*.

Beneš, J., Konvička, M., Dvořák, J., Fric, Z., Havelda, Z., Pavlíčko, A., *et al.* (2005). Hesperioidea & Papilionoidea (denní motýli). *Červený seznam ohrožených druhů České republiky. Bezobratlí*, 219–223.

Ebert, G. & Rennwald, E. (1991). Die Schmetterlinge Baden-Württembergs. Band 2: Tagfalter II. *Ulmer, Stuttgart*.

García-Barros, E. (2000). Body size, egg size, and their interspecific relationships with ecological and life history traits in butterflies (Lepidoptera: Papilionoidea, Hesperioidea). *Biol. J. Linn. Soc.*, 70, 251–284.

Higgins, L.G. & Riley, N.D. (1970). A field guide to the butterflies of Britain and Europe. *A F. Guid. to butterflies Britain Eur.*

Chytrý, M., Kučera, T., Kočí, M., Grulich, V. & Lustyk, P. (2010). Katalog biotopů České republiky.

Reinhardt, R., Sbieschne, H., Settele, J., Fischer, U. & Fiedler, G. (2007). Tagfalter von Sachsen (In: Beiträge zur Insektenfauna Sachsen bd. 6, Eds: B. Klausnitzer, R. Reinhardt)–Entomologische Nachrichten und Berichte 11.

Tolman, T. & Lewington, R. (2008). The most complete guide to the butterflies of Britain and Europe.

#### 96) N46TAT : Insects from Switzerland – van Klink *et al*. 2019

**Data owner(s)**: Roel van Klink

**Description:** The authors conducted a five-year experiment, replicated at 12 sites across the Swiss lowlands, applying three different mowing regimes to low- intensity hay meadows: (1) first cut of the year not earlier than 15 June (control regime); (2) the first cut delayed until 15 July; and (3) leaving an uncut grass refuge on 10–20% of the mea- dow area (after earliest first cut on 15 June).

**Geographical scope:**	Switzerland

**Taxonomical scope:** Insects (larvae of Lepidoptera and sawflies, and adults of moths, parasitoid wasps, wild bees, hoverflies, ground beetles, and rove beetles).

**Ecological scope:** Terrestrial

**Number of sampling units:** 35

**Minimum distance between sampling units:** 0.54 kilometres

**Maximum distance between sampling units:** 192 kilometres

**Environmental variables:**

| Code | Variable | Unit | Unit or factor levels | Description |
| --- | --- | --- | --- | --- |
| Precipitation | Total annual precipitation | mm | millimeter | Mean precipitation per year |
| Elevation | Elevation | m a.s.l. | Metres above sea level | Elevation above sea level per meadow |
| Forest | Forest | % | Degree | Percentage forest in 500m circle |
| Slope | Slope | % | Degree | Slope of meadow |
| TEMP4_9 | Temperature (April-September) | ºC | Degree celsius | Mean temperature April - September |
| Plant_biomass.m2 | Plant biomass m² | g/m² | Grams per square meter | Plant biomass in 2015 per 1 m2 |
| Plant_sp_richness | Plant species richness | n/16 m² | Number of species per sixten squere meters | Number of plan species per 16m2 |

**Functional traits:**

| Entry | Variable | Unit | Unit or factor levels |
| --- | --- | --- | --- |
| size_min_f | Female minimum body length | mm |  |
| size_max_f | Female maximum body length | mm |  |
| size_min_m | Male minimum body length | mm |  |
| size_max_m | Male maximum body length | mm |  |
| flight1_min | Start time of adult activity | month number |  |
| Nesting_guild | Nesting guild |  | categorical: below-ground or above-ground (= in tree holes etc) |
| larval_substrate | Larval substrate |  |  |
| literature_size | source of size data |  |  |
| literature_flight | source of phenologica data |  |  |

**References:** Amiet, F. & Gesellschaft, S.E. (1996). *Insecta Helvetica. A, Fauna: 12. Hymenoptera. Apidae.-T. 1. Allgemeiner Teil, Gattungsschlüssel, Gattungen Apis, Bombus und Psithyrus*. Musée d’Histoire naturelle.

Amiet, F., Herrmann, M., Müller, A. & Neumeyer, R. (2001). Fauna Helvetica 9. Apidae 3: Lasioglossum, Halictus. *Neuchatel, Switz. Cent. Suisse Cartogr. la Faune*.

Amiet, F., Herrmann, M., Müller, A. & Neumeyer, R. (2004). Fauna Helvetica 9. Apidae 4: Anthidium, Chelostoma, Coelioxys, Dioxys, Heriades, Lithurgus, Megachile, Osmia, Stelis. *Fauna Helv. 9. Apidae 4 Anthidium, Chelostoma, Coelioxys, Dioxys, Heriades, Lithurgus, Megachile, Osmia, Stelis.*

Amiet, F., Herrmann, M., Müller, A. & Neumeyer, R. (2007). *Apidae 5: Ammobates, Ammobatoides, Anthophora, Biastes, Ceratina, Dasypoda, Epeoloides, Epeolus, Eucera, Macropis, Melecta, Melitta, Nomada, Pasites, Tetralonia, Thyreus, Xylocopa*. Centre suisse de cartographie de la faune.

Amiet, F., Herrmann, M., Müller, A. & Neumeyer, R. (2010). Andrena, Melitturga, Panurginus, Panurgus. *Fauna Helv.*, 26.

Amiet, F., Müller, A. & Neumeyer, R. (1999). *Apidae 2: Colletes, Dufourea, Hylaeus, Nomia, Nomioides, Rhophitoides, Rophites, Sphecodes, Systropha*. Schweizerische Entomologische Gesellschaft.

van Klink, R., Menz, M.H.M., Baur, H., Dosch, O., Kühne, I., Lischer, L., *et al.* (2019). Larval and phenological traits predict insect community response to mowing regime manipulations. *Ecol. Appl.*, 29, e01900.

Westrich, P. (1989). *Die Wildbienen Baden-Württembergs/die Gattungen und Arten*. E. Ulmer.

#### 97) N46FFI: Fish communities from the lake Drouin, Canada. - Brind’Amour *et al.* 2011.

**Data owner(s)**: Anik Brind'Amour

**Description:** The fish community and the environmental variables were quantified at 90 (Lake Drouin) and 60 sites (Lake Paré) that covered the complete perimeter of the littoral zones of the lakes. The sampling sites were defined as an area that possessed fairly homogenous attributes with respect to a combination of environmental variables (i.e., substrate, macrophyte density). The present dataset corresponds to the Lake Drouin.

**Geographical scope:**	Quebec (QC), Canada.

**Taxonomical scope:** Fish

**Ecological scope:** Freshwaters

**Number of sampling units:** 90

**Minimum distance between sampling units:** 0.01 kilometres

**Maximum distance between sampling units:** 0.86 kilometres

**Environmental variables:**

| Code | Variable | Unit or factor levels |
| --- | --- | --- |
| Litt | Mean littoral slope | quantitative |
| Rip | Riparian slope | presence/absence |
| Z | Mean depth | quantitative |
| Sand | Substrate: Sand | presence/absence |
| Rock | Substrate: Rock | presence/absence |
| Boulders | Substrate: Boulders | presence/absence |
| Bedrock | Substrate: Bedrock | presence/absence |
| WoodDebris | Substrate: Woody debris | presence/absence |
| Cottage | Riparian use: Cottage / brick wall | presence/absence |
| Forest | Riparian use: Forest | presence/absence |
| Beach | Riparian use: Beach | presence/absence |
| Bush | Riparian use: Bush | presence/absence |
| Tree | Riparian trees | presence/absence |
| Emerg | Macrophytes: mean density of emergent | quantitative |
| Subm | Macrophytes: mean density of submersed | quantitative |
| cover | Macrophytes: bottom cover (M. spicatum) | percentage |
| Fetch | Fetch | m |
| Trib | Distance to tributary | m |
| Size | Surface of a sampling site | m^2 |

**Functional traits:**

| Code | Variable | Unit or factor levels |
| --- | --- | --- |
| DietPl | Type of diet | Plant |
| DietZoob | Type of diet | Zoobenthos |
| DietZoop | Type of diet | Zooplankton |
| FeedWns | Type of diet | Insect larvae |
| DietFish | Type of diet | Fish |
| FeedBenth | Feeding strata | Benthic |
| FeedWat | Feeding strata | Water column |
| FeedSurf | Feeding strata | Surface |
| BodyFusi | Body morphology | Fusiform |
| BodyComp | Body morphology | Compressed |
| BodyCyl | Body morphology | Cylindrical |
| MigraDai | Migration | Daily |
| MigraSea | Migration | Seasonal |
| MouthInf | Mouth position | Inferior |
| MouthSup | Mouth position | Superior |
| MouthTer | Mouth position | Terminal |
| Temp10-15 | Temperature | 10–15°C |
| Temp15-20 | Temperature | 15–20°C |
| Temp20-25 | Temperature | 20–25°C |
| Oxy7-8 | Dissolved oxygen | 7–8 mg/L |
| Oxy5-7 | Dissolved oxygen | 5–7 mg/L |
| Oxy0 | Dissolved oxygen | < 0,2 mg/L |
| ActDiur | Activity | Diurnal |
| ActNoct | Activity | Nocturnal |

**References:** Brind’Amour, A., Daniel, B., Dray, S. & Legendre, P. (2011). Relationships between species feeding traits and environmental conditions in fish communities: A three-matrix approach. *Ecol. Appl.*, 21, 363–377.

#### 98) N47TAT: Carabid beetles from Switzerland – van Klink *et al*. 2019

**Data owner(s)**: Roel van Klink

**Description:** The authors conducted a five-year experiment, replicated at 12 sites across the Swiss lowlands, applying three different mowing regimes to low- intensity hay meadows: (1) first cut of the year not earlier than 15 June (control regime); (2) the first cut delayed until 15 July; and (3) leaving an uncut grass refuge on 10–20% of the mea- dow area (after earliest first cut on 15 June).

**Geographical scope:**	Switzerland

**Taxonomical scope:** Insects (larvae of Lepidoptera and sawflies, and adults of moths, parasitoid wasps, wild bees, hoverflies, ground beetles, and rove beetles).

**Ecological scope:** Terrestrial

**Number of sampling units:** 33

**Minimum distance between sampling units:** 0.54 kilometres

**Maximum distance between sampling units:** 192 kilometres

**Environmental variables:**

| Variable | Unit | Unit or factor levels | Description |
| --- | --- | --- | --- |
| Landscape unit | none | Name | Municipality / region where three meadows are located |
| Treatment | none | Delayed, refuge, control | mowing regime imposed on meadows |
| Total annual precipitation | mm | Millimeter | Mean precipitation per year |
| Elevation | m a.s.l. | Metres above sea level | Elevation above sea level per meadow |
| Forest | % | Degree | Percentage forest in 500m circle |
| Slope | % | Degree | Slope of meadow |
| Temperature (April-September) | ºC | Degree celsius | Mean temperature April - September |
| Plant biomass m² | g/m² | Grams per square meter | Plant biomass in 2015 per 1 m2 |
| Plant species richness | n/16 m² | Number of species per sixteen squere meters | Number of plant species per 16m2 |

**Functional traits:**

| Variable | Unit | Unit or factor levels |
| --- | --- | --- |
| Abrreviation used in original species list | cm | Centimeter |
| Full species name | cm | Centimeter |
| Minimum body length | none | Winged, dimorphic, short-winged/wingless |
| Maximum body length | none | Predator, herbivore, omnivore |
| Winged, dimorphic or wingless | none | Larva, imago |
| Predator, herbivore or omnivore |  | Predator, herbivore, omnivore |
| Overwintering as egg, latva or adult (Imago) |  | egg, larva, imago |
| Start of adult activity as middle of the month (original dat were given per month) | month number |  |
| End of adult activity | month number |  |

**References:** van Klink, R., Menz, M.H.M., Baur, H., Dosch, O., Kühne, I., Lischer, L., *et al.* (2019). Larval and phenological traits predict insect community response to mowing regime manipulations. *Ecol. Appl.*, 29, e01900.

Luka, H., Marggi, W., Huber, C., Gonseth, Y. & Nagel, P. (2009). *Coleoptera, Carabidae: ecology, atlas*. Centre suisse de cartographie de la faune.

Muilwijk, J., Felix, R.F.F.L., Dekoninck, W. & Bleich, O. (2015). *De loopkevers van Nederland en België (Carabidae)*. Nederlandse Entomologische Vereniging.

Homburg, K., Homburg, N., Schäfer, F., Schuldt, A. & Assmann, T. (2014). Carabids. org–a dynamic online database of ground beetle species traits (Coleoptera, Carabidae). *Insect Conserv. Divers.*, 7, 195–205.

Turin, H. (2000). *De Nederlandse loopkevers: verspreiding en oecologie (Coleoptera: Carabidae)*. Nationaal Natuurhistorisch Museum.

#### 99) N47TAT_2: Coleoptera from Switzerland – van Klink *et al*. 2019

**Data owner(s)**: Roel van Klink

**Description:** The authors conducted a five-year experiment, replicated at 12 sites across the Swiss lowlands, applying three different mowing regimes to low- intensity hay meadows: (1) first cut of the year not earlier than 15 June (control regime); (2) the first cut delayed until 15 July; and (3) leaving an uncut grass refuge on 10–20% of the mea- dow area (after earliest first cut on 15 June).

**Geographical scope:**	Switzerland

**Taxonomical scope:** Insects (larvae of Lepidoptera and sawflies, and adults of moths, parasitoid wasps, wild bees, hoverflies, ground beetles, and rove beetles).

**Ecological scope:** Terrestrial

**Number of sampling units:** 32

**Minimum distance between sampling units:** 0.54 kilometres

**Maximum distance between sampling units:** 192 kilometres

**Environmental variables:**

| Entry | Variable | Unit | Unit or factor levels | Description |
| --- | --- | --- | --- | --- |
| Landscape_Unit | Landscape unit | name | Name | Municipality / region where three meadows are located |
| Treatment | Treatment | none | Delayed, refuge, control | mowing regime imposed on meadows |
| Precipitation | Total annual precipitation | mm | millimeter | Mean precipitation per year |
| Elevation | Elevation | m a.s.l. | Metres above sea level | Elevation above sea level per meadow |
| Forest | Forest | % | Degree | Percentage forest in 500m circle |
| Slope | Slope | % | Degree | Slope of meadow |
| TEMP4_9 | Temperature (April-September) | ºC | Degree celsius | Mean temperature April - September |
| Plant_biomass.m2 | Plant biomass m² | g/m² | Grams per square meter | Plant biomass in 2015 per 1 m2 |
| Plant_sp_richness | Plant species richness | n/16 m² | Number of species per sixteen square meters | Number of plant species per 16m2 |

**Functional traits:**

| Entry | Variable | Unit or factor levels |
| --- | --- | --- |
| Body_length_min | Minimal body length | mm |
| Body_length_max | Maximal body length | mm |
| Humidity_preference | Humidity preference | sx = steno-xerophil, x = xerophil, m = mesophil, h = hygrophil, sh = steno-hygrophil |
| Phen_start | start of adult activity (mid month) | month number |

**References:**

Luka, H., Nagel, P., Luka, A. & Gonseth, Y. (in press) Staphylinidae Der Schweiz, Ein Beitrag Zur Ökologie. Centre Suisse de cartographie de la faune, Neuchâtel.

Horion, A. (1963) *Faunistik der Mitteleuropäischen Käfer. Band IX: Staphylinidae, 1. Teil Micropeplinae bis Euaestethinae.* Überlingen – Bodensee, 412 pp.

Horion, A. (1965) *Faunistik der Mitteleuropäischen Käfer. Band X: Staphylinidae, 2. Teil Paederinae bis Staphylininae.* Überlingen – Bodensee, 335 pp.

Horion, A. (1967) *Faunistik der Mitteleuropäischen Käfer. Band XI: Staphylinidae, 3. Teil Habrocerinae bis Aleocharinae (ohne Subtribus Athetae).* Überlingen – Boden-see, 419 pp.

#### 100) N47TAT_3: Diptera from Switzerland – van Klink *et al*. 2019

**Data owner(s)**: Roel van Klink

**Description:** The authors conducted a five-year experiment, replicated at 12 sites across the Swiss lowlands, applying three different mowing regimes to low- intensity hay meadows: (1) first cut of the year not earlier than 15 June (control regime); (2) the first cut delayed until 15 July; and (3) leaving an uncut grass refuge on 10–20% of the mea- dow area (after earliest first cut on 15 June).

**Geographical scope:**	Switzerland

**Taxonomical scope:** Insects (larvae of Lepidoptera and sawflies, and adults of moths, parasitoid wasps, wild bees, hoverflies, ground beetles, and rove beetles).

**Ecological scope:** Terrestrial

**Number of sampling units:** 34

**Minimum distance between sampling units:** 0.54 kilometres

**Maximum distance between sampling units:** 192 kilometres

**Environmental variables:**

| Variable | Unit | Unit or factor levels | Description |
| --- | --- | --- | --- |
| Landscape unit | name | Name | Municipality / region where three meadows are located |
| Treatment | none | Delayed, refuge, control | mowing regime imposed on meadows |
| Size of meadow in Ha | mm | millimeter | Mean precipitation per year |
| Elevation | m a.s.l. | Metres above sea level | Elevation above sea level per meadow |
| Forest | % | Degree | Percentage forest in 500m circle |
| Slope | % | Degree | Slope of meadow |
| Temperature (April-September) | ºC | Degree celsius | Mean temperature April - September |
| Plant biomass m² | g/m² | Grams per square meter | Plant biomass in 2015 per 1 m2 |
| Plant species richness | n/16 m² | Number of species per sixteen square meters | Number of plant species per 16m2 |

**Functional traits:**

| Variable | Unit | Unit or factor levels | Description |
| --- | --- | --- | --- |
| minimum body size | mm |  |  |
| maximum body size | mm |  |  |
| Start time of adult activity (months) | month number |  |  |
| end of adult activity (months) | month number |  |  |
| larval diet |  | Aphidophagous, Phytophagous | What the larvae feed on |
| larval substrate |  |  | as for bees derived from Larval.feeding.guild, here divided in species that feed on detritus or live in trees (outside meadow), and mostly aphidophagous ones, (in vegetation) |

**References:**

Reemer, M., Renema, W., van Steenis, W., Zeegers, T., Barendregt, A., Smit, J.T. *et al.* (2009) De Nederlandse Zweefvliegen. KNNV Uitgeverij, Zeist, The Netherlands.

Reemer, M. (2000) *Zweefvliegen veldgids (Diptera, Syrphidae).* Jeugdbondsuitgeverij

#### 101) N47TAT_4: Butterflies from Switzerland – van Klink *et al*. 2019

**Data owner(s)**: Roel van Klink

**Description:** The authors conducted a five-year experiment, replicated at 12 sites across the Swiss lowlands, applying three different mowing regimes to low- intensity hay meadows: (1) first cut of the year not earlier than 15 June (control regime); (2) the first cut delayed until 15 July; and (3) leaving an uncut grass refuge on 10–20% of the mea- dow area (after earliest first cut on 15 June).

**Geographical scope:**	Switzerland

**Taxonomical scope:** Insects (larvae of Lepidoptera and sawflies, and adults of moths, parasitoid wasps, wild bees, hoverflies, ground beetles, and rove beetles).

**Ecological scope:** Terrestrial

**Number of sampling units:** 35

**Minimum distance between sampling units:** 0.54 kilometres

**Maximum distance between sampling units:** 194 kilometres

**Environmental variables:**

| Variable | Unit | Unit or factor levels | Description |
| --- | --- | --- | --- |
| Landscape unit | name |  | Municipality / region where three meadows are located |
| Treatment | none | Delayed, refuge, control | mowing regime imposed on meadows |
| Total annual precipitation | mm | Millimeter | Mean precipitation per year |
| Elevation | m a.s.l. | Metres above sea level | Elevation above sea level per meadow |
| Forest | % | Percentage | Percentage forest in 500m circle |
| Slope | % | Degree | Slope of meadow |
| Temperature (April-September) | ºC | Degree celsius | Mean temperature April - September |
| Plant biomass m² | g/m² | Grams per square meter | Plant biomass in 2015 per 1 m2 |
| Plant species richness | n/16 m² | Number of species/16 m² | Number of plant species per 16m2 |

**Functional traits:**

| Code | Variable | Unit | Description |
| --- | --- | --- | --- |
| Synonym | Synonym name |  | Several species have also synonym names |
| Family | Family |  |  |
| Wingspan_min | Minimum recorded wing span | mm |  |
| Wingspan_max | Maximum wing span | mm |  |
| Source_wingspan | Source of wing span information |  | See Notes |
| flight_ad_min | Start of adult flight period | month number |  |
| flight_ad_peak | peak of adult flight period | month number |  |
| flight_ad_max | end of adult flight period | month number |  |
| Larval_substrate | Larval substrate | month number | Larvae feed on Trees or other plants that don't occur in meadows (= not in meadow) OR species feed on plants that occur in meadows (vegetation) |
| Source_phen | source of phenological and host information |  | See Notes |

**References:** Ebert, G., Steiner, A., Esche, T., Herrmann, R., Hofmann, A., Lussi, H. *et al*. (1994-2003) *Die Schmetterlinge Baden-Württembergs Band 3-9* (ed G Ebert). Ulmer, Stuttgart.

Steiner, A., Ratzel, U., Top-Jensen, M. & Fibiger, M. (2014) *Die Nachtfalter Deutschlands. Ein Feldführer*. Bugbook Publishing, Østermarie.

Willner, W. (2017) *Taschenlexikon der Schmetterlinge Europas Nachtfalter*. Quelle & Meyer Verlag, Wiebelsheim

#### 102) N47TTP_4: Weed communities from France – Fried, Kazakou & Gaba 2012

**Data owner(s)**: Guillaume Fried

**Description:** Vegetation data were extracted from two national large-scaleweed surveys conducted in France during the 1970s and the 2000s. The first survey was done between 1973 and 1976 and sampled a total of 2170 field. In this dataset, only the frequency and mean abundance of the 32 most frequent weed species were available. Data from the Biovigilance Flore survey (Fried *et al.* 2008) were used for the 2003–2006 period which involved 816 winter wheat fields out of the 2773 fields available and 206 weed species.

**Geographical scope:**	France

**Taxonomical scope:** Terrestrial vascular plants (Weed)

**Ecological scope:** Terrestrial

**Number of sampling units:** 218

**Minimum distance between sampling units:** < 0.01 kilometres

**Maximum distance between sampling units:** 889 kilometres

**Environmental variables:**

| Variable | Unit | Unit or factor levels | Description |
| --- | --- | --- | --- |
| Temperature | °C | Degree celsius | Annual mean temperature |
| Preciptiation | mm | Millimeter | Annual mean rainfall (rainfall over the 30 years) |
| Soil pH | pH | pH |  |
| Sowing date |  |  | Sowing or (planting) is the process of casting handfuls of seed over prepared ground, or broadcasting (Wiki) |
| Tillage depth | cm | Centimeters |  |
| Treatment frequency index |  |  | Ttreatment Frequency Index (TFX) is the sum of the ratio of the applied dose to the recommended dose of all the treatments applied in a year |
| Number of herbicide plants | none | Quantity |  |
| Number of HRAC families | none | Quantity | Herbicide classification according to primary site of action |
| Preceding crop | none | Maize, wcer, sunflower, orape, pea, sbeet |  |
| Tillage system | none | (CT) Conventional tillage, (MT) minimum tillage, (NT) no tillage |  |
| Soil texture | none | (1) Clay, (2) clay loam, (3) sandy clay, (4) silty loam, (5) silt clay, (6) S]sandy loam, (7) sand |  |

**Functional traits:**

| Variable | Unit | Unit or factor levels |
| --- | --- | --- |
| Plant life form | none | (Th) therophytes, (Geo) geophytes, (Hcr) hemicryptophytes |
| Mode of species dispersal | none | Gravity, animal, wind |
| Plant class | none | (Geo) geophytes, (Gr) grasses, (Bl) broadleaf plant |
| Plant height | cm | Centimeter |
| Seed weight | mg | Milligrams |
| Specific Leaf Area | mm²/mg | Squere millimetrs per gram |
| Germination start | month | Coded from 1 (January) till 12 (December) |
| Germination duration | month | Coded from 1 (January) till 12 (December) |
| Flowering onset | month | Coded from 1 (January) till 12 (December) |
| Flowering duration | month | Coded from 1 (January) till 12 (December) |

**References:** Fried, G., Norton, L.R. & Reboud, X. (2008). Environmental and management factors determining weed species composition and diversity in France. *Agric. Ecosyst. Environ.*

Fried, G., Kazakou, E. & Gaba, S. (2012). Trajectories of weed communities explained by traits associated with species’ response to management practices. *Agric. Ecosyst. Environ.*, 158, 147–155.

#### 103) N48FAB: Amphibian from France – Jeliazkov *et al.* 2014

**Data owner(s)**: Alienor Jeliazkov

**Description:** Amphibian biodiversity relationships in 150 ponds in an intensive agricultural landscape in Seine-et-Marne (France)

**Geographical scope:**	Brie, within the Seine-et- Marne department of France (Eastern Ile-de-France; 48.6°N/3.2°E)

**Taxonomical scope:** Amphibian

**Ecological scope:** Freshwaters

**Number of sampling units:** 135

**Minimum distance between sampling units:** 0.02 kilometres

**Maximum distance between sampling units:** 36.76 kilometres

**Environmental variables:**

| Variable | Unit or factor levels | Description |
| --- | --- | --- |
| Fish presence | 0 absence, 1 presence |  |
| Water Quality Index | %; 0% very bad quality - 100% very good quality | calculated from the following measured parameters: 02, temperature, pH, NH4, NH3, NO3,NO2, Si, PO4, chlorophyll; see Jeliazkov et al. 2014 for details |
| Proportion of wooded habitat | % | Proportion of surface occupied by wooded area in a 1000-m buffer around the focal pond |
| Pond density | integer | Number of ponds in a 1000-m buffer around the focal pond |
| Pond area | meters square |  |
| Agricultural gradient | numeric, PCA scores | PCA scores of the ponds along the Axis1 (driven by crops) extracted from a PCA performed on the proportion of surface covered by different land-cover types in 200-m buffer around ponds (see Jeliazkov et al. 2014) |
| Urban gradient | numeric, PCA scores | same as above with Axis 2; driven by urban habitat |
| Semi-natural gradient | numeric, PCA scores | same as above with Axis 3; driven by grasslands |

**Functional traits:**

| Variable | Unit or factor levels |
| --- | --- |
| Species identifier | none |
| Vertical foraging stratum: fossorial | binary |
| Vertical foraging stratum: terrestrial | binary |
| Vertical foraging stratum: aquatic | binary |
| Vertical foraging stratum: arboreal | binary |
| Diet: Arthropods | binary |
| Diet: Vertebrates | binary |
| Diel period active: diurnal | binary |
| Diel period active: nocturnal | binary |
| Diel period active: crepuscular | binary |
| Maximum adult body mass | grams |
| Minimum age at maturation/sexual maturity | years |
| Maximum age at maturation/sexual maturity | years |
| Maximum adult body size | millimeters |
| Maximum life span | years |
| Minimum no. of offspring or eggs per clutch | number |
| Maximum no. of offspring or eggs per clutch | number |

**References:**

Jeliazkov, A., Chiron, F., Garnier, J., Besnard, A., Silvestre, M. & Jiguet, F. (2014). Level-dependence of the relationships between amphibian biodiversity and environment in pond systems within an intensive agricultural landscape. *Hydrobiologia*, 723, 7–23.

Oliveira, B.F., São-Pedro, V.A., Santos-Barrera, G., Penone, C. & Costa, G.C. (2017). AmphiBIO, a global database for amphibian ecological traits. *Sci. Data*, 4, 170123.

#### 104) N49TAT: Continental-scale flea data – Krasnov *et al*. 2015

**Data owner(s)**: Boris R. Krasnov

**Description:** Continental data on flea species composition in diff erent regions from published surveys that reported the number of fleas of a given species found on a given number of individuals of a given species of a small mammal (Erinaceomorpha, Soricomorpha, Rodentia and Lagomorpha) in the northern and temperate Palearctic.

**Geographical scope:**	northern and temperate Palearctic

**Taxonomical scope:** Insects (Flea)

**Ecological scope:** Terrestrial

**Number of sampling units:** 45 sites

**Minimum distance between sampling units:** 77.63 kilometres

**Maximum distance between sampling units:** 7939.32 kilometres

**Environmental variables:**

| Variable | Unit | Unit or factor levels |
| --- | --- | --- |
| Size of the area | km² | Square kilometre |
| Mean altitude | m a.s.l. | Meters above sea level |
| Minimal altitude | m a.s.l. | Meters above sea level |
| Maximal altitude | m a.s.l. | Meters above sea level |
| NDVI for autumn |  | Normalized Difference Vegetation Indices (NDVI) for autumn |
| NDVI for spring |  | NDVI for spring |
| NDVI for summer |  | NDVI for summer |
| NDVI for winter |  | NDVI for winter |
| Mean autumn precipitation | mm | millimetre |
| Mean spring precipitation | mm | millimetre |
| Mean summer precipitation | mm | millimetre |
| Mean winter precipitation | mm | millimetre |
| Annual temperature ranges | °C | Degree celsius |
| Monthly temperature ranges | °C | Degree celsius |
| Maximal temperature | °C | Degree celsius |
| Mean temperature | °C | Degree celsius |
| Minimal temperature | °C | Degree celsius |

**Functional traits:**

| Variable | Unit | Unit or factor levels | Description |
| --- | --- | --- | --- |
| Abundance |  |  | Mean characteristic abundance (that is, mean number of fleas per individual host) |
| Host number |  |  | Mean size of the host spectrum (that is, mean number of host species on which a given fl ea species was recorded) |
| Number of host exploited across regions |  |  |  |
| Number of host exploited across continents |  |  |  |
| Host number correlation |  |  | Host number correlated with sampling effort (i.e., number of all host individuals examined; in log-log space, r2=0.03, F1,982=28.3, p<0.001). |
| Regional PHS |  |  | Phylogenetic host specificity (PHS) of a flea within a region |
| Continental PHS |  |  | Phylogenetic host specificity (PHS) of a flea within a continent |
| Living on the host body | none | (1)Yes, (0) no | Preference to spending most of the time on the host body |
| Living in the host nest | none | (1)Yes, (0) no | Preference to spending most of the time in the host burrow or nest |
| Living on the host body and nest | none | (1)Yes, (0) no | Preference to spending most of the time on the body of a host and in its burrow or nest |
| Reproduciton season in summer | none | (1)Yes, (0) no |  |
| Reproduciton season in winter | none | (1)Yes, (0) no |  |
| Reproduciton seasaonin the year round | none | (1)Yes, (0) no |  |

**References:** Krasnov, B.R., Shenbrot, G.I., Khokhlova, I.S., Stanko, M., Morand, S. & Mouillot, D. (2015). Assembly rules of ectoparasite communities across scales: Combining patterns of abiotic factors, host composition, geographic space, phylogeny and traits. *Ecography (Cop.).*, 38, 184–197.

#### 105) N49TBT: Bat communities from Europe – Charbonnier *et al*. 2016

**Data owner(s)**: Luc Barbaro

**Description:** The FunDivEurope Exploratory Platform was set up to investigate the effects of tree diversity on multiple for- est ecosystem functions in a network of 209 forest plots in Europe. Bat and bird communities were sampled in all 209 plots.

**Geographical scope:**	Europe

**Taxonomical scope:** Mammals (Bats)

**Ecological scope:** Terrestrial

**Number of sampling units:**	175

**Minimum distance between sampling units:** 0.04 kilometres

**Maximum distance between sampling units:** 3274.26 kilometres

**Environmental variables:**

| Variable | Unit | Unit or factor levels | Description |
| --- | --- | --- | --- |
| Plot altitude | m a.s.l. | none | Metres above sea level |
| Forest composition | none | Forest species composition (Quercus , Pinus, etc.) | Tree mixture identity based on mixture levels of main target tree species |
| Mean temperature | °C | Degree celsius |  |
| Mean precipitation | mm | Millimeter |  |
| Deciduous cover | % | Percentage | A deciduous cover is a biome dominated by deciduous trees which lose their leaves seasonally |
| Stratification index | none | none | Stratification index refers to the arrangement of the vertical vegetation in layers |
| Undestory cover | % |  | The understory is the underlying layer of vegetation in a forest or wooded area |
| Rao tree diversity |  |  | Rao's Q index of functional tree diversity based on leaf trait measurement |
| Mean basal areas | m²/ha | Squere meter per hectare | [Basal area is the area of a given section of land that is occupied by the cross-section of tree trunks and stems at the base](https://en.wikipedia.org/wiki/Tree) |

**Functional traits:**

| Variable | Unit | Unit or factor levels | Description |
| --- | --- | --- | --- |
| Foraging behaviours | none | Edge, open, glean, mixed | Foraging is searching for wild food resource |
| Diet type | none | Spec, interm, gener |  |
| Nest or roost site location | none | Tree, build, cave |  |
| Migration status | none | Resident, short, long, |  |
| Breeding date | none | May, jun, july |  |
| Home range size | none | Small, middle, large, |  |
| Body mass | gm | Gram |  |
| Number of pups | none | none | Mean number of pups per female |
| Species Thermal Index | ° C | Degree celsius | Mean temperature at the centroid of species range |
| Species Specialization Index | none | none | Coefficient of variation of densities of a given species across habitats |

**References:** Charbonnier, Y.M., Barbaro, L., Barnagaud, J.Y., Ampoorter, E., Nezan, J., Verheyen, K., *et al.* (2016). Bat and bird diversity along independent gradients of latitude and tree composition in European forests. *Oecologia*, 182, 529–537.

#### 106) N49TBI: Bird communities from Europe – Charbonnier *et al*. 2016

**Data owner(s)**: Luc Barbaro

**Description:** The FunDivEurope Exploratory Platform was set up to investigate the effects of tree diversity on multiple for- est ecosystem functions in a network of 209 forest plots in Europe. Bat and bird communities were sampled in all 209 plots.

**Geographical scope:**	Europe

**Taxonomical scope:** Birds

**Ecological scope:** Terrestrial

**Number of sampling units:** 208

**Minimum distance between sampling units:** 0.04 kilometres

**Maximum distance between sampling units:** 3274.26 kilometres

**Environmental variables:**

| Variable | Unit | Unit or factor levels | Description |
| --- | --- | --- | --- |
| Plot altitude | m a.s.l. | none | Metres above sea level |
| Forest composition | none | Forest species composition (Quercus, Pinus, etc.) | Tree mixture identity based on mixture levels of main target tree species |
| Mean temperature | °C | Degree celsius |  |
| Mean precipitation | mm | Millimeter |  |
| Deciduous cover | % | Percentage | A deciduous cover is a biome dominated by deciduous trees which lose their leaves seasonally |
| Stratification index | none | none | Stratification index refers to the arrangement of the vertical vegetation in layers |
| Undestory cover | % |  | The understory is the underlying layer of vegetation in a forest or wooded area |
| Rao tree diversity |  |  | Rao's Q index of functional tree diversity based on leaf trait measurement |
| Mean basal areas | m²/ha | Squere meter per hectare | [Basal area is the area of a given section of land that is occupied by the cross-section of tree trunks and stems at the base](https://en.wikipedia.org/wiki/Tree) |

**Functional traits:**

| *Variable* | *Unit* | *Unit or factor levels* | *Description* |
| --- | --- | --- | --- |
| Foraging behaviours | none | Under, ground, conopy, bark, prober, | Foraging is searching for wild food resource |
| Diet type | none | Insect, seeds, mixed |  |
| Nest or roost site location | none | Shrub, ground, tree, cavity |  |
| Migration status | none | Resident, short, long, |  |
| Breeding date | none | March, early april, late april,may, jun |  |
| Home range size | none | Small, middle, large, |  |
| Eggs number/mass | none | none |  |
| Body mass | gm | Gram |  |
| Species Thermal Index | ° C | Degree celsius | Mean temperature at the centroid of species range |
| Species Specialization Index | none | none | Coefficient of variation of densities of a given species across habitats |

**References:** Charbonnier, Y.M., Barbaro, L., Barnagaud, J.Y., Ampoorter, E., Nezan, J., Verheyen, K., *et al.* (2016). Bat and bird diversity along independent gradients of latitude and tree composition in European forests. *Oecologia*, 182, 529–537.

#### 107) N50TAT_5: Butterflies from the Czech Republic – Robinson *et al.* 2014

**Data owner(s)**: Natalie Robinson

**Description:** The authors used previously collected data (Kadlec *et al.* 2008; Konvicka & Kadlec 2011) of butterfly species from the Czech Republic.

**Geographical scope:**	Czech Republic

**Taxonomical scope:** Butterflies

**Ecological scope:** Terrestrial

**Number of sampling units:** 20

**Minimum distance between sampling units:** 0.52 kilometres

**Maximum distance between sampling units:** 16.98 kilometres

**Environmental variables:**

| Variable | Unit or factor levels | Description |
| --- | --- | --- |
| Habitat area | ha |  |
| Shape complexity | number of points | Number of Shape Characterizing Points |
| Edge permeability |  | see formula in Robinson et al. 2018, Table 1 |
| Proportion of open habitat | % |  |
| Water availability | Absent / Present | Water availability |
| Habitat heterogeneity | number of types | Number of land cover types |
| Open habitat in patch buffer | ha | Total open habitat in patch buffer; isolation metric |

**Functional traits:**

| Variable | Unit or factor levels |
| --- | --- |
| Average wing length | mm |
| Eggs per batch | Single, Small (2–50), Medium (51–200), Large (>200) |
| Voltinism | Univoltine (1 generation/yr.), Bivoltine (2 generations/yr.), Multivoltine (>2 generations/yr.) |
| Diapause strategy | Egg (or larva inside egg),  Larva, Pupa, Adult |
| Diet breadth | Monophagous, Oligophagous, Polyphagous |
| Flight period | Short (1–2 mo.), Medium (3–4 mo.), Long (>4 mo.) |

**References:** Robinson, N., Kadlec, T., Bowers, M.D. & Guralnick, R.P. (2014). Integrating species traits and habitat characteristics into models of butterfly diversity in a fragmented ecosystem. *Ecol. Modell.*, 281, 15–25.

Kadlec, T., Benes, J., Jarosik, V. & Konvicka, M. (2008). Revisiting urban refuges: Changes of butterfly and burnet fauna in Prague reserves over three decades. *Landsc. Urban Plan.*, 85, 1–11.

Konvicka, M. & Kadlec, T. (2011). How to increase the value of urban areas for butterfly conservation? A lesson from Prague nature reserves and parks. *Eur. J. Entomol.*, 108, 219–229.

#### 108) N51TAT_9: Insects from the floodplains of the river Elbe – Dziock *et al*. 2011

**Data owner(s)**: Frank Dziock

**Description:** Orthoptera species abundances sampled between August and September 2006 on sunny days with a minimum temperature of 20º C and low wind speed.

**Geographical scope:**	Sachsen-Anhalt, Germany.

**Taxonomical scope:** Insects (Orthoptera)

**Ecological scope:** Terrestrial

**Number of sampling units:** 34 sites

**Minimum distance between sampling units:** 0.05 kilometres

**Maximum distance between sampling units:** 8.46 kilometres

**Environmental variables:**

| Variable | Unit | Unit or factor levels | Description |
| --- | --- | --- | --- |
| Elevation | m | Meters | Above sea level |
| Distance to the river | m | Meters |  |
| Class of litter cover | none | From 1 (none) to 5 (very high) |  |
| Vegetation height | cm | Centimeters |  |
| Land use and intensity | none | (0) no management, (1) one annual cut, (2) two annual cuts, (3) two annual cuts and grazing by sheep afterwards | Class of land use management and intensity |
| Modelled duration of inundation | days | 0 days (very dry) – 104 days (very wet) | Calculated by one-dimensional hydrodynamic model based on field measurements and river gauges (Böhnke and Follner, 2002; Follner and Henle, 2006; Follner et al., 2010) |

**Functional traits:**

| Variable | Unit | Unit or factor levels | Description |
| --- | --- | --- | --- |
| Dispersal ability | none | High, med, low |  |
| Passive dispersal ability | none | High, med, low | Dispersal of eggs by being attached on a certain object (bark, leave, etc.) |
| Ovarioles number | none | 1-10 (low), 11-25 (moderate), >25 (high) |  |
| Body size | none | 16-18 mm (small), 19-21 mm (medium), >21 mm (large) |  |
| Oviposition in plant material | none | Yes, no | Oviposition in plant material (leafs, plant stems, tree bark, etc.) |
| Oviposition at the ground level | none | Yes, no | Oviposition at the ground level (directly into the ground or at the base of grass roots) |

**References:** Dziock, F., Gerisch, M., Siegert, M., Hering, I., Scholz, M. & Ernst, R. (2011). Reproducing or dispersing? Using trait-based habitat templet models to analyse Orthoptera response to flooding and land use. *Agric. Ecosyst. Environ.*, 145, 85–94.

Klaiber, J., Altermatt, F., Birrer, S., Chittaro, Y., Dziock, F., Gonseth, Y., *et al.* (2017). Fauna Indicativa. *WSL Berichte*, 54.

#### 109) N51TAT_10: Beetles from Scotland – Ribera *et al.* 2001

**Data owner(s)**: Ignacio Ribera

**Description:** Beetles were sampled with two parallel rows of nine pitfall traps. A total of eighty-seven sites were sampled, each site during one season. The whole study was carried within three years, between 1995 to 1997.

**Geographical scope:**	Scotland

**Taxonomical scope:** Insects (beetles)

**Ecological scope:** Terrestrial

**Number of sampling units:** 87

**Minimum distance between sampling units: <** 0.01 kilometre

**Maximum distance between sampling units:** 380.19 kilometres

**Environmental variables:**

| Variable | Unit or factor levels | Description |
| --- | --- | --- |
| Texture | 1, peat; 2, peaty loam; 3, loamy sand; 4, sandy loam; 5, sandy clay loam; 6, sandy silt loam; 7, silty clay |  |
| organic content | % loss of organic content on ignition | logl0 transformed |
| soil pH |  |  |
| available P | mg/L | logl0 transformed |
| available K | mg/L |  |
| Moist percentage moisture content |  |  |
| Bare percentage cover estimate of bare ground in 11 1-iM2 quadrats, arcsine transformed |  |  |
| Litter percentage cover |  | estimate of litter cover in 11 1-iM2 quadrats, log10 transformed |
| Bryophyte percentage cover |  | estimate of bryophytes in 11 1-iM2 quadrats, arcsine transformed |
| Plants/M2 number of reproductive stems |  | (flowering or fruiting) in 11 1-iM2 quadrats |
| Canopy height |  | canopy height (cm) in 11 -iM2 quadrats |
| Stem density number of stems |  | (ramets) in 100 cm2 |
| Biom 0-5 dry mass (g) of biomass 0-5 cm from soil surface in 400 cm2 |  |  |
| Biom 5+ dry mass (g) of biomass >5 cm from soil surface in 400 cm2, logl0 transformed |  |  |
| Repro biom biomass of reproductive parts (flowers and fruits) in 100 cm2, logl0 transformed |  |  |
| Elevation | m a.s.l. |  |
|  |  | (see Materials and Methods: Environmental data) |

**Functional traits:**

| Variable | Unit or factor levels |
| --- | --- |
| diameter of the eye, measured from above |  |
| length of the antenna |  |
| maximum width of the pronotum |  |
| maximum depth ("vaulting") of the pronotum |  |
| maximum width of the elytra |  |
| length of the metafemur | (with the articulation segments), from the coxa to the apex |
| length of the metatrochanter |  |
| length of the metatarsi |  |
| maximum width of the metafemur |  |
| total length | (length of the pronotum in the medial line plus length of the elytra, from the medial ridge of the scutellum to the apex) |
| color of the legs | 1, pale; 2, black; 3, metallic |
| color of the body | 1, pale; 2, black; 3, metallic |
| wing development | 1, apterous or brachypterous; 2, dimorphic; 3, macropterous |
| shape of the pronotum | 1, oval; 2, cordiform; 3, trapezoidal |
| overwintering | 1, only adults; 2, adults and larvae or only larvae |
| food of the adult | 1, mostly Collembola; 2, generalist predator; 3, mostly plant material |
| daily activity | 1, only diurnal; 2, nocturnal |
| breeding season | 1, spring; 2, summer; 3, autumn or winter |
| main period of emergence of the adults | 1, spring; 2, summer; 3, autumn |
| main period of adult activity | 1, autumn; 2, summer only |

**References:** Ribera, I., Dolédec, S., Downie, I.S. & Foster, G.N. (2001). Effect of land disturbance and stress on species traits of ground beetle assemblages. *Ecology*, 82, 1112–1129.

#### 110) N51TTP_3: Woody plants from Poland – Chmura *et al.* 2016

**Data owner(s)**: Damian Chmura

**Description:** A total of 364 decaying logs were studied in terms of their characteristics and the cover of vascular plants colonising them.

**Geographical scope:**	Karkonosze Mts, Sudeten Mts, Poland

**Taxonomical scope:** terrestrial vascular plants (woody plants)

**Ecological scope:** Terrestrial

**Number of sampling units:** 364 plots

**Minimum distance between sampling units: <** 0.01 kilometre

**Maximum distance between sampling units:** 80.21 kilometres

**Environmental variables:**

| Variable | Unit | Unit or factor levels | Description |
| --- | --- | --- | --- |
| Bryophytes cover | % | Percentage | The total percentage cover of all bryophytes including mosses and liverworts |
| Decomposition stage | none | 1-8 degree scale | Decomposition stage of downed trees |
| Length of a log | m | Meter | The length of the log |
| Area of a log surface | m² | Squere meter |  |
| Moisture of a log | none | (1) dry, completely dry to the touch, (2) intermediate, slightly perceptible moisture to the touch, (3) moist, water flows when pressure is applied | Moisture of a log |
| Shade | none | (1) <30% of canopy cover, (2) 30–40%, (3) –40–60%, (4) 60–80%, (5) >80% | Shadiness above a decaying log, was based on the canopy cover, which was visually estimated |
| Forest protection status | none | (1) protection, (0) management | The status of the forest |
| Altitude | m a.s.l. | Metres above sea level |  |
| Species log | none | (FAGUS) Fagus sylvatica, (PICEA) Picea abies | Tree speacies of the dead logs |
| Type of community | none | (FAGET) Fagetalia, (AP) Abieti Picetum, (CVP) Calamagrostio villosae Piceetum, (PAM) planted forest with Picea abies | The type of forest community in which a log was laying |

**Functional traits:**

| Variable | Unit | Unit or factor levels | Description |
| --- | --- | --- | --- |
| Species Leaf Area (mean value) | mm²/g | Square millimeter per gram |  |
| Plant height (mean value) | m | Meter |  |
| Seed mass | mg | (0) no seed, (>0) mean value [milligram] |  |
| Seed dispersal type | none | (AN) anemochory, (AUB) autochory, (BA) barochory, (ZOO) zoochory | The way how plant disperse it’s own seeds. There can be few types: (anemochory) by the wind; (autochory) self-dispersal; (barochory) by the gravity: (zoochory) by the animals |
| Rosette (leaf arrangement) | none | (1) yes, (0) no | Circular leaf arrangement |
| Semirosette (leaf arrangement) | none | (2) yes, (0) no | Semi-circular leaf arrangement |
| Regular (leaf arrangement) | none | (3) yes, (0) no | Regular leaf arrangement |
| Competitors (grime strategy) | none | (1) yes, (0) no | The survival of the individual using traits that maximize resource acquisition and resource control in consistently productive niches (Grime strategy is the adaptive plant strategy theory developed by the J.Philip Grime. The theory depicts adaptive strategies throughout the plant life, with extreme strategies facilitating the survival of genes) |
| Stress tolerators (grime strategy) | none | (1) yes, (0) no | Plant species with individual survival via maintenance of metabolic performance in variable and unproductive niches |
| Intermediate strategy (grime strategy) | none | (1) yes, (0) no | Plant species that have intermediate strategy characteristics in between stress tolerator, competitors and ruderal |
| Stress competitor (grime strategy) | none | (1) yes, (0) no | Plant species that survives by using traits that maximize resource acquisition and resource control in consistently productive niches due to stress |
| Competitive ruderal (grime strategy) | none | (1) yes, (0) no | Plant species with rapid gene propagation via rapid completion of the life-cycle and regeneration in niches where events are frequently lethal to the individual due to competition |
| Stress ruderal (grime strategy) | none | (1) yes, (0) no | Plant species with rapid gene propagation via rapid completion of the life-cycle and regeneration in niches where events are frequently lethal to the individual due to stress |
| Annuals (life span) | none | (1) yes, (0) no | Plant that completes its life cycle, from germination to the production of seeds, within one year, and then dies |
| Biannuals (life span) | none | (1) yes, (0) no | Flowering plant that takes two years to complete its biological life-cycle |
| Herb perennials (life span) | none | (1) yes, (0,5) half herb perennial, (0) no | Plant with herbal structure that lives more than two years |
| Woody perennials (life span) | none | (1) yes, (0,5) half woody perennials, (0) no | Plant with wooden structure that lives more than two years |

**References:** Chmura, D., Żarnowiec, J. & Staniaszek-Kik, M. (2016). Interactions between plant traits and environmental factors within and among montane forest belts: A study of vascular species colonising decaying logs. *For. Ecol. Manage.*, 379, 216–225.

#### 111) N53TTP: Plants data from the Netherlands – Jamil *et al*., 2013

**Data owner(s)**: Cajo ter Braak, Ruut Wegman

**Description:** The trait data were taken from the LEDA database by Jamil et al 2013. The spatial coordinates were retreived by georeferencing in GIS of the maps in Batterink & Wijffels (1993) by Ruut Wegman and Cajo ter Braak

**Geographical scope:**	Terschelling island, dune meadow, Netherlands

**Taxonomical scope:** Terrestrial vascular plants

**Ecological scope:** Terrestrial

**Number of sampling units:** 20 plots

**Minimum distance between sampling units:** 0.07	kilometres

**Maximum distance between sampling units:** 14.79 kilometres

**Environmental variables:**

| Variable | Description |
| --- | --- |
| A1 horizon thickness | Thickness of the A1 horizon |
| Moisture | Moisture content f the soil |
| Grassland management type | SF=standard farming, BF=biological farming, HF=hobby farming, NM=nature conservation management |
| Use | Agricultural grassland use (1) hay production (2) intermediate (3) grazing |
| Manure | Quantity of manure applied based on N and P manuring (N/P class in B&W 1983) |

**Functional traits:**

| Variable | Description |
| --- | --- |
| Specific Leaf Area | The trait data are taken from the LEDA database |
| Canopy height of a shoot |  |
| Lead dry matter content |  |
| Seed mass |  |
| Life span | Nominal; annual vs. perennial |

**References:**

Kleyer, M., Bekker, R.M., Knevel, I.C., Bakker, J.P., Thompson, K., Sonnenschein, M., *et al.* (2008). The LEDA Traitbase: a database of life‐history traits of the Northwest European flora. *J. Ecol.*, 96, 1266–1274.

Jongman, E. (1995). *Data analysis in community and landscape ecology*. Cambridge university press.

Jongman, R.G.H., Ter Braak, C.J.F. & Van Tongeren, O.F.R. (1987). Data Analysis in Community and Landscape Ecology, –Pudoc, Wageningen, 299pp.

Jamil, T., Ozinga, W.A., Kleyer, M. & ter Braak, C.J.F. (2013). Selecting traits that explain species–environment relationships: a generalized linear mixed model approach. *J. Veg. Sci.*, 24, 988–1000.

Batterink, M. (1983). Een vergelijkend vegetatiekundig onderzoek naar de typologie en invloeden van het beheer van 1973 tot 1982 in de duinweilanden op Terschelling.

#### 112) N55TBP: Bryophytes from Europe – Robroek *et al*. 2017

**Data owner(s)**: Bjorn Robroek

**Description:** Vegetation data from 56 Sphagnum-dominated peat bogs across Europe between 2010 and 2011.

**Geographical scope:**	Peat bogs, Western Europe.

**Taxonomical scope:** Bryophytes

**Ecological scope:** Terrestrial

**Number of sampling units:** 56 plots

**Minimum distance between sampling units:** 2.16 kilometres

**Maximum distance between sampling units:** 2688.27 kilometres

**Environmental variables:**

| Variable | Unit |
| --- | --- |
| Altitude | masl |
| Mean annual temperature | (°C) |
| Seasonality in temperature | (°C) |
| Mean annual precipitation | (mm) |
| Seasonality in precipitation | % |
| Total sulphur deposition SOx | mg m-2 yr-1 |
| Total oxidized nitrogen deposition NOy | mg m-2 yr-1 |
| Total reduced nitrogen deposition NHx | mg m-2 yr-1 |
| Lang’s moisture index | mm °C |

**Functional traits:**

| Variable | Unit |
| --- | --- |
| plant length | (mm) |
| spore diameter | (µm) |
| spore capsule diameter | (µm) |
| tissue carbon content | mgg-1 |
| tissue nitrogen content | mgg-1 |
| tissue phosphorous content | mgg-1 |
| Productivity |  |
| stem width | mm |
| length of hyalin cells | μm |
| width of hyalin cells | μm |
| length of stem leaves | mm |
| width of stem leaves | mm |

**References:** Robroek, B.J.M., Jassey, V.E.J., Payne, R.J., Martí, M., Bragazza, L., Bleeker, A., *et al.* (2017). Taxonomic and functional turnover are decoupled in European peat bogs. *Nat. Commun.*, 8, 1161.

#### 113) N55TAT : Carabid beetles from coniferous plantations in UK – Spake *et al*. 2016

**Data owner(s)**: Nadia Barsoum, Rebecca Spake

**Description:** Conifer plantation stands at 12 sites across the UK were

selected for study. Five traps were positioned 10 m apart on a north–south transect through the centre of each 1-ha plot and trapping was carried out over a 20-week period from May to September for two consecutive years (1995 - 1997) and emptied at fortnightly intervals.

**Geographical scope:**	Coniferous plantations, UK

**Taxonomical scope:** Insects (Carabid beetles)

**Ecological scope:** Terrestrial

**Number of sampling units:** 44 sites

**Minimum distance between sampling units: <** 0.01 kilometres

**Maximum distance between sampling units:** 759.80 kilometres

**Environmental variables:**

| **Variable** | | **Unit or factor levels** |
| --- | --- | --- |
| Percentage cover of open semi-natural area | 0–50; continuous | |
| Field, 10 cm – 1.9 m high | 0–75; continuous | |
| Shrub, 2–5 m high | | 0–40; continuous |
| Lower canopy, 5.1–15 m high | | 0–55; continuous |
| Upper canopy, 15.1–20 m high | | 0–30; continuous |
| Ground vegetation diversity | | 0.000–0.144; continuous |

**Functional traits:**

| Variable | Unit or factor levels |
| --- | --- |
| Body length | Continuous/mm 2.95–30 |
| Adult feeding guild | Categorical, Collembola specialist/generalist predator/phytophagous/omnivorous |
| Hind-wing morphology | Categorical, Macropterous/dimorphic/apterous or brachypterous |
| Activity pattern | Categorical, Diurnal/nocturnal |
| Adult habitat affinity | Categorical, Forest/open/generalist |
| Overwinter type | Categorical, Adult only/larvae or adult |

**References:** Spake, R., Barsoum, N., Newton, A.C. & Doncaster, C.P. (2016). Drivers of the composition and diversity of carabid functional traits in UK coniferous plantations. *For. Ecol. Manage.*, 359, 300–308.

#### 114) N55TTP_2: Plants from Europe – Robroek *et al*. 2017

**Data owner(s)**: Bjorn Robroek

**Description:** Vegetation data from 56 Sphagnum-dominated peat bogs across Europe between 2010 and 2011.

**Geographical scope:**	Peat bogs, Western Europe.

**Taxonomical scope:** Terrestrial vascular plants

**Ecological scope:** Terrestrial

**Number of sampling units:** 56 sites

**Minimum distance between sampling units:** 2.16 kilometres

**Maximum distance between sampling units:** 2688.27 kilometres

**Environmental variables:**

| Variable | Unit |
| --- | --- |
| Altitude | masl |
| Mean annual temperature | (°C) |
| Seasonality in temperature | (°C) |
| Mean annual precipitation | (mm) |
| Seasonality in precipitation | % |
| Total sulphur deposition SOx | mg m-2 yr-1 |
| Total oxidized nitrogen deposition NOy | mg m-2 yr-1 |
| Total reduced nitrogen deposition NHx | mg m-2 yr-1 |
| Lang’s moisture index | mm °C |

**Functional traits:**

| Variable | Unit |
| --- | --- |
| Specific leaf area | (cm2 g-1) |
| Canopy_height | m |
| Leaf dry matter content | (g g-1) |
| Seed_mass | g |
| Seed_number | numeric |

**References:** Robroek, B.J.M., Jassey, V.E.J., Payne, R.J., Martí, M., Bragazza, L., Bleeker, A., *et al.* (2017). Taxonomic and functional turnover are decoupled in European peat bogs. *Nat. Commun.*, 8, 1161.

#### 115) S05MFI: Fish communities from the Seychelles archipelago – Chong-Seng *et al*. 2012

**Data owner(s)**: Karen Chong-Seng

**Description:** Twenty-one carbonate fringing reefs within a 3600 km² area around the inner Seychelles islands (4 309S, 55 309E) were surveyed in October 2010.

**Geographical scope:**	Seychelles archipelago

**Taxonomical scope:** Fishes

**Ecological scope:** Marine

**Number of sampling units:** 79 sites

**Minimum distance between sampling units:** < 0.01 kilometres

**Maximum distance between sampling units:** 88.03 kilomettres

**Environmental variables:**

| Variable | Unit or factor levels | Description |
| --- | --- | --- |
| Acropora coral | proportional cover along 50m transects | Small polyp stony coral |
| Crustose coralline algae | proportional cover along 50m transects | encrusing Algal speceis that is characterized by calcareous deposits contained within the cell walls |
| Leathery macroalgae | proportional cover along 50m transects | Macroalgae that have leathery body surface |
| Non biological particles | proportional cover along 50m transects | non-living - e.g. sand, bare exposed rock surfaces |
| Other benthic organisms | proportional cover along 50m transects | Noncoral or algae benthic organisms (corallimorphs, sponges and zoanthids) |
| Other scleractinian corals (hard corals) | proportional cover along 50m transects | All other genera (hard corals) scleractinian corals (branching acroporids, massive Porites, and favids coral) |
| Other macroalgae | proportional cover along 50m transects | all other macroalgae (algae >5cm high/diameter) e.g. Dictyota, Halimeda |
| Massive poritiid corals | proportional cover along 50m transects | Poritiid corals present stony corals |
| Turf algae | proportional cover along 50m transects | "a ubiquitous, multispecific assemblage of diminutive algae" (1-10mm height) cf. Steneck 1988 - can provide pdf if necessary |
| Total scleractinian corals (hard corals) | proportional cover along 50m transects | this is a result of sum(Acropora, OtherHC, PoritiidMassive) |
| Rugosity | score from 0.5 up to 5 | Rugosity is a simple measurement of the surface roughness that has been used routinely by coral reef biologists |
| Holes (number or size) | number of holes | Number of small refuge holes, <10 cm diameter, along two 10x1 m sub-transects |
| Consolidated rubble (underlaying substrate) | proportional cover along 50m transects | Consolidated rubble in the underlaying substratum present layer below recorded benthic cover or the top 10 mm of sand/sediment |
| Bommie (underlaying substrate) | proportional cover along 50m transects | Isolated coral outcrops |
| Pavement (underlaying substrate) | proportional cover along 50m transects | Solid carbonate pavement (hard surface) |
| Rubble (underlaying substrate) | proportional cover along 50m transects | The piles of broken stones |
| Sand (underlaying substrate) | proportional cover along 50m transects |  |
| Reef name | AC, AP, BO, AS, BT, BV, CE, CF CL, GC, GP, GS, MY, NE, PL, PS, SA, SP | Reef names. Each abreviation presents one reef |

**Functional traits:**

| Variable | Unit | Description |
| --- | --- | --- |
| Fish functional group | Herbivore.browser, herbivore.nonbrowsing, planktivore, invertivore, piscivore, omnivore, corallivore, carnivore | Funcitonal groups based on feeding characteristics of a fish |
| Fishing pressure | (N) not fished, (O) occasional by-catch, (I) important by-catch, (P) primary target | Level of fish species exploitation through fishing |

**References:** Chong-Seng, K.M., Mannering, T.D., Pratchett, M.S., Bellwood, D.R. & Graham, N.A.J. (2012). The influence of coral reef benthic condition on associated fish assemblages. *PLoS One*, 7, 1–10.

#### 116) S05MFI_2: Fish communities from the Seychelles archipelago – Chong-Seng *et al*. 2012

**Data owner(s)**: Karen Chong-Seng

**Description:** Twenty-one carbonate fringing reefs within a 3600 km² area around the inner Seychelles islands (4 309S, 55 309E) were surveyed in October 2010.

**Geographical scope:**	Seychelles archipelago

**Taxonomical scope:** Fishes

**Ecological scope:** Marine

**Number of sampling units:** 79 sites

**Minimum distance between sampling units:** < 0.01 kilometres

**Maximum distance between sampling units:** 88.03 kilomettres

**Environmental variables:**

| Variable | Unit or factor levels | Description |
| --- | --- | --- |
| Acropora coral | proportional cover along 50m transects | Small polyp stony coral |
| Crustose coralline algae | proportional cover along 50m transects | encrusing Algal speceis that is characterized by calcareous deposits contained within the cell walls |
| Leathery macroalgae | proportional cover along 50m transects | Macroalgae that have leathery body surface |
| Non biological particles | proportional cover along 50m transects | non-living - e.g. sand, bare exposed rock surfaces |
| Other benthic organisms | proportional cover along 50m transects | Noncoral or algae benthic organisms (corallimorphs, sponges and zoanthids) |
| Other scleractinian corals (hard corals) | proportional cover along 50m transects | All other genera (hard corals) scleractinian corals (branching acroporids, massive Porites, and favids coral) |
| Other macroalgae | proportional cover along 50m transects | all other macroalgae (algae >5cm high/diameter) e.g. Dictyota, Halimeda |
| Massive poritiid corals | proportional cover along 50m transects | Poritiid corals present stony corals |
| Turf algae | proportional cover along 50m transects | "a ubiquitous, multispecific assemblage of diminutive algae" (1-10mm height) cf. Steneck 1988 - can provide pdf if necessary |
| Total scleractinian corals (hard corals) | proportional cover along 50m transects | this is a result of sum(Acropora, OtherHC, PoritiidMassive) |
| Rugosity | score from 0.5 up to 5 | Rugosity is a simple measurement of the surface roughness that has been used routinely by coral reef biologists |
| Holes (number or size) | number of holes | Number of small refuge holes, <10 cm diameter, along two 10x1 m sub-transects |

**Functional traits:**

| Variable | Unit or factor levels |
| --- | --- |
| Functional group | Herbivore.browser; Herbivore.nonbrowsing; Planktivore; Invertivore; Piscivore; Omnivore; Corallivore; Carnivore |
| FishingPressure | N=not fished, O=occasional by-catch, I=important by-catch, P=primary target |

**References:** Chong-Seng, K.M., Mannering, T.D., Pratchett, M.S., Bellwood, D.R. & Graham, N.A.J. (2012). The influence of coral reef benthic condition on associated fish assemblages. *PLoS One*, 7, 1–10.

#### 117) N59TTP: Vegetation from Sweden - Marteinsdóttir & Eriksson 2014.

**Data owner(s)**: Bryndis Marteinsdottir

**Description:** Six grazed ex-arable fields and eight semi-natural grasslands in south- east Sweden.

**Geographical scope:**	southeast Sweden

**Taxonomical scope:** Terrestrial vascular plants

**Ecological scope:** Terrestrial

**Number of sampling units:** 14 sites

**Minimum distance between sampling units:** < 0.01 kilometre

**Maximum distance between sampling units:** 3.68 kilometres

**Environmental variables:**

| Variable | Unit | Unit or factor levels | Description |
| --- | --- | --- | --- |
| pH | pH |  | Soil pH |
| Ammonium | mg 100¯¹ | milligram per cent | Soil Amonium |
| Phosphorus | mg 100¯¹ | milligram per cent | Soil Phosphorus |
| Moisture | % | percentage | Soil Moisture |
| Potassium | mg 100¯¹ | milligram per cent | Soil Potassium |
| Magnesium | mg 100¯¹ | milligram per cent | Soil Magnesium |
| Calcium | mg 100¯¹ | milligram per cent | Soil Calcium |

**Functional traits:**

| Variable | Unit | Unit or factor levels | Description |
| --- | --- | --- | --- |
| Specific leaf area | mm² mg¯¹ | millimetre squere pre milligram | Specific leaf area (SLA) is the one-sided area of a fresh leaf, divided by its oven-dry mass. |
| Leaf drymatter content | mg g¯¹ | milligram per gram | Leaf drymatter content (LDMC) is the oven-dry mass (mg) of a leaf, divided by its water-saturated fresh mass (g). |
| Seed mass | mg | milligram |  |

**References:** Marteinsdóttir, B. & Eriksson, O. (2014). Plant community assembly in semi-natural grasslands and ex-arable fields: A trait-based approach. *J. Veg. Sci.*, 25, 77–87.

#### 118) N51TBI: Birds from German wasteland – Meffert *et al*. 2013.

**Data owner(s)**: Peter J. Meffert

**Description:** A set of 54 wasteland sites of early successional stages that scattered over the entire urban area of Berlin, Germany.

**Geographical scope:**	Urban wasteland, Berlin, Germany

**Taxonomical scope:** Birds

**Ecological scope:** Terrestrial

**Number of sampling units:** 54 sites

**Minimum distance between sampling units:** 0.15 kilometres

**Maximum distance between sampling units:** 32.53 kilometres

**Environmental variables:**

| Variable | Unit | Unit or factor levels | Description |
| --- | --- | --- | --- |
| Population within 50 m | Inhabitants/ha | Number of inhabitant per hectare | Human populaiton denisty within 50 m (Inhabitants/ha) |
| Population within 200 m | Inhabitants/ha | Number of inhabitant per hectare | Human populaiton denisty within 200 m (Inhabitants/ha) |
| Sealing within 50 m | % | degree | Degree of sealing within 50 m |
| Sealing within 2000 m | % | degree | Degree of sealing within 2000 m |

**Functional traits:**

| Variable | Unit | Unit or factor levels | Description |
| --- | --- | --- | --- |
| Food type | none | insectivorous, omnivorous, granivorous |  |
| Foraging technique | none | ground gleaner, tree/foliage gleaner, sit-and-wait predator, |  |
| Adult survival | % | Percentage |  |
| Innovation rate | % | Percentage | A novel behaviour is a behaviour that the person would not have engaged in before while engaging in the targeted behaviour. Often these novel behaviour can be aggressive or more emotional (Miltenberger, 2008).It was calculated from numbers of novel behaviours reported in journals that are included in the Zoological Records’ web index published by Overington et al. (2009). |
| Migration strategy | none | long (long distance migrants, usually to Africa), short (short distance), sedentary |  |
| Body mass | g | grams |  |
| Nesting location | none | ground, shrub or tree, building |  |
| Nestling period | none | number of days | Time period that young bird is not yet old enough to leave the nest; The nestling period is the average time in days that chicks spend in the nest. |

**References:** Meffert, P.J. & Dziock, F. (2013). The influence of urbanisation on diversity and trait composition of birds. *Landsc. Ecol.*, 28, 943–957.

#### 119) S01TBI: Birds communities from Indonesia – Cleary *et al.* 2007

**Data owner(s)**: Daniel Cleary, Nicole de Voogd, Willem Renema

**Description:** The authors assessed bird species composition within a logging concession in Central Kalimantan, Indonesian Borneo. Within the study area (196 km2) a total of 9747 individuals of 177 bird species were recorded.

**Geographical scope:**	Mentaya river, Central Kalimantan province, Borneo, Indonesia

**Taxonomical scope:** Birds

**Ecological scope:** Terrestrial

**Number of sampling units:** 37 sites

**Minimum distance between sampling units:** 0.17 kilometres

**Maximum distance between sampling units:** 13.96 kilometres

**Environmental variables:**

| Variable | Unit | Unit or factor levels | Description |
| --- | --- | --- | --- |
| Logging | ordinal var | (1) primary unlooged, (2) logged in 1989, (3) logged in 1993 |  |
| Slope position | none | (1) low, (2) middle, (3) upper |  |
| Elevation | m | m a.s.l. | Meters above see level |
| Short saplings | none | density | Saplings < 5 m and < 5 cm dbh (diameter at breast hight) |
| Tall saplings | none | density | Saplings > 5 m and > 5 cm dbh |
| Short poles | none | density | Poles heigh <10 m and 5–10 cm dbh. Poles are long, relatively slender, generally rounded piece of wood. |
| Tall poles | none | density | Poles heigh>10 m and 5–10 cm dbh |
| Tree density | none | density | Number of all trees per unit area with dbh >10cm of the tree of average basal area |
| Ferns cover | none | abundance |  |
| Grass cover | none | abundance |  |
| Herbs cover | none | abundance |  |
| Seedlings | none | abundance |  |
| Large woody lianas abundance | none | abundance | Large woody lianas (stem diameter > 5 cm) |
| Small woody lianas | none | abundance | Small woody lianas (stem diameter < 5 cm) |
| Non-woody liana abundance | none | abundance |  |
| Epiphytes | none | abundance | Organism that grows on the surface of a plant and derives its moisture and nutrients from the air, rain, water or from debris accumulating around it |
| Bryophytes | none | abundance | Bryophytes are an informal group consisting of three divisions of non-vascular land plants: the liverworts, hornworts and mosses |
| Dead wood, state 1 | none | volume | Fresh dead wood |
| Dead wood, state 2 | none | volume | Dead wood with sound wood but flaking bark, dead wood |
| Dead wood, state 3 | none | volume | Dead wood with sound wood but no bark |
| Dead wood, state 4 | none | volume | Dead wood with rotting wood but firm |
| Dead wood, state 5 | none | volume | Dead wood with wood rotten and soft |
| Dead wood, falen | none | volume | Fallen dead wood |
| Dead wood, standings | none | volume | Standing dead wood |
| Dead wood total | none | volume | Dead wood total |
| Mean litter depth | mm | milimeter | Dead plant material, such as leaves, bark, and twigs, that has fallen to the ground |
| Small woody chips | abundance | abundance | Dead wood with diameter <10 cm |
| Mesophyll leaf litter cover | mm² | square millimeters | Mesophyll leaf litter cover with size from 4500 to 20 000 mm² |
| Notophylls leafs litter cover | mm² | square millimeters | Notophylls leafs litter cover with size from 2000 to 4500 mm² |
| Microphyll leafs litter cover | mm² | square millimeters | Microphyll leaves litter cover with size up to 2000 mm² |
| Conopy cover | cover |  | The aboveground portion of a plant community, formed by the collection of individual plant crowns. |
| Diameter at breast height | cm | centimeter |  |
| Tree height | m | meter |  |
| Bifurcation index | none | none |  |
| Tree crown depth | m | meter | The crown dept of a plant refers to dsitance from the lowest point of the crown till the top of the tree |
| Tree crown radius | m | meter | Rarius of the tree crown |

**Functional traits:**

| Variable | Unit or factor levels | Description |
| --- | --- | --- |
| Feeding Guild | (Rap) raptors, (C-Bar-In) canopy break-gleaning insectivores, (C-Fol-Fr) canopy foliage-gleaning frugivores, (C-Fol-IF) canopy foliage-gleaning insectivore/frugivores, (C-Fol-In) canopy foliage-gleaning insectivores, (C-Nec) canopy nectarivores, (C-S-In) canopy sallying insectivores, (Fr-Car) frugivore/carnivores, (T-In) terrestrial insectivores, (U-Bar-In) understory bark-gleaning insectivores, (U-Fol-IF) understory foliage-gleaning insectivore/frugivores, (U-Fol-In) understory foliage-gleaning insectivores, (Ug-In) undergrowth specialist insectivores, (U-Nec) understory nectarivores, (U-S-In) understory sallying insectivores, (U-SS-In) understory sallying substrate insectivores | Any group of species that exploit the same resources, or that exploit different resources in related ways |
| Global distribution | (D-End) endemics, (D-Non) non-endemics |  |
| Size | (S-VS) very small [<10 cm], (S-S) small [10 cm<x<20 cm], (S-M) medium [20 cm<x<30 cm], (S-L) large [30 cm<x<60 cm], (S-VL) very large [>60cm] | Classiciation of the bird species size according to wing size |
| Conservation status | (R-T) threatened, (R-NT) not threatened |  |

**References:** Cleary, D.F.R., Boyle, T.J.B., Setyawati, T., Anggraeni, C.D., Van Loon, E.E. & Menken, S.B.J. (2007). Bird species and traits associated with logged and unlogged forest in Borneo. *Ecol. Appl.*, 17, 1184–1197.

#### 120) S02TBT: Amazonian bats – Farneda *et al.* 2015

**Data owner(s)**: Fábio Z. Farneda, Adrià López-Baucells, Christoph Meyer, Ricardo Rocha	.

**e-mail**:

**Description:** Bats were captured over two years in eight forest fragments, nine control sites in continu- ous forest, and in the secondary forest matrix at the Biological Dynamics of Forest Frag- ments Project, Central Amazon, Brazil.

**Geographical scope:**	Biological Dynamics of Forest Fragments Project (BDFFP) located ca. 80 km north of Manaus, Central Amazon, Brazil.

**Taxonomical scope:** Mammals (Bats)

**Ecological scope:** Terrestrial

**Number of sampling units:** 17

**Minimum distance between sampling units:** 0.48	kilometres

**Maximum distance between sampling units:** 41.28 kilometres

**Environmental variables:**

| Code | Variable | Unit or factor levels | Description |
| --- | --- | --- | --- |
| NT | Average number of trees | numeric | Number of trees ≥ 10 cm diameter at breast height (DBH) |
| BT | Average tree diameter | cm | Average DBH of trees ≥ 10 cm |
| VSVD | Average vertical stratification | m | Vertical stratification in vegetation density; visually classified at 5-m intervals across each plot as the sum of the values obtained by the estimation at seven height intervals [0–1, 1–2, 2–4, 4–8, 8–16, 16–24 and 24–32 m] using six categories varying from no to very dense vegetation |
| L | Average number of lianas | numeric | Number of lianas |
| R | Average number of potential roosts | numeric | Number of potential roosts |
| VC | Average number of Vismia and Cecropia trees | numeric | Number of Vismia and Cecropia trees |
| TH | Average mean tree height | m | Mean tree height (m) |
| FC | Average forest cover | % | Forest cover (%) within a 1.5 km buffer centered on each sampling site |
| Habitat | Habitat type | 4 categories: fragment of 1ha, fragment of 10ha, fragment of 100ha, cotinuous forest |  |

**Functional traits:**

| Code | Variable | Unit or factor levels | Description |
| --- | --- | --- | --- |
| trophic_level | trophic_level | animalivorous or phytophagous |  |
| habitat_classification | habitat_classification |  |  |
| body_mass | body_mass | grams | Based on the average body mass of each species |
| dietary_specialization | dietary_specialization | low, intermediate, high | Based on the percentage of the contribution of each food item to the total dietary records for each species |
| vertical_stratification | vertical_stratification |  | Based on capture rate/mnh using the proportion of captures in ground vs. canopy nets in interior of fragments and of continuous forest sites. |
| mobility | mobility |  | Based on the relationship between the mean and the maximum recapture distance; species were grouped into three categories of mobility: low, intermediate and high |
| aspect_ratio | aspect_ratio |  | wingspan2/wing area |
| relative_wing_loading | wing morphology |  | wing loading [body mass multiplied by the gravitational acceleration, 9.81 ms2/wing area]/body mass0.33 |

**References:** Farneda, F.Z., Rocha, R., López-Baucells, A., Groenenberg, M., Silva, I., Palmeirim, J.M., *et al.* (2015). Trait-related responses to habitat fragmentation in Amazonian bats. *J. Appl. Ecol.*, 52, 1381–1391.

#### 121) S33TAT: Australian spiders – Gibb *et al.* 2015

**Data owner(s)**: Heloise Gibb

**Description:** Data of foliage-living spider assemblages associated with *Themeda triandra* grasslands along a 900 km climatic gradient in south-eastern Australia. The authors used sweep-netting to collect T. triandra-associated spiders and counted juveniles and identified adults.

**Geographical scope:**	Themeda grasslands, south-east Australia

**Taxonomical scope:** Insects (Spiders)

**Ecological scope:** Terrestrial

**Number of sampling units:** 36 sites

**Minimum distance between sampling units:** 6.90	kilometres

**Maximum distance between sampling units:** 1196.70 kilometres

**Environmental variables:**

| Variable | Unit or factor levels |
| --- | --- |
| Elevation | m |
| Temperature | °C |
| Precipitation | mm |
| Grass height |  |
| Intra |  |
| Inter |  |
| Disturbance | level of disturbance |

**Functional traits:**

| Variable | Unit or factor levels |
| --- | --- |
| Sex | M/F |
| Body length | mm |
| Abdomen length | mm |
| Abdomen width | mm |
| Cephalothorax width | mm |
| Fang length (left) | mm |
| Fang length (right) | mm |
| Tibia length (left) | mm |
| Tibia length (right) | mm |
| Asymmetry |  |
| Pilosity |  |

**References:** Gibb, H., Muscat, D., Binns, M.R., Silvey, C.J., Peters, R.A., Warton, D.I., *et al.* (2015). Responses of foliage-living spider assemblage composition and traits to a climatic gradient in Themeda grasslands. *Austral Ecol.*, 40, 225–237.

#### 122) S21TAT: Brazilian spiders – Gonçalves-Souza *et al*. 2014

**Data owner(s)**: Thiago Gonçalves-Souza

**Description:** The authors sampled 12 localities of restinga vegetation along 2040 km of the Brazilian coast.

**Geographical scope:**	Open restingas, Atlantic rainforest, Brazil

**Taxonomical scope:** Insects (Spiders)

**Ecological scope:** Terrestrial

**Number of sampling units:** 17 sites

**Minimum distance between sampling units:** 0.48 kilometres

**Maximum distance between sampling units:** 41.28 kilometres

**Environmental variables:**

| Variable | Unit | Unit or factor levels |
| --- | --- | --- |
| Prosoma height | cm | Centimeter |
| Prosoma width | cm | Centimeter |
| Prosoma length | cm | Centimeter |
| Opistosoma length | cm | Centimeter |
| Body compression | cm | Centimeter |
| Orbicular web type | none | (1) yes, (0) no |
| Sheet web type | none | (1) yes, (0) no |
| Space cob wer type | none | (1) yes, (0) no |
| Ambush hunter | none | (1) yes, (0) no |
| Active hunter | none | (1) yes, (0) no |
| Shelter | none | (1) yes, (0) no |
| Diurnal | none | (1) yes, (0) no |
| Nocturnal | none | (1) yes, (0) no |
| Diuturno | none | (1) yes, (0) no |
| Dorsal eyes | none | (1) yes, (0) no |
| Dorsofrontal | none | (1) yes, (0) no |
| Tapetum | none | (1) yes, (0) no |
| Claw tuft | none | (1) yes, (0) no |
| Scopula tuft | none | (1) yes, (0) no |
| None of mentioned | none | (1) yes, (0) no |

**Functional traits:**

| Variable | Unit | Unit or factor levels |
| --- | --- | --- |
| Plant species | none | AechmeaLingulata, BROMELIA1, ByrsonimaMicrophora, DodoniaViscosa, GRANDE, HohenbergiaLitorallis, MEDIA, MIUDA, NeoregeliaCruenta, PALMEIRA, QuesneliaArvensis, REDONDA, TibouchinaClavatium, VriseaProcera |
| Crown height | cm | centimeter |
| Higher crown length | cm | centimeter |
| Lower crown length | cm | centimeter |
| Leaf length | cm | centimeter |
| Leaf width | cm | centimeter |
| Distance between the second and third leaves | cm | centimeter |
| Ratio between leaf width and length | cm | centimeter |
| Volume | cm | centimeter |
| Biomass | g | gram |

**References:** Gonҫalves-Souza, T., Romero, G.Q. & Cottenie, K. (2014). Metacommunity versus biogeography: A case study of two groups of neotropical vegetation-dwelling arthropods. *PLoS One*, 9, 1–20.

#### 123) S28TAT: Insects from Australia – Yates *et al*. 2014

**Data owner(s)**: Nigel Andrew

**Description**: Ant assemblages were collected at nine paired pasture and remnant sites fromwithin three areas along a 300 km distance.

**Geographical scope:** Pasture vs remnant vegetation, North east of New South Wales, Australia

**Taxonomical scope:** Insects (Ants)

**Ecological scope:** Terrestrial

**Number of sampling units:** 18 sites

**Minimum distance between sampling units:** < 0.01 kilometres

**Maximum distance between sampling units:** 340 kilometres

**Environmental variables:**

| Variable | Unit or factor levels |
| --- | --- |
| soil carbon-nitrogen ratio |  |
| soil p content |  |
| tall grass cover | proportion |
| short grass cover | proportion |
| herb cover | proportion |
| leaf litter cover | proportion |
| bare ground cover | proportion |
| Average ambient daily temperature | degrees Celsius |

**Functional traits:**

| Variable | Description |
| --- | --- |
| eye width | ability to see laterally |
| head length | may be indicative of diet; longer head length may indicate herbivory |
| mandible length | indicative of diet; longer mandibles could allow predation of larger prey |
| top tooth length | may function to cut and masticate. Longer top teeth may increase funcitonal cpomlexity, increasin gability to cut and break down plant paterial |
| scape lengtjh | mechano and chemoreception |
| max. hait length (alitrunk) | heirs may increase tolerance to dehydration, they may funtion in thermo-regulation or relatve to mechanoreception |
| mid-femur length | linked to locomotion and climbing ability |

**References:** Yates, M.L., Andrew, N.R., Binns, M. & Gibb, H. (2014). Morphological traits: Predictable responses to macrohabitats across a 300 km scale. *PeerJ*, 2014, 1–20.

#### 124) S22TAT Brazilian spiders – Gonçalves-Souza *et al*. 2014

**Data owner(s)**: Thiago Gonçalves-Souza

**Description:** The authors sampled 12 localities of restinga vegetation along 2040 km of the Brazilian coast.

**Geographical scope:**	Open restingas, Atlantic rainforest, Brazil

Taxonomical scope: Insects (Spiders)

**Ecological scope:** Terrestrial

**Number of sampling units:** 309 sites

**Minimum distance between sampling units:** < 0.01 kilometre

**Maximum distance between sampling units:** 1991.14 kilometres

**Environmental variables:**

| Variable | Unit | Unit or factor levels |
| --- | --- | --- |
| Plant species | none | AechmeaLingulata, BROMELIA1, ByrsonimaMicrophora, DodoniaViscosa, GRANDE, HohenbergiaLitorallis, MEDIA, MIUDA, NeoregeliaCruenta, PALMEIRA, QuesneliaArvensis, REDONDA, TibouchinaClavatium, VriseaProcera |
| Crown height | cm | centimeter |
| Higher crown length | cm | centimeter |
| Lower crown length | cm | centimeter |
| Leaf length | cm | centimeter |
| Leaf width | cm | centimeter |
| Distance between the second and third leaves | cm | centimeter |
| Ratio between leaf width and length | cm | centimeter |
| Volume | cm | centimeter |
| Biomass | g | gram |

**Functional traits:**

| Variable | Unit | Unit or factor levels |
| --- | --- | --- |
| Guild | none | Hunting, web building |
| Prosoma height | cm | centimeters |
| Prosoma width | cm | centimeters |
| Prosoma length | cm | centimeters |
| Opistosoma length | cm | centimeters |
| Body compression | cm | centimeters |
| Orbicular web type | none | (1) yes, (0) no |
| Sheet web type | none | (1) yes, (0) no |
| Space cob wer type | none | (1) yes, (0) no |
| Ambush hunter | none | (1) yes, (0) no |
| Active hunter | none | (1) yes, (0) no |
| Shelter | none | (1) yes, (0) no |
| Diurnal | none | (1) yes, (0) no |
| Nocturnal | none | (1) yes, (0) no |
| Diuturno | none | (1) yes, (0) no |
| Dorsal eyes | none | (1) yes, (0) no |
| Dorsofrontal | none | (1) yes, (0) no |
| Tapetum | none | (1) yes, (0) no |
| Claw tuft | none | (1) yes, (0) no |
| Scopula tuft | none | (1) yes, (0) no |
| None of mentioned | none | (1) yes, (0) no |

**References:** Gonҫalves-Souza, T., Romero, G.Q. & Cottenie, K. (2014). Metacommunity versus biogeography: A case study of two groups of neotropical vegetation-dwelling arthropods. *PLoS One*, 9, 1–20.

#### 125) S34TAT: Spiders along an urbanization gradient in Sydney, Australia – Lowe *et al*. 2018

**Data owner(s)**: Elizabeth C Lowe, Jonas Wolff

**Description: S**amples were taken along a gradient of urbanisation extending 50 km north and south of the Sydney Central Business District (CBD) and around 20 km wide.

**Geographical scope:**	Urban gradient, Sydney, Australia

**Taxonomical scope:** Insects (Spider)

**Ecological scope:** Terrestrial

**Number of sampling units:** 115 sites

**Minimum distance between sampling units:** 0.01	kilometres

**Maximum distance between sampling units:** 53.80 kilometres

**Environmental variables:**

| Variable | Unit | Unit or factor levels | Description |
| --- | --- | --- | --- |
| Land use type | none | Bush, Remnant, Park, Garden |  |
| Microhabitat (0-50 cm) | % | Percentage | Vegetastion habitat complexity that is quatified at 0 - 50 cm, calculated by using a visual estimation of the percentage of vegetation cover to the nearest |
| Microhabitat (50-100 cm) | % | Percentage | Vegetation habitat complexity that is quatified at 50-100 cm, calculated by using a visual estimation of the percentage of vegetation cover to the nearest |
| Microhabitat (100-200 cm) | % | Percentage | Vegetation habitat complexity that is quatified at 100-200 cm, calculated by using a visual estimation of the percentage of vegetation cover to the nearest |
| Leaf litter in microhabitat | % | Percentage | Percentige of the leaf litter in microhabitat |
| Rocks and sticks in microhabitat | % | Percentage | Percentige of the rocks and sticks in microhabitat |
| Native vegetation in microhabitat | % | Percentage | Percentige of the native vegetation in microhabitat |
| Hard surface in microhabitat | % | Percentage | Percentige of the hard surface in microhabitat |
| Grass in microhabitat | % | Percentage | Percentige of the grass in microhabitat |
| Vegetation in microhabitat | % | Percentage | Percentage of the vegetation in microhabitat |
| Native vegetation within 500 m | % | Percentage | Landscape within 500 m covered by native vegetation. Calculated as the amount per hectare. |
| Housing within 500 m | amount/ha | Percentage | Landscape within 500 m used for housing. Percent cover . |
| Human population denisty within 500 m | residents/ ha | residents per hectare | Population number within 500 m. Calculated as the amount per hectare. |
| Comerc and hospital within 500 m | % | Percentage | Landscape within 500 m used the comercial/hospital purposes. Percent cover . |
| Eduaction within 500 m | % | Percentage | Landscape within 500 m used for the educational purposes. Percent cover . |
| Industry within 500 m | % | Percentage | Landscape within 500 m used for the undustrial purposes. Percent cover . |
| Residence within 500 m | % | Percentage | Landscape within 500 m used for the residentail purposes. Percent cover . |
| Parks within 500 m | % | Percentage | Landscape within 500 m used for the parks/agrycultural purposes. Percent cover . |
| Percentage of vegetation within 500 m | % | Percentage | Percent coverage of vegetatoin within 500 m |
| Percentage of hard land within 500 m | % | Percentage | Percent coverage of hard land within 500 m |
| Percentage of parks within 500 m | % | Percentage | Percent coverage of parks within 500 m |
| Native vegetation are within 2 km | % | Squere meters | Area within 2 km coverd by the native vegetation |
| Commercial are within 2 km | % | Percentage | Area within 2 km used for the comerce purposes |
| Education are within 2 km | % | Percentage | Area within 2 km used for the educaitonal purposes |
| Hospital/medical are within 2 km | % | Percentage | Area within 2 km used for hosputals/medical purpose |
| Industrial are within 2 km | % | Percentage | Area within 2 km used for the industial purposes |
| Other purposes area within 2 km | % | Percentage | Area within 2 km used for other purposes |
| Park/agryculture are within 2 km | % | Percentage | Area within 2 km used for the park purposes |
| Residential are within 2 km | % | Percentage | Area within 2 km used for the residential purposes |
| Transportation are within 2 km | % | Percentage | Area within 2 km used for the transportation purposes |
| Water areas within 2 km | amount/ha | Percentage | Area within 2 km coverd by the water |
| Dwellings within 2 km | amount/ha | amount per hectare | Numebr of dwellings within 2 km |
| Human population denisty within 2 km | residents/ ha | residents per hectare | Number of people within 2 km |

**Functional traits:**

| Variable | Unit | Unit or factor levels | Description |
| --- | --- | --- | --- |
| Guild | none | Web builder, Hunter | (Web builder) Builds silken snare to capture prey; (Hunter) Hunts without the aid of a web (but may actively deploy silk to immobilize prey) |
| Hunting style | none | Orb web, Cobweb, Reduced web, Sheet web, Lace web, Ambusher, Foliage runner, Ground runner, Kleptoparasite | (Orb web) builds an orb web to capture prey; (Cobweb) Builds a space web with knock-down- or tangle-lines, occasionally with gumfooted; (Sheet web) Build suspended or substrate bound horizontal sheets to capture prey; (Lace web) Builds cribellar lace webs to capture prey; (Reduced web) Strongly modified web, which is used in conjunction with a special hunting strategy; (Foliage runner) Non-web building hunter that actively searches and pursues prey in above-ground habitats; may use sticky silk to immobilize prey; (Ground runner) Non-web building hunter that actively searches and pursues prey on the ground; may use sticky silk to immobilize prey; (Ambusher) Non-web building hunter that waits for prey to come by; (Kleptoparasite) Steals food from webs of other spiders |
| Capture lines | none | Cribellar silk, Viscid silk, Non-adhesive, None | (Cribellar silk) Utilizes cribellar silk (dry nanofiber-bundles) for prey capture; (Viscid silk) Utilizes glue coated capture lines; (Non-adhesive) Adhesive threads are absent in web; (None) Does not build webs for prey capture |
| Body Size | none | Small, Medium, Large | (Small) Body length <5 mm; (Medium) Body length 5-10 mm; (Large) Body length > 10 mm |
| Period of activity | none | Durnal, Nocturnal | (Diurnal) Active during day; (Nocturnal) Active at night |
| Type of Dispersal | none | Ballooning, No ballooning | (Ballooning) Performs aerial dispersal via silk-aided wind drift (often only as a juvenile; inferred from family); (No ballooning) Not known to perform aerial dispersal or aerial dispersal rare (inferred from family) |
| Possession of foot pads | none | Yes, No | (Yes) Possesses adhesive foot pads to assist climbing; (None) No adhesive foot pads |

**References:** Lowe, E.C., Threlfall, C.G., Wilder, S.M. & Hochuli, D.F. (2018). Environmental drivers of spider community composition at multiple scales along an urban gradient. *Biodivers. Conserv.*, 27, 829–852.

#### 126) S34TTP: Vegetation across an aridity gradient – Pekin *et al.* 2011

**Data owner(s)**: Burak K. Pekin

**Description:** Plant species abundance and soil nutrients were determined at 16 forest sites with variable fire histories across an aridity gradient.

**Geographical scope:**	Walpole and Albany, SW Australia

**Taxonomical scope:** Terrestrial vascular plants

**Ecological scope:** Terrestrial

**Number of sampling units:** 16 sites

**Minimum distance between sampling units:** 0.68 kilometres

**Maximum distance between sampling units:** 22.16 kilometres

**Environmental variables:**

| Variable | Unit or factor levels |
| --- | --- |
| Soil type |  |
| Fire interval sequence |  |
| Fire frequency |  |
| Mean annual precip. | mm |
| Potential evapotrans. | mm |
| Aridity index |  |
| Total species richness |  |
| Herbaceous species richness |  |
| Woody species richness |  |
| Soil NO3- | μg g-1 |
| Soil NH4+ | μg g-1 |
| Total soil N (%) | % |
| Total soil C (%) | % |
| Soil C: N | ratio |
| Inorganic soil P | μg g-1 |
| Organic soil P | μg g-1 |
| Soil N: P | ratio |

**Functional traits:** Life cycle, Regeneration strategy, Root structure, N-fixing ability.

**References:** Pekin, B.K., Wittkuhn, R.S., Boer, M.M., Macfarlane, C. & Grierson, P.F. (2011). Plant functional traits along environmental gradients in seasonally dry and fire-prone ecosystem. *J. Veg. Sci.*, 22, 1009–1020.

#### 127) S34TAT_2: Spiders along an urbanization gradient in Sydney, Australia – Lowe *et al*. 2018

**Data owner(s)**: Elizabeth C Lowe, Jonas Wolff

**Description: S**amples were taken along a gradient of urbanisation extending 50 km north and south of the Sydney Central Business District (CBD) and around 20 km wide.

**Geographical scope:**	Urban gradient, Sydney, Australia

**Taxonomical scope:** Insects (Spider)

**Ecological scope:** Terrestrial

**Number of sampling units:** 65 sites

**Minimum distance between sampling units:** 0.01 kilometres

**Maximum distance between sampling units:** 52.80 kilometres

**Environmental variables:**

| Variable | Unit or factor levels |
| --- | --- |
| Stoires number |  |
| Adjoining backyards |  |
| Distance between the bushes | meter |
| Distance between fragments | meter |
| Total area of the property | square meter |
| Base area of house | meter |
| Area surveyed | hectare |
| Fenced gardens | quantity |
| Free standings gardens | quantity |
| Outdoor garden furniture | quantity |
| Washing lines | quantity |
| Shed pr bubby | quantity |
| Garden structures | quantity |
| Pool or ponds number | quantity |
| Pool number | quantity |
| Ponds/drains number | quantity |
| Water sources number | quantity |
| Composts number | quantity |
| Pets number in residence | quantity |
| Pet details | quantity |
| Kids number in residence | quantity |
| Trees number | quantity |
| Tree DBH | centimeter |
| Native vegetation | percentage |
| Hard surfaces | percentage |
| Grass | percentage |
| Garden beds | percentage |
| Microhabitat (0-50 cm) | percentage |
| Microhabitat (50-100 cm) | percentage |
| Microhabitat (100-200 cm) | percentage |
| Litter cover | percentage |
| Rocks/ sticks | percentage |
| Vegetation area (50 m) | percentage |
| Building area (50 m) | percentage |
| Grass area (50 m) | percentage |
| Hard surface (50 m) | percentage |
| Water area (50 m) | percentage |

**Functional traits:**

| Variable | Unit or factor levels | Description |
| --- | --- | --- |
| Guild | Web builder, Hunter | (Web builder) Builds silken snare to capture prey; (Hunter) Hunts without the aid of a web (but may actively deploy silk to immobilize prey) |
| Hunting style | Orb web, Cobweb, Reduced web, Sheet web, Lace web, Ambusher, Foliage runner, Ground runner, Kleptoparasite | (Orb web) builds an orb web to capture prey; (Cobweb) Builds a space web with knock-down- or tangle-lines, occasionally with gumfooted; (Sheet web) Build suspended or substrate bound horizontal sheets to capture prey; (Lace web) Builds cribellar lace webs to capture prey; (Reduced web) Strongly modified web, which is used in conjunction with a special hunting strategy; (Foliage runner) Non-web building hunter that actively searches and pursues prey in above-ground habitats; may use sticky silk to immobilize prey; (Ground runner) Non-web building hunter that actively searches and pursues prey on the ground; may use sticky silk to immobilize prey; (Ambusher) Non-web building hunter that waits for prey to come by; (Kleptoparasite) Steals food from webs of other spiders |
| Capture lines | Cribellar silk, Viscid silk, Non-adhesive, None | (Cribellar silk) Utilizes cribellar silk (dry nanofiber-bundles) for prey capture; (Viscid silk) Utilizes glue coated capture lines; (Non-adhesive) Adhesive threads are absent in web; (None) Does not build webs for prey capture |
| Body Size | Small, Medium, Large | (Small) Body length <5 mm; (Medium) Body length 5-10 mm; (Large) Body length > 10 mm |
| Period of activity | Durnal, Nocturnal | (Diurnal) Active during day; (Nocturnal) Active at night |
| Type of Dispersal | Ballooning, No ballooning | (Ballooning) Performs aerial dispersal via silk-aided wind drift (often only as a juvenile; inferred from family); (No ballooning) Not known to perform aerial dispersal or aerial dispersal rare (inferred from family) |
| Possession of foot pads | Yes, No | (Yes) Possesses adhesive foot pads to assist climbing; (None) No adhesive foot pads |

**References:** Lowe, E.C., Threlfall, C.G., Wilder, S.M. & Hochuli, D.F. (2018). Environmental drivers of spider community composition at multiple scales along an urban gradient. *Biodivers. Conserv.*, 27, 829–852.

#### 128) S43TBI: Bird assemblages from New Zealand – Barbaro *et al.* 2012

**Data owner(s)**: Luc Barbaro

**Description:** Bird assemblages were sampled by conducting 15 min point-counts at paired edge and interior plots in 13 forest fragments of increasing size (0.5–141 ha).

**Geographical scope:**	Fragmented native forests, volcanic banks peninsula, Canterbury, South Island, New Zealand

**Taxonomical scope:** Birds

**Ecological scope:** Terrestrial

**Number of sampling units:** 26

**Minimum distance between sampling units:** < 0.01 kilometres

**Maximum distance between sampling units:** 24.87 kilometres

**Environmental variables:**

| Variable | Unit | Unit or factor levels | Description |
| --- | --- | --- | --- |
| Plot elevation | m | Meters | Meters above sea level |
| Patch area size | ha | Hectare | Size of the habitat patch |
| Forest cover | % | Percentage | Forest percentage cover around a patch |
| Grassland cover | % | Percentage | Grassland percentage cover around a patch |
| ShannonLandscapeDiv |  |  | [Quantitative measure that shows how many different patch types there are in a landscape, and simultaneously takes into account how evenly the patches are distributed among those types](https://en.wikipedia.org/wiki/Species) |

**Functional traits:**

| Variable | Unit | Unit or factor levels |
| --- | --- | --- |
| Biogeographic origin | none | (1) endemic, (2) native, (3) exotic |
| Foraging method | none | (1) canopy gleaner, (2) understorey gleaner, (3) ground gleaner, (4) ground prober |
| Body mass | g | (1) <15 g(gram), (2) 15-29 g, (3) 30-100 g, (4) >100 g |
| Adult diet | none | (1) omnivorous, (2) insectivorous, (3) mixed diet |
| Mobility | none | (1) resident, (2) migrant or local movements |
| Nest location | none | (1) open on ground, (2) open on tree, (3) cavity |
| Clutch size | none | (1) 1-2 eggs, (2) 3 eggs, (3) 4 eggs, (4) > 5 eggs |

**References:** Barbaro, L., Brockerhoff, E.G., Giffard, B. & van Halder, I. (2012). Edge and area effects on avian assemblages and insectivory in fragmented native forests. *Landsc. Ecol.*, 27, 1451–1463.

#### 129) S06MCO: Coral reefs from Indonesia – Rachello *et al*. 2007

**Data owner(s)**: Daniel Cleary, Nicole de Voogd, Willem Renema

**Description:** The present study assessed the Pulau Seribu reef system through the study of 20 mid-to-offshore patch reefs, and seven inshore patch reefs, or what remains of them, within and just outside Jakarta Bay.

**Geographical scope:**	Coral reefs, Jakarta, Indonesia

**Taxonomical scope:** Corals

**Ecological scope:** Marine

**Number of sampling units:** 27 sites

**Minimum distance between sampling units:** 0.79 kilometres

**Maximum distance between sampling units:** 65.27 kilometres

**Environmental variables:**

| Entry | Variable | Unit | Unit or factor levels |
| --- | --- | --- | --- |
| DistJak | Distance to Jakarta | km | Kilometre |
| DistMain | Distance to mainland | km | Kilometre |
| Alg_ass | Algal assemblage cover |  |  |
| Alg_cor | Coraline algae cover |  |  |
| Alg_tur | Turf algae cover |  |  |
| Alg_hal | Halimeda cover |  |  |
| Alg_mac | Macroalgae cover |  |  |
| Deadcor | Dead coral with algae cover |  |  |
| Dea_alg | Dead coral cover |  |  |
| Rubble | Rubble cover |  |  |
| Sand | Sand cover |  |  |
| Softcor | Soft coral cover |  |  |
| Sponge | Sponge cover |  |  |
| Coral | Live coral cover |  |  |
| Temp | Temperature |  |  |
| DO | Dissolved oxygen | mg/l |  |
| pH | pH | pH |  |
| Turb | Water turbidity |  |  |
| Sal | Water salinity |  |  |
| CdSea | Cd in sea water | µg1ˉ¹ | Microgram per liter |
| CuSea | Cd in sea water | µg1ˉ¹ | Microgram per liter |
| CrSea | Cr in sea water | µg1ˉ¹ | Microgram per liter |
| NiSea | Ni in sea water | µg1ˉ¹ | Microgram per liter |
| PbSea | Pb in sea water | µg1ˉ¹ | Microgram per liter |
| ZnSea | Zn in sea water | µg1ˉ¹ | Microgram per liter |
| AsSed | As in sediment | mgkgˉ¹ or % | Milligram per kilogram or percentage |
| AlSed | Al in sediment | mgkgˉ¹ or % | Milligram per kilogram or percentage |
| CaSed | Ca in sediment | mgkgˉ¹ or % | Milligram per kilogram or percentage |
| CoSed | Co in sediment | mgkgˉ¹ or % | Milligram per kilogram or percentage |
| CrSed | Cr in sediment | mgkgˉ¹ or % | Milligram per kilogram or percentage |
| CuSed | Cu in sediment | mgkgˉ¹ or % | Milligram per kilogram or percentage |
| FeSed | Fe in sediment | mgkgˉ¹ or % | Milligram per kilogram or percentage |
| KSed | K in sediment | mgkgˉ¹ or % | Milligram per kilogram or percentage |
| MgSed | Mg in sediment | mgkgˉ¹ or % | Milligram per kilogram or percentage |
| MnSed | Mn in sediment | mgkgˉ¹ or % | Milligram per kilogram or percentage |
| MoSed | Mo in sediment | mgkgˉ¹ or % | Milligram per kilogram or percentage |
| NiSed | Ni in sediment | mgkgˉ¹ or % | Milligram per kilogram or percentage |
| PSed | P in sediment | mgkgˉ¹ or % | Milligram per kilogram or percentage |
| PbSed | Pb in sediment | mgkgˉ¹ or % | Milligram per kilogram or percentage |
| SiSed | Si in sediment | mgkgˉ¹ or % | Milligram per kilogram or percentage |
| SnSed | Sn in sediment | mgkgˉ¹ or % | Milligram per kilogram or percentage |
| SrSed | Sr in sediment | mgkgˉ¹ or % | Milligram per kilogram or percentage |
| ThSed | Th in sediment | mgkgˉ¹ or % | Milligram per kilogram or percentage |
| TiSed | Ti in sediment | mgkgˉ¹ or % | Milligram per kilogram or percentage |
| VSed | V in sediment | mgkgˉ¹ or % | Milligram per kilogram or percentage |
| ZnSed | Zn in sediment | mgkgˉ¹ or % | Milligram per kilogram or percentage |
| ZrSed | Zr in sediment | mgkgˉ¹ or % | Milligram per kilogram or percentage |

**Functional traits:**

| Code | Variable | Unit | Unit or factor levels |
| --- | --- | --- | --- |
| ColoShape | Colony shape |  | Branching, encrusting, free, laminar, massive, branching |
| ColoForm | Colony form |  | Plocoid, cerioid, meandroid, solitary, phaceloid |
| ReproStrat | Reproductive strategy |  | Broadcast-spawn, brooder, broadcast-spawn/brooder |
| CoralSize_ORDINAL | Corallite size | classe of size (mm) | (<2mm) very small, (2-5 mm) small, (5-8 mm) medium, (8-11 mm) large, (>11m) very large |
| AdaptStrat_ORDINAL | Adaptive strategy | class of tolerance | based on growth speed and subsequent fragility |
| ShapeCat | Shape category |  | Thick branching, thin branching, encrusting, free, laminar, massive, branching |
| ShapeCat-b | Shape category b |  | Branching, encrusting, free, laminar, massive, branching |
| Corallites-calice-valleysCat | Corallites-calice-valleys categories | classe of size (mm) | Very large, large, medium, small, very small |
| CoralSize_ORDINAL | Corallite size | mm | (<2mm) very small, (2-5 mm) small, (5-8 mm) medium, (8-11 mm) large, (>11m) very large |
| MOR-FORM | Colony form | none | Plocoid, cerioid, meandroid, solitary, phaceloid |
| REPRODUCTIVE MODE | Reproductive mode | none | Broadcast-spawn, brooder, broadcast-spawn/brooder |
| DistCat | Distinktive categorie |  | Small, moderate, wide, very wide |
| Shape2 | Shape |  | From 1 to 5 |
| Cor-Size2 | Size 2 |  | From 1 to 5 |
| Mod-form2 | Colony form |  | From 1 to 5 |
| Repr2 | Reproductive mode 2 |  | 1, 2 |
| DistCat2 | Distinktive categorie 2 |  | From 1 to 4 |

**References:** Rachello-Dolmen, P.G. & Cleary, D.F.R. (2007). Relating coral species traits to environmental conditions in the Jakarta Bay/Pulau Seribu reef system, Indonesia. *Estuar. Coast. Shelf Sci.*, 73, 816–826.

#### 130) N57TTP: Vegetation from Scotland – Pakeman 2011

**Data owner(s)**: Robin Pakeman

**Description:** Thirty sites were chosen within a small area on the west coast of Scotland. The area was selected to have a large range of land uses and intensities within a small area (5 3 7 km, centered on 57.318 N, 5.668 W) sharing the same geology and climate.

**Geographical scope:**	Drumbuie, Scotland

**Taxonomical scope:** Terrestrial vascular plants

**Ecological scope:** Terrestrial

**Number of sampling units:** 30 sites

**Minimum distance between sampling units:** 0.01	kilometres

**Maximum distance between sampling units:** 6.99 kilometres

**Environmental variables:**

| Code | Variable | Unit or factor levels | Description |
| --- | --- | --- | --- |
| SoilN | SoilN | %/w | Soil total nitrogen |
| SoilC | SoilC | %/w | Soil total carbon |
| SoilCN | Soil_C: N | none | soil carbon: nitrogen ratio |
| MoistureLoss | MoistureLoss |  | moisture loss at 105 Celsius degrees |
| LossOnIgnition | LossOnIgnition |  | loss of ignition at 405 Celsius degrees |
| SoilSizeFraction0.01-2 | SoilSizeFraction0.01-2 | % | soil texture, relative composition in fractions 0.01-2 micrometres |
| SoilSizeFraction2-20 | SoilSizeFraction2-20 | % | soil texture, relative composition in fractions 2-20 micrometres |
| SoilSizeFraction20-60 | SoilSizeFraction20-60 | % | soil texture, relative composition in fractions 20-60 micrometres |
| SoilSizeFraction60-2000 | SoilSizeFraction60-2000 | % | soil texture, relative composition in fractions 60-2000 micrometres |
| Field Capacity | Field Capacity |  | estimated based on the soil texture above |
| Soil type | Soil type |  |  |
| pH_H2O | pH(H2O) | none | soil pH in water |
| pH_CaCl2 | pH(CaCl2) | none | soil pH |
| ResinBagNH4-N | ResinBagNH4-N |  | soil ammonium relase |
| ResinBagNO2_3-N | ResinBagNO2/3-N |  | soil nitrates/nitrites release |
| ResinBagtotal-N | ResinBagtotal-N |  | soil total nitrogen release based on the two measures above |
| VegN | Veg_%N | % |  |
| VegC | Veg_%C | % |  |
| VegCN | Veg_C: N |  |  |
| VegCa | VegCa | g/100g |  |
| VegK | VegK | g/100g |  |
| VegMg | VegMg | g/100g |  |
| VegP | VegP | g/100g |  |
| VegNP | Veg N: P |  |  |
| SoilCa | SoilCa | mg/kg |  |
| SoilK | soilK | mg/kg |  |
| SoilMg | SoilMg | mg/kg |  |
| SoilP | SoilP | mg/kg |  |
| Township | Township | factor |  |
| Sum_graz | Sum_graz | 0 or 1 | summer grazing |
| Win_graz | Win_graz | 0 or 1 | winter grazing |
| Cropped | Cropped | 0 or 1 |  |
| Silage | Silage | 0 or 1 |  |
| Fallow | Fallow | 0 or 1 |  |
| Woodland | Woodland | 0 or 1 |  |
| height | height | metres | vegetation height |
| Disturbance_intensity | Disturbance_intensity | numeric, 0-60 | index of disturbance intensity based on several parameters: frequency/intensity of disturbance, biomass loss at disturbance, belowground disturbance and start of disturbance in weeks (see Pakeman 2011) |
| Description | Description | short text | Site description, management information |
| disturb_freq | Disturbance frequency | 0-5 | see: Kühner, A. & Kleyer, M. (2008). A parsimonious combination of functional traits predicting plant response to disturbance and soil fertility. Journal of Vegetation Science, 19, 681–692. |
| biom_loss | Biomass loss | 0-100 |  |
| below_disturb | Belowground disturbance | 0 = no belowground disturbance in previous 5 years; 0.5 = plowing 1–5 years previously; 1 = plowing in year of sampling |  |
| disturb_start | Start of disturbance | 1–53 | first disturbance event of year measured from start of growing season where week 53 equates to undisturbed |
| ANPP | Annual net primary production |  |  |
| LiveBiomass | Live biomass |  |  |
| DeadBiomass | Dead biomass |  |  |
| Standingmass | Total aboveground biomass |  |  |
| Offtake | Offtake | % |  |
| massloss | Mass loss from standard litter | % |  |
| PLFA_fungbac | Fungi-to-bacteria ratio |  |  |
| NNI | Nitrogen nutrition index |  |  |
| NPI | Phosphorus nutrition index |  |  |

**Functional traits:**

**References:** Pakeman, R.J. (2011). Multivariate identification of plant functional response and effect traits in an agricultural landscape. *Ecology*, 92, 1353–1365.

#### 131) S05MFI_3: Fishes from Indonesia – Cleary *et al*. 2016.

**Data owner(s)**: Daniel Cleary

**Description:** Fishes were visually assessed along six transects (30m long) at each of the two studied depths (3 and 5m). Individuals observed within 5 m on either side of the transect were identified to species, if possible, and recorded.

**Geographical scope:**

**Taxonomical scope:** Fishes

**Ecological scope:** Marine

**Number of sampling units:** 27 sites

**Minimum distance between sampling units:** 0.47 kilometres

**Maximum distance between sampling units:** 65.33 kilometres

**Environmental variables:**

| Variable | Unit | Unit or factor levels |
| --- | --- | --- |
| Water transparency | m | Metres |
| Temperature | °C | Degree celsius |
| Acidity | pH | pH |
| Dissolved oxygen | μg /L | Microgamrs per liter |
| Salinity | ppt or % | Parts per thousand or percentage |
| Zones | none | (Of) Offshore, (In) Inshore, (Mid) Midshore |
| Acropora_corals | % | Mean percentage cover |
| Branching_corals | % | Mean percentage cover |
| Encrusting_corals | % | Mean percentage cover |
| Foliose_corals | % | Mean percentage cover |
| Massive_corals | % | Mean percentage cover |
| Mushroom_corals | % | Mean percentage cover |
| Submassive_corals | % | Mean percentage cover |
| Algal_assemblages | % | Mean percentage cover |
| Coraline_algaes | % | Mean percentage cover |
| Dead_corals | % | Mean percentage cover |
| Halimeda_algae | % | Mean percentage cover |
| Macroalgae | % | Mean percentage cover |
| Rubble | % | Mean percentage cover |
| Sand | % | Mean percentage cover |

**Functional traits:** Traits were partly deduced from McClanahan, T. R., 2019.

| Code | Variable |
| --- | --- |
| Lmx | maximum length? |
| Linf | minimum length? |
| K | growth rate? |
| Yrs | life span? |
| Mort | natural mortality? |
| Life | life span? |
| Gen | generation time? |
| AgeM | age at first maturity? |
| Lm | length at first maturity? |
| Lopt | optimum length? |
| Res |  |
| Anim | Animal? |
| Plan | Plant? |
| Both | Both (Animal and plant)? |
| Trop | trophic level? |

**References:** Cleary, D.F.R., Polónia, A.R.M., Renema, W., Hoeksema, B.W., Rachello-Dolmen, P.G., Moolenbeek, R.G., *et al.* (2016). Variation in the composition of corals, fishes, sponges, echinoderms, ascidians, molluscs, foraminifera and macroalgae across a pronounced in-to-offshore environmental gradient in the Jakarta Bay–Thousand Islands coral reef complex. *Mar. Pollut. Bull.*, 110, 701–717.

#### 132) S05MFR: Foraminifera from Indonesia – Cleary & Renema 2007

**Data owner(s)**: Daniel Cleary

**Description:** Foraminifera species and environmental data were sampled from July to October 1997. Sample stations were selected so that a wide range of microhabitats were covered along an in-to-offshore gradient.

**Geographical scope:**	Coral reefs, Spermonde Archipelago, Makassar, southwest Sulawesi, Indonesia

**Taxonomical scope:** Foraminifera

**Ecological scope:** Marine

**Number of sampling units:** 31 sites

**Minimum distance between sampling units:** < 0.01 kilometres

**Maximum distance between sampling units:** 39.59 kilometres

**Environmental variables:**

| Variable | Unit | Unit or factor levels | Description |
| --- | --- | --- | --- |
| Maximal sea depth | m | Meter |  |
| Maximal distanace | km |  |  |
| Exposure to oceanic currents | none | Categorical intensity from 1 to 4 | Exposure to wave action and water currents |
| Sedimentation | abundance |  |  |
| Coral formation | abundance |  |  |
| Sparse coral | abundance |  |  |
| Density on the reef flat | abundance |  | The reef flat is the shoreward, flat, broadest area of the reef |
| Scattered on the reef flat | abundance |  |  |
| Human settlement | abundance |  | Number of human settlements |
| Mean stream velocity | m/sec | Meter per second |  |
| Mean salinity | g/kg | Gram per kilogram |  |
| Mean temperature | ºC | Degree celsius |  |
| Mean suspensions (nutrients) | mg/l | Milligram per liter |  |
| Standard stream velocity | m/sec | Meter per second |  |
| Standard salinity | g/kg | Gram per kilogram |  |
| Standard temperature | ºC | Degree celsius |  |
| Standard suspensions (nutrients) | mg/l | Milligram per liter |  |
| Water visibility | m | Meter |  |

**Functional traits:**

| Variable | Unit or factor levels |
| --- | --- |
| Species form | Spines, Lenticular, Flat, Spindle |
| Symbiont-bearing foraminifera | Dinoflagellates, Chloroplasts, Rhodophytes, Diatoms, Chlorophytes |
| Skeletal structure | Hyaline, Imperphorat |

**References:** Cleary, D. & Renema, W. (2007). Relating species traits of foraminifera to environmental variables in the Spermonde Archipelago, Indonesia. *Mar. Ecol. Prog. Ser.*, 334, 73–82.

### V. Datasets compiled from the Marine microplankton diversity database

**Data owner:** Sofia Sal

**DOI:** [https: //doi.org/10.1890/13-0236.1](https://doi.org/10.1890/13-0236.1)

**Description:** Data were compiled for 788 stations during different oceanographic cruises in temperate, polar and subtropical regions. Stations sampled cover a wide range of marine ecosystems, ranging from coastal to open ocean. North Atlantic Ocean samples include the Irminger Sea, Norwegian Sea, North Sea, and Iceland Basin. Samples were also collected on cruises along the South Atlantic Ocean (mainly Atlantic Meridional Transect cruises), Benguela current, Indian Ocean, and West Coast of the North Pacific Ocean. One of the experiments, ''Bergen'', took place in open-air mesocosms at the Espegrend Marine Biological Station (University of Bergen). The database included biovolume values at species-level. However, other traits e.g., presence of flagella or silica were drawn from the literature, and included for the distance decay 2.0. We selected data from six different areas, particularly:

#### 133) N18MPP: Phytoplankton from the SONNE cruise

**Number of sampling units:** 18 sites

**Minimum distance between sampling units:** 1.88 kilometres

**Maximum distance between sampling units:** 586.22 kilometres

**Environmental variables:** Temperature, Surface photosynthetic active radiation, Kd490, Phosphate,	mixed layer depth, Silicate, and day length

#### 134) N54MPP: Phytoplankton from the North Sea

**Number of sampling units:** 44 sites

**Minimum distance between sampling units:** 0.13 kilometres

**Maximum distance between sampling units:** 397.64 kilometres

**Environmental variables:** Temperature, Surface photosynthetic active radiation, Kd490, Phosphate,	mixed layer depth.

#### 135) N59MPP: Phytoplankton from the PRIME cruise

**Number of sampling units:** 50 sites

**Minimum distance between sampling units:** < 0.01 kilometres

**Maximum distance between sampling units:** 37.17 kilometres

**Environmental variables:** Temperature, Surface photosynthetic active radiation, Kd490, Phosphate,	mixed layer depth.

#### 136) N63MPP: Phytoplankton from the FISHES cruise

**Number of sampling units:** 25 sites

**Minimum distance between sampling units:** < 0.01 kilometres

**Maximum distance between sampling units:** 895.36 kilometres

**Environmental variables:** Temperature, surface photosynthetic active radiation, Kd490, phosphate, nitrate, silicate, mixed layer depth.

#### 137) S22MPP: Phytoplankton from the Benguela current

**Number of sampling units:** 54 sites

**Minimum distance between sampling units:** < 0.01 kilometres

**Maximum distance between sampling units:** 773.00 kilometres

**Environmental variables:** Temperature, surface photosynthetic active radiation, Kd490, phosphate, nitrate, silicate, mixed layer depth.

#### 138) S05MPP: Phyoplankton from the Atlantic Meridional Transect

**Number of sampling units:** 25 sites

**Minimum distance between sampling units:** 326.62 kilometres

**Maximum distance between sampling units:** 11635.00 kilometres

**Environmental variables:** Temperature, surface photosynthetic active radiation, Kd490, PARz, phosphate, nitrite, nitrate, silicate, mixed layer depth.

**References:** Sal, S., López-Urrutia, Á., Irigoien, X., Harbour, D.S. & Harris, R.P. (2013). Marine microplankton diversity database. *Ecology*, 94, 1658.
