## Supplemental information 4 for "Distance decay 2.0 – a global synthesis of taxonomic and functional turnover in ecological communities"

*Appendix S4*

**Results of occurrence-based replacement and richness differences components**

Organismal variables and dataset features

There results of replacement and richness differences were similar for the organismal variables and dataset features (Fig. S3-S4). Actively dispersed taxa had the flattest slopes of replacement and richness differences along spatial and environmental distances (Fig. S3-S4). Larger-bodied organisms had the flattest slopes along space and the steepest slope along environment (Fig. S3-S4). In contrast, body size had very small effect in the decay of functional similarities (Fig. S3-S4). Datasets with smaller number of species had the steepest slopes along the space and the flattest slope along the environment (Fig. S3-S4). Higher functional γ-diversity resulted in flatter slopes along the space while a U-shaped relationship was observed along environmental distances (Fig. S3-S4). Datasets with smaller number of study sites had the steepest slopes while the datasets with smaller number of environmental variables had the flattest slopes (Fig. S3-S4).

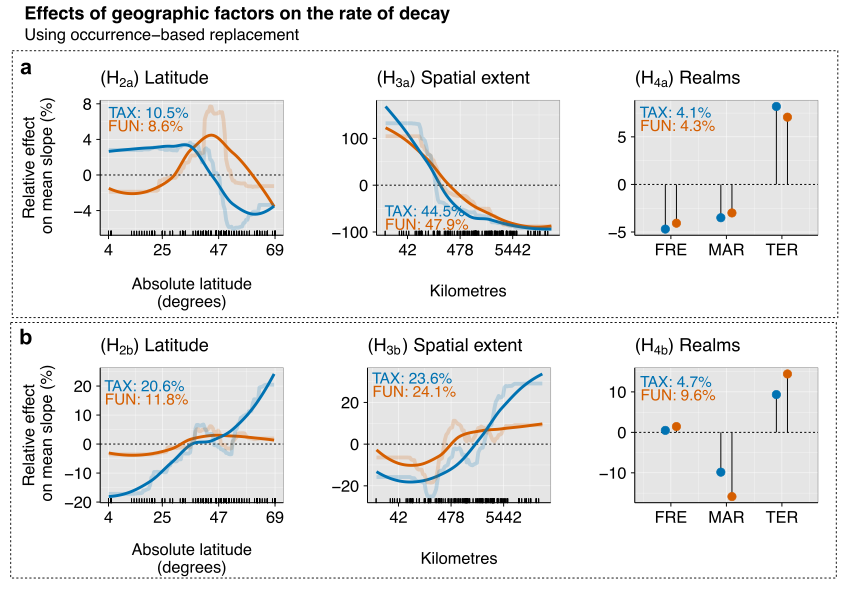

Figure S1. Relative effects (%) of geographic factors on the rate of decay along spatial (a) and environmental (b) distance decay of the replacement component of taxonomic (TAX - blue) and functional (FUN - orange) similarities using occurrence data across datasets. Partial dependence plots show the effects of a predictor variable on the response variable after accounting for the average effects of all other variables in the model. Semi-transparent lines represent the actual predicted effects; solid lines represent LOESS fits to predicted values from BRT. We show here only the variables related to the specific hypotheses, i.e., latitude, spatial extent, and realms (FRE = Freshwater, TER = Terrestrial, MAR = Marine).

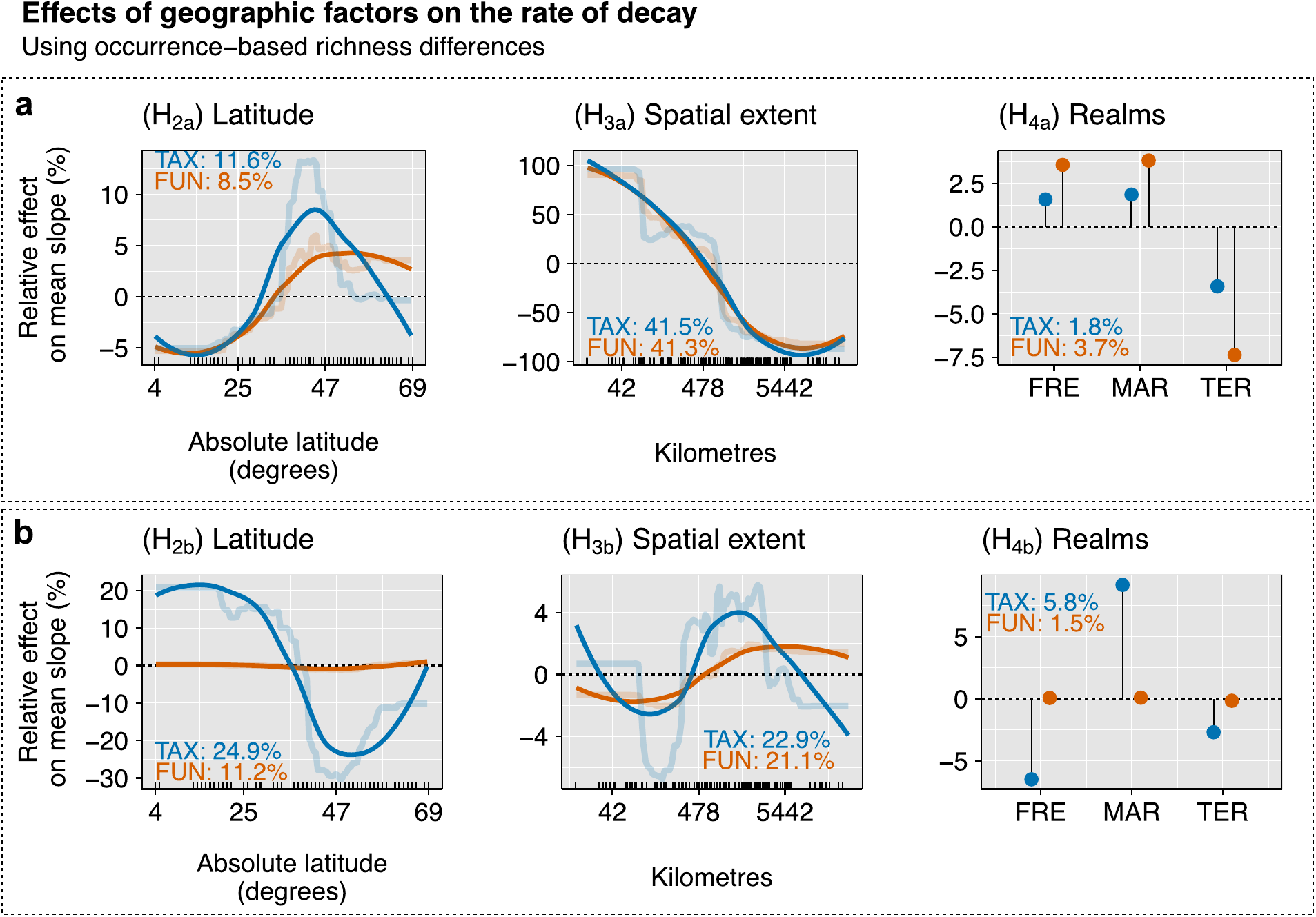

Figure S2. Relative effects (%) of geographic factors on the rate of decay along spatial (a) and environmental (b) distance decay of the richness differences component of taxonomic (TAX - blue) and functional (FUN - orange) similarities using occurrence data across datasets. Partial dependence plots show the effects of a predictor variable on the response variable after accounting for the average effects of all other variables in the model. Semi-transparent lines represent the actual predicted effects; solid lines represent LOESS fits to predicted values from BRT. We show here only the variables related to the specific hypotheses, i.e., latitude, spatial extent, and realms (FRE = Freshwater, TER = Terrestrial, MAR = Marine).

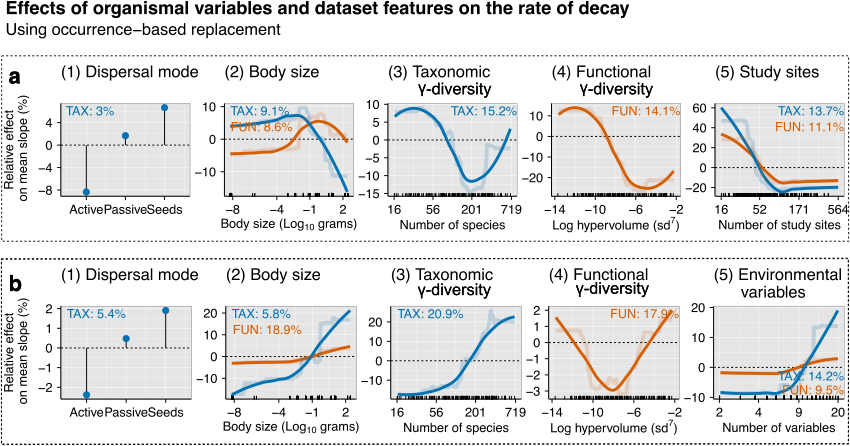

Figure S3. Relative effects (%) of organismal variables and dataset features on the rate of decay along spatial (a) and environmental (b) distance considering the replacement component of taxonomic (blue lines) and functional (orange lines) similarities using occurrence data across datasets. Partial dependence plots show the effects of a predictor variable on the response variable after accounting for the average effects of all other variables in the model. Semi-transparent lines represent the actual predicted effects; solid lines represent LOESS fits to predicted values from BRT. We show here the organismal variables and the variables related to the dataset features.

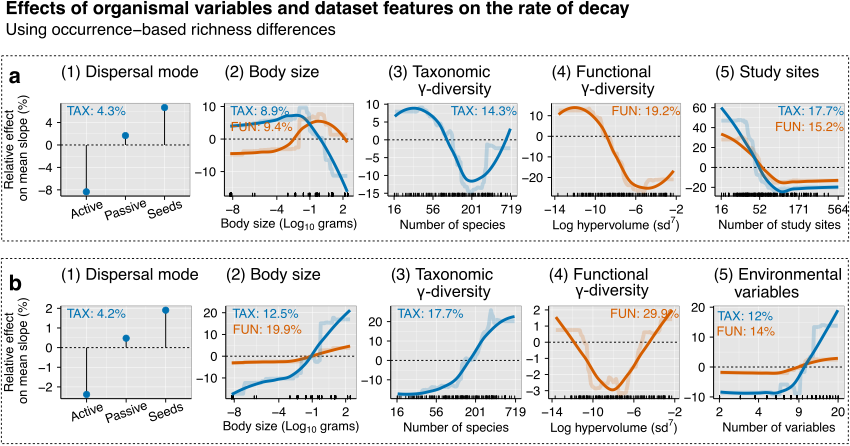

Figure S4. Relative effects (%) of organismal variables and dataset features on the rate of decay along spatial (a) and environmental (b) distance considering the richness differences component of taxonomic (blue lines) and functional (orange lines) similarities using occurrence data across datasets. Partial dependence plots show the effects of a predictor variable on the response variable after accounting for the average effects of all other variables in the model. Semi-transparent lines represent the actual predicted effects; solid lines represent LOESS fits to predicted values from BRT. We show here the organismal variables and the variables related to the dataset features.
