## Supplemental information 3 for "Distance decay 2.0 – a global synthesis of taxonomic and functional turnover in ecological communities"

*Appendix S3*

**Results using abundance-based total similarities**

*Strength of the distance decay*

The distance decay of similarity had an overall exponential shape, but a few datasets showed a linear decay (Fig. S1). The distance decay of taxonomic similarity was stronger than the distance decay of functional similarity along both spatial distances (t = 6.917, df = 142, p < 0.001; Fig. S1a) and environmental distances (t = 6.956, df = 139, p < 0.001; Fig S1b). Considering abundance-based similarities, 26 datasets had stronger distance decay of functional similarities than taxonomic similarities along environmental distance.

*Rate of the distance decay*

The mean slope of the spatial distance decay was 0.018 (sd ± 0.086) for taxonomic similarities, and 0.009 (sd ± 0.054) for functional similarities. For environmental distances, the mean slope of the distance decay was 1.455 (sd ± 1.220) for taxonomic similarities and 0.522 (sd ± 0.404) for functional similarities. Regarding the biotic groups, aquatic plants had the steepest slopes along spatial distance both for taxonomic and functional similarities (Fig. S2a). Along environmental distance, corals had the steepest slopes (Fig. S2b).

Across datasets, BRT explained 40.81% of the deviance of the slopes of the spatial distance decay for taxonomic similarities, and 41.51% for functional similarities using occurrence data. For the distance decay along environmental gradients, BRT explained 13.41% of the deviance of the slopes of the decay of taxonomic similarities and 8.27% for functional similarities. Spatial extent contributed most to the variation in slopes considering spatial distance or environmental distance while γ-diversity (taxonomic and functional) and latitude alternate as the second most important variable (Fig. S3 – S4).

Latitudinal patterns

The slopes of spatial distance decay of both taxonomic and functional similarities were the steepest in datasets centred at ca. 35º-45º and decreased sharply for datasets at higher latitudes (Fig S3a). The slopes of environmental distance decay were the flattest in datasets centred at ca. 50º, above which it got steeper (Fig. S3b).

Spatial extent

The distance decay of taxonomic and functional similarities was flatter in the datasets that covered larger spatial extent (Fig. S3a). For environmental distances, distance decay was steeper in the datasets that covered larger spatial extents for taxonomic similarities but had little effect on the slopes of functional similarities (Fig S3b).

Realms

Marine ecosystems had flatter slopes compared to freshwater or terrestrial ecosystems considering spatial and environmental distances, but only for taxonomic similarities. The importance of realm in BRTs was overall low, however (Fig S3).

Organismal variables and dataset features

The slopes of distance decay were the flattest for actively dispersed organisms for both spatial and environmental gradients (Fig S4). Large-bodied organisms had steeper decay of taxonomic similarity along spatial distance, whereas small-bodied organisms had steeper decay of functional similarity (Fig S4). Taxonomic γ-diversity had a U-shaped relationship with slopes for distance decay along spatial and environmental gradients (Fig. S4). Datasets with higher functional γ-diversity showed flatter slopes along spatial gradients (Fig S4a), but steeper slopes along environmental gradient (Fig S4b). Generally, slopes were steeper in the datasets where the number of sampling sites was higher (Fig. S4a), and flatter when datasets comprised only a few environmental variables (Fig S4b).

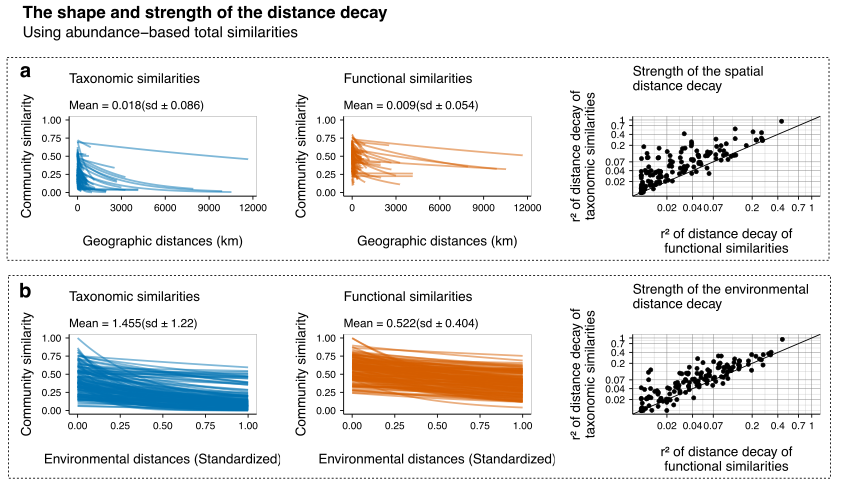

Figure S1. The distance decay along (a) spatial distance, and (b) environmental distance using abundance data. Each line in the panels of left and middle columns shows the shape of the distance decay of an individual dataset. The mean and standard deviation of slopes are given in the plots. The blue lines show the distance decay of taxonomic similarity while the orange line shows the distance decay of functional similarity. The panels on the right column show the strength of the distance decay of taxonomic (y-axis) and functional (x-axis) similarity. The 1:1 line marks the equivalence of r² between taxonomic and functional similarities. The dots below the line indicate a dataset with stronger decay of functional than taxonomic similarity, whereas circles above the line indicates stronger decay of taxonomic than functional similarities.

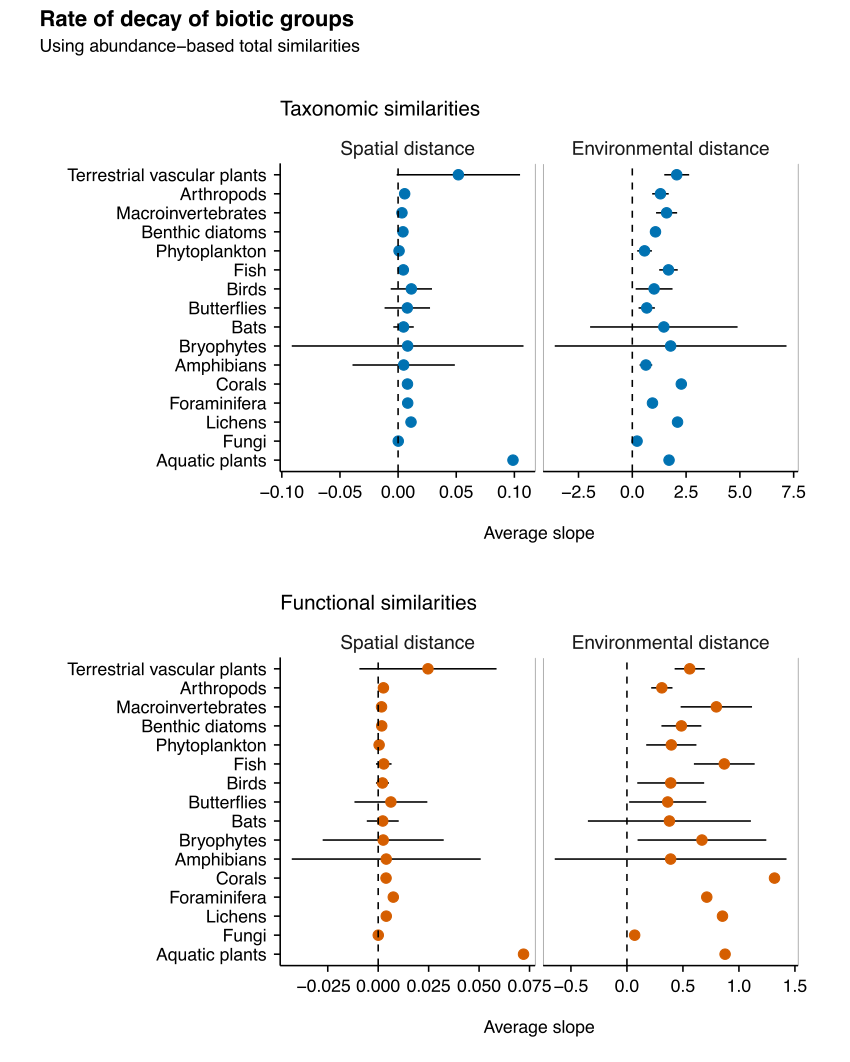

Figure S2. The average rate of decay of biotic groups using abundance data along spatial and environmental distances. The vertical dotted lines highlight the zero rate (absence of decay) and the horizontal lines indicate the 95% confidence interval of the mean. The blue circles show the rate of decay of taxonomic similarities while the orange circles show the rate of decay of functional similarities.

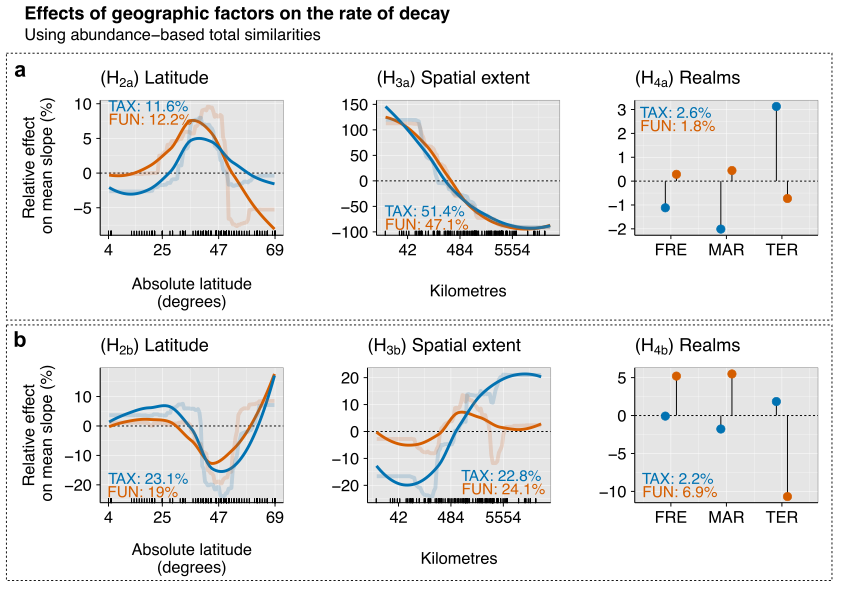

Figure S3. Relative effects (%) of the predictors on the mean slopes of spatial (a) and environmental (b) distance decay of the total component of taxonomic (TAX - blue) and functional (FUN - orange) similarities using abundance data across datasets. Partial dependence plots show the effects of a predictor variable on the response variable after accounting for the average effects of all other variables in the model. Semi-transparent lines represent the actual predicted effects; solid lines represent LOESS fits to predicted values from BRT. We show here only the variables related to the specific hypotheses, i.e., latitude, spatial extent, and realms (FRE= Freshwaters, MAR = Marine, TER = Terrestrial).

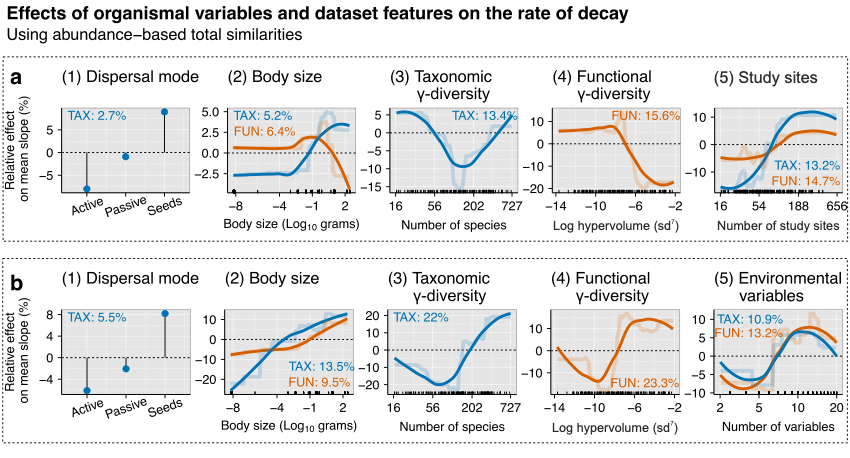

Figure S4. Relative effects (%) of the predictors on the rate of decay along spatial (a) and environmental (b) distance considering the total component of taxonomic (blue lines) and functional (orange lines) similarities using abundance data across datasets. Partial dependence plots show the effects of a predictor variable on the response variable after accounting for the average effects of all other variables in the model. Semi-transparent lines represent the actual predicted effects; solid lines represent LOESS fits to predicted values from BRT. We show here the organismal variables and the variables related to the dataset features.
