## Supplemental Information 3 for "Distance decay 2.0 – a global synthesis of taxonomic and functional turnover in ecological communities"

*Appendix S2*

**The partition of taxonomic and functional similarities based on occurrence and abundance data through dendrogram comparisons following Cardoso *et al.* (2014).**

Traditionally, β-diversity has been measured by comparing the matching/mismatching components of pairs of communities^1^. For an occurrence-based β-diversity, these components are the number of features, e.g. species, (a) shared by both communities, (b) present in the first community only, (c) present in the second community only and (d) absent in both communities but present elsewhere in the region. For an abundance-based β-diversity, we use the upper-case letter (A) to designate the sum of the abundances of the multiple features; (B) to represent the sum of abundances at the first community minus A; and (C) as the sum of abundances at the second community minus A. Therefore, the upper and lower-case components are analogous to each other and represent both the intersections and community-specific features of the studied sites. β-diversity can thus be calculated and partitioned into total, replacement and richness differences components by applying some dissimilarity index (see ref.^2^ for a review of indices) such as Sørensen index^3^ on occurrence data or equivalently the percentage differences index^4^ on abundance data as shown in Table S1.

**Table S1** Equations used for the estimation of total similarities and replacement and richness differences components using occurrence and abundance information.

|  | Ocurrence data | Abundance data |
| --- | --- | --- |
| Total | $1-(b+c)/(2a+b+c)$ | $1-(B+C)/(2A+B+C)$ |
| Replacement | $1-(2 \times min(b,c))/(2a+b+c)$ | $1-(2 \times min(B,C))/(2A+B+C)$ |
| Richness differences | $1-(\vert b-c\vert)/(2a+b+c)$ | $1-(\vert B-C\vert)/(2A+B+C)$ |

**References**

1.         Legendre, P. Interpreting the replacement and richness difference components of beta diversity. *Global Ecology and Biogeography* **23**, 1324–1334 (2014).

2.         Baselga, A. & Leprieur, F. Comparing methods to separate components of beta diversity. *Methods in Ecology and Evolution* **6**, 1069–1079 (2015).

3.         Sørensen, T. A. A method of establishing groups of equal amplitude in plant sociology based on similarity of species content, and its application to analyses of the vegetation on Danish commons. *Kongelige Danske Videnskabernes Selskabs Biologiske Skrifter* **5**, (1948).

4.         Odum, E. P. Bird Populations of the Highlands (North Carolina) Plateau in Relation to Plant Succession and Avian Invasion. *Ecology* **31**, (1950).
